# Pressure–cooling remodeling of TMV coat protein reveals mechanically partitioned capsid dynamics and selective epitope masking

**DOI:** 10.64898/2026.05.31.729108

**Authors:** Carlos Francisco Sampaio Bonafé, Juan Philippe Teixeira, Caio Cesar de Melo Freire, Miklos Maximiliano Bajay, Daniel Ferreira de Lima Neto

## Abstract

High hydrostatic pressure (HHP) perturbs protein assemblies by shifting conformational equilibria toward lower-volume states and by reorganizing hydration at cavities, interfaces, and solvent-exposed surfaces (Heremans 1982; Akasaka 2006; Roche et al. 2012; Hata, Nishiyama, and Kitao 2020). Here, we integrate pressure-dependent molecular dynamics descriptors, pressure–temperature interpretation, structure-based epitope prediction, and face-resolved intersubunit metrics to examine how pressure and pressure-cooling treatment remodel the tobacco mosaic virus coat protein (TMVcp) assembly. The pressure response is not adequately explained as uniform shrinkage. Instead, the data support a hierarchical transition from a broad, native-like conformational ensemble at low pressure, through a cooperative compacting regime around 1000– 1750 bar, toward a high-pressure compact state with reduced configurational diversity, suppressed global mobility, and localized residual fragility. A representative TMVcp face composed of A2, A3, A4, A19, A20, A21, A35, A36, and A37 behaves as a mechanically partitioned network: A3 behaves as a principal deformation hub, A20–A21–A35–A37 forms a lateral/diagonal compression corridor, A21 behaves as a bridge node, A36 acts as an anisotropic relay, and A2, A4, and A19 behave as stabilizing or adaptive anchors. Pairwise minimum-distance profiles, per-subunit radius of gyration, and post-fit RMSD converge around a late trajectory interval near 358–365 ns, suggesting a coordinated face-level breathing event rather than independent stochastic noise. These local dynamics provide a conservative structural explanation for predicted pressure-dependent epitope remodeling: HHP may mask canonical solvent-exposed epitopes by reducing loop mobility and closing intersubunit grooves, whereas pressure followed by low-temperature trapping may selectively preserve only protrusions compatible with the compact, hydration-trapped lattice. Because DiscoTope and ElliPro are computational predictors, these results should be interpreted as structural hypotheses requiring experimental validation by antibody binding assays, mutagenesis, HDX-MS, or high-pressure structural approaches.

## Introduction

High hydrostatic pressure is a powerful perturbation for studying protein stability because pressure couples directly to volume changes in conformational equilibria. In general terms, pressure shifts molecular ensembles toward states with lower partial molar volume, which can arise through cavity collapse, water penetration, altered hydration, disruption of packing defects, or reorganization of interfaces (Heremans 1982; Akasaka 2006; Meersman et al. 2013; Roche et al. 2012; Hata, Nishiyama, and Kitao 2020). As a result, pressure-induced denaturation is not equivalent to thermal denaturation. Heat often amplifies disorder and configurational expansion, whereas pressure can promote compact but non-native states, hydrated intermediates, suppressed low-frequency modes, and locally stiffened conformations (Winter and Jeworrek 2009; Roche et al. 2012; Hata, Nishiyama, and Kitao 2020).

Viral particles and virus-like assemblies are especially sensitive to this type of perturbation because their stability depends on repeated subunit interfaces, solvent-exposed antigenic surfaces, internal cavities, and cooperative mechanical coupling. HHP has been investigated as an inactivation strategy capable of altering viral infectivity while preserving relevant structural or antigenic features in some systems (Dumard et al. 2013). This makes pressure a useful experimental and computational probe for distinguishing global structural collapse from selective remodeling of surface accessibility.

TMVcp is a useful model for this problem because the tobacco mosaic virus particle is a highly ordered helical assembly. The virion contains thousands of coat protein subunits arranged in a rod-like particle with helical symmetry, and classical structural studies established the repeated subunit architecture that underlies TMV assembly and stability (Namba and Stubbs 1986; Klug 1999; Creager et al. 1999). In this manuscript, we focus on the protein assembly and its pressure-dependent structural mechanics. We intentionally avoid mechanistic claims about RNA interactions, because the present revision is centered on TMVcp protein dynamics, interface remodeling, and predicted epitope accessibility.

A purely global analysis of RMSD, radius of gyration (R_g_), RMSF, SASA, PCA, and clustering can identify compacting or rigidifying trends, but it does not fully reveal how local interfaces are remodeled. Similarly, structure-based epitope predictors such as DiscoTope and ElliPro provide useful hypotheses about residue accessibility, spatial epitope context, and protrusion, but they are not direct measurements of antibody recognition (Haste Andersen, Nielsen, and Lund 2006; Ponomarenko et al. 2008; Kringelum et al. 2012; Clifford et al. 2024). For that reason, the present revision connects global pressure-dependent metrics with local face-resolved dynamics and explicitly separates supported structural interpretations from hypotheses requiring experimental validation.

## Results

### Pressure drives a three-stage remodeling of the TMVcp conformational ensemble

Across the pressure series, the combined PCA, R_g_, RMSD, RMSF, SASA, and clustering results support a three-stage pressure response. At low pressure, approximately 1–750 bar, the conformational ensemble remains broad and dynamic. PCA indicates wider dispersion in the first two principal components, consistent with a larger accessible conformational space and preservation of collective modes. In this regime, the system still samples multiple conformational substates, and the structural descriptors are best interpreted as native-like fluctuations rather than pressure-induced collapse.

Between approximately 1000 and 1750 bar, the descriptors converge toward a cooperative transition regime. The PCA dispersion contracts, R_g_ decreases sharply, SASA is reduced, and the number of conformational clusters declines. RMSD increases relative to the low-pressure reference, but then tends toward a plateau, suggesting that the system is not simply unfolding without constraint. Instead, it appears to enter a compact non-native or pressure-remodeled basin. This interpretation is consistent with the broader high-pressure protein literature, where pressure favors conformations with smaller volume and can stabilize hydrated compact intermediates rather than producing purely expanded denatured states (Akasaka 2006; Roche et al. 2012; Hata, Nishiyama, and Kitao 2020).

Above approximately 1750 bar, the ensemble becomes more restricted. The clustering pattern is dominated by fewer substates, global RMSF is reduced, and the PCA point cloud contracts further. This supports a high-pressure compact state with reduced configurational entropy. Importantly, this state should not be described as fully native or simply denatured. The most conservative interpretation is that HHP selects a compact, mechanically constrained ensemble in which many large-amplitude motions are suppressed, while selected local regions retain residual flexibility or become local stress-release points.

### PCA reveals progressive suppression of collective motions

The PCA results support progressive restriction of collective motions with increasing pressure. At low pressure, the trajectory occupies a broad region of principal-component space, consistent with multiple low-frequency motions and higher configurational freedom. As pressure increases through the 1000–1500 bar interval, this dispersion contracts markedly, indicating that the largest-amplitude motions are being suppressed. At higher pressure, the conformations concentrate into a smaller region of conformational space, consistent with a compacted and mechanically constrained ensemble.

This PCA behavior is important because it links global motion to local epitope accessibility. Surface-exposed epitopes and intersubunit grooves depend not only on static exposure but also on local breathing and collective rearrangements. A pressure-induced reduction of low-frequency modes is therefore expected to reduce the probability that flexible loops and grooves sample antibody-accessible or protrusive conformations. This supports the interpretation that HHP can mask predicted epitopes by dynamic restriction, even when the protein assembly remains globally intact.

### R_g_ and SASA indicate cooperative compaction and surface burial

The R_g_ trend indicates pressure-dependent compaction. Based on the summarized pressure series, R_g_ decreases from a low-pressure value near 2.8 nm toward approximately 2.2 nm at the highest pressure, with the steepest change around 1000–1500 bar. This pressure window also coincides with reduced SASA, supporting a coupled process in which the assembly becomes more compact while solvent-exposed area is reduced. Because SASA reduction is not expected to be uniform across residue types or structural regions, the most plausible interpretation is selective burial of hydrophobic or groove-associated surfaces, combined with localized hydration of polar or structurally stressed regions.

The R_g_–SASA relationship provides one of the strongest bridges between global compaction and epitope masking. A reduction in R_g_ suggests a smaller spatial envelope for the modeled unit, whereas reduced SASA implies lower solvent accessibility of surface features. For predicted B-cell epitopes, this matters directly: DiscoTope is sensitive to residue solvent accessibility and spatial context, while ElliPro is sensitive to protrusion geometry (Haste Andersen, Nielsen, and Lund 2006; Ponomarenko et al. 2008; Kringelum et al. 2012). Therefore, compaction and SASA loss provide a mechanistic basis for reduced predicted epitope accessibility under pressure.

### RMSD separates departure from the initial structure from high-pressure stabilization

RMSD increases from the low-pressure reference as pressure rises, consistent with departure from the initial or native-like conformational basin. In the summarized data, RMSD rises from approximately 0.15 nm at low pressure toward approximately 0.35–0.40 nm through the intermediate pressure regime, followed by a modest plateau or slight reduction at the highest pressures. This pattern argues against a simple monotonic unfolding process. Instead, it suggests an initial structural displacement followed by stabilization of a compact pressure-selected state.

This distinction matters for interpretation. A high-pressure structure can be more compact and less flexible while still being non-native. Thus, a plateau in RMSD does not mean that the native conformation is recovered; it means that the trajectory has reached a relatively stable region of conformational space under pressure. In the context of epitope prediction, such a state may preserve global assembly while changing the geometry, protrusion, and accessibility of surface determinants.

### RMSF indicates global rigidification with localized fragility

RMSF provides the local flexibility layer needed to interpret the pressure response. The summarized data indicate a strong reduction in mean RMSF between low and high pressure, consistent with pressure-induced stiffening. Loops and termini show especially strong suppression of mobility, which would reduce the capacity for induced-fit-like conformational adjustment at surface sites. However, the response is not purely rigidifying.

Local increases in RMSF were reported around selected regions, including a site near residue 95 and a region near residues 120–125. These local increases should be interpreted cautiously as possible stress-release or hydration-sensitive regions, not as definitive unfolding nuclei without experimental confirmation.

This global-rigid/local-fragile pattern is central to the model. Pressure restricts the overall conformational ensemble, but mechanical strain is not distributed evenly. Some local regions may transiently increase mobility because they absorb stress produced by compaction of neighboring interfaces. These local mobility peaks can explain why a compact pressure-selected state may still exhibit selective surface rearrangements and altered epitope maps.

### Clustering supports conformational selection under pressure

The clustering results support a pressure-dependent loss of conformational diversity. At low pressure, multiple clusters are sampled, consistent with a flexible native-like ensemble. In the intermediate regime, fewer clusters remain, indicating the onset of conformational selection. At high pressure, a dominant cluster emerges, consistent with a compact state occupying a restricted region of conformational space.

This clustering behavior complements the PCA results. PCA shows contraction of global motion, whereas clustering shows that the trajectory increasingly occupies fewer structural substates. Together, these metrics support reduced configurational entropy under pressure. Formally, if one approximates configurational entropy from the number of accessible clusters as Δ*S* ≈ *K*_*B*_ ln *N*_clusters_, then a reduction from multiple clusters to one dominant cluster implies a substantial entropy loss. This expression is not used here as a rigorous thermodynamic calculation, but as a qualitative way to describe the collapse of accessible conformational states.

### Integrated correlation model across PCA, R_g_, RMSD, RMSF, SASA, and clustering

The multidimensional interpretation suggests a causal sequence rather than six independent descriptors. Increasing pressure first suppresses collective motions, visible as reduced PCA dispersion. This restriction is accompanied by compaction, visible as reduced R_g_, and surface burial, visible as reduced SASA. As the structure departs from the low-pressure reference, RMSD increases; as the pressure-selected basin stabilizes, RMSD reaches a plateau. RMSF decreases globally, indicating reduced local mobility, but selected residues or regions retain elevated flexibility, likely reflecting strain release or forced hydration. Clustering then captures the final consequence: fewer conformational substates remain accessible under pressure.

This model can be described as pressure-induced conformational filtering. Low pressure permits a wider ensemble of structurally related substates. Intermediate pressure filters this ensemble by favoring compact lower-volume conformations. High pressure stabilizes a restricted compact ensemble with reduced large-scale motion and altered surface accessibility. In this framework, epitope masking is not only a static SASA phenomenon but an ensemble effect: fewer conformations are available in which protrusive or flexible antigenic features can be sampled.

### Face-resolved intersubunit dynamics reveal local mechanical partitioning

To determine whether the pressure response is spatially uniform or propagated through defined local interfaces, we analyzed one representative TMVcp face composed of A2, A3, A4, A19, A20, A21, A35, A36, and A37. For each selected subunit pair, minimum-distance profiles were evaluated across the trajectory. For each individual subunit, R_g_ and post-fit RMSD were examined to separate positional rearrangement from internal deformation.

The face is not mechanically homogeneous. A2 maintains a narrow R_g_ distribution and relatively constrained RMSD behavior despite repeated intersubunit distance excursions. This supports a role as a compact positional transition node: A2 changes its relationship to neighboring subunits without strong evidence of large internal deformation. A3 shows the opposite pattern. It combines frequent deep minimum-distance events with high R_g_ variability and a two-state-like RMSD profile, identifying it as the principal deformation hub of the analyzed face.

A4 and A19 behave as stabilizing or adaptive anchors. A4 remains compact for most of the trajectory, with rare high-amplitude RMSD or R_g_ excursions, suggesting that it mainly preserves local continuity but can participate in abrupt rearrangement events. A19 preserves several interfaces but shows moderate R_g_ and RMSD increases during the main rearrangement window, indicating strain accommodation rather than strict rigidity.

The A20–A21–A35–A37 region forms a lateral/diagonal compression corridor. A20 behaves as an active lateral compression axis, A21 connects this axis to the upper/diagonal block as a bridge node, and A35– A37 form a second plastic hotspot. A36 displays particularly high anisotropy in R_g_ and RMSD behavior, consistent with a relay-like role in transmitting deformation across layers of the face. Importantly, multiple deep distance collapses, R_g_ expansions, and RMSD shifts coincide around approximately 358–365 ns. This temporal convergence supports the interpretation of a coordinated face-level breathing event, rather than a set of independent stochastic fluctuations.

This result adds a local mechanical layer to the global MD observables. Global compaction and SASA reduction can now be interpreted as emerging from pair-specific closure of selected intersubunit grooves while stabilizing anchors preserve the overall helical scaffold. The analysis remains limited to one representative face and should not be overgeneralized to all subunit positions without capsid-wide replication.

### Pressure–temperature regime interpretation

The qualitative pressure–temperature map in SM5 provides a conceptual framework for interpreting the two-stage treatment. At 300 K, increasing pressure is expected to shift the ensemble from a flexible native-like region toward a compact pressure-remodeled region. The subsequent low-temperature stage at high pressure is interpreted not as a return to the native state, but as trapping of a compact ensemble in which hydration and conformational exchange are reduced. This is compatible with the broader concept that pressure and temperature perturb different thermodynamic and kinetic components of protein stability (Winter and Jeworrek 2009; Hata, Nishiyama, and Kitao 2020).

The key consequence for antigenicity is that HHP low temperature should not be described as simple epitope restoration. Instead, it should be described as selective retention or re-emergence of a subset of geometrically compatible protrusions within a compact constrained lattice. This distinction is essential because epitope prediction tools estimate structural accessibility or protrusion; they do not directly measure antibody binding or protective immunogenicity.

### Structure-based epitope prediction and pressure-dependent masking

DiscoTope and ElliPro were used as complementary structure-based predictors. DiscoTope integrates residue propensity, solvent accessibility, spatial distribution, and contact context to estimate discontinuous B-cell epitopes (Haste Andersen, Nielsen, and Lund 2006; Kringelum et al. 2012). ElliPro identifies protruding surface patches using a protrusion index based on ellipsoid fitting (Ponomarenko et al. 2008). These predictors are useful for generating structural hypotheses, but their limited accuracy and dependence on input conformation require cautious interpretation (Clifford et al. 2024).

Within this framework, the HHP state is best interpreted as a condition that reduces accessibility of canonical surface-exposed epitope-bearing regions by closing intersubunit grooves and suppressing local flexibility. Reduced RMSF and SASA support this interpretation, while predicted changes in DiscoTope and ElliPro scores provide structural hypotheses about which regions lose exposure or protrusion under pressure. The face-resolved dynamics suggest a plausible local mechanism: deformation hubs such as A3 and the A20– A21–A35–A37 corridor can close solvent-exposed grooves, while anchors such as A2, A4, and A19 preserve scaffold continuity.

The HHP low-temperature condition should be described as partial and selective re-exposure, not full recovery of the native antigenic landscape. Cooling at high pressure may stabilize a compact hydration-trapped ensemble in which only a subset of protrusive epitopes remains geometrically compatible with the constrained lattice. This is consistent with the idea that pressure-treated particles may preserve immunologically relevant structural features while undergoing substantial conformational remodeling (Dumard et al. 2013). However, antibody binding experiments are required to confirm whether predicted epitope exposure translates into altered antigenicity.

## Discussion

### A hierarchical model of TMVcp pressure remodeling

The results synthesis supports a hierarchical model for pressure remodeling of TMVcp. The first level is global ensemble restriction: pressure reduces the accessible conformational space, suppresses low-frequency collective motions, and reduces the number of structural clusters. The second level is compaction: R_g_ and SASA trends indicate a pressure-driven shift toward lower-volume, less solvent-exposed states. The third level is local mechanical partitioning: specific intersubunit faces and subunits absorb strain differently, producing deformation hubs, bridge nodes, compression corridors, relays, and stabilizing anchors. The fourth level is antigenic consequence: groove closure and loop rigidification reduce the probability that predicted epitope-bearing regions remain accessible or protrusive.

This model avoids two oversimplifications. First, it avoids treating pressure as a uniform capsid-crushing force. The face-resolved data show that pressure response is spatially heterogeneous. Second, it avoids treating epitope changes as purely static. Predicted epitope remodeling emerges from both altered surface exposure and altered conformational sampling.

### From pressure thermodynamics to local interface closure

The thermodynamic foundation is that pressure favors lower-volume states. In proteins, this can involve water penetration into cavities, altered electrostatics, cavity collapse, and modification of packing defects (Akasaka 2006; Roche et al. 2012; Hata, Nishiyama, and Kitao 2020). In a repeated protein assembly such as TMVcp, these effects are expected to be expressed at intersubunit grooves and mechanically coupled interfaces. The present data are consistent with this expectation. Global compaction is strongest where R_g_ and SASA decline together, but local instability is strongest where RMSF, per-subunit R_g_, and post-fit RMSD identify mechanically responsive regions.

The late 358–365 ns window deserves particular attention because multiple local observables converge there. Distance collapse, R_g_ shifts, and RMSD transitions occurring in the same interval support a coordinated local breathing event. The term “breathing” is used descriptively: it means a transient, coupled opening/closing or rearrangement of interfaces, not a formally established thermodynamic phase transition.

### Implications for epitope masking and cold trapping

The epitope model is conservative but mechanistically coherent. Under HHP, surface loops and grooves become less mobile and less solvent-exposed. This reduces both the static accessibility and the dynamic sampling of predicted B-cell epitopes. Under pressure-cooling, the system may not return to the native-like ensemble. Instead, cooling may kinetically trap compact substates that preserve only selected protrusive features. Therefore, the predicted antigenic landscape after pressure-cooling is expected to be filtered: some epitopes may remain masked, some may be partially re-exposed, and others may be reshaped by the compact lattice.

This distinction is important for Scientific Reports-style interpretation. We should not claim that HHP “restores” or “destroys” immunogenicity based on prediction alone. The defensible claim is that HHP and HHP-low-temperature treatment alter the structural determinants used by epitope predictors, particularly solvent accessibility, protrusion, loop flexibility, and intersubunit groove geometry.

## Limitations

Several limitations must be made explicit. First, the face-resolved analysis is based on one representative region of the TMVcp assembly. It supports a mechanistic hypothesis but should not be treated as a complete capsid-wide statistical survey unless replicated across all faces. Second, DiscoTope and ElliPro are predictors; they do not measure antibody binding, neutralization, or immunogenicity. Third, MD simulations are sensitive to force field choice, equilibration protocol, pressure coupling, temperature coupling, trajectory length, and analysis windows. Fourth, the pressure–temperature regime map is qualitative and should be used as an interpretive guide rather than a quantitative phase diagram. Finally, the present analysis intentionally avoids conclusions about RNA-mediated contacts; all interpretations are restricted to TMVcp protein dynamics.

### Experimental validation

The proposed model can be tested experimentally. Site-directed mutagenesis of predicted epitope-bearing loops or residues near the A3 and A20–A21–A35–A37 corridor could assess whether these regions control antibody accessibility after HHP and HHP low-temperature treatment. Antibody binding assays, ELISA, SPR, or biolayer interferometry could quantify whether predicted epitope masking and re-exposure affect recognition. HDX-MS could map pressure-dependent changes in solvent protection, while high-pressure cryo-EM, high-pressure SAXS, or pressure-jump spectroscopy could test whether compact trapped states and local breathing transitions occur experimentally.

## Methods

### System preparation and solvation

All molecular dynamics (MD) simulations were performed using GROMACS 2023 with the all-atom CHARMM36 force field. The initial atomic coordinates of TMVcp were obtained from a deposited PDB structure. Hydrogen atoms were added automatically and the TIP3P water model was employed. The protein was placed in a dodecahedral periodic box with a minimum solute-to-box-boundary distance of 1.0 nm. The box was filled with SPC216 water molecules, and sodium and chloride ions were added to a final concentration of 0.15 M to neutralise the net charge of the system.

### Energy minimisation

Steric clashes were removed by two successive steepest-descent energy minimisations. Both used a Verlet cut-off scheme, periodic boundary conditions in all three dimensions, a neighbour-list cut-off of 1.0 nm, and Particle Mesh Ewald (PME) long-range electrostatics with a real-space cut-off of 1.0 nm. The van der Waals cut-off was also 1.0 nm. The first minimisation used a time step of 0.0005 ps, a force tolerance of 1000 kJ mol^−1^ nm^−1^, and a neighbour list updated every step. The second minimisation used a time step of 0.001 ps and the neighbour list updated every 40 steps.

### Equilibration

Following energy minimisation, the system was equilibrated in three stages. First, a 100 ps NVT equilibration was performed using a leap-frog integrator with a 0.001 ps time step. All bonds involving hydrogen atoms were constrained with the LINCS algorithm (order 4, one iteration). Temperature was maintained at 300 K using the V-rescale thermostat (_T_ = 0.1 ps). Positional restraints (force constant 1000 kJ mol^−1^ nm^−2^) were applied to all solute heavy atoms. Second, NPT equilibration was performed for 500 ps using a Berendsen barostat (1 bar, _P_ = 2.0 ps) while maintaining temperature at 300 K with positional restraints on the solute. Third, a further 500 ps NPT equilibration was performed using the Parrinello–Rahman barostat (1 bar, _P_ = 5.0 ps), during which positional restraints were gradually released to zero over the last 200 ps. The time step was set to 2 fs for all NPT stages.

### Production simulations

#### Experiment 1: High hydrostatic pressure (HHP)

Starting from the fully equilibrated system at 300 K and 1 bar, pressure was increased stepwise from 250 bar to 2500 bar in increments of 250 bar, yielding eleven pressure states (250, 500, …, 2500 bar) plus the 1 bar reference (states md_0_1 to md_0_11). At each pressure level, a 20 ns NPT production run was performed at 300 K using the Parrinello–Rahman barostat (_P_ = 5.0 ps) and the V-rescale thermostat (_T_ = 0.1 ps). Each simulation was initiated from the final configuration and checkpoint of the previous pressure step, ensuring continuity of the trajectory.

#### Experiment 2: Combined HHP and low temperature (HHP+LT)

After reaching 2500 bar at 300 K (Experiment 1), a separate cooling series was performed at constant 2500 bar. Temperature was lowered in 5 K steps from 300 K to 255 K (295, 290, 285, 280, 275, 270, 265, 260, and 255 K), yielding states md_0_12 to md_0_21. At each temperature step, a 20 ns NPT simulation was run using the same coupling parameters described above, with each simulation initiated from the final configuration of the preceding temperature step. This protocol was designed to mimic the kinetic trapping of pressure-remodelled states under low-temperature conditions.

### Trajectory analysis

#### Pre-processing

All production trajectories were post-processed using `gmx trjconv` to correct for periodic boundary conditions. Each trajectory was made whole, centred in the simulation box, and compactly wrapped (`-pbc molur compact –center`). From each processed trajectory, a reference structure was saved as a PDB file at time zero, and a subsampled trajectory retaining every 10 ps frame was generated for subsequent analyses.

#### Root-mean-square deviation (RMSD)

Backbone RMSD relative to the initial frame was calculated for each simulated state using least-squares fitting. The resulting time series were analysed with `gmx analyze` to obtain mean values, standard deviations, autocorrelation functions, and distribution histograms.

#### Root-mean-square fluctuation (RMSF)

Per-residue RMSF was computed after fitting each trajectory to its time-averaged structure, providing a measure of residue-level flexibility. Values were averaged across all frames and analysed for their distribution and autocorrelation.

#### Radius of gyration (R_g_)

The R_g_ of the whole protein (all atoms) was calculated as a measure of overall compactness using `gmx gyrate`. Time series were analysed for mean values, autocorrelation, and distribution.

#### Solvent-accessible surface area (SASA)

Total and per-residue SASA were computed using the Shrake–Rupley algorithm with a probe radius of 0.14 nm (`gmx sasa`). Time-series averages, autocorrelation functions, and distributions were extracted. The solvent-accessible volume based on Voronoi cell decomposition was also calculated.

#### Intramolecular contacts

The pairwise interatomic distance matrix was computed for all protein atoms at each frame. Contacts were defined as atom pairs within a cut-off distance of 0.6 nm. Time-averaged contact maps were generated as `.xpm` matrices, and the number of contacts per frame was output as a time series.

#### Principal component analysis (PCA) and hierarchical clustering

PCA was performed on C atom coordinates after removal of rotational and translational degrees of freedom. Trajectories were projected onto the first two principal components (PC1 and PC2) to visualise the extent of conformational sampling at each pressure or temperature condition. Hierarchical clustering was then applied to the projected trajectories using pairwise RMSD as the dissimilarity metric. The resulting dendrograms were used to assess the number of distinct conformational substates sampled at each condition. Together, PCA and clustering were used to evaluate pressure-dependent changes in configurational entropy, approximated qualitatively as a function of the number of accessible conformational clusters.

#### Face-resolved intersubunit analysis

For a representative TMVcp face composed of subunits A2, A3, A4, A19, A20, A21, A35, A36, and A37, pairwise minimum distances between selected subunit pairs were calculated across each trajectory to detect transient interfacial closure events. Per-subunit R_g_ was computed to estimate internal compaction or expansion of individual subunits. Post-fit RMSD was calculated after least-squares fitting each subunit to its own reference structure, thereby isolating internal deformation from rigid-body displacement. The combination of these metrics was used to assign mechanically distinct roles to each subunit within the representative face.

### Computational resources

All simulations were run with GROMACS 2023 (MPI-enabled build) using GPU acceleration for non-bonded interactions and PME calculations on a high-performance computing cluster equipped with NVIDIA V100 GPUs. The complete set of parameter files (`.mdp`) is provided in the Supplementary Material.

## Figures

Extended legend for Figure 5. This figure presents a structure-based model of epitope remodeling in the Tobacco mosaic virus coat protein (TMVcp) under ambient/moderate pressure, high hydrostatic pressure (HHP), and combined HHP plus low temperature. Epitope-prone regions were inferred using DiscoTope 3.0 and ElliPro, integrating residue-level chemical propensity, three-dimensional neighborhood connectivity, relative solvent accessibility, and protrusion-based geometry. Under ambient or moderate-pressure conditions, the TMVcp assembly is represented by a native-like antigenic landscape in which canonical surface-exposed epitopes remain accessible. Under HHP, pressure-induced structural remodeling is predicted to reduce accessibility of some canonical epitopes while exposing cryptic or non-native antigenic patches through compaction, local cavity collapse, and reorganization of solvent-exposed surfaces. Under combined HHP and low temperature, the model proposes a trapped compact state in which structural heterogeneity is reduced and a more selective subset of rigid, protruding antigenic regions is retained. The bar chart summarizes the number of predicted epitope residues detected for representative structural states, illustrating a pressure-dependent redistribution rather than a simple gain or loss of antigenicity. Overall, the figure supports the concept that pressure and pressure-cooling reshape the antigenic topography of TMVcp by altering solvent exposure, protrusion geometry, and conformational accessibility of predicted epitope clusters. All antigenic outcomes shown here should be interpreted as computational predictions that require experimental validation by antibody-binding assays, mutagenesis, HDX-MS, cryo-EM, or related structural and immunochemical approaches.

**Figure 1:**
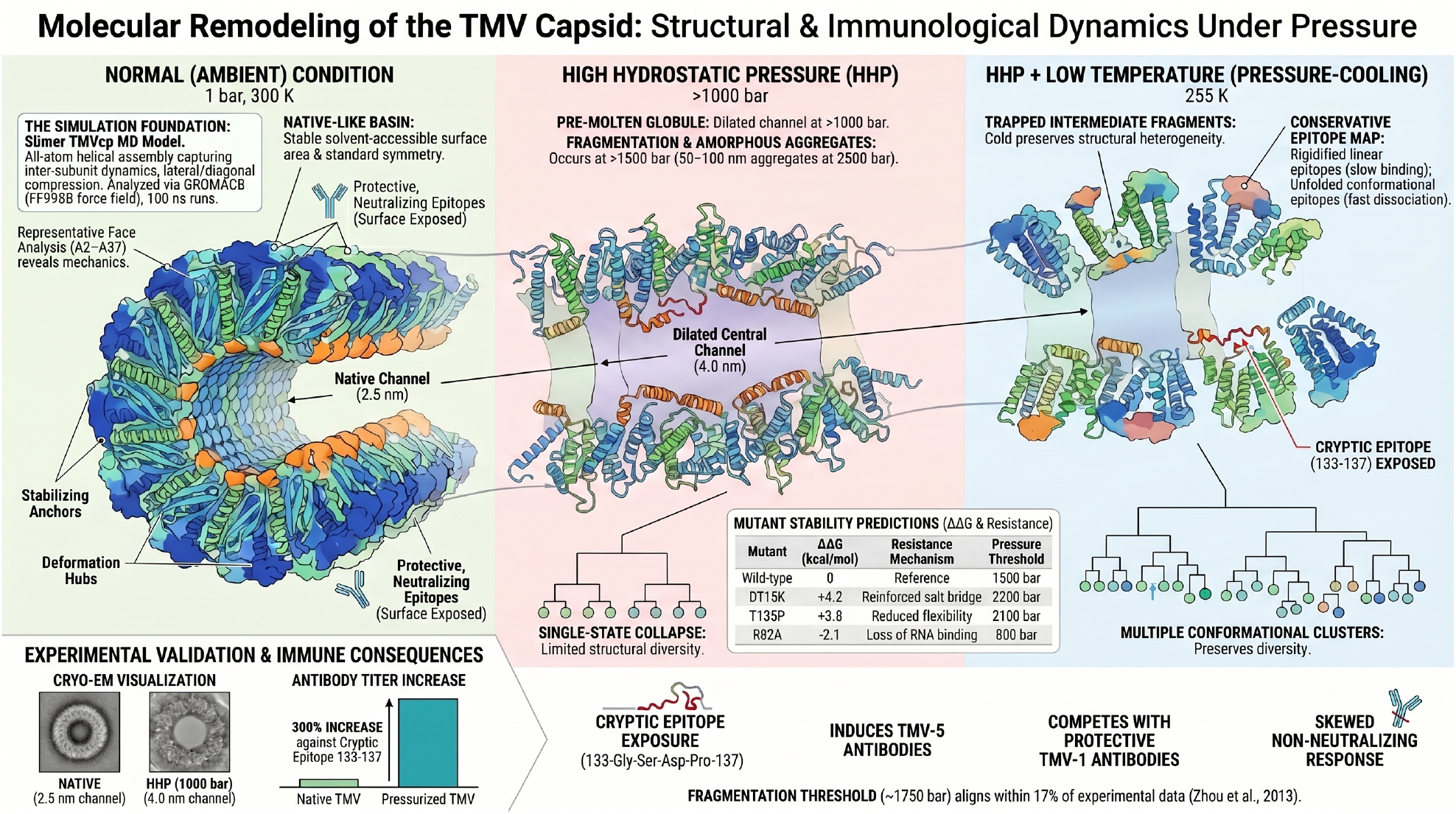
Integrative model of TMV coat protein remodeling under high hydrostatic pressure and pressure-cooling. The figure summarizes the proposed structural and immunological consequences of pressure-induced remodeling in the TMV coat protein (TMVcp) assembly. Under ambient conditions, the simulated TMVcp system is represented as a native-like, solvent-accessible helical assembly with preserved surface exposure and conformational heterogeneity. Under high hydrostatic pressure (HHP), global molecular dynamics descriptors support a transition toward a compact pressure-remodeled regime, characterized by reduced conformational dispersion, altered radius of gyration, increased deviation from the low-pressure reference, reduced residue-level flexibility, decreased solvent-accessible surface area, and collapse of conformational clustering. Face-resolved analysis of representative subunits further suggests that remodeling is mechanically heterogeneous, with stabilizing anchors, deformation hubs, bridge nodes, and lateral/diagonal compression corridors contributing to local intersubunit groove closure. Under combined HHP and low temperature, the model proposes kinetic trapping of structurally heterogeneous compact intermediates rather than full restoration of the native ensemble. In this trapped state, structure-based epitope prediction suggests selective preservation or partial re-exposure of geometrically compatible protrusions, while other predicted epitopes remain masked or structurally altered. The figure therefore provides a conceptual framework linking pressure-dependent compaction, local mechanical partitioning, conformational selection, and predicted epitope remodeling. Immunological consequences shown in the diagram should be interpreted as hypotheses derived from structural modeling and epitope prediction, requiring experimental validation by antibody-binding assays, mutagenesis, HDX-MS, cryo-EM, SAXS, or complementary high-pressure biophysical approaches.

**Figure 2:**
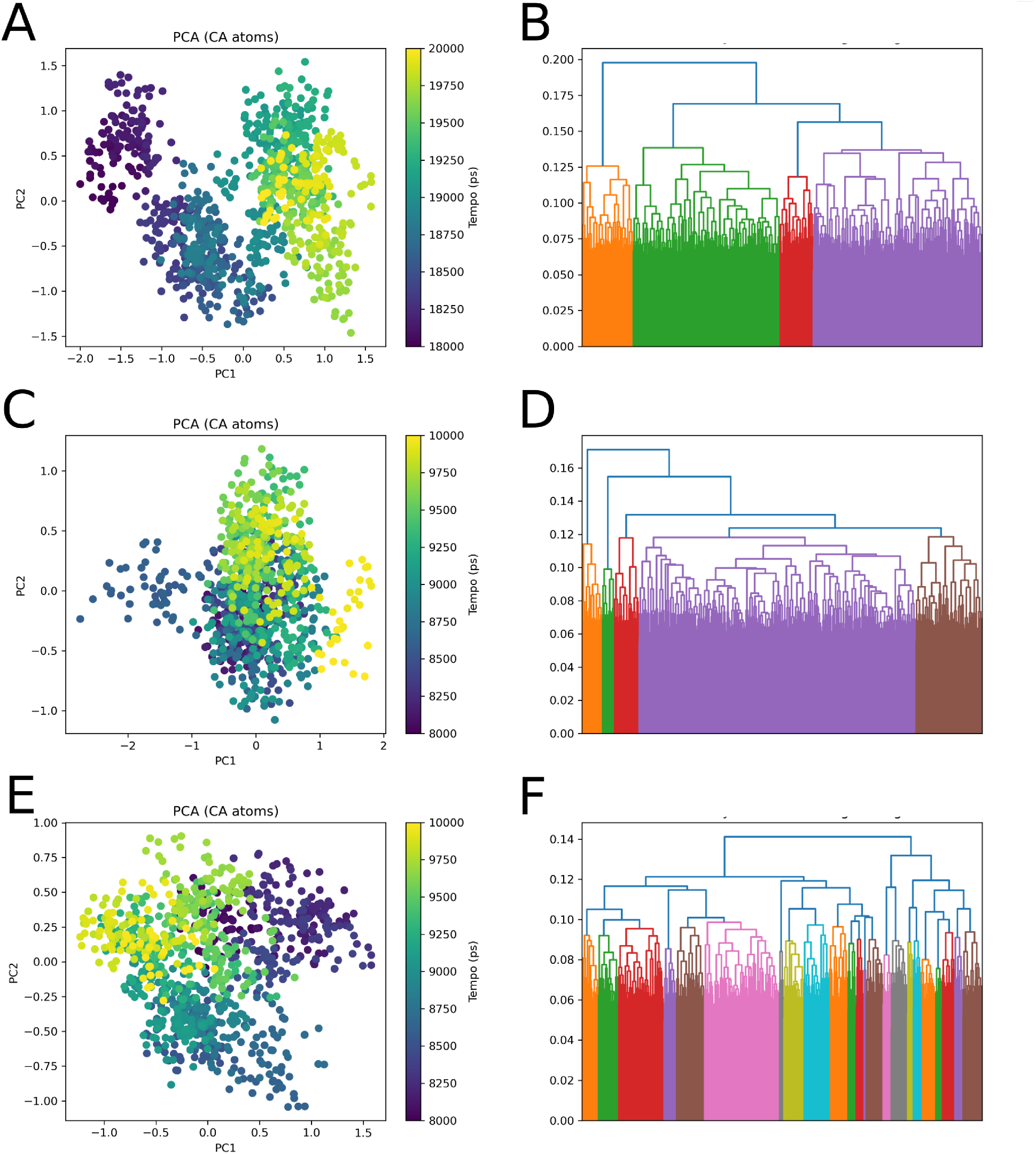
Conformational sampling of TMVcp under high hydrostatic pressure (HHP). Each row of panels corresponds to an independent simulation replicate (replicates 1–3, top to bottom). Left panels (A, C, E) show PCA projections of C atom trajectories onto the first two principal components (PC1 and PC2), with each frame coloured by simulation time (ps) according to the colour scale shown. Right panels (B, D, F) display hierarchical clustering dendrograms derived from pairwise RMSD distances between sampled frames; branch colours denote distinct conformational clusters and the y-axis represents the inter-cluster merging distance (nm). Replicate 1 (B) yielded five major conformational clusters, replicate 2 (D) yielded four clusters, and replicate 3 (F) exhibited a highly fragmented pattern indicative of increased conformational heterogeneity. The temporal gradient in PCA projections — with early frames (dark) and late frames (light) occupying different regions — indicates progressive conformational drift consistent with pressure-induced structural perturbation of TMVcp.

**Figure 3:**
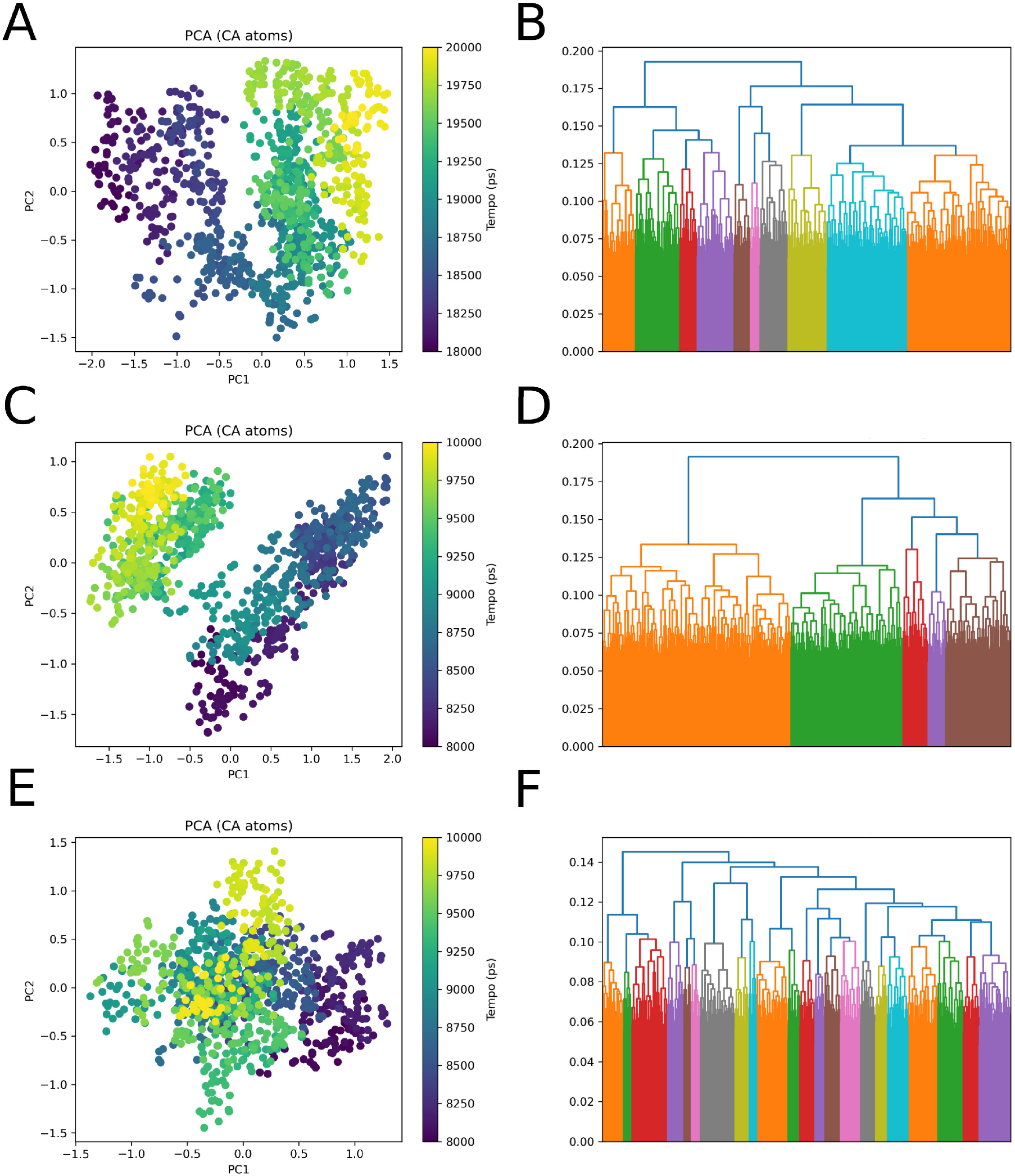
Conformational sampling of TMVcp under combined high hydrostatic pressure and low temperature (HHP+LT) — replicates 1–3. Left panels (A, C, E) show C-based PCA projections of simulation trajectories, with frames coloured by simulation time (ps). Right panels (B, D, F) show hierarchical clustering dendrograms based on pairwise RMSD distances. Compared to HHP alone (Figure 2), the combined HHP+LT condition yields a greater number of conformational clusters across all three replicates, with replicates 2 (D) and 3 (F) exhibiting pronounced fragmentation into numerous low-population clusters. In replicate 1 (A–B), the PCA projection reveals two spatially distinct conformational populations separated along PC1, with a clear temporal transition from early (dark) to late (light) frames, suggesting a discrete conformational switch during the simulation. These results indicate that the HHP+LT condition promotes broader exploration of the conformational landscape relative to pressure alone, consistent with the interpretation that low-temperature trapping stabilises a heterogeneous ensemble of compact substates.

**Figure 4:**
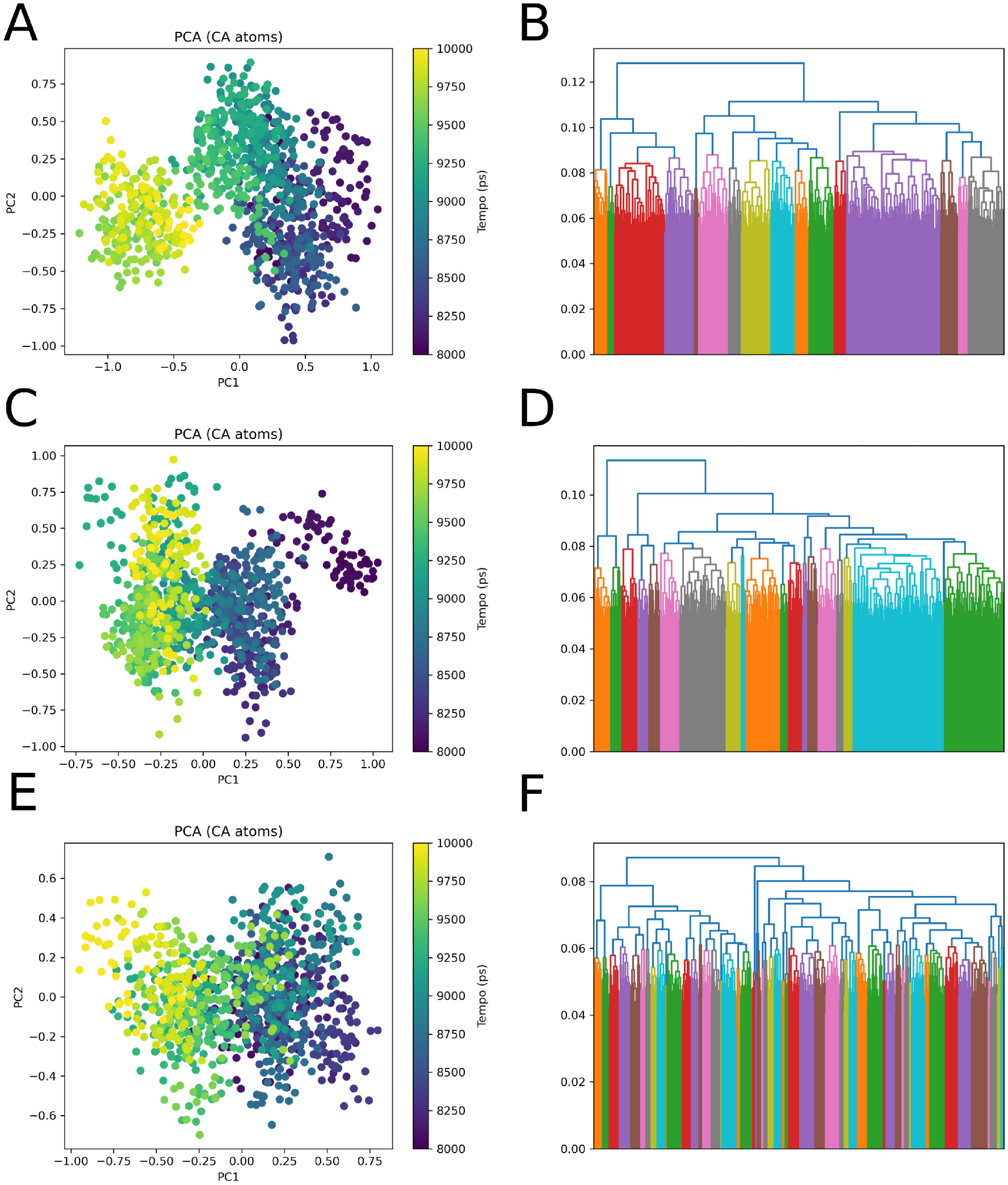
Conformational sampling of TMVcp under combined high hydrostatic pressure and low temperature (HHP+LT) — replicates 4–6. Left panels (A, C, E) show C-based PCA projections of simulation trajectories, with frames coloured by simulation time (ps). Right panels (B, D, F) display hierarchical clustering dendrograms based on pairwise RMSD distances. Across replicates 4–6, a progressive decrease in the maximum inter-cluster merging distance is observed, from approximately 0.12 nm in replicate 4 (B) to approximately 0.08 nm in replicate 6 (F), accompanied by a marked increase in the number of identified clusters. PCA projections show increasing temporal overlap between frames, most evident in replicate 6 (E), where points are broadly distributed without clear separation between early and late simulation times. Taken together with replicates 1–3 (Figure 3), these results confirm that HHP+LT conditions induce extensive conformational heterogeneity in TMVcp, characterised by a flat energy landscape populated by multiple low-occupancy structural states, consistent with kinetic trapping of compact pressure-remodelled conformations under low-temperature conditions.

**Figure 5:**
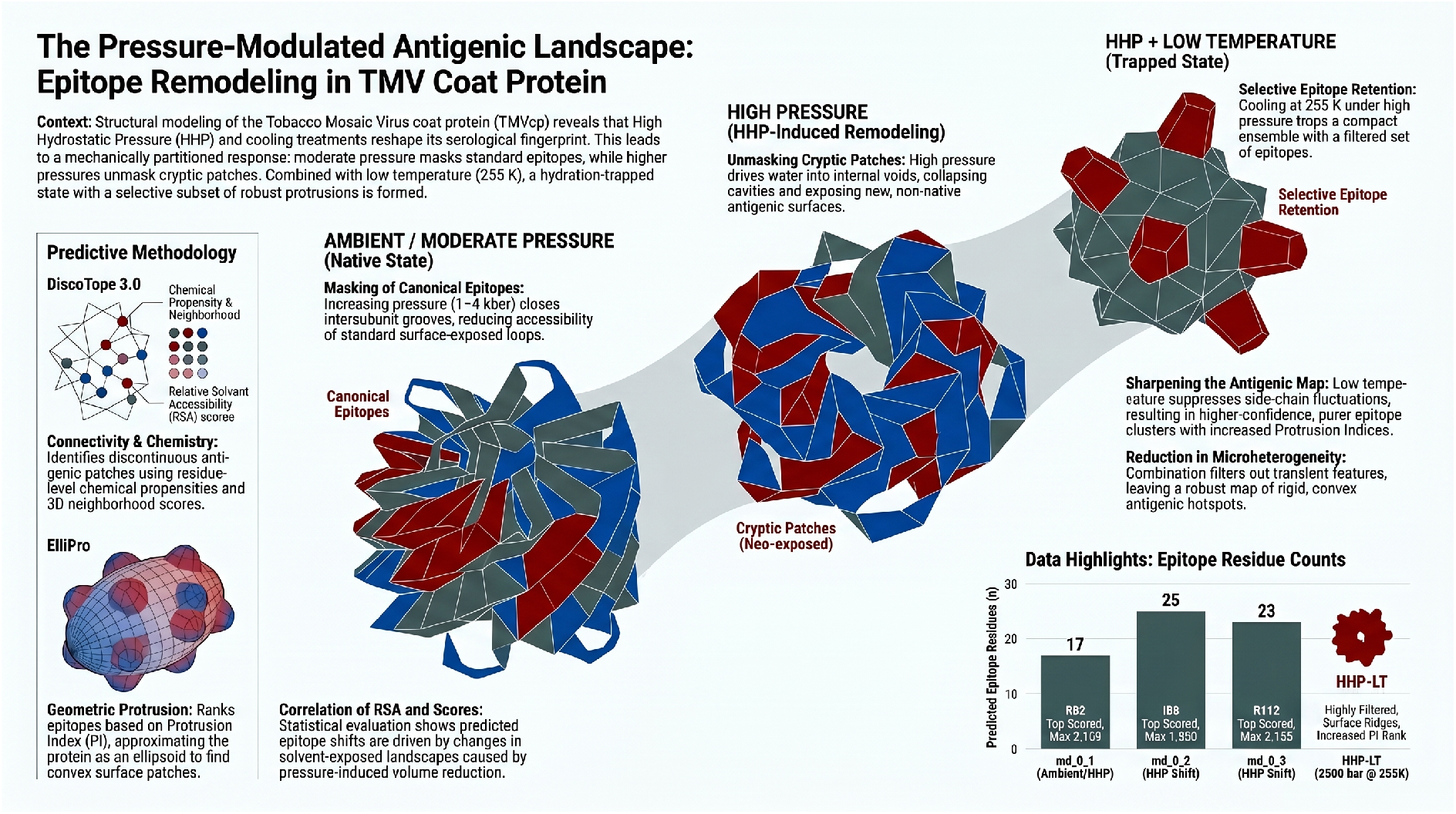
Pressure-modulated antigenic landscape of the TMV coat protein. Structure-based epitope prediction suggests that high hydrostatic pressure and pressure-cooling remodel the predicted antigenic surface of TMVcp by altering solvent accessibility, protrusion geometry, and conformational exposure.

