## Supplementary material for "Pressure–cooling remodeling of TMV coat protein reveals mechanically partitioned capsid dynamics and selective epitope masking": SM4

2026-05-31

### A Complete results

#### RMSD

##### HHP

##### HHP and low temperature

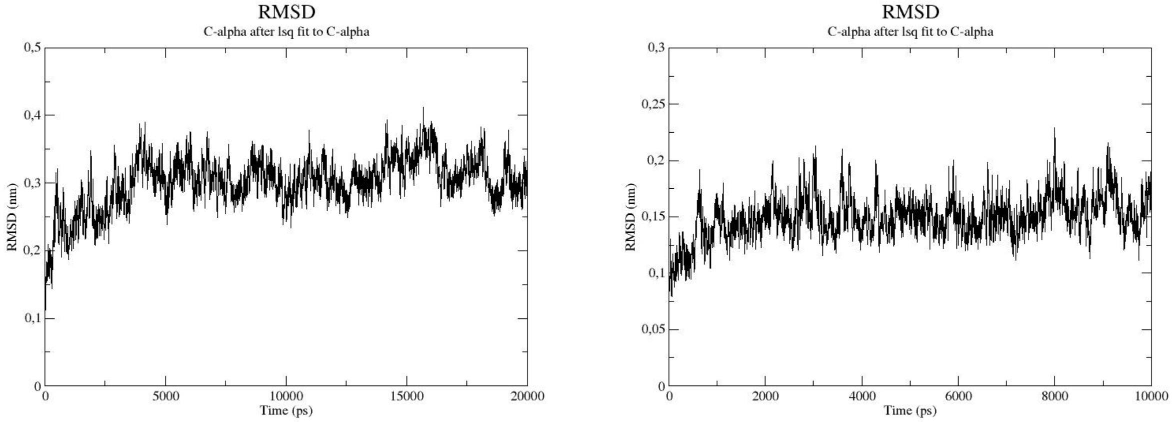

A) 1 bar B) 250 bar

Figure 1: RMSD side-by-side - High Hydrostatic Pressure

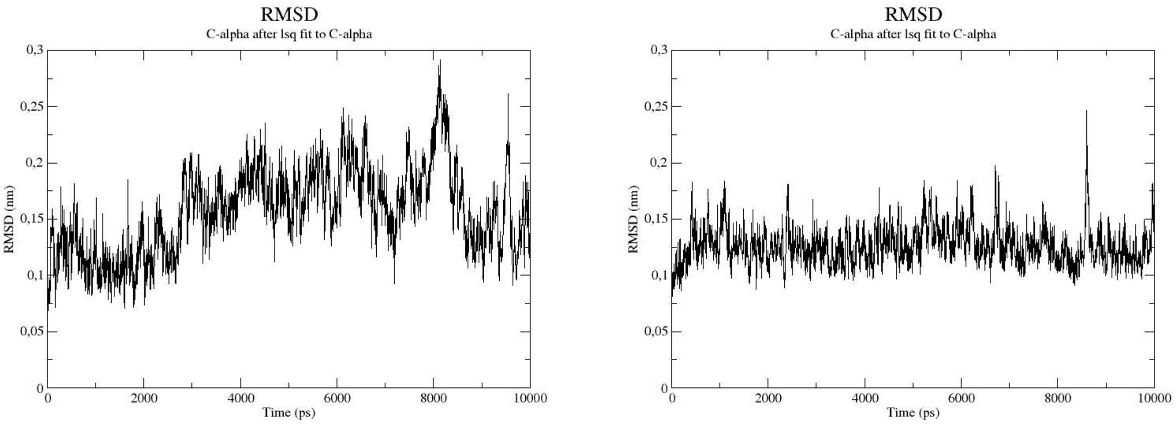

A) 500 bar B) 750 bar

Figure 2: RMSD side-by-side - High Hydrostatic Pressure

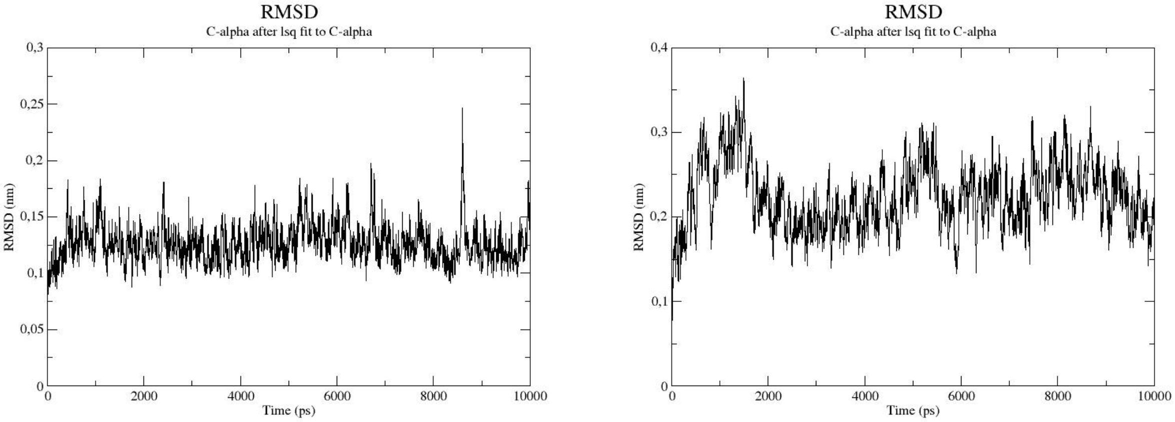

A) 1000 bar B) 1250 bar

Figure 3: RMSD side-by-side - High Hydrostatic Pressure

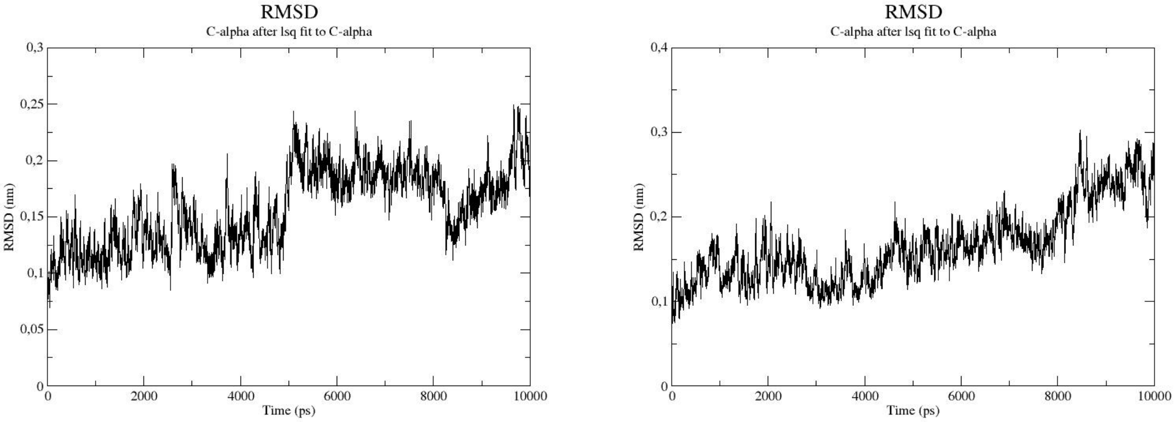

A) 1500 bar B) 1750 bar

Figure 4: RMSD side-by-side - High Hydrostatic Pressure

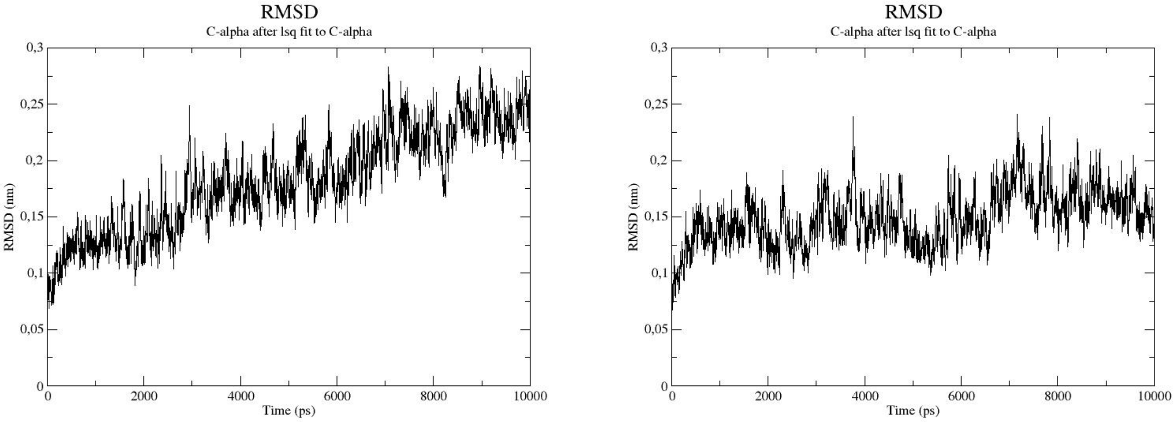

A) 2000 bar B) 2250 bar

Figure 5: RMSD side-by-side - High Hydrostatic Pressure

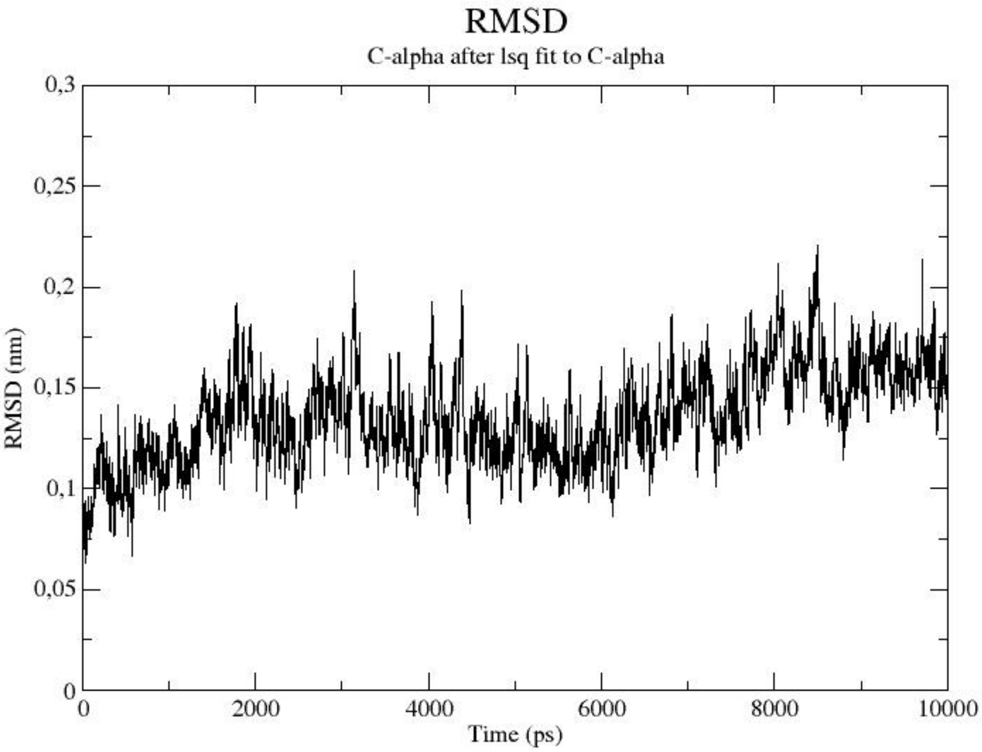

A) 2500 bar

Figure 6: RMSD side-by-side - High Hydrostatic Pressure

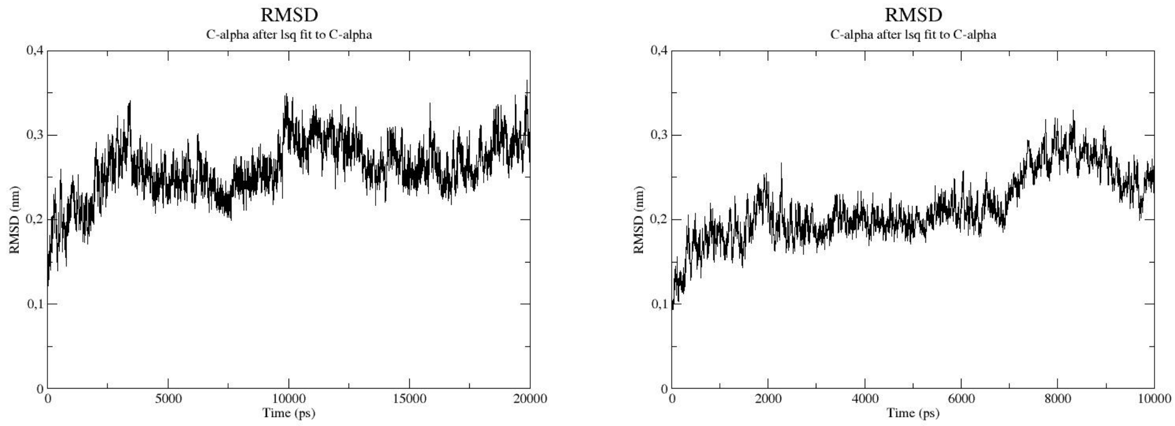

A) 1 bar B) 250 bar

Figure 7: RMSD side-by-side - High Hydrostatic Pressure and Low Temperature

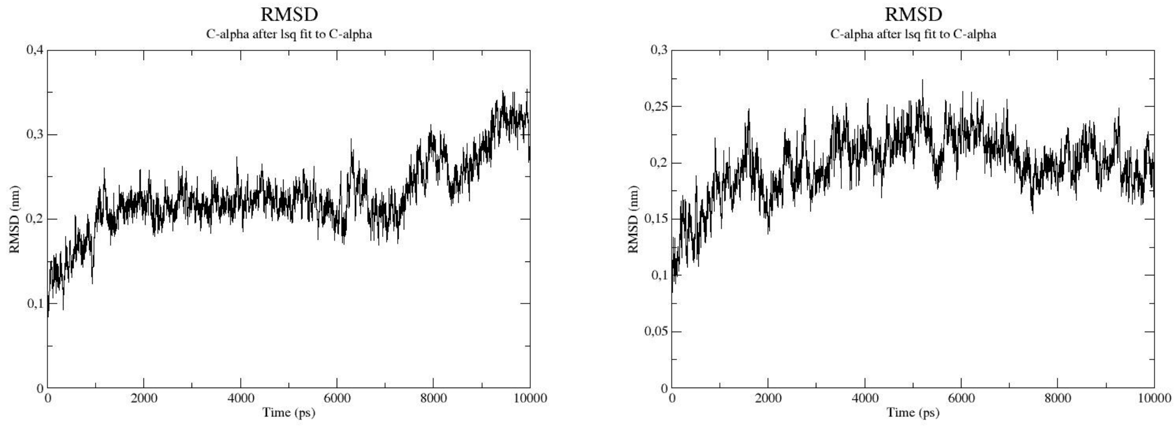

A) 500 bar B) 750 bar

Figure 8: RMSD side-by-side - High Hydrostatic Pressure and Low Temperature

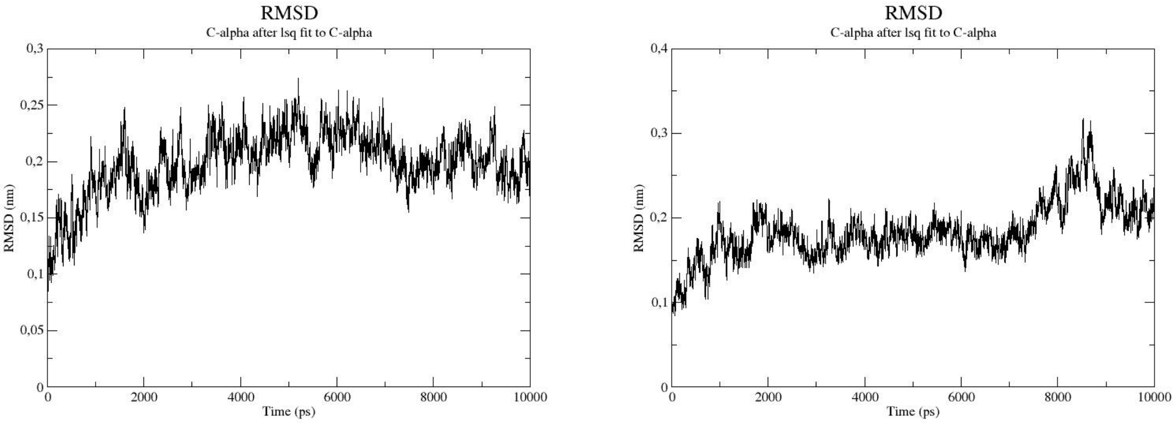

A) 1000 bar B) 1250 bar

Figure 9: RMSD side-by-side - High Hydrostatic Pressure and Low Temperature

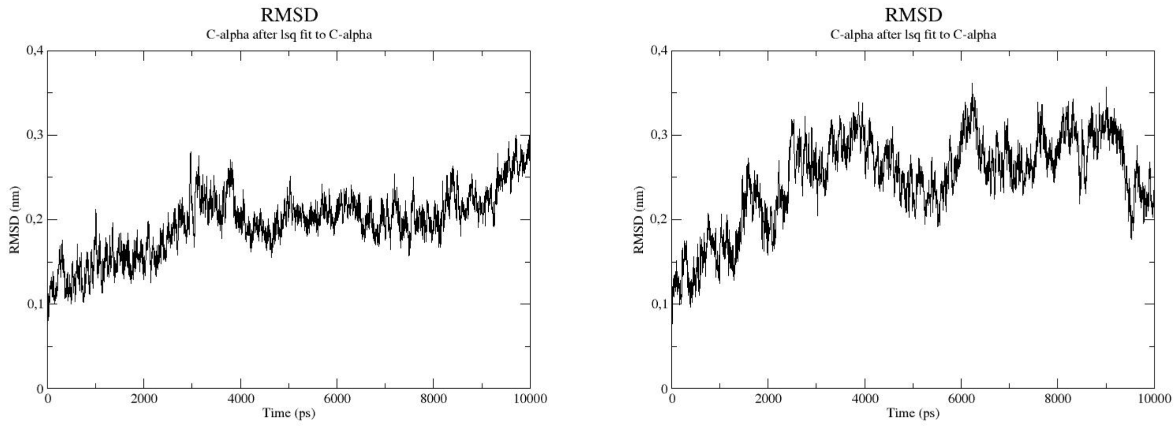

A) 1500 bar B) 1750 bar

Figure 10: RMSD side-by-side - High Hydrostatic Pressure and Low Temperature

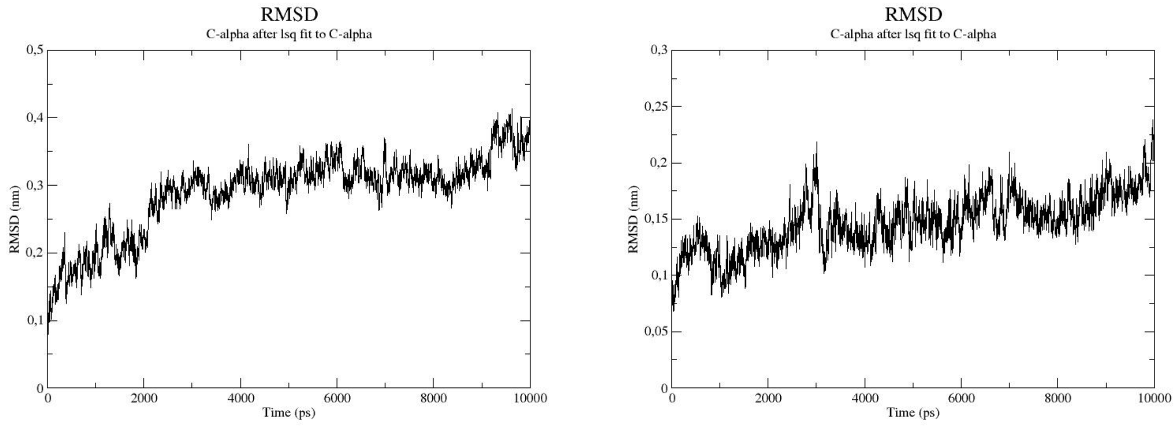

A) 2000 bar B) 2250 bar

Figure 11: RMSD side-by-side - High Hydrostatic Pressure and Low Temperature

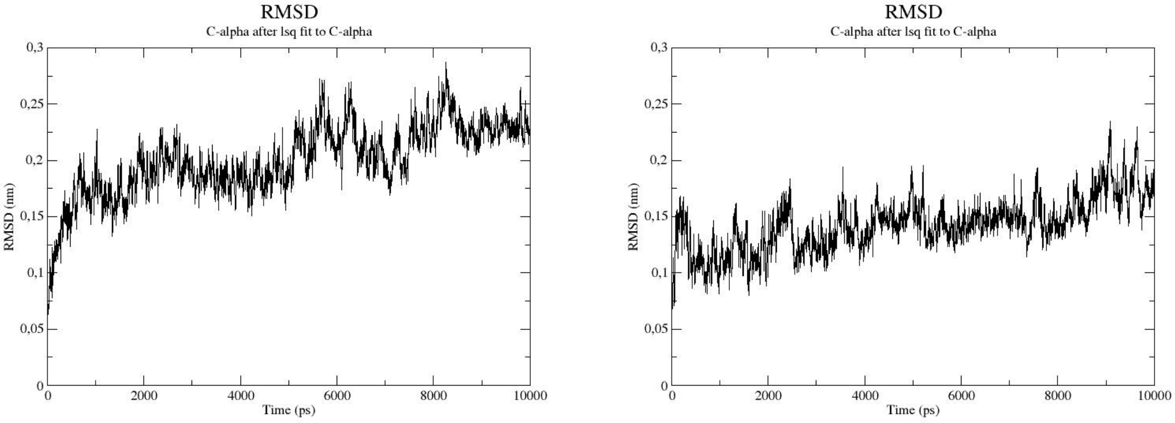

A) 2500 bar @ 300 K

B) 2500 bar @ 295 K

Figure 12: RMSD side-by-side - High Hydrostatic Pressure and Low Temperature

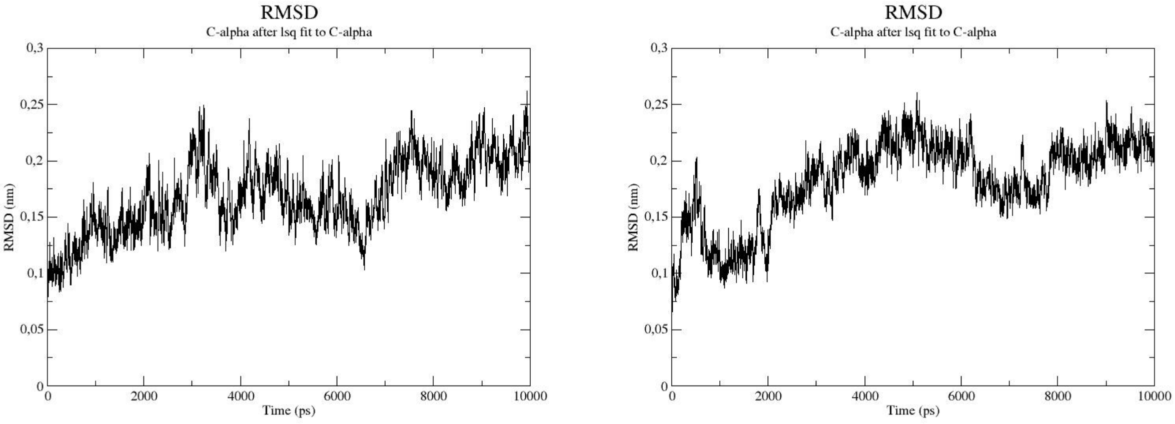

A) 2500 bar @ 290 K

B) 2500 bar @ 285 K

Figure 13: RMSD side-by-side - High Hydrostatic Pressure and Low Temperature

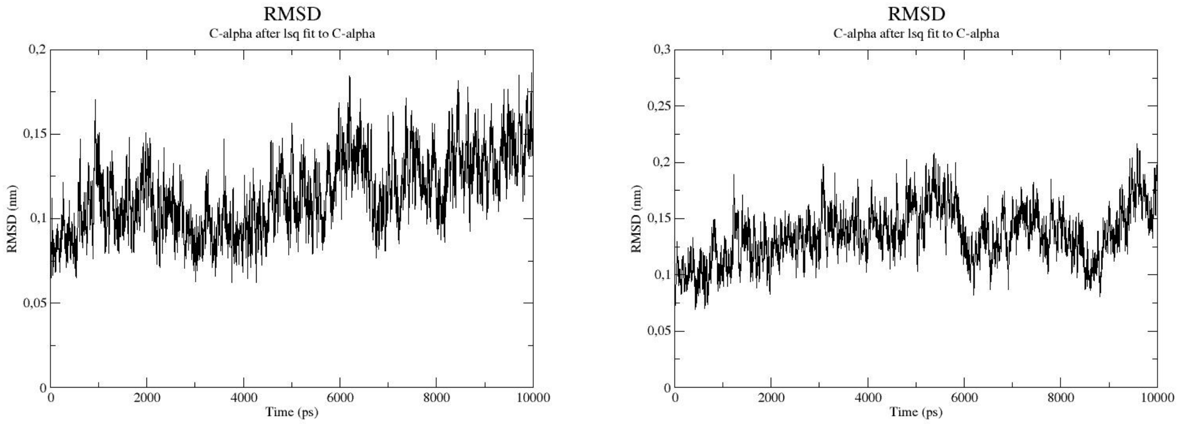

A) 2500 bar @ 280 K B) 2500 bar @ 275 K

Figure 14: RMSD side-by-side - High Hydrostatic Pressure and Low Temperature

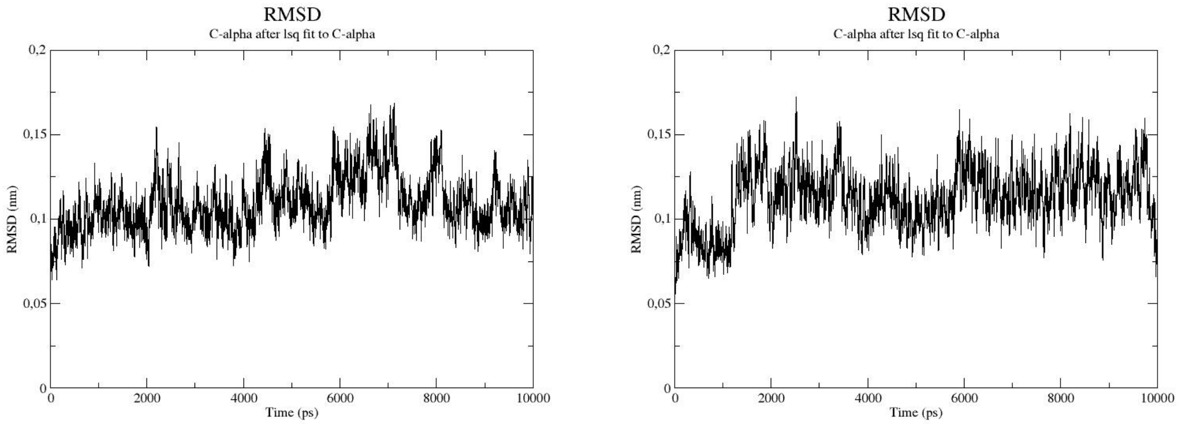

A) 2500 bar @ 270 K B) 2500 bar @ 265 K

Figure 15: RMSD side-by-side - High Hydrostatic Pressure and Low Temperature

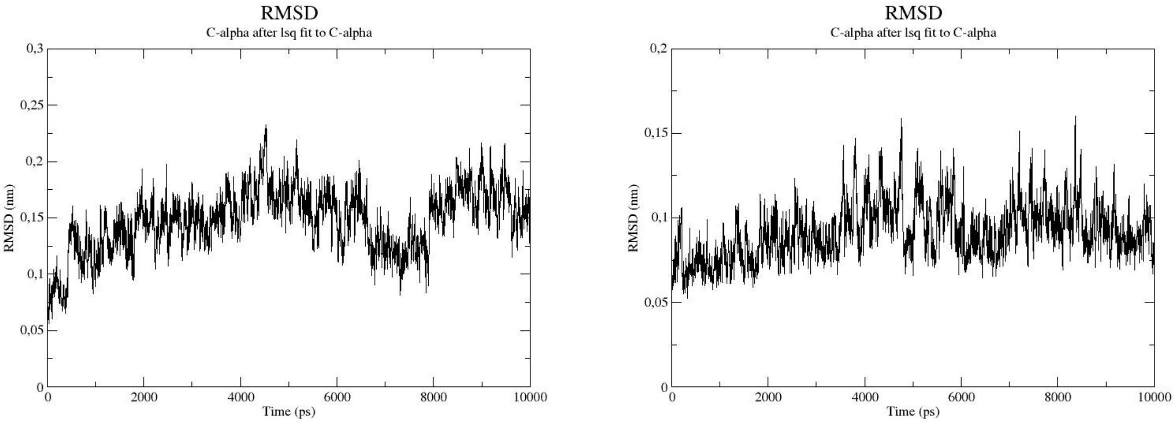

A) 2500 bar @ 260 K B) 2500 bar @ 255 K

Figure 16: RMSD side-by-side - High Hydrostatic Pressure and Low Temperature

#### RMSF

##### HHP

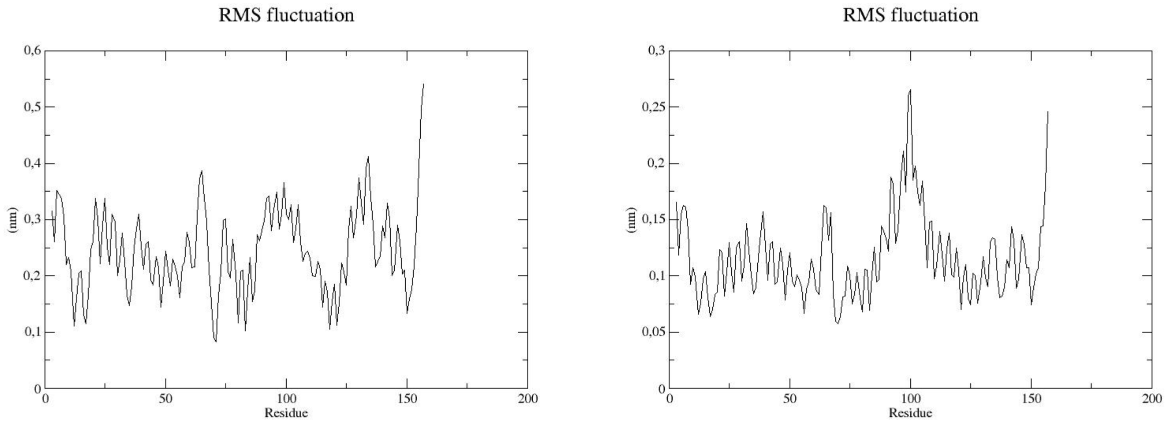

A) 1 bar B) 250 bar

Figure 17: RMSF side-by-side - High Hydrostatic Pressure

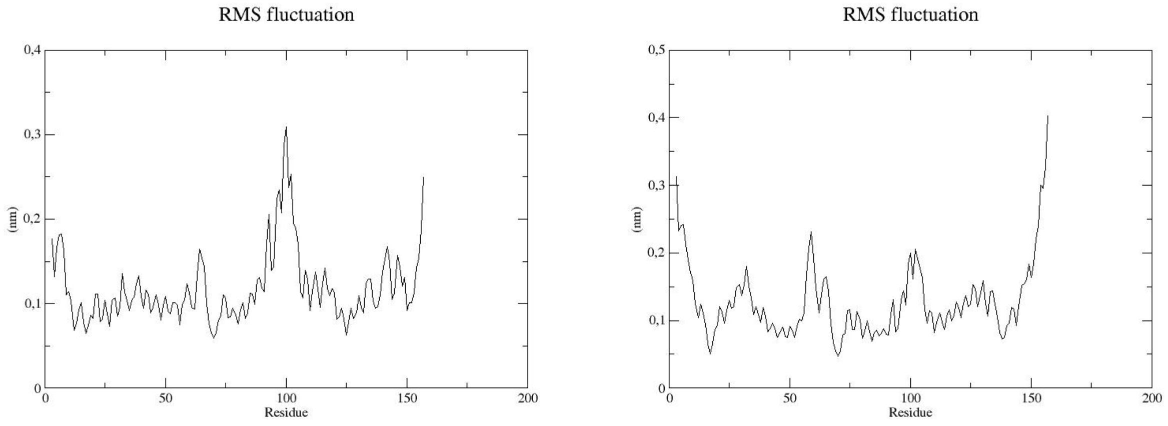

A) 500 bar B) 750 bar

Figure 18: RMSF side-by-side - High Hydrostatic Pressure

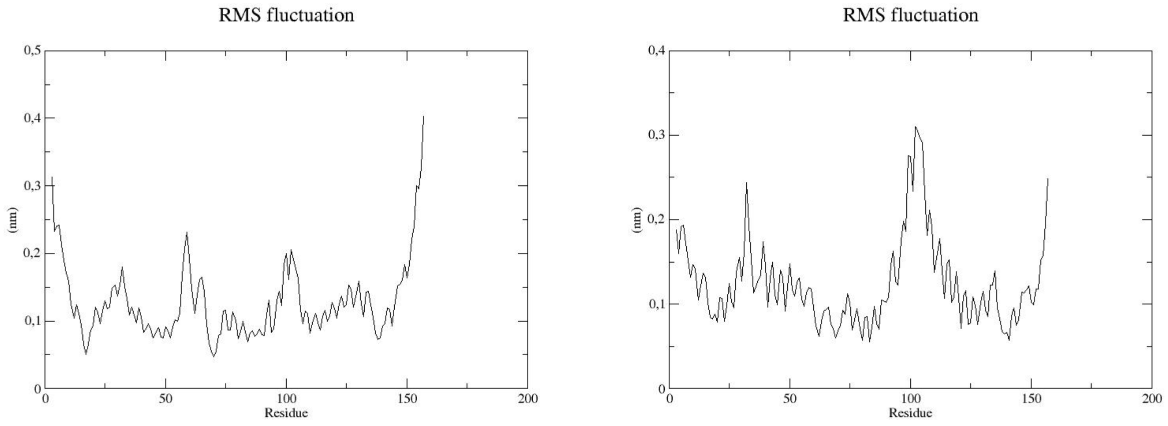

A) 1000 bar B) 1250 bar

Figure 19: RMSF side-by-side - High Hydrostatic Pressure

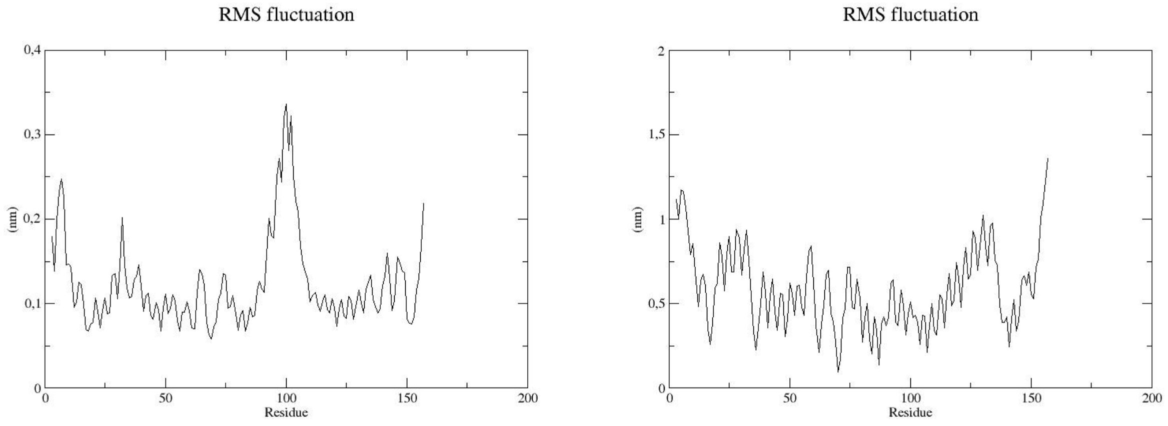

A) 1500 bar B) 1750 bar

Figure 20: RMSF side-by-side - High Hydrostatic Pressure

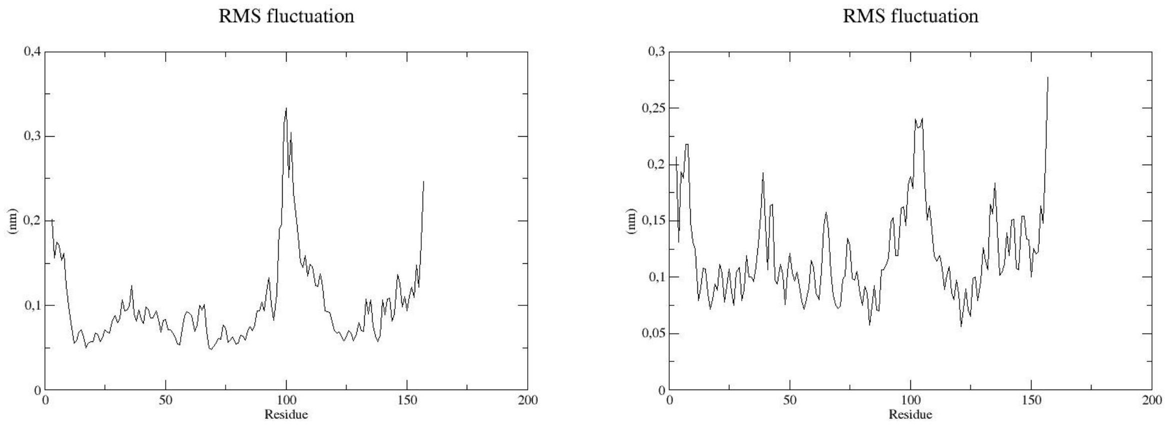

A) 2000 bar B) 2250 bar

Figure 21: RMSF side-by-side - High Hydrostatic Pressure

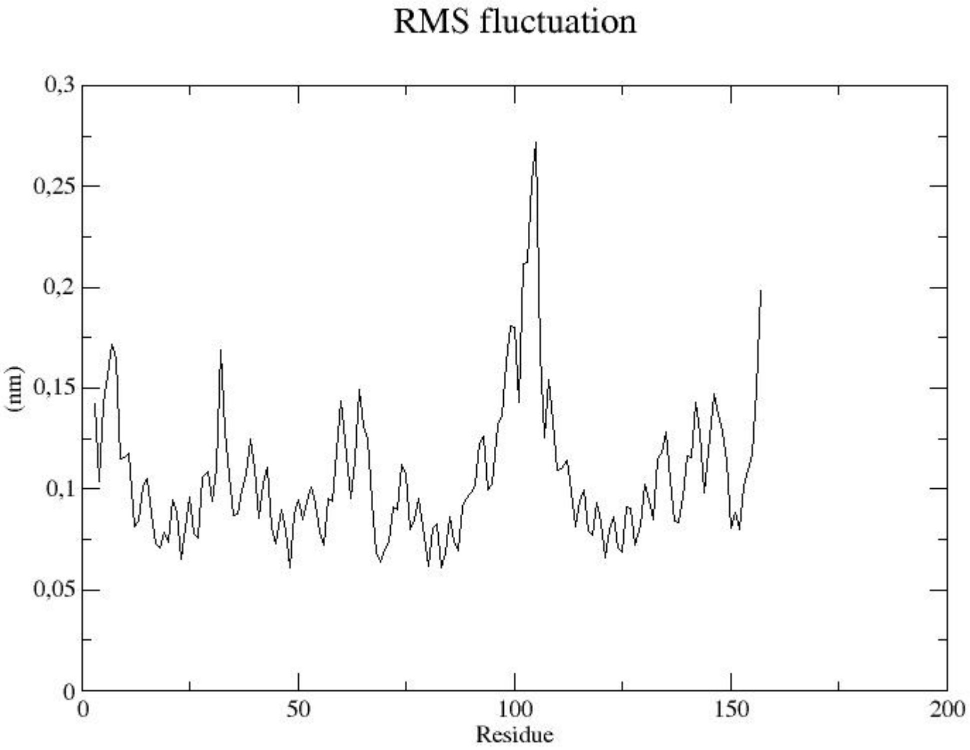

A) 2500 bar

Figure 22: RMSF side-by-side - High Hydrostatic Pressure

**HHP and low temperature**

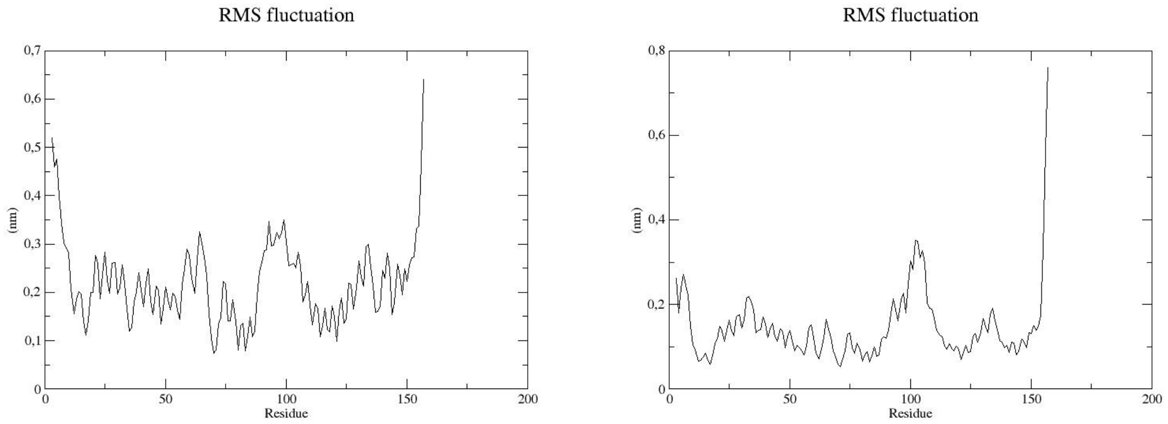

A) 1 bar B) 250 bar

Figure 23: RMSF side-by-side - High Hydrostatic Pressure and Low Temperature

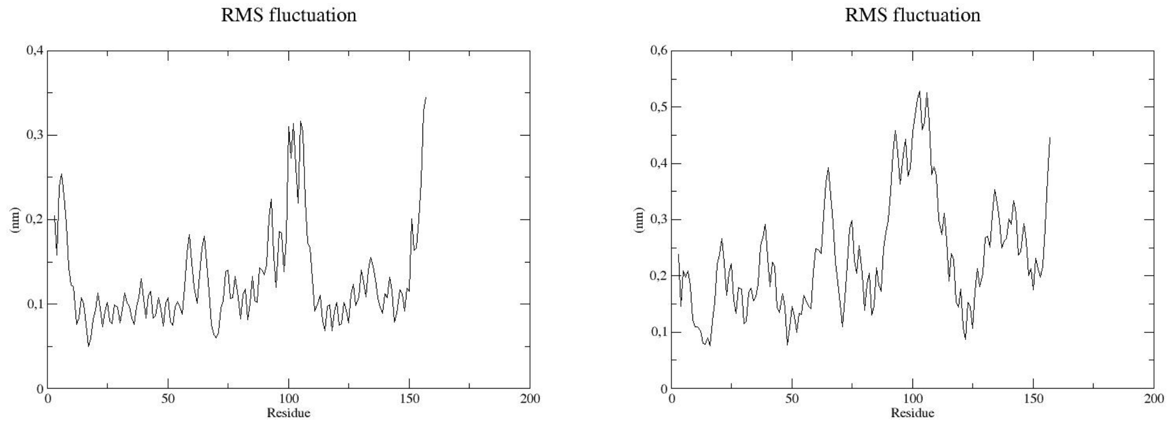

A) 500 bar B) 750 bar

Figure 24: RMSF side-by-side - High Hydrostatic Pressure and Low Temperature

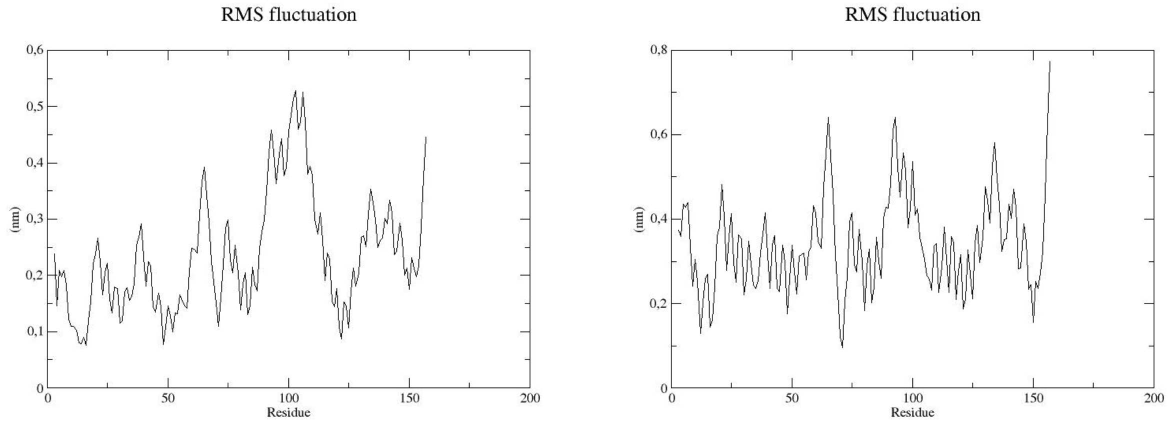

A) 1000 bar B) 1250 bar

Figure 25: RMSF side-by-side - High Hydrostatic Pressure and Low Temperature

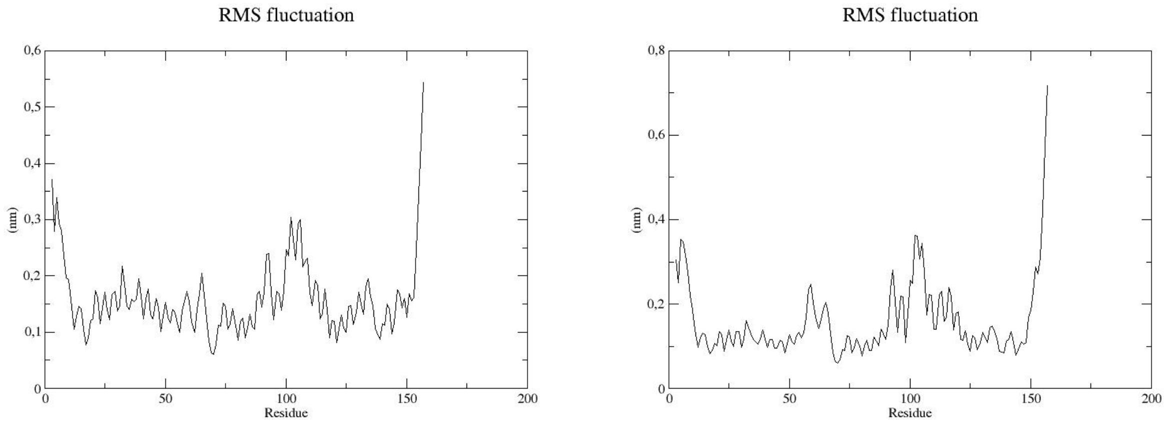

A) 1500 bar B) 1750 bar

Figure 26: RMSF side-by-side - High Hydrostatic Pressure and Low Temperature

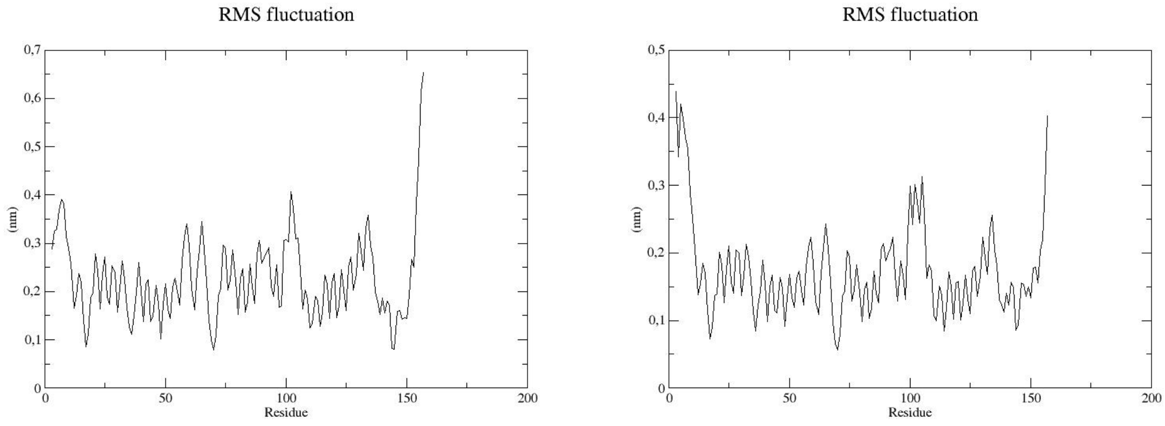

A) 2000 bar B) 2250 bar

Figure 27: RMSF side-by-side - High Hydrostatic Pressure and Low Temperature

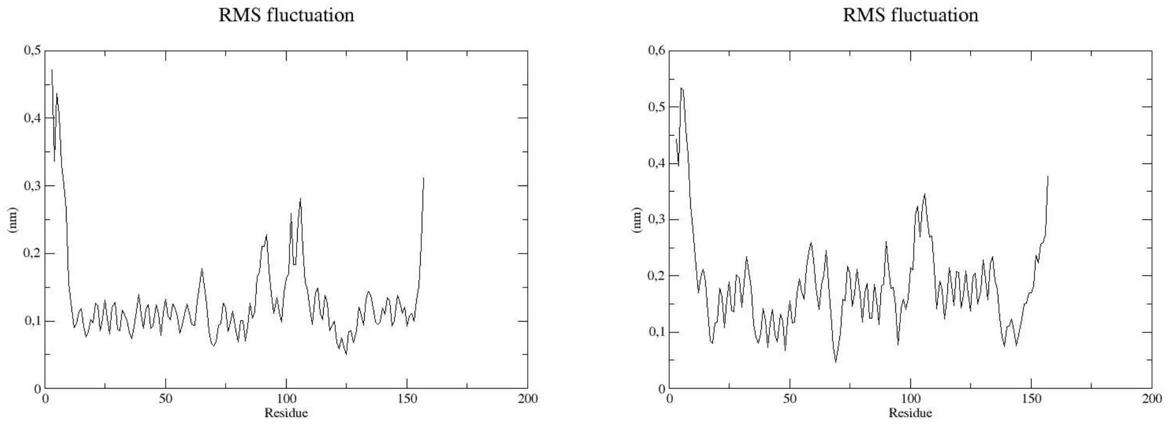

A) 2500 bar @ 300 K B) 2500 bar @ 295 K

Figure 28: RMSF side-by-side - High Hydrostatic Pressure and Low Temperature

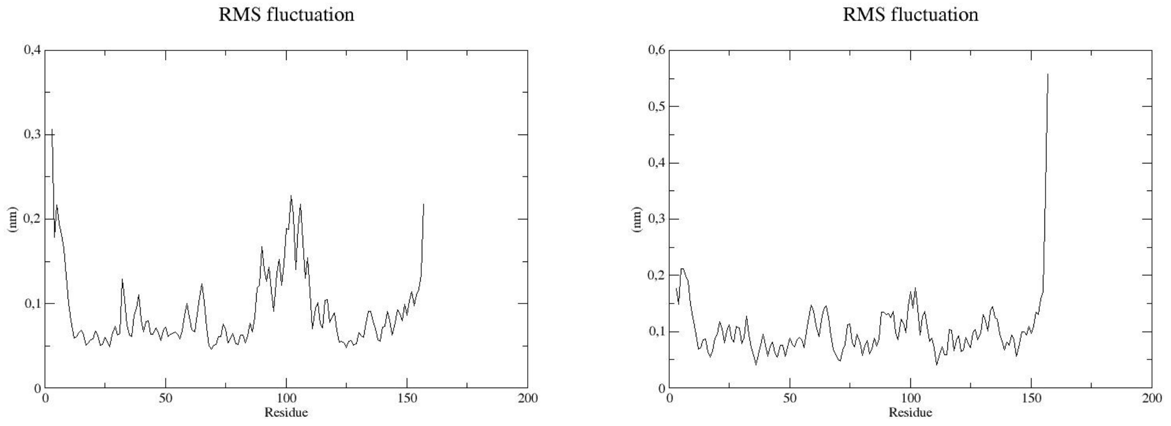

A) 2500 bar @ 290 K B) 2500 bar @ 285 K

Figure 29: RMSF side-by-side - High Hydrostatic Pressure and Low Temperature

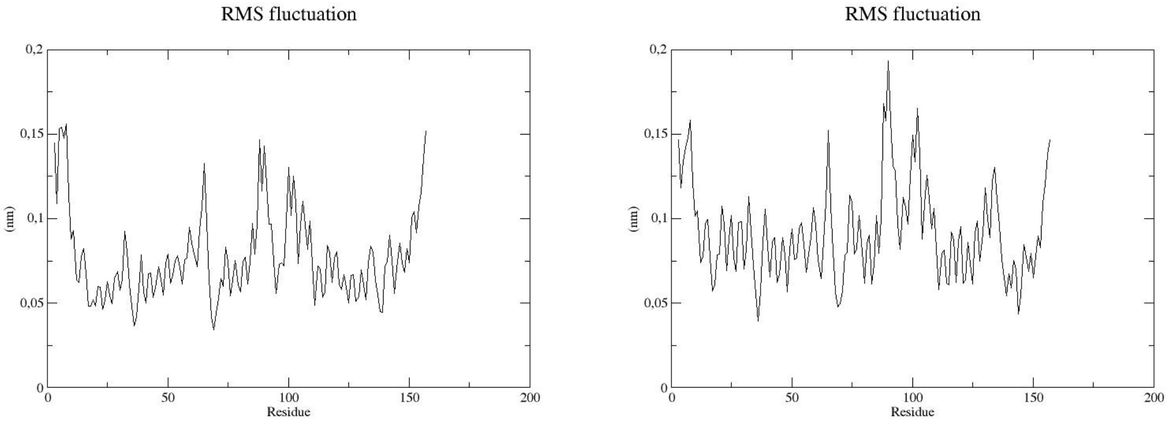

A) 2500 bar @ 280 K B) 2500 bar @ 275 K

Figure 30: RMSF side-by-side - High Hydrostatic Pressure and Low Temperature

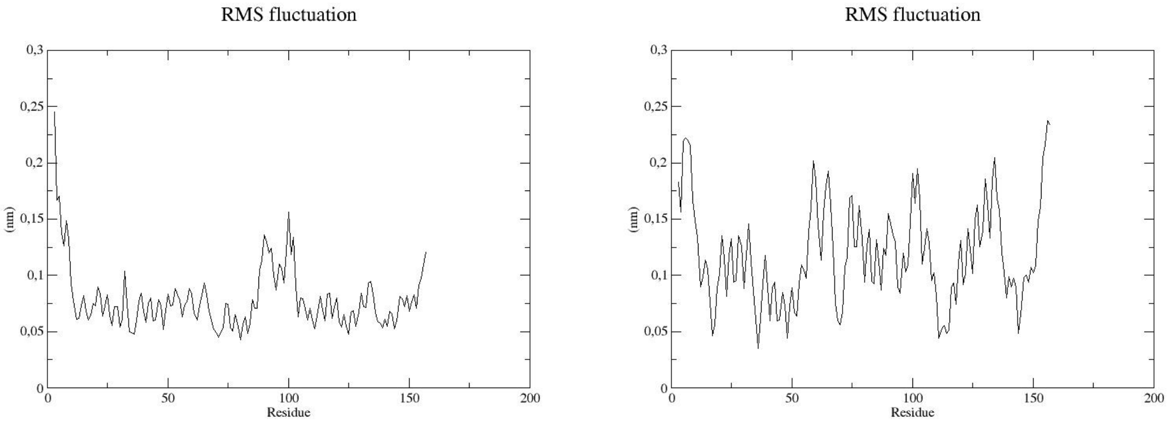

A) 2500 bar @ 270 K B) 2500 bar @ 265 K

Figure 31: RMSF side-by-side - High Hydrostatic Pressure and Low Temperature

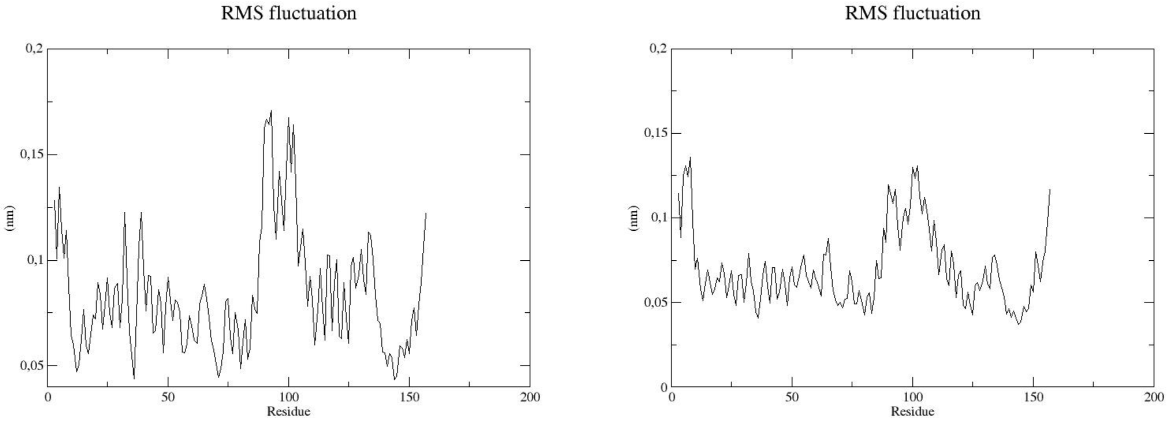

A) 2500 bar @ 260 K

B) 2500 bar @ 255 K

Figure 32: RMSF side-by-side - High Hydrostatic Pressure and Low Temperature

## Rg

##### HHP

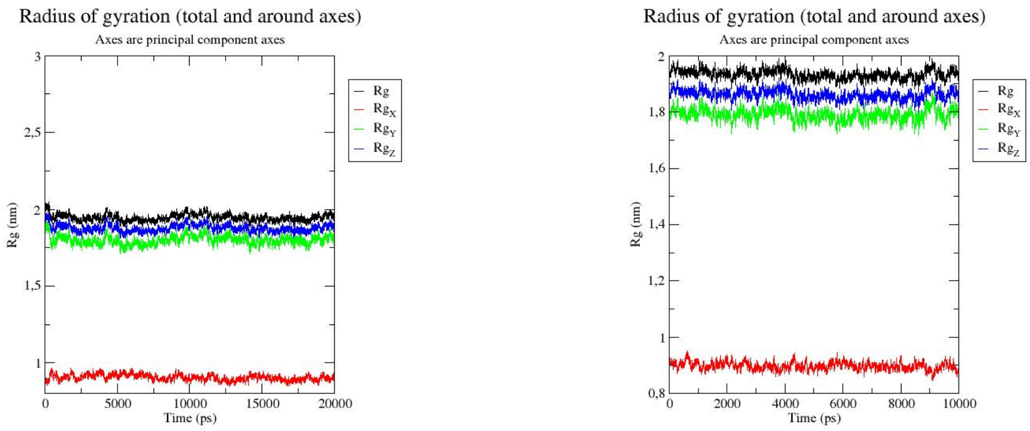

A) 1 bar B) 250 bar

Figure 33: Rg side-by-side - High Hydrostatic Pressure

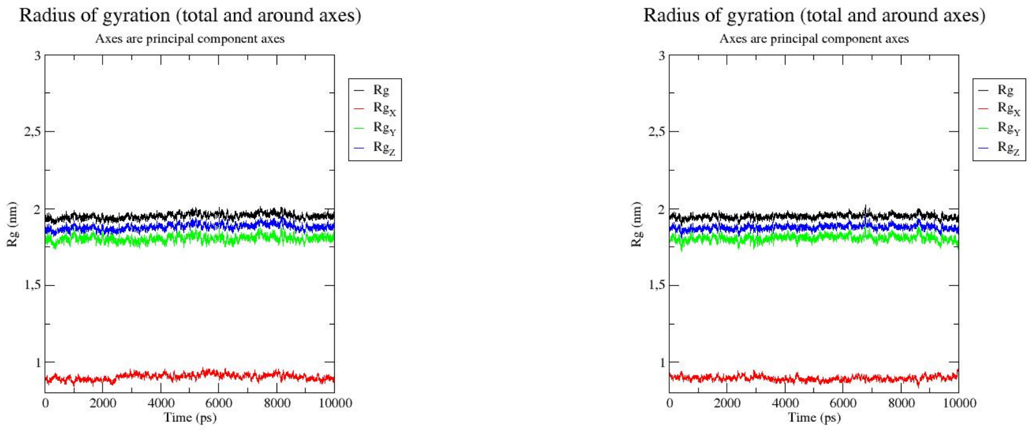

A) 500 bar B) 750 bar

Figure 34: Rg side-by-side - High Hydrostatic Pressure

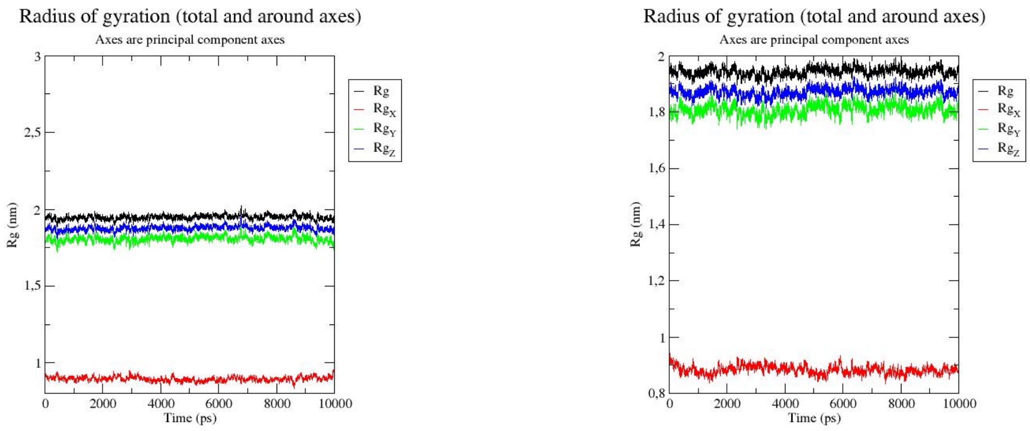

A) 1000 bar B) 1250 bar

Figure 35: Rg side-by-side - High Hydrostatic Pressure

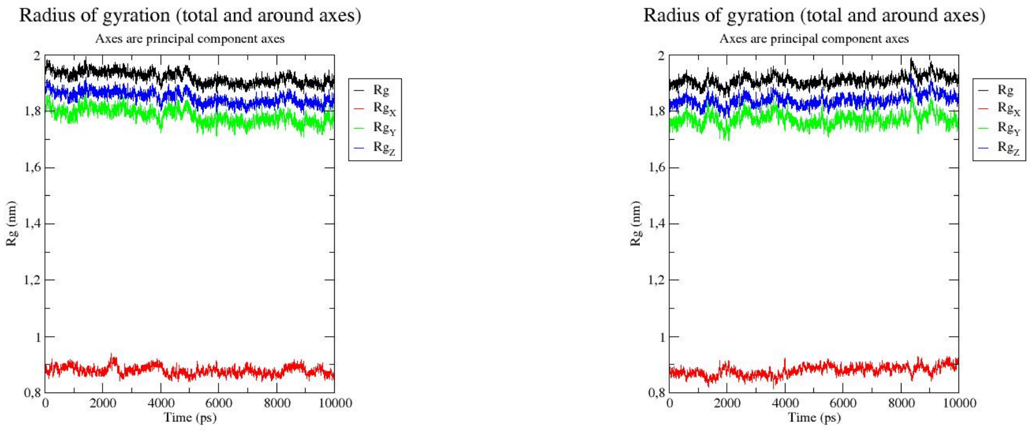

A) 1500 bar B) 1750 bar

Figure 36: Rg side-by-side - High Hydrostatic Pressure

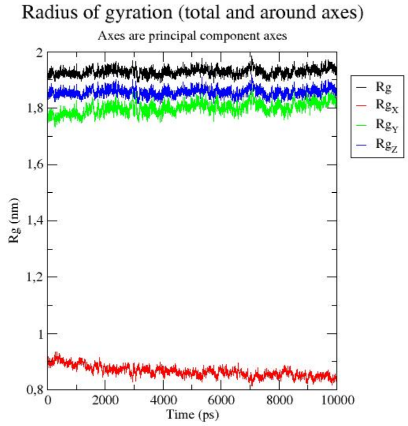

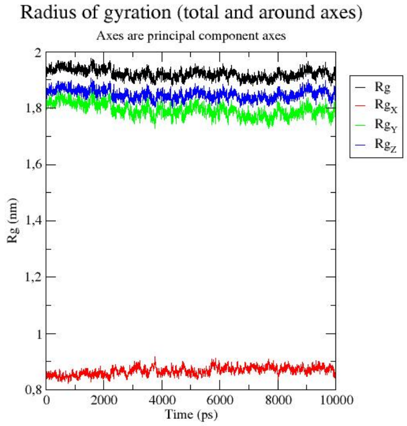

A) 2000 bar

B) 2250 bar

Figure 37: Rg side-by-side - High Hydrostatic Pressure

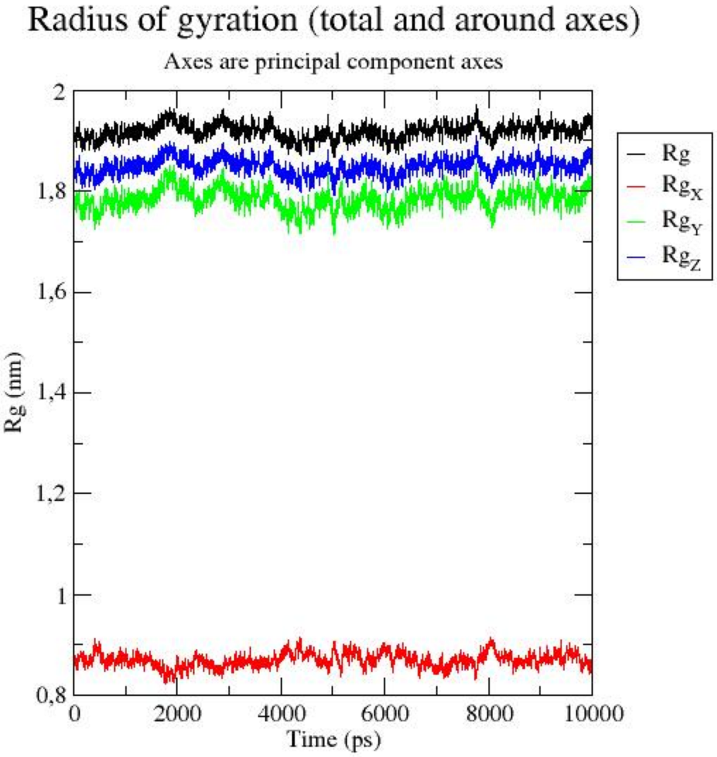

A) 2500 bar

Figure 38: Rg side-by-side - High Hydrostatic Pressure

##### HHP and low temperature

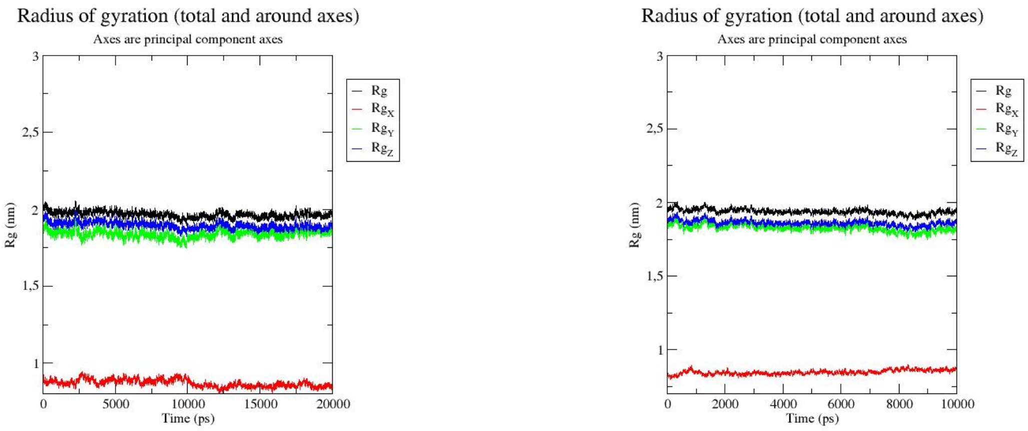

A) 1 bar B) 250 bar

Figure 39: Rg side-by-side - High Hydrostatic Pressure and Low Temperature

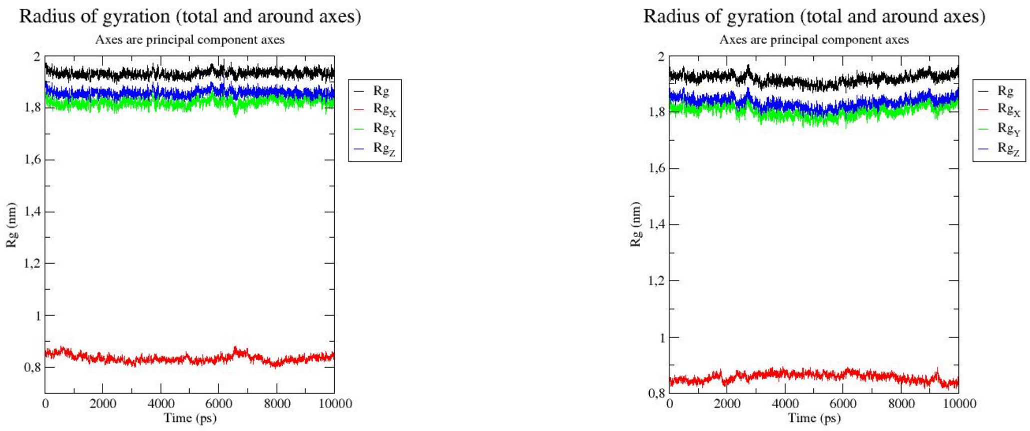

A) 500 bar

B) 750 bar

Figure 40: Rg side-by-side - High Hydrostatic Pressure and Low Temperature

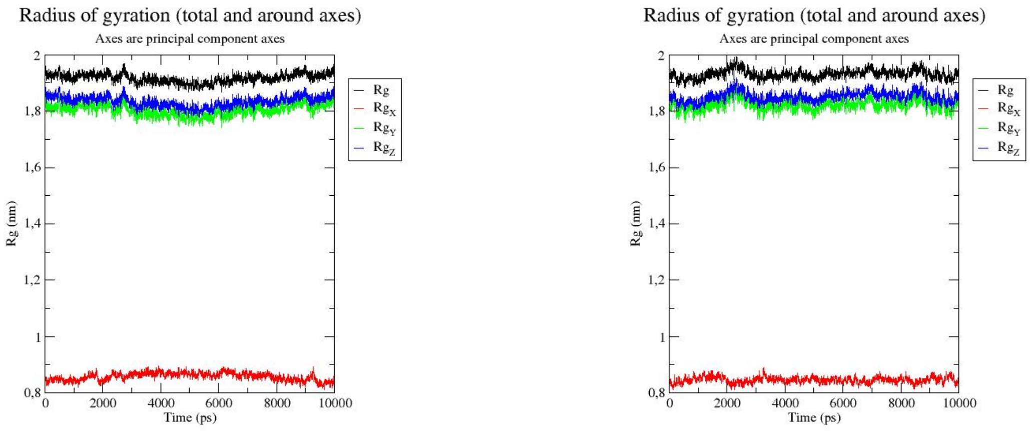

A) 1000 bar

B) 1250 bar

Figure 41: Rg side-by-side - High Hydrostatic Pressure and Low Temperature

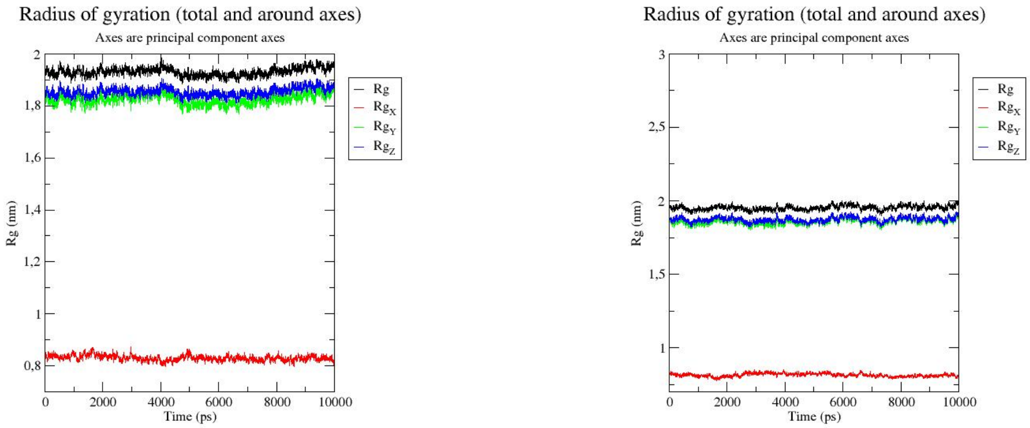

A) 1500 bar B) 1750 bar

Figure 42: Rg side-by-side - High Hydrostatic Pressure and Low Temperature

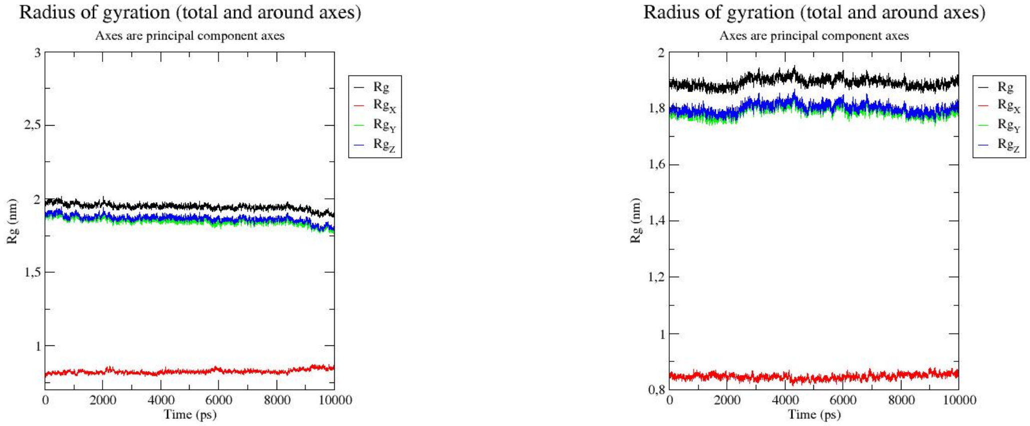

A) 2000 bar B) 2250 bar

Figure 43: Rg side-by-side - High Hydrostatic Pressure and Low Temperature

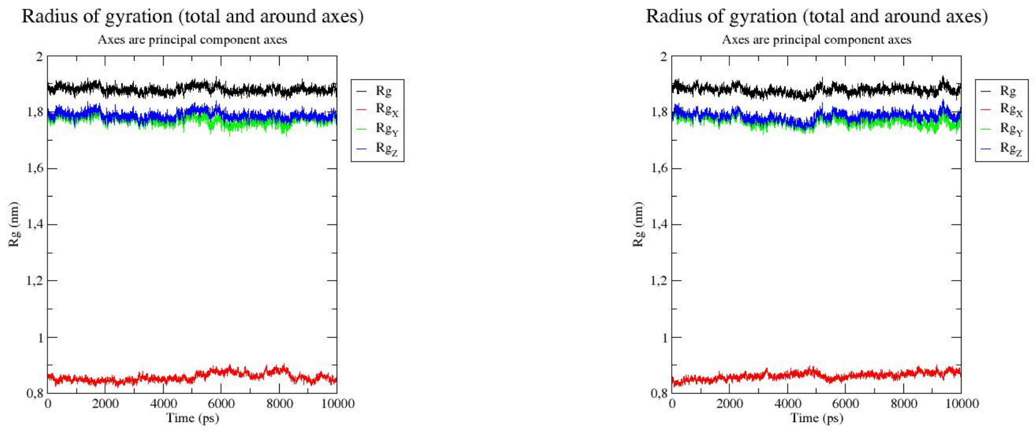

A) 2500 bar @ 300 K B) 2500 bar @ 295 K

Figure 44: Rg side-by-side - High Hydrostatic Pressure and Low Temperature

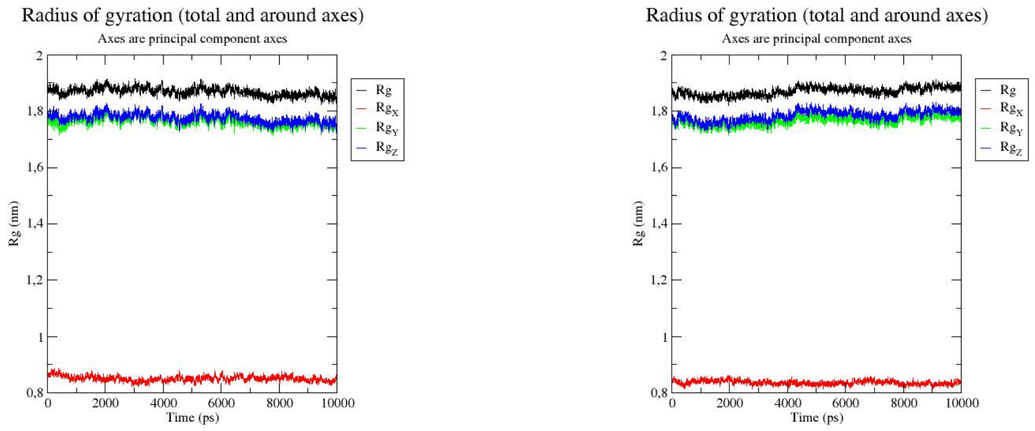

A) 2500 bar @ 290 K B) 2500 bar @ 285 K

Figure 45: Rg side-by-side - High Hydrostatic Pressure and Low Temperature

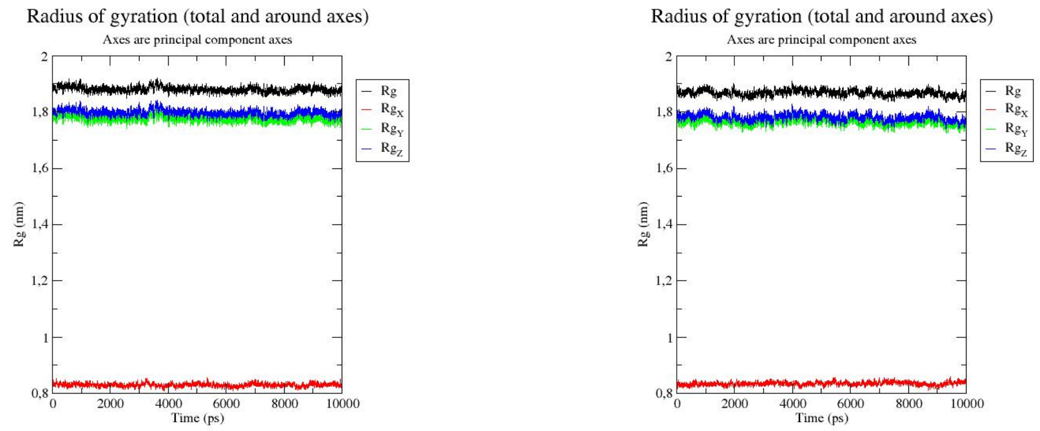

A) 2500 bar @ 280 K B) 2500 bar @ 275 K

Figure 46: Rg side-by-side - High Hydrostatic Pressure and Low Temperature

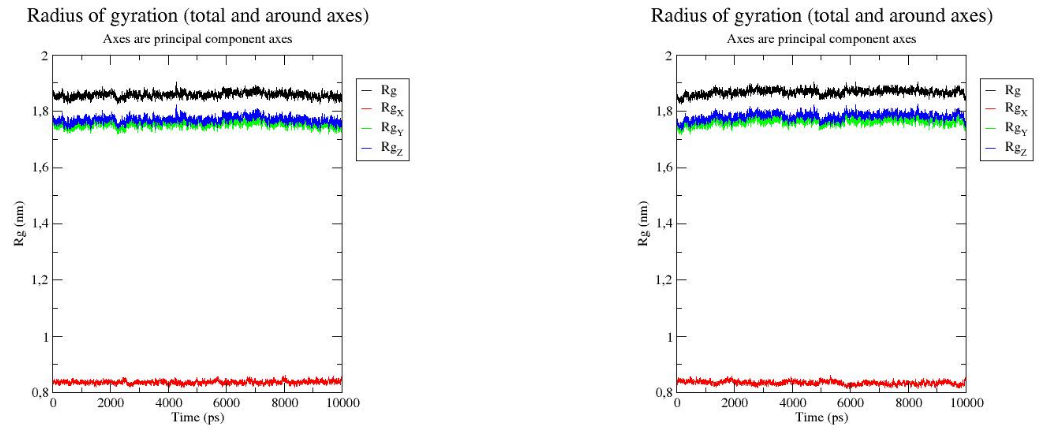

A) 2500 bar @ 270 K B) 2500 bar @ 265 K

Figure 47: Rg side-by-side - High Hydrostatic Pressure and Low Temperature

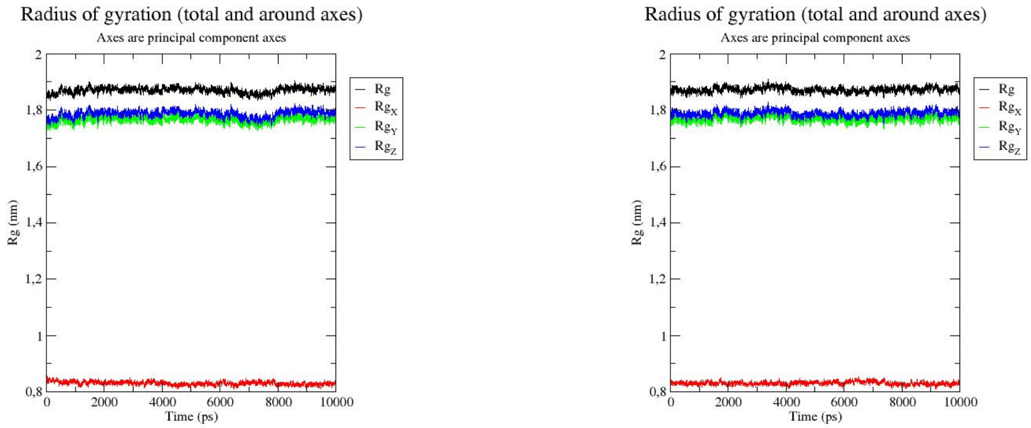

A) 2500 bar @ 260 K    B) 2500 bar @ 255 K

Figure 48: Rg side-by-side - High Hydrostatic Pressure and Low Temperature

#### SASA

##### HHP

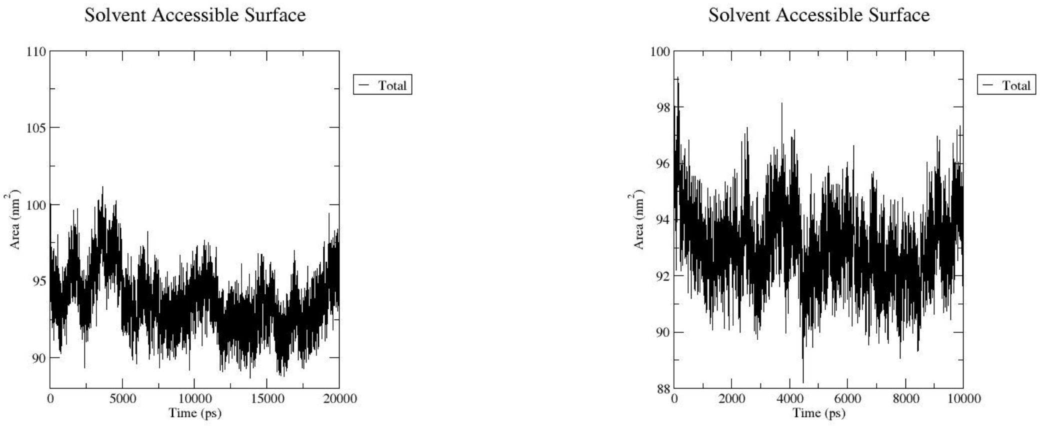

A) 1 bar B) 250 bar

Figure 49: SASA side-by-side - High Hydrostatic Pressure

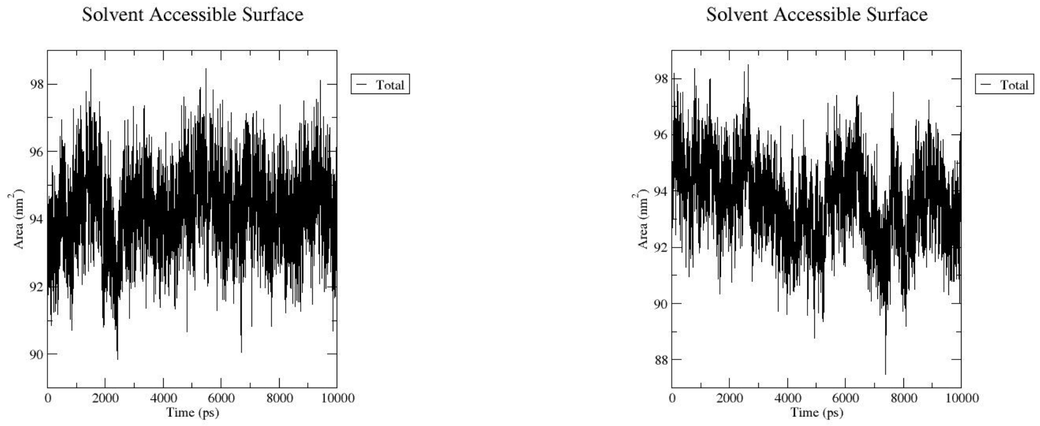

A) 500 bar B) 750 bar

Figure 50: SASA side-by-side - High Hydrostatic Pressure

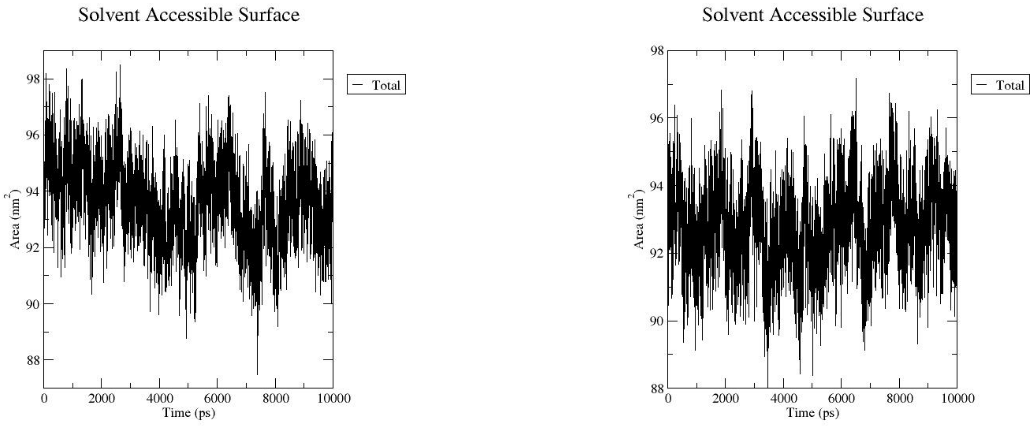

A) 1000 bar B) 1250 bar

Figure 51: SASA side-by-side - High Hydrostatic Pressure

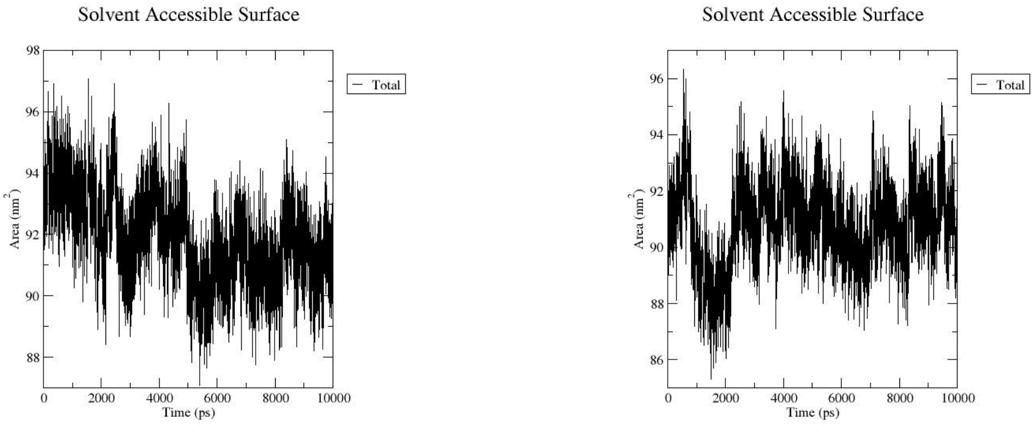

A) 1500 bar B) 1750 bar

Figure 52: SASA side-by-side - High Hydrostatic Pressure

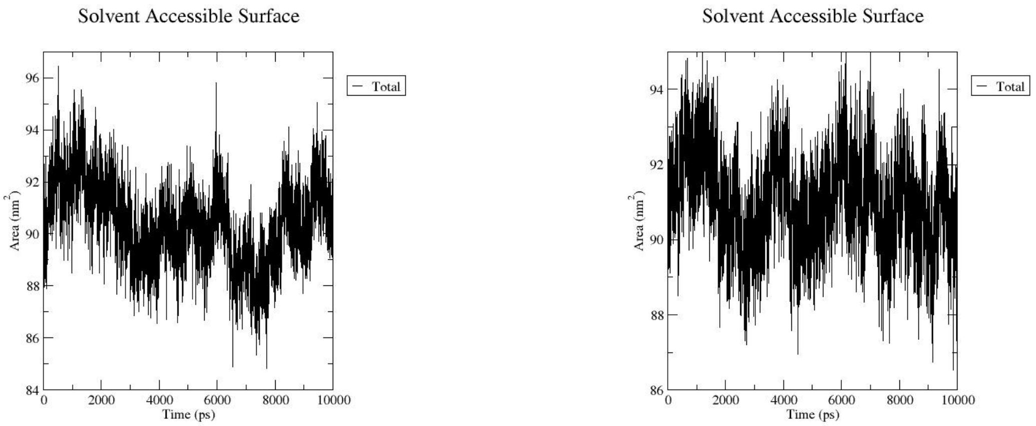

A) 2000 bar B) 2250 bar

Figure 53: SASA side-by-side - High Hydrostatic Pressure

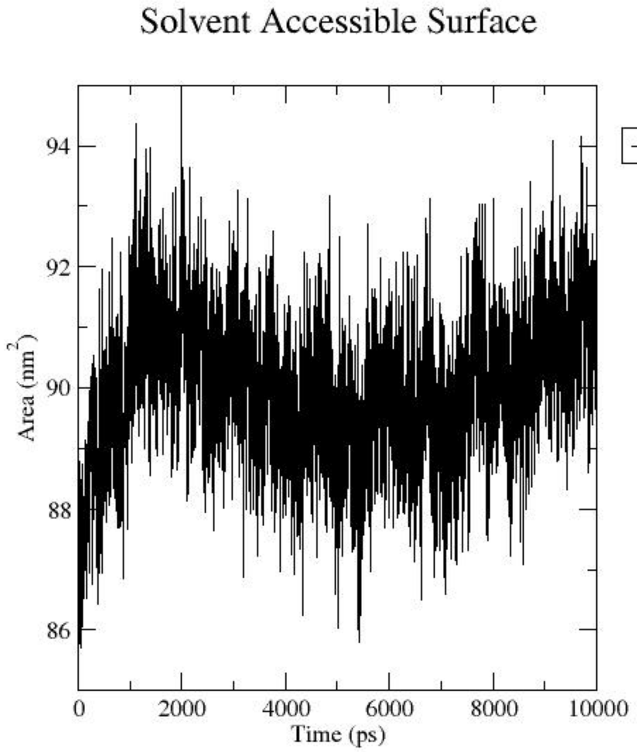

A) 2500 bar

Figure 54: SASA side-by-side - High Hydrostatic Pressure

**HHP and low temperature**

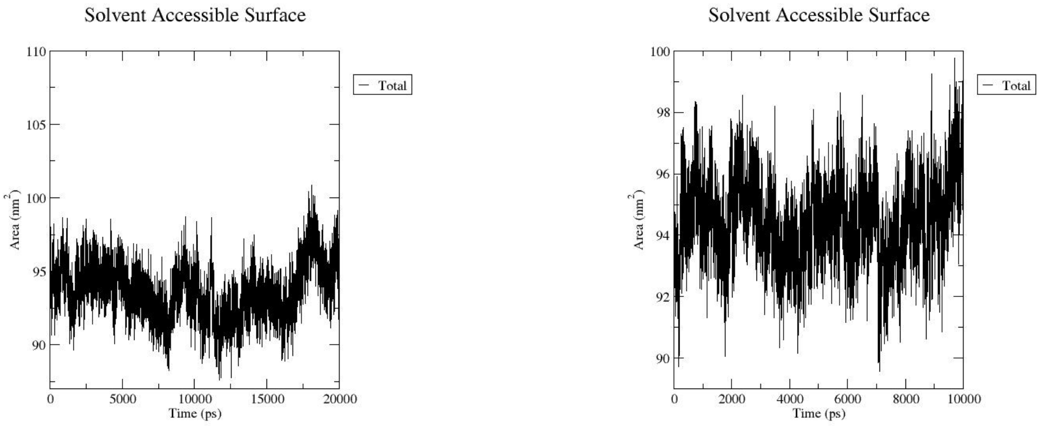

A) 1 bar B) 250 bar

Figure 55: SASA side-by-side - High Hydrostatic Pressure and Low Temperature

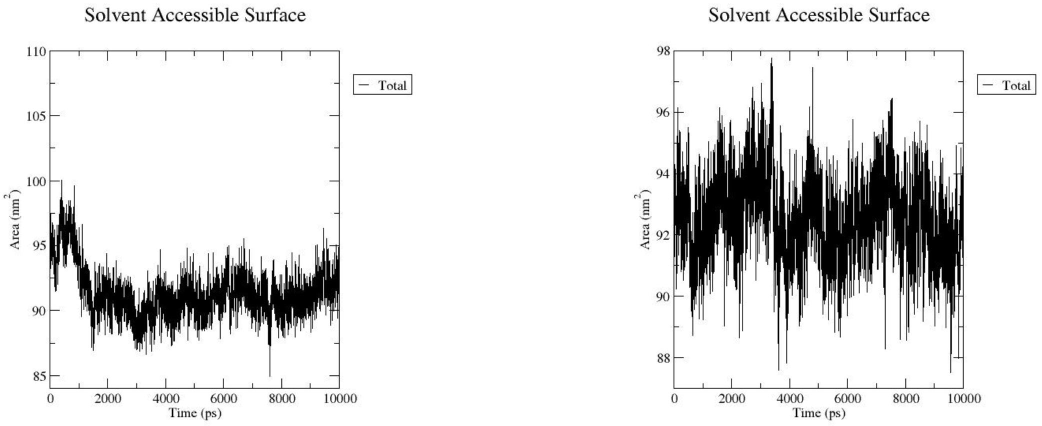

A) 500 bar B) 750 bar

Figure 56: SASA side-by-side - High Hydrostatic Pressure and Low Temperature

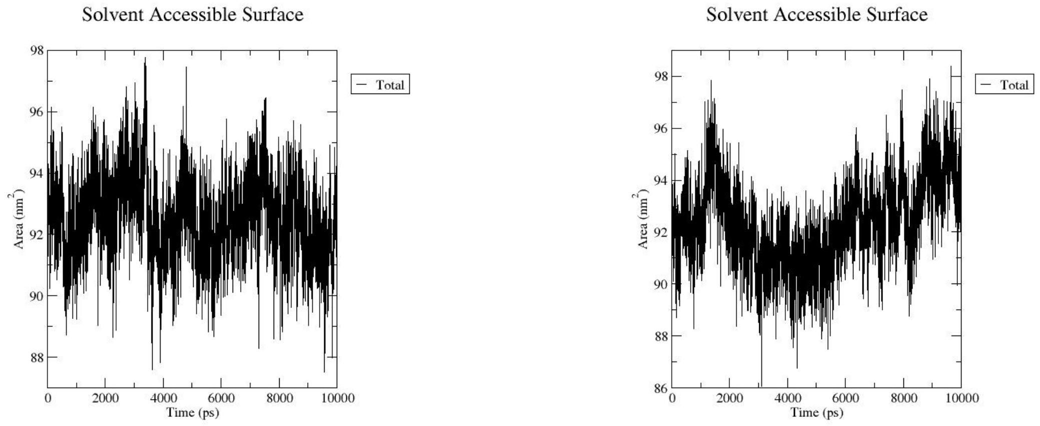

A) 1000 bar B) 1250 bar

Figure 57: SASA side-by-side - High Hydrostatic Pressure and Low Temperature

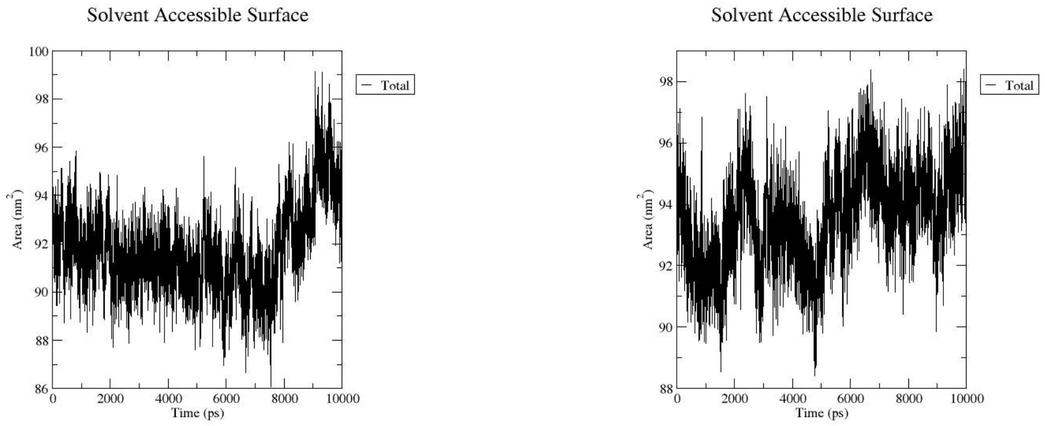

A) 1500 bar B) 1750 bar

Figure 58: SASA side-by-side - High Hydrostatic Pressure and Low Temperature

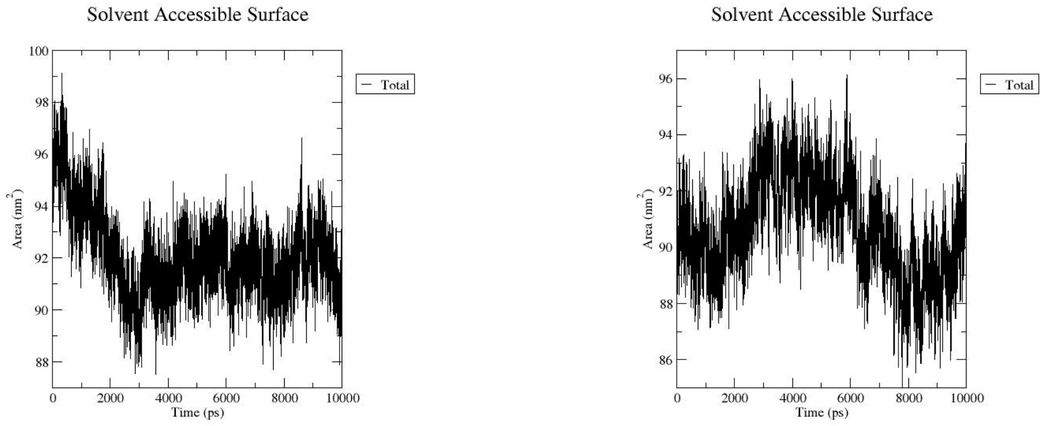

A) 2000 bar B) 2250 bar

Figure 59: SASA side-by-side - High Hydrostatic Pressure and Low Temperature

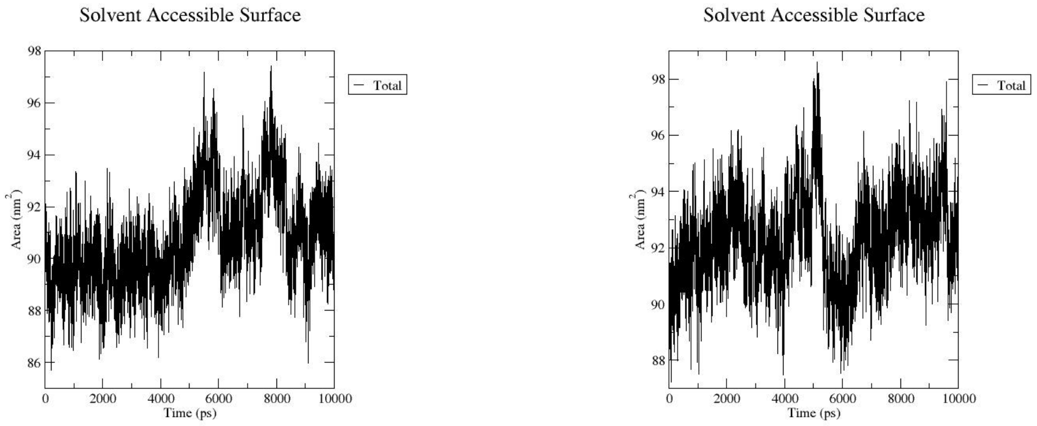

A) 2500 bar @ 300 K B) 2500 bar @ 295 K

Figure 60: SASA side-by-side - High Hydrostatic Pressure and Low Temperature

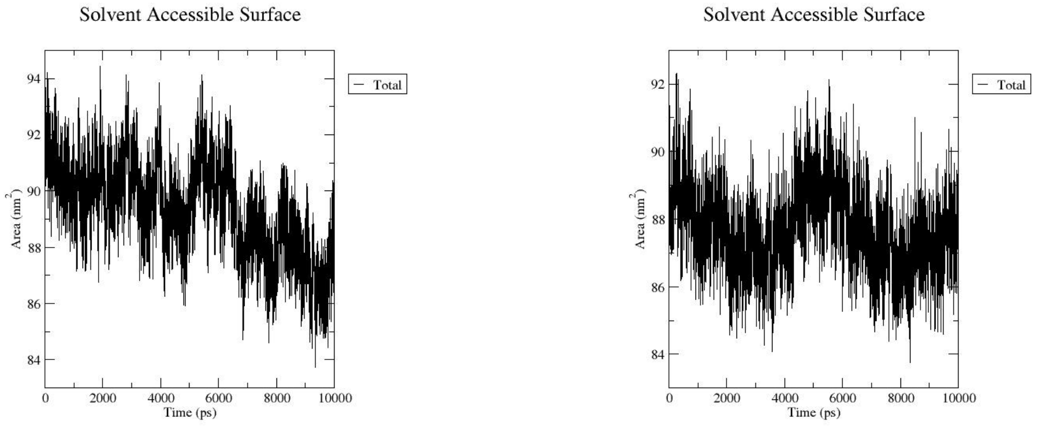

A) 2500 bar @ 290 K B) 2500 bar @ 285 K

Figure 61: SASA side-by-side - High Hydrostatic Pressure and Low Temperature

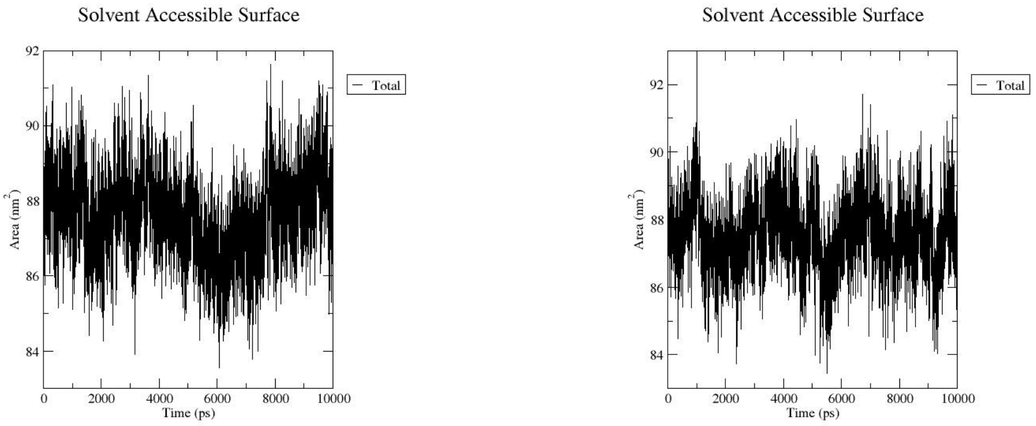

A) 2500 bar @ 280 K B) 2500 bar @ 275 K

Figure 62: SASA side-by-side - High Hydrostatic Pressure and Low Temperature

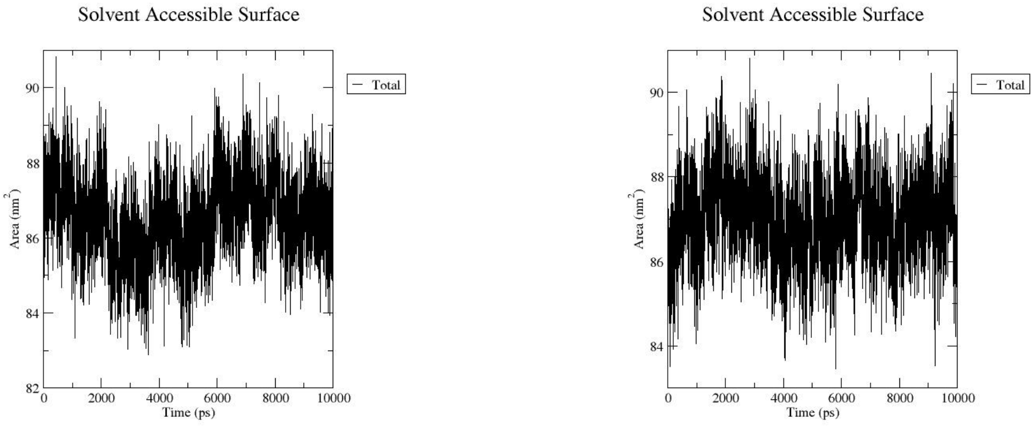

A) 2500 bar @ 270 K B) 2500 bar @ 265 K

Figure 63: SASA side-by-side - High Hydrostatic Pressure and Low Temperature

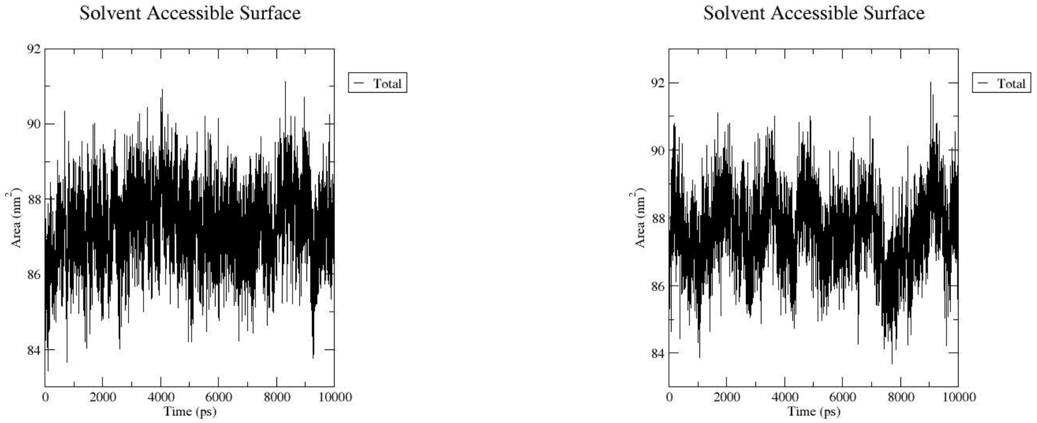

A) 2500 bar @ 260 K

B) 2500 bar @ 255 K

Figure 64: SASA side-by-side - High Hydrostatic Pressure and Low Temperature

#### Hbonds

##### HHP

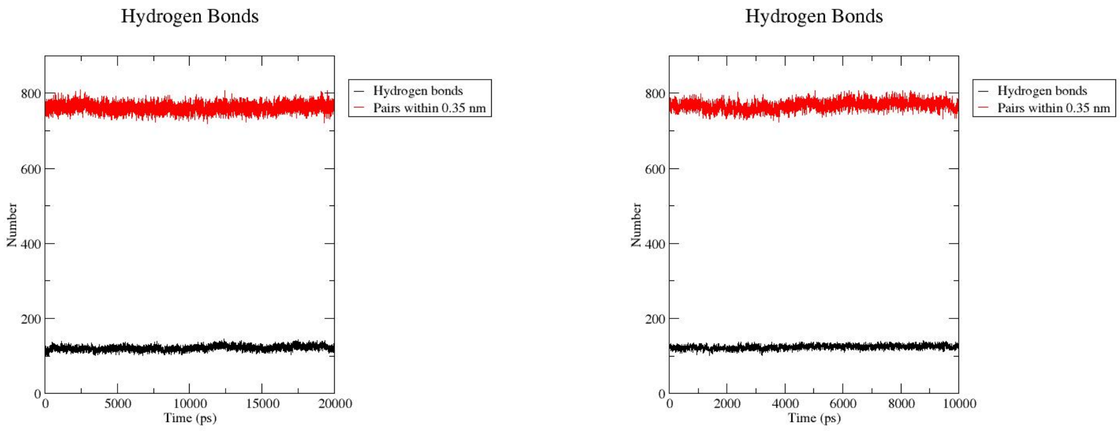

A) 1 bar

B) 250 bar

Figure 65: Hbond side-by-side - High Hydrostatic Pressure

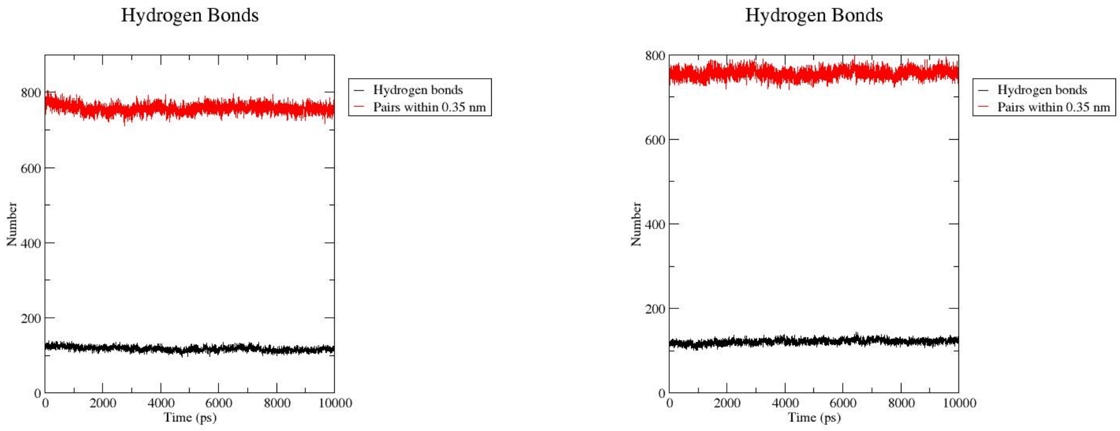

A) 500 bar

B) 750 bar

Figure 66: Hbond side-by-side - High Hydrostatic Pressure

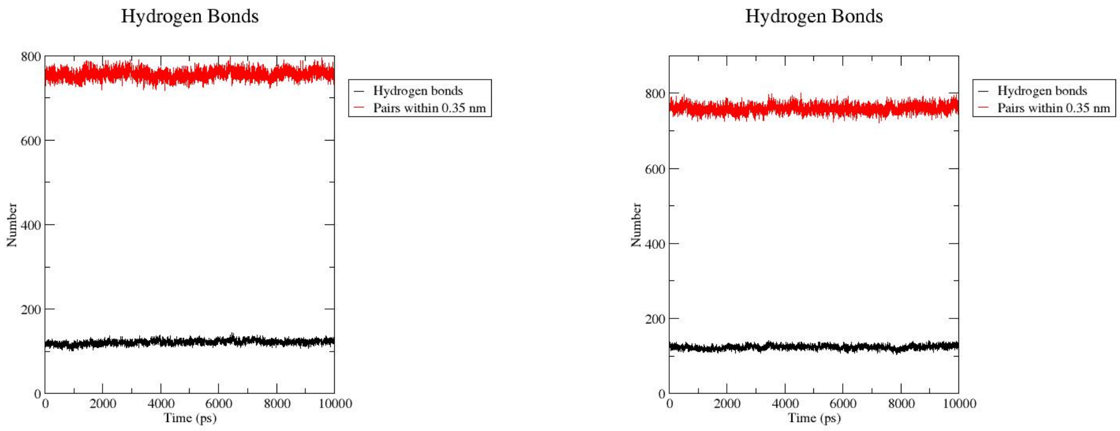

A) 1000 bar B) 1250 bar

Figure 67: Hbond side-by-side - High Hydrostatic Pressure

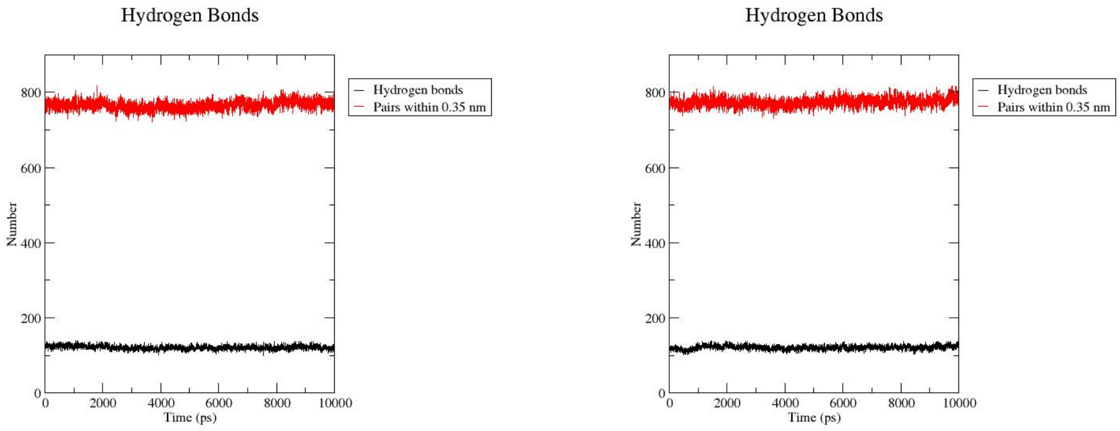

A) 1500 bar B) 1750 bar

Figure 68: Hbond side-by-side - High Hydrostatic Pressure

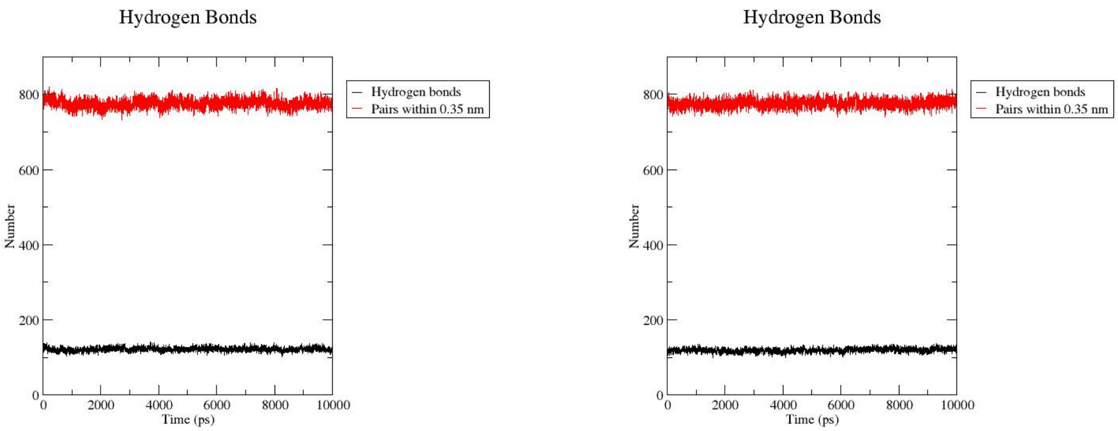

A) 2000 bar B) 2250 bar

Figure 69: Hbond side-by-side - High Hydrostatic Pressure

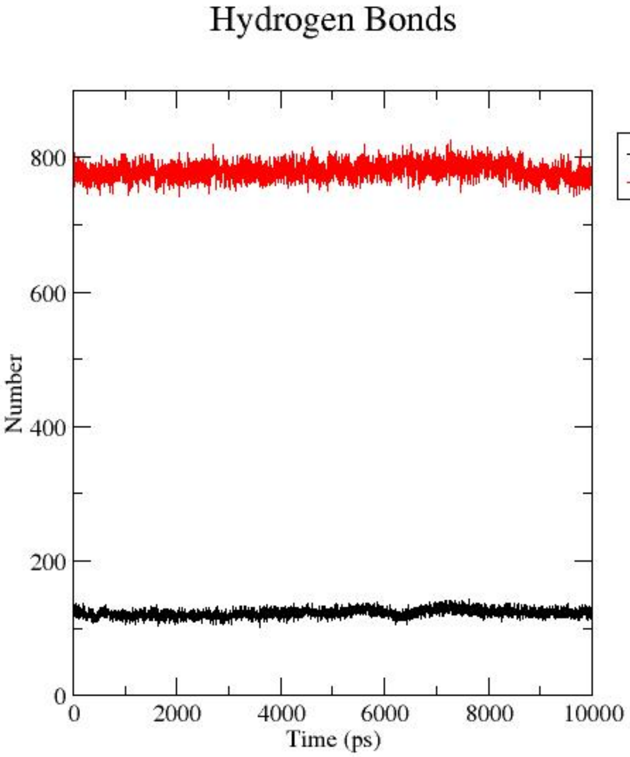

A) 2500 bar

Figure 70: Hbond side-by-side - High Hydrostatic Pressure

**HHP and low temperature**

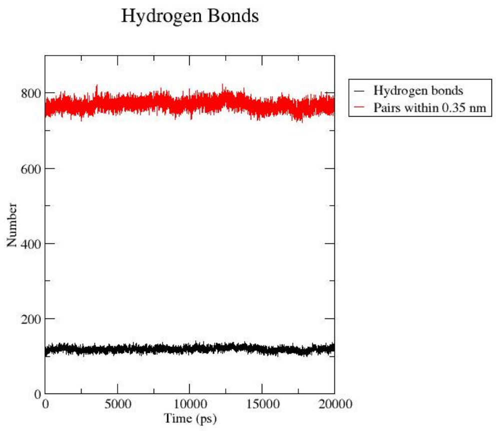

A) 1 bar B) 250 bar

Figure 71: HBond side-by-side - High Hydrostatic Pressure and Low Temperature

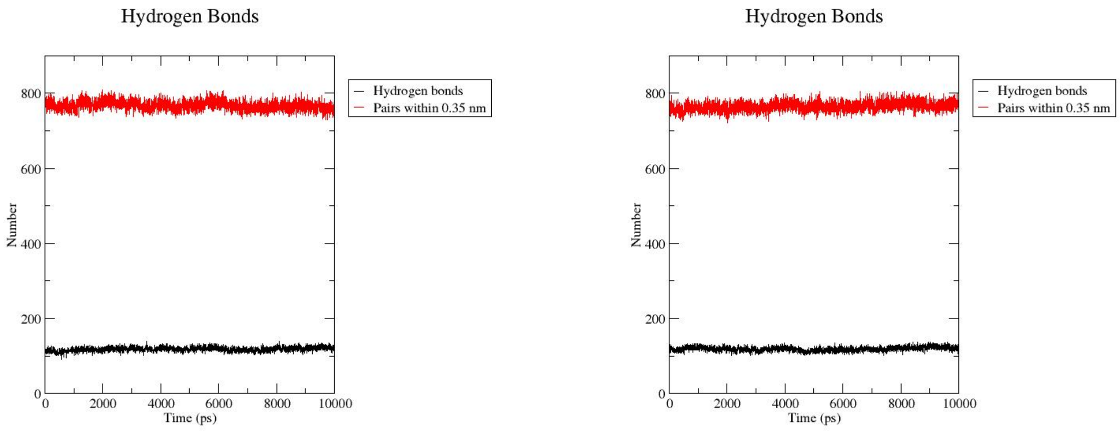

A) 500 bar B) 750 bar

Figure 72: HBond side-by-side - High Hydrostatic Pressure and Low Temperature

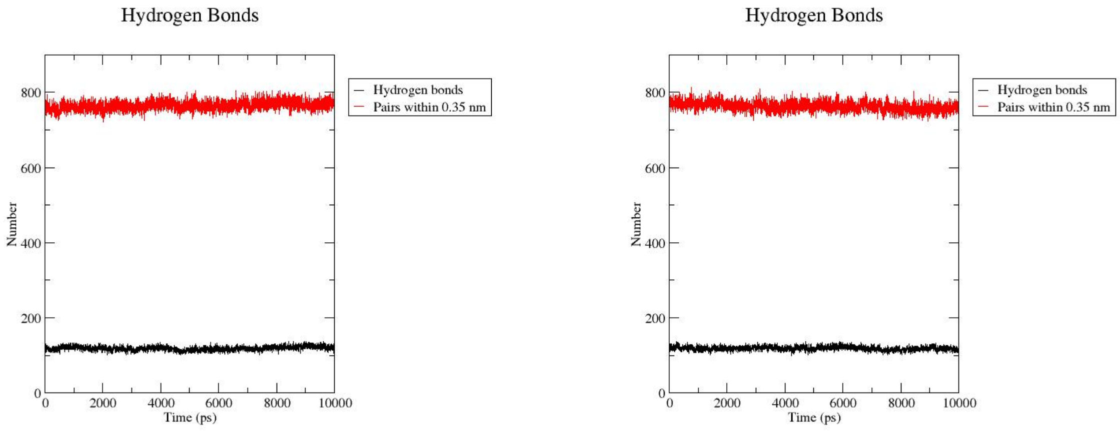

A) 1000 bar B) 1250 bar

Figure 73: HBond side-by-side - High Hydrostatic Pressure and Low Temperature

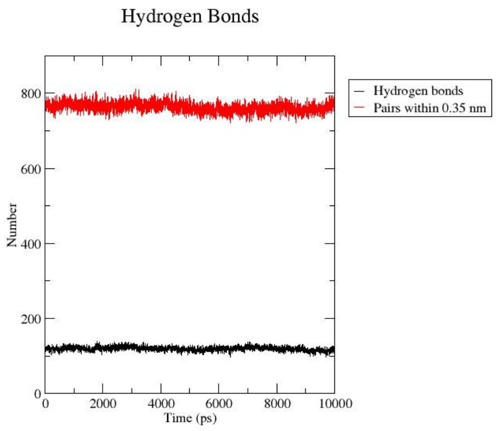

A) 1500 bar

B) 1750 bar

Figure 74: HBond side-by-side - High Hydrostatic Pressure and Low Temperature

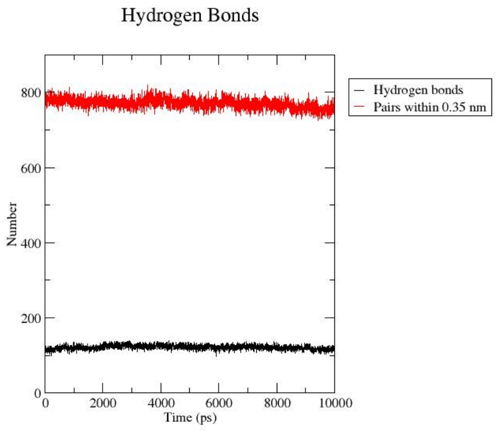

A) 2000 bar

B) 2250 bar

Figure 75: HBond side-by-side - High Hydrostatic Pressure and Low Temperature

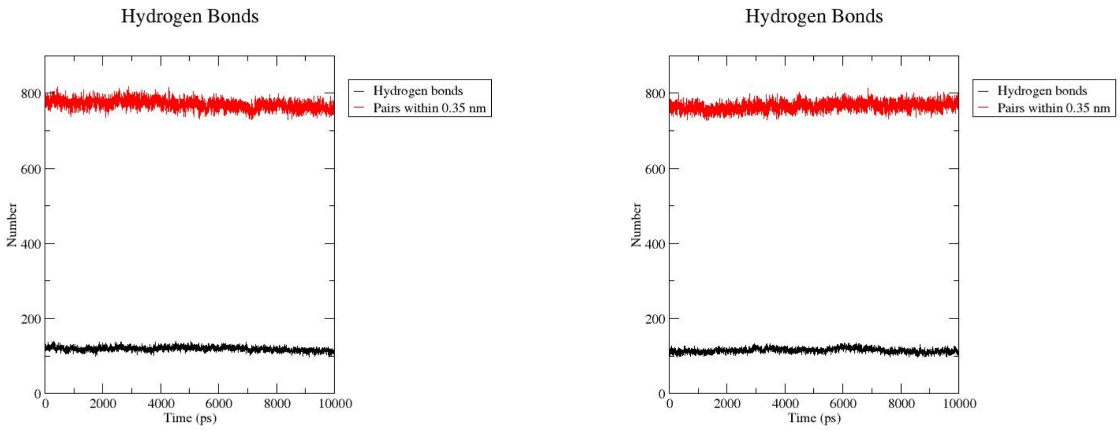

A) 2500 bar @ 300 K B) 2500 bar @ 295 K

Figure 76: HBond side-by-side - High Hydrostatic Pressure and Low Temperature

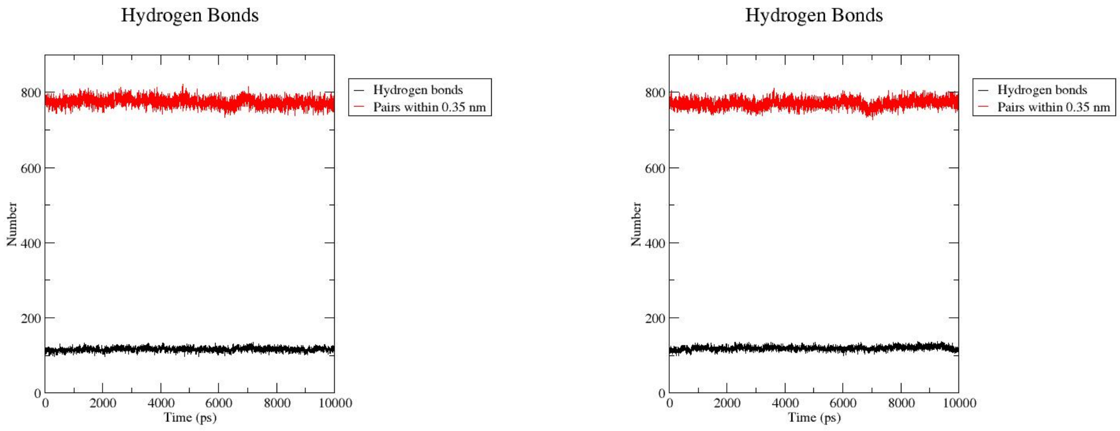

A) 2500 bar @ 290 K B) 2500 bar @ 285 K

Figure 77: HBond side-by-side - High Hydrostatic Pressure and Low Temperature

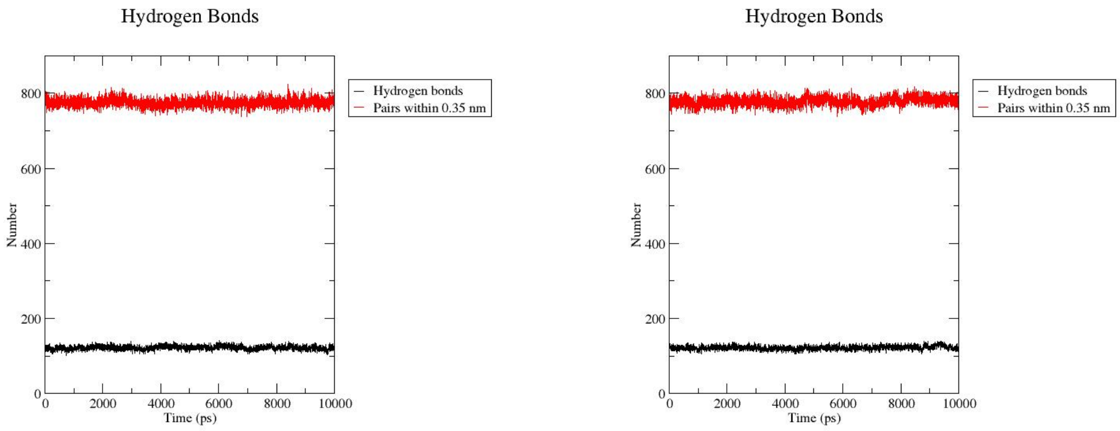

A) 2500 bar @ 280 K B) 2500 bar @ 275 K

Figure 78: HBond side-by-side - High Hydrostatic Pressure and Low Temperature

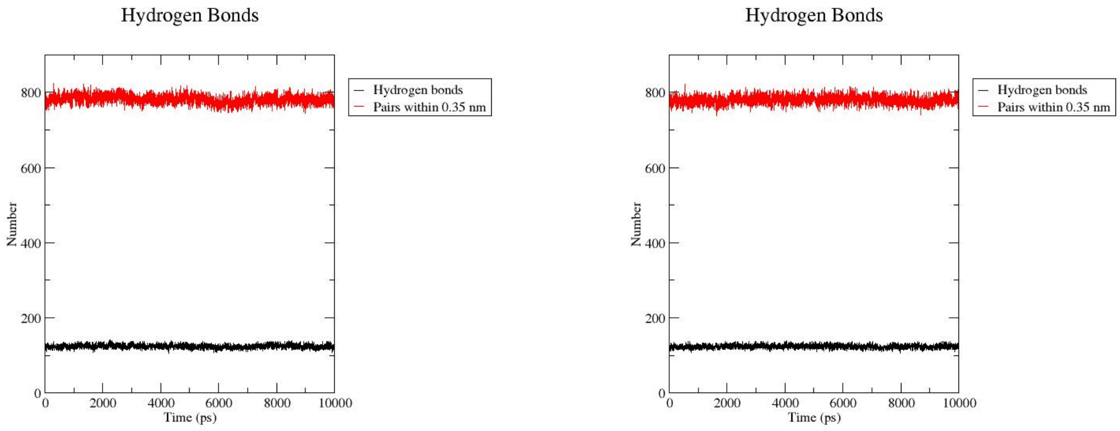

A) 2500 bar @ 270 K B) 2500 bar @ 265 K

Figure 79: HBond side-by-side - High Hydrostatic Pressure and Low Temperature

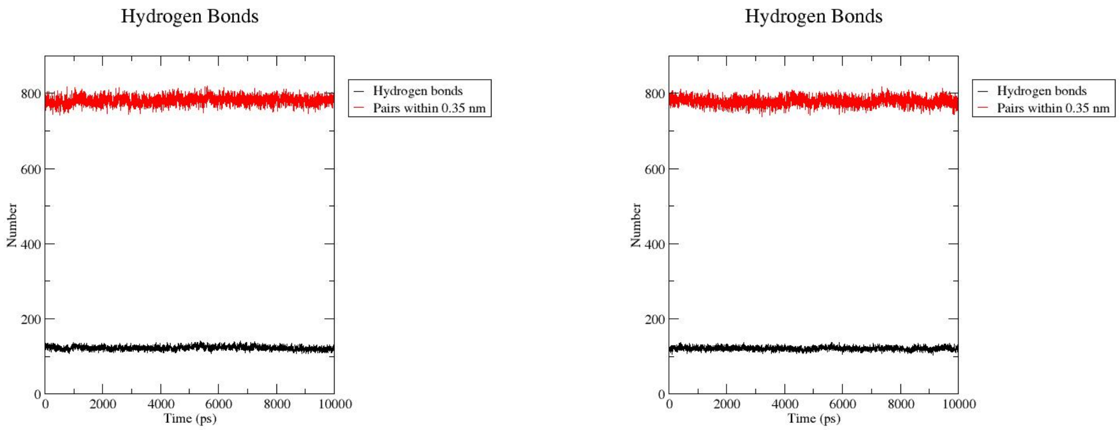

A) 2500 bar @ 260 K    B) 2500 bar @ 255 K

Figure 80: HBond side-by-side - High Hydrostatic Pressure and Low Temperature

#### Hierarchical Clustering

##### HHP

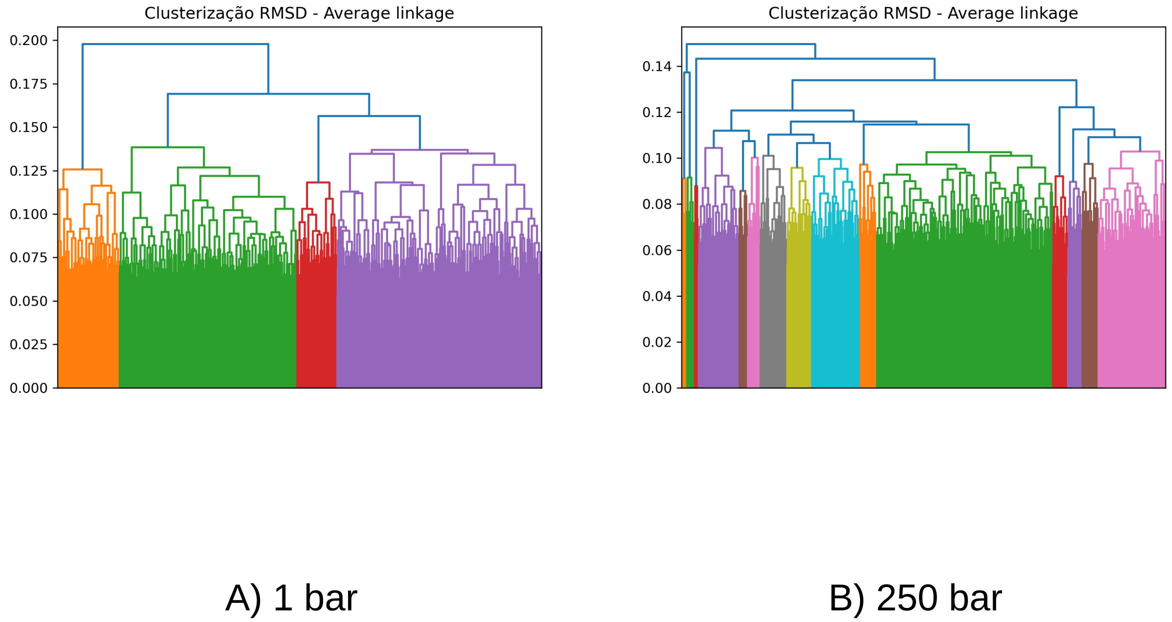

Figure 81: Hierachical Clustering side-by-side - High Hydrostatic Pressure

**HHP and low temperature**

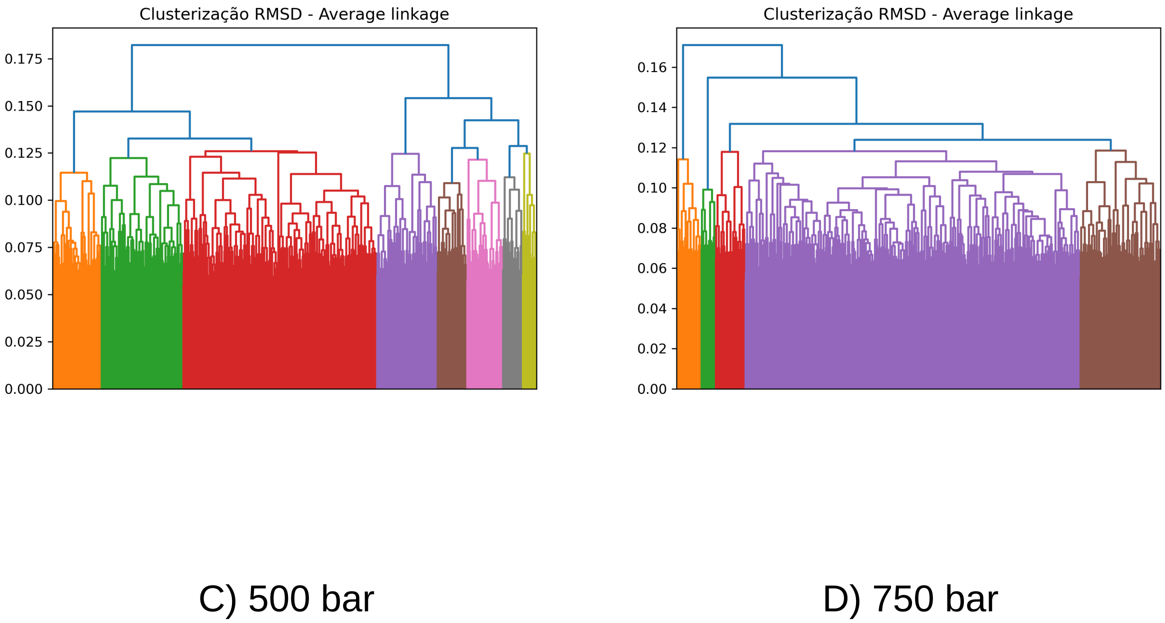

Figure 82: Hierachical Clustering side-by-side - High Hydrostatic Pressure

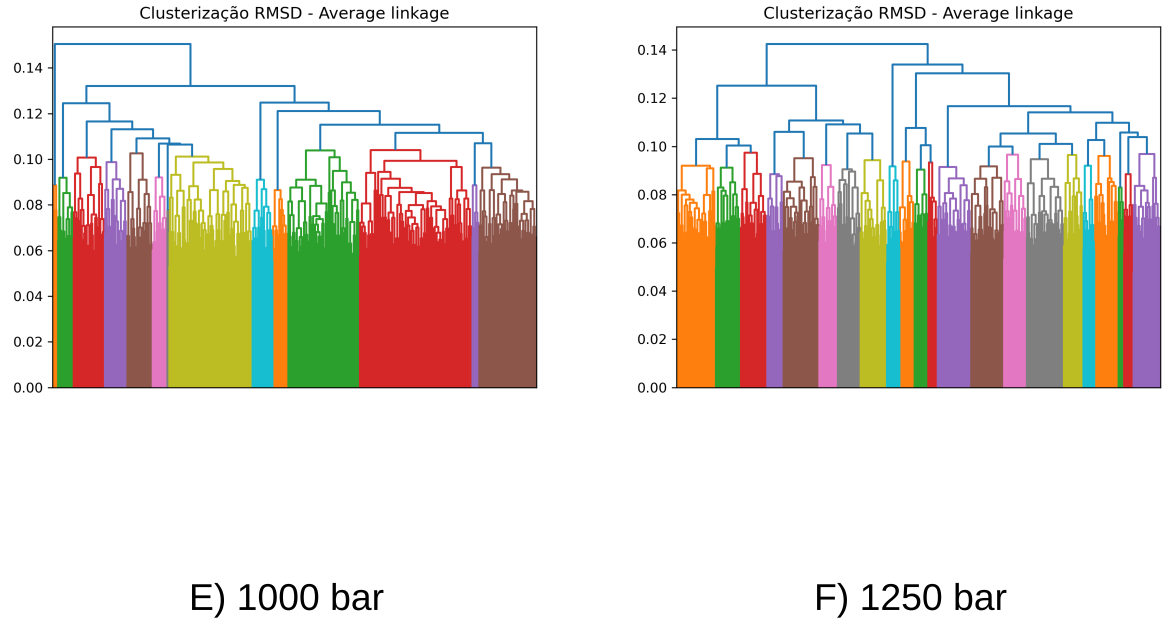

Figure 83: Hierachical Clustering side-by-side - High Hydrostatic Pressure

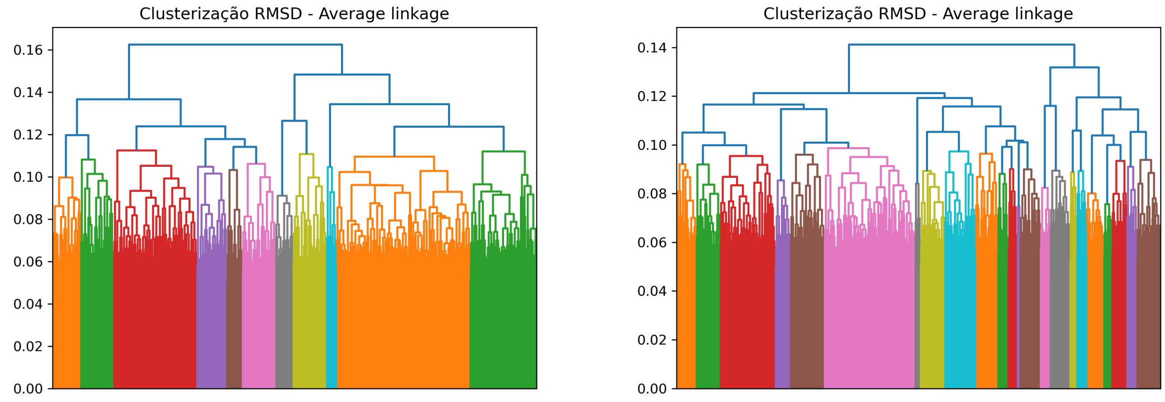

G) 1500 bar H) 1750 bar

Figure 84: Hierachical Clustering side-by-side - High Hydrostatic Pressure

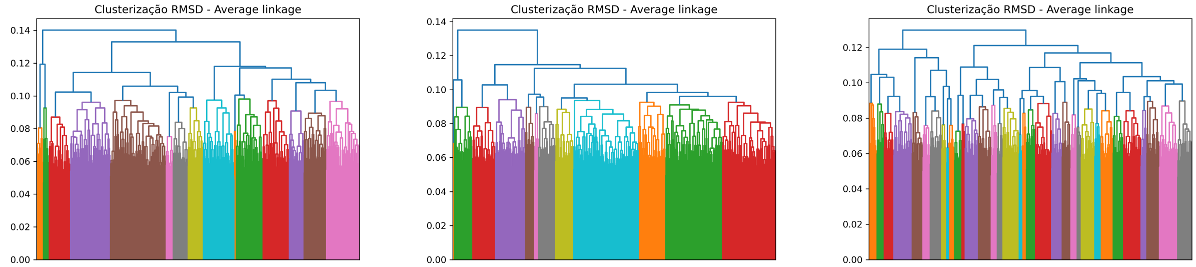

I) 2000 bar J) 2250 bar K) 2500 bar

Figure 85: Hierachical Clustering side-by-side - High Hydrostatic Pressure

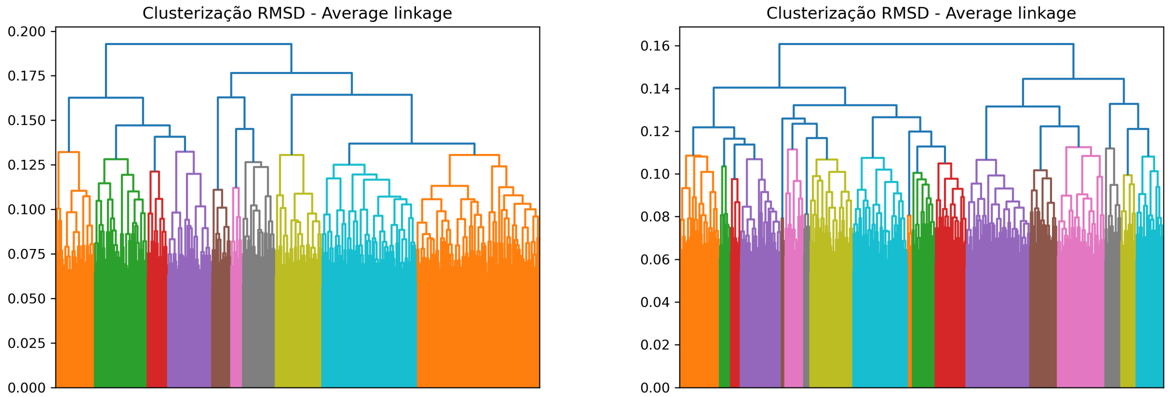

A) 1 bar

B) 250 bar

Figure 86: Hierachical Clustering side-by-side - High Hydrostatic Pressure and Low Temperature

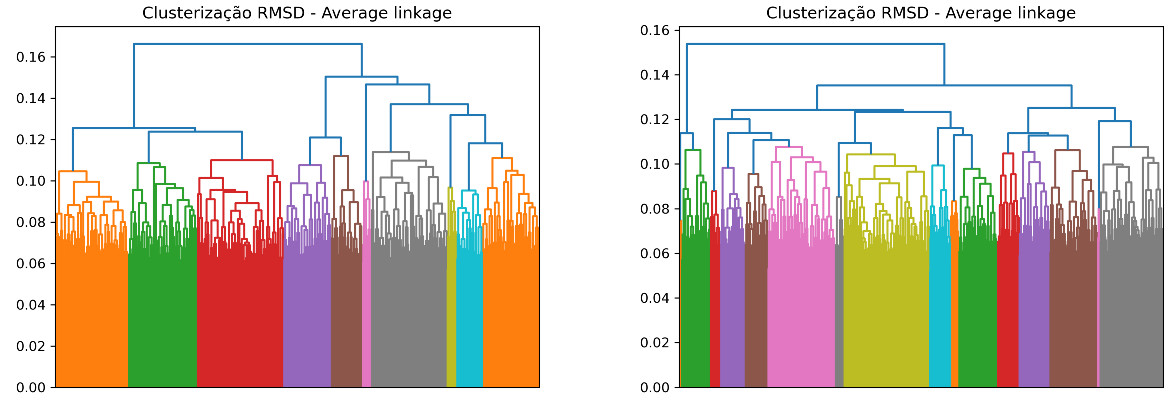

C) 500 bar

D) 750 bar

Figure 87: Hierachical Clustering side-by-side - High Hydrostatic Pressure and Low Temperature

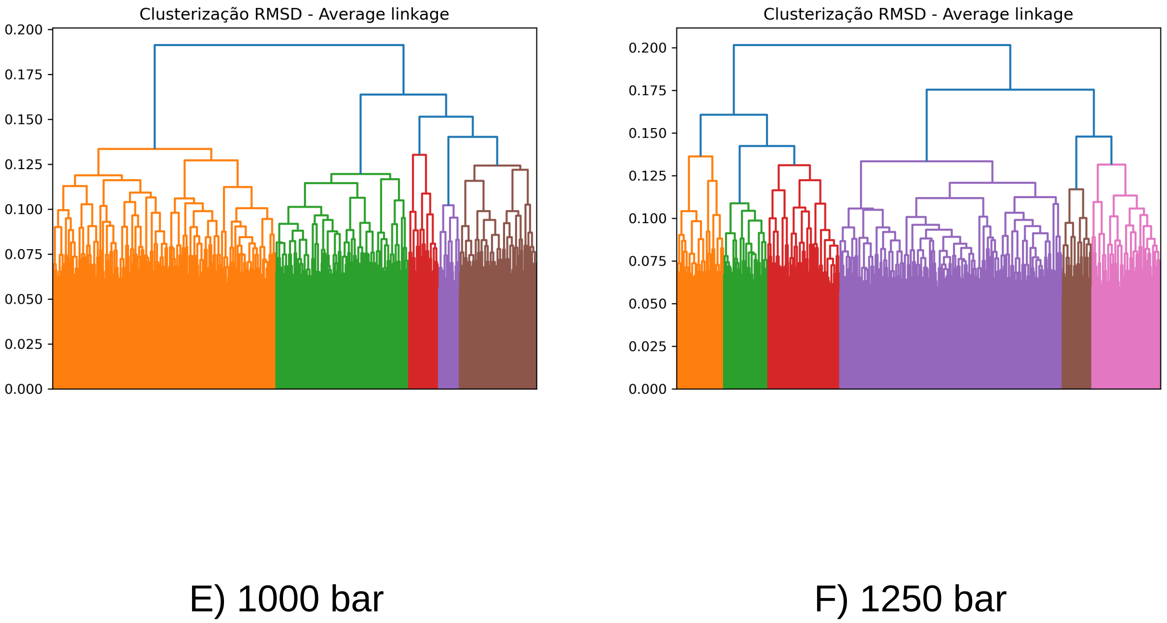

Figure 88: Hierachical Clustering side-by-side - High Hydrostatic Pressure and Low Temperature

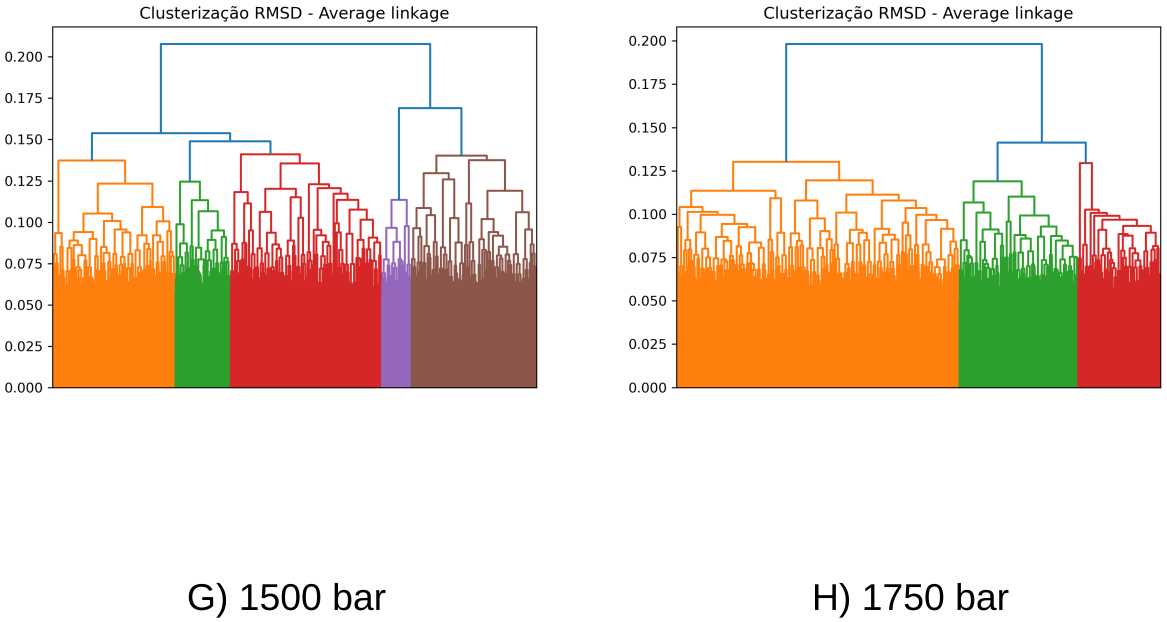

Figure 89: Hierachical Clustering side-by-side - High Hydrostatic Pressure and Low Temperature

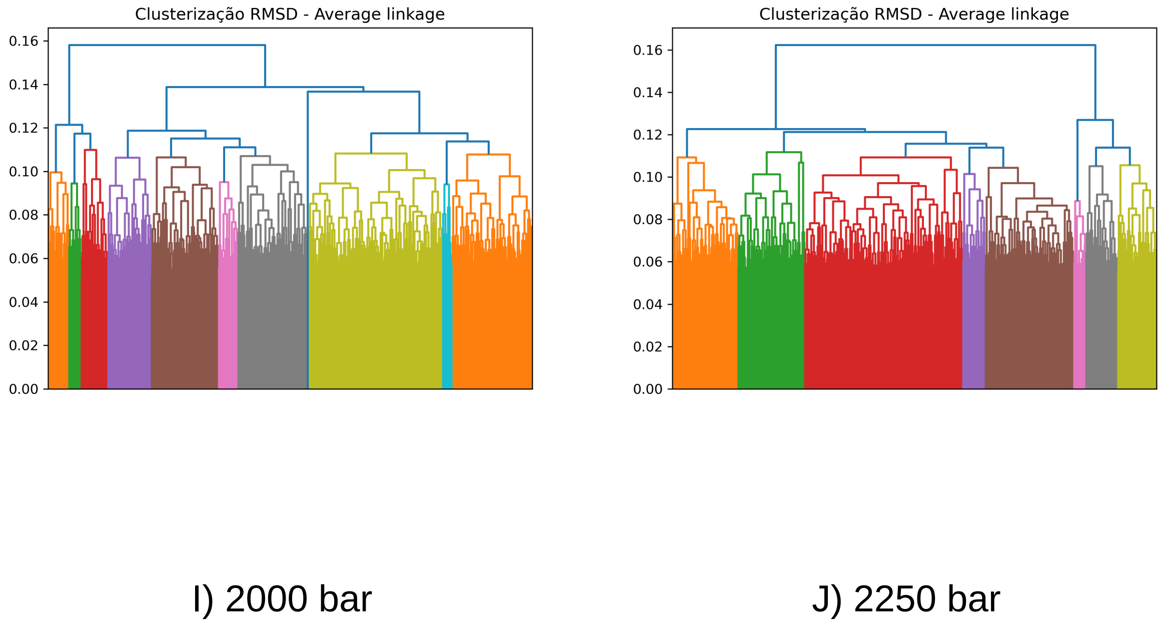

Figure 90: Hierachical Clustering side-by-side - High Hydrostatic Pressure and Low Temperature

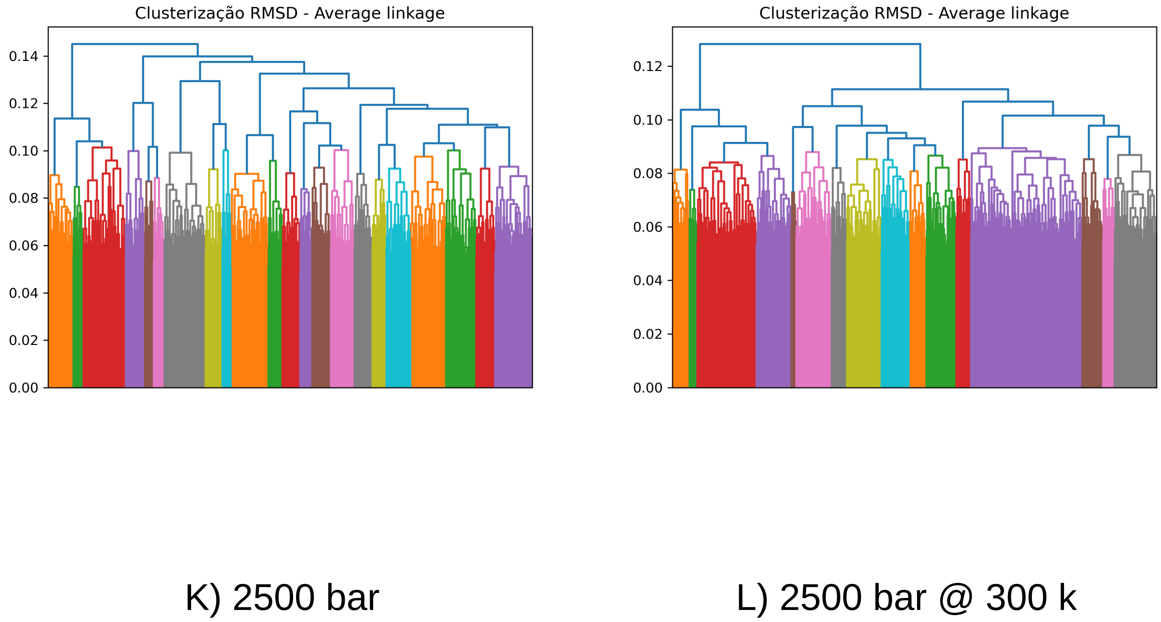

Figure 91: Hierachical Clustering side-by-side - High Hydrostatic Pressure and Low Temperature

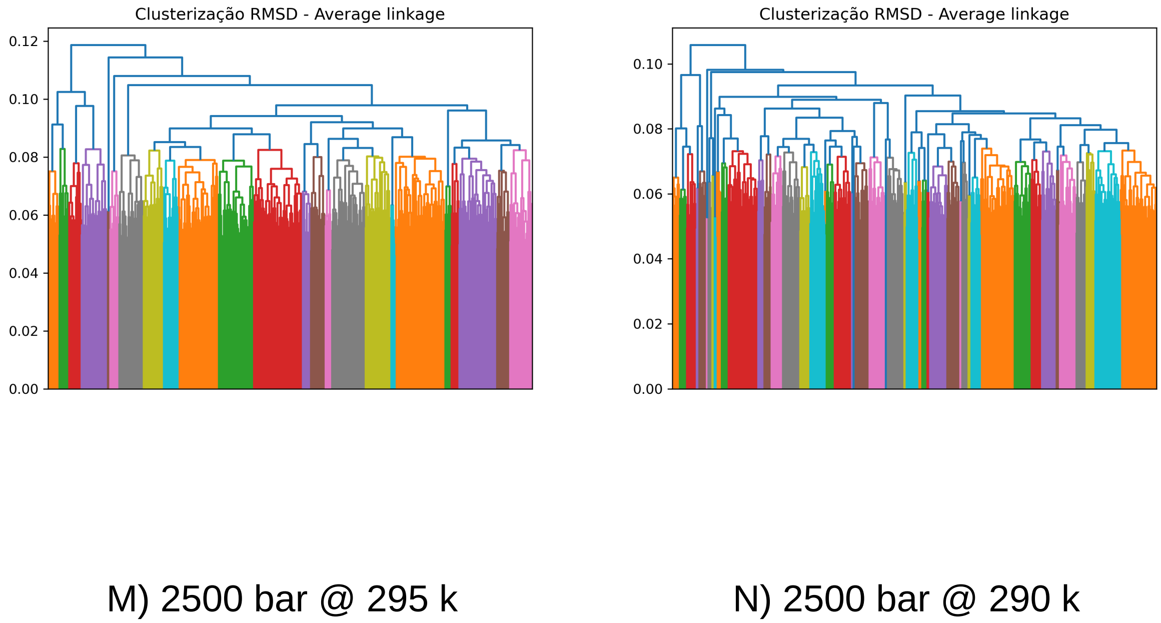

Figure 92: Hierachical Clustering side-by-side - High Hydrostatic Pressure and Low Temperature

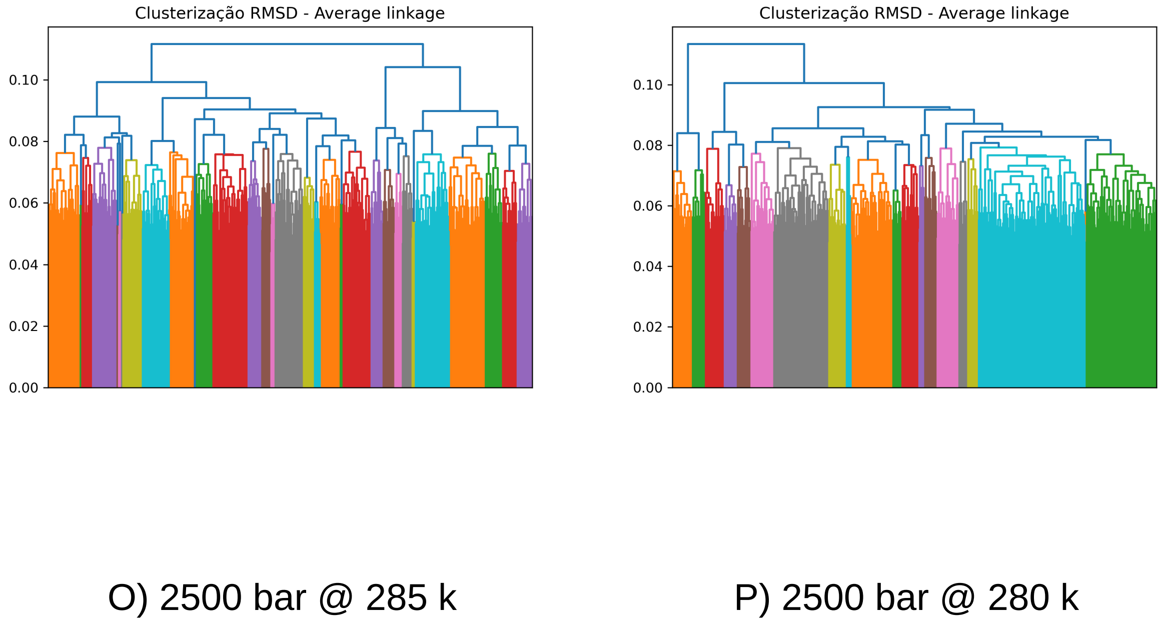

Figure 93: Hierachical Clustering side-by-side - High Hydrostatic Pressure and Low Temperature

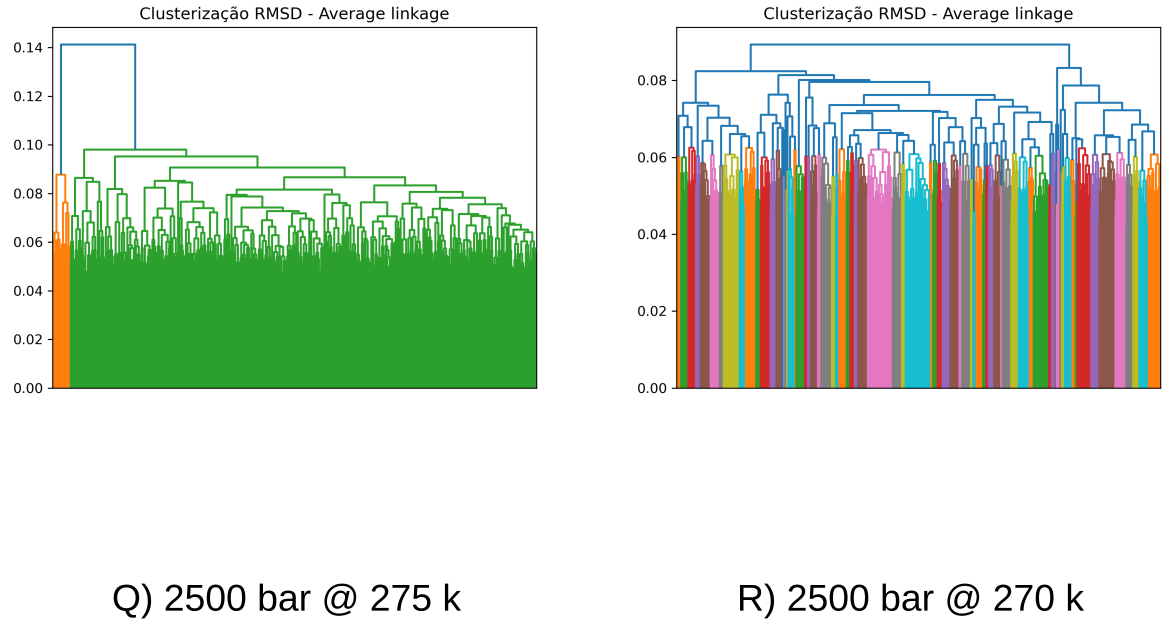

Figure 94: Hierachical Clustering side-by-side - High Hydrostatic Pressure and Low Temperature

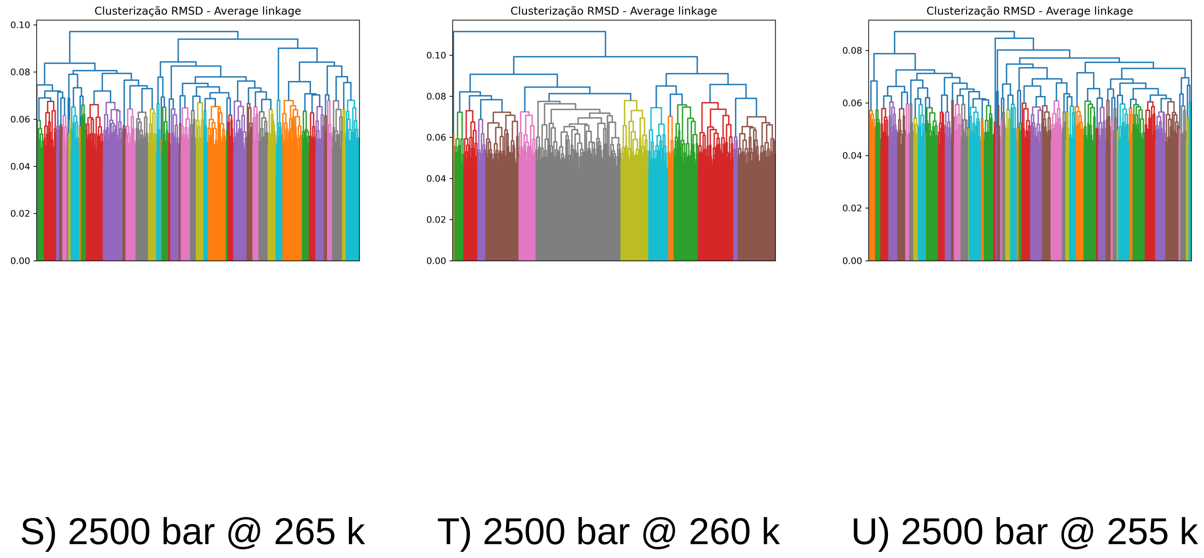

Figure 95: Hierachical Clustering side-by-side - High Hydrostatic Pressure and Low Temperature

### PCA

#### HHP

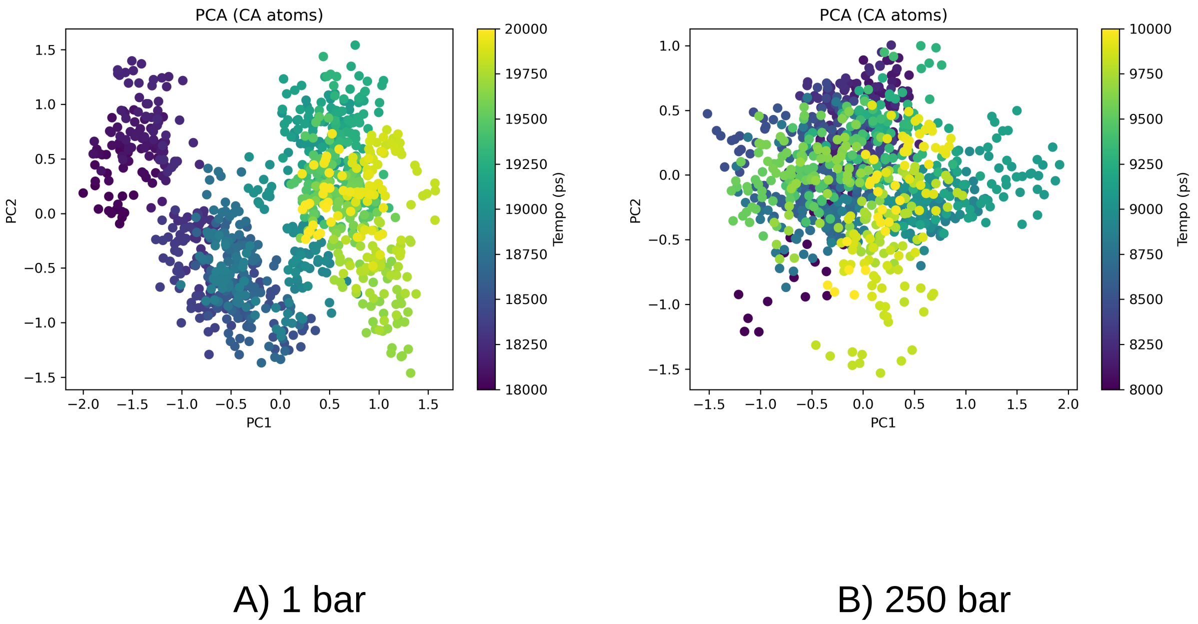

Figure 96: Principal Component Analysis side-by-side - High Hydrostatic Pressure

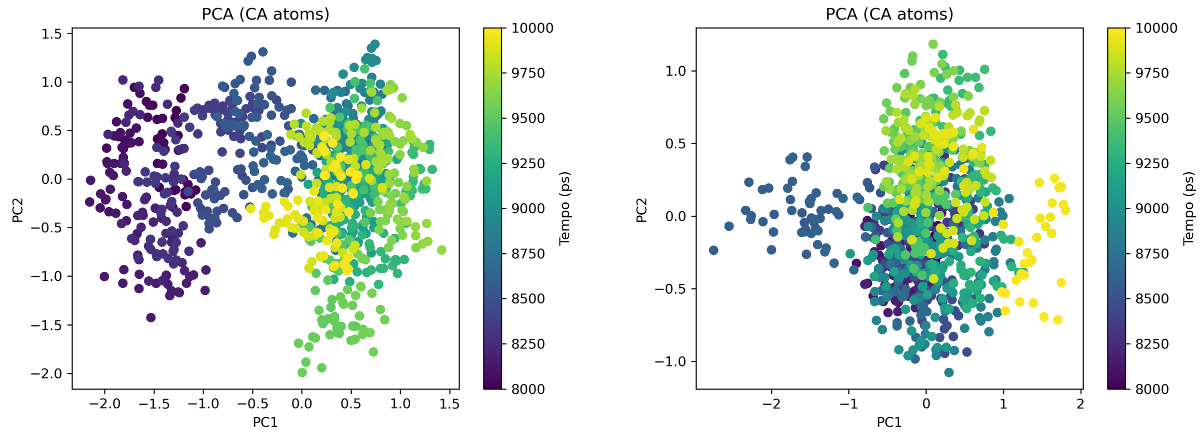

C) 500 bar D) 750 bar

Figure 97: Principal Component Analysis side-by-side - High Hydrostatic Pressure

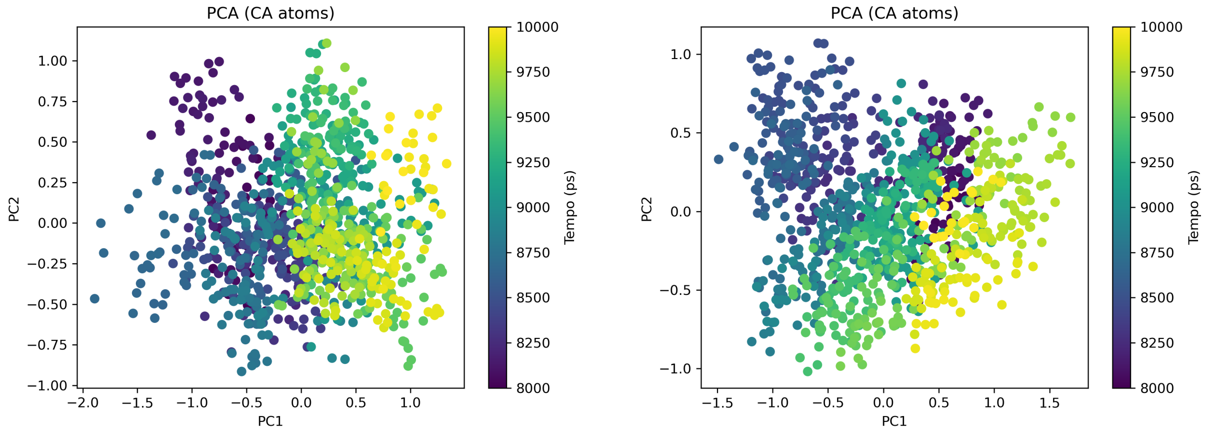

E) 1000 bar F) 1250 bar

Figure 98: Principal Component Analysis side-by-side - High Hydrostatic Pressure

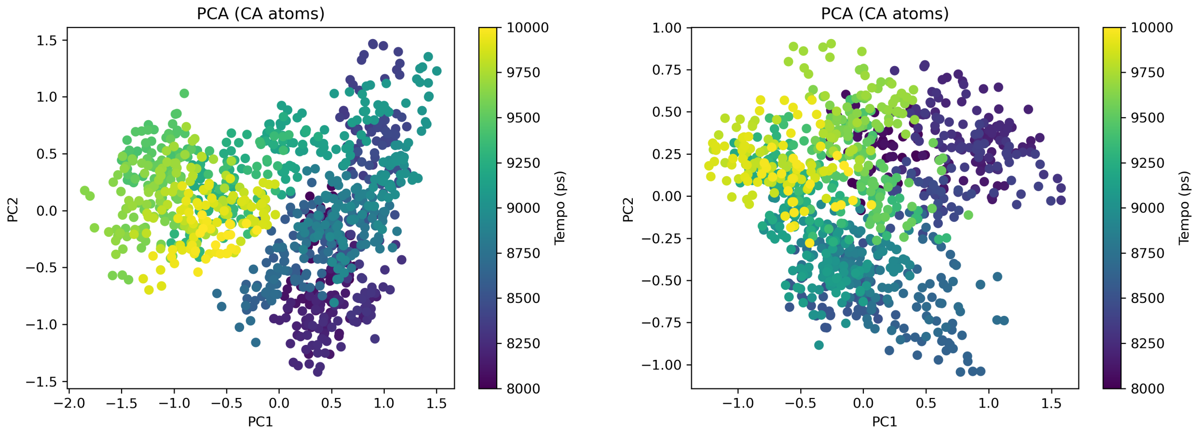

G) 1500 bar

H) 1750 bar

Figure 99: Principal Component Analysis side-by-side - High Hydrostatic Pressure

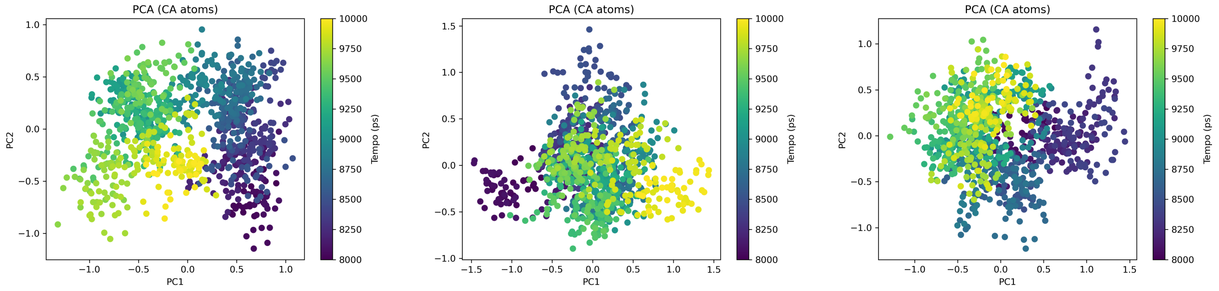

I) 2000 bar

J) 2250 bar

K) 2500 bar

Figure 100: Principal Component Analysis side-by-side - High Hydrostatic Pressure

**HHP and low temperature**

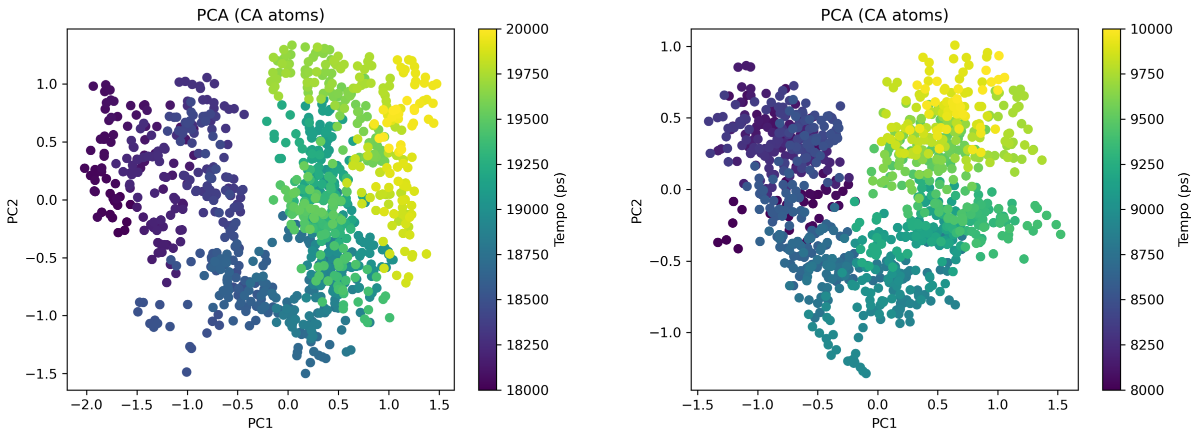

A) 1 bar

B) 250 bar

Figure 101: Principal Component Analysis side-by-side - High Hydrostatic Pressure and Low Temperature

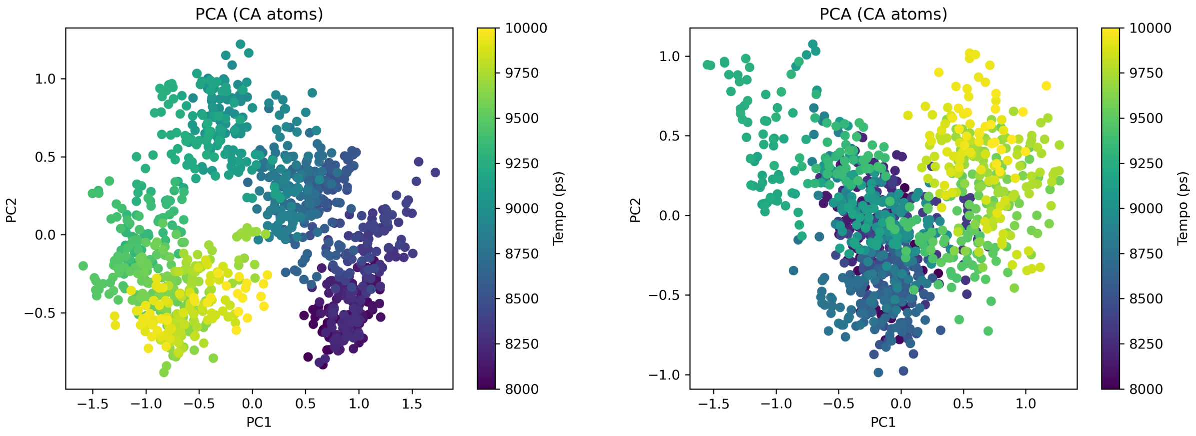

C) 500 bar

D) 750 bar

Figure 102: Principal Component Analysis side-by-side - High Hydrostatic Pressure and Low Temperature

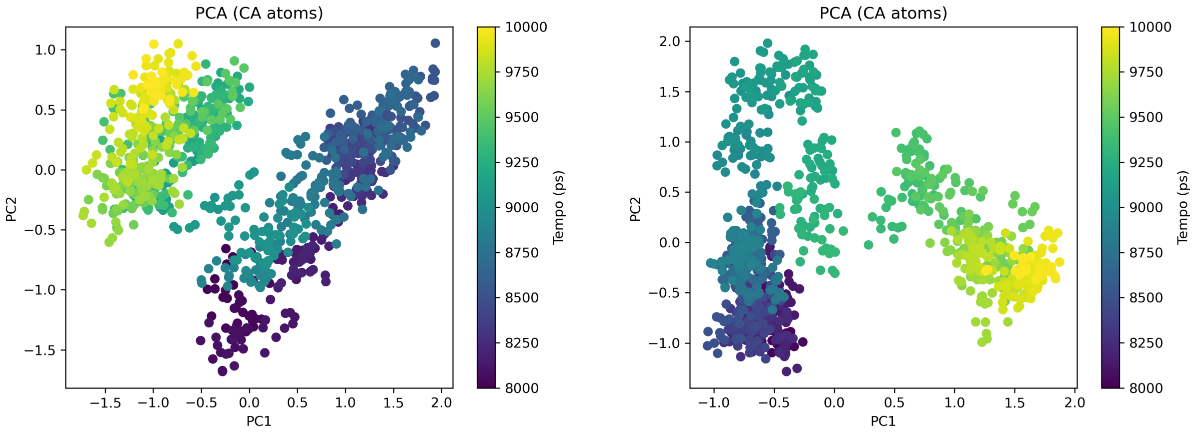

E) 1000 bar

F) 1250 bar

Figure 103: Principal Component Analysis side-by-side - High Hydrostatic Pressure and Low Temperature

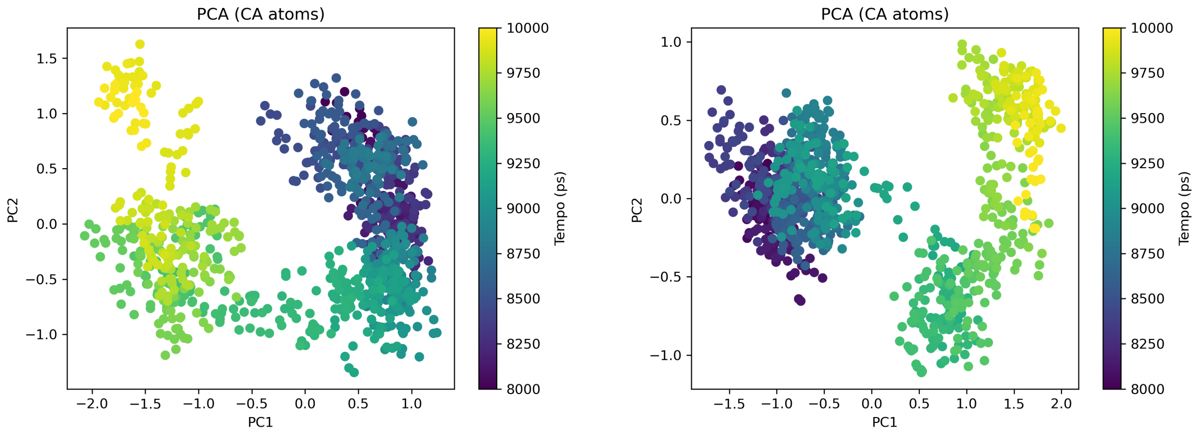

G) 1500 bar

H) 1750 bar

Figure 104: Principal Component Analysis side-by-side - High Hydrostatic Pressure and Low Temperature

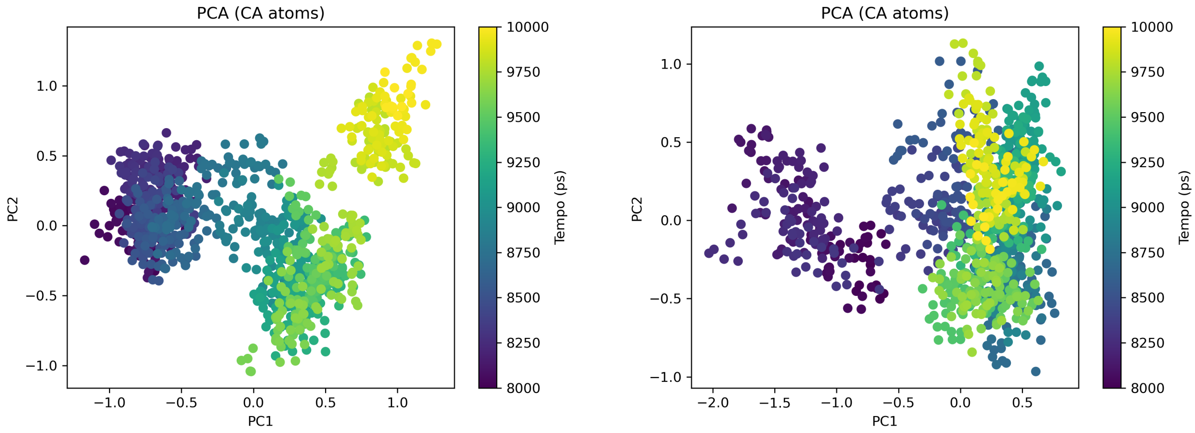

I) 2000 bar

J) 2250 bar

Figure 105: Principal Component Analysis side-by-side - High Hydrostatic Pressure and Low Temperature

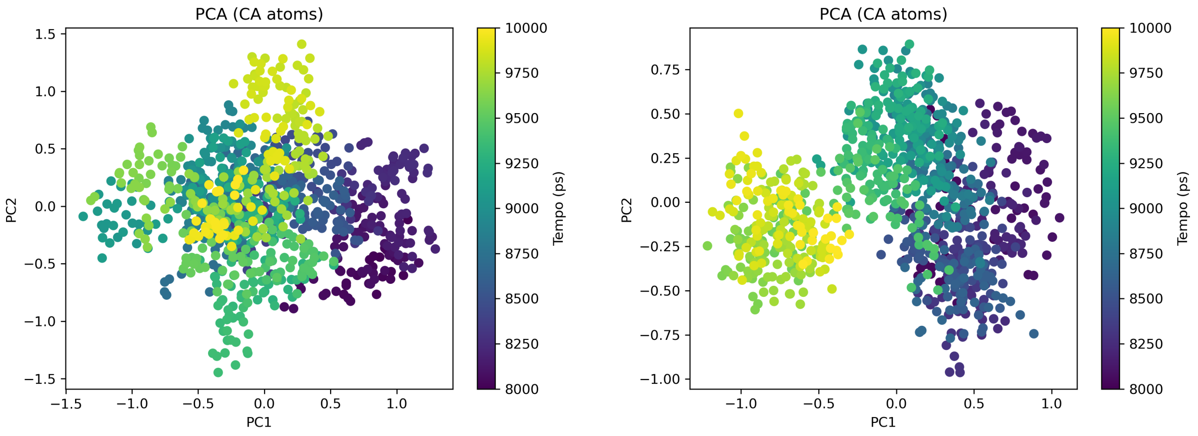

K) 2500 bar

L) 2500 bar @ 300 k

Figure 106: Principal Component Analysis side-by-side - High Hydrostatic Pressure and Low Temperature

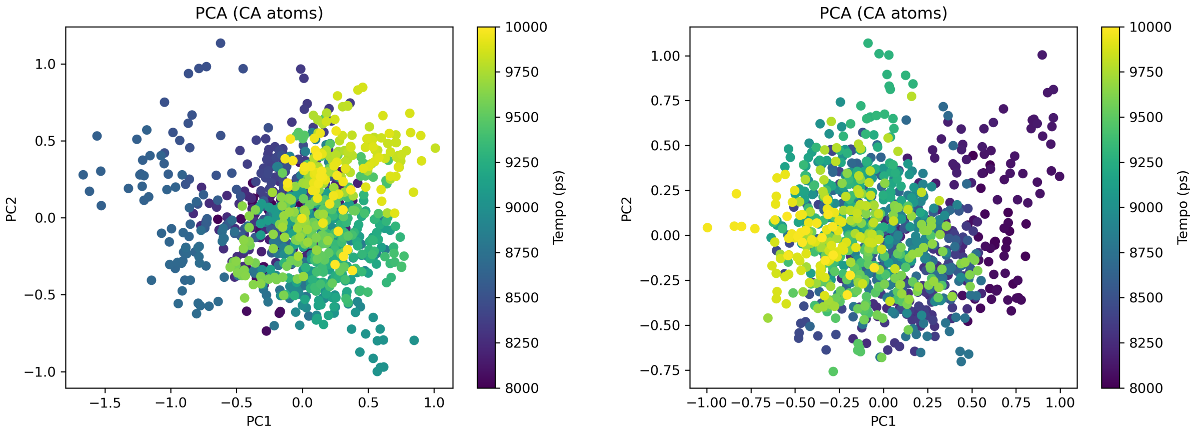

M) 2500 bar @ 295 k

N) 2500 bar @ 290 k

Figure 107: Principal Component Analysis side-by-side - High Hydrostatic Pressure and Low Temperature

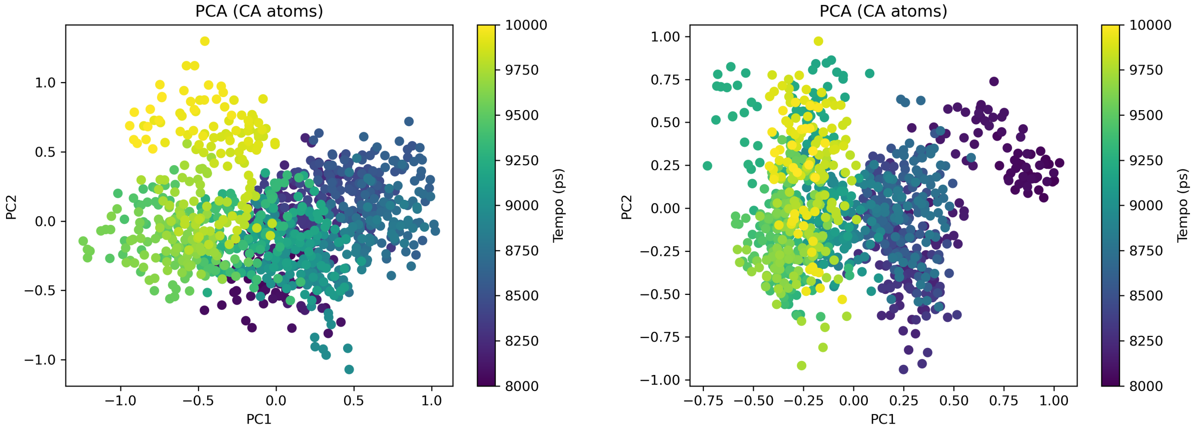

O) 2500 bar @ 285 k

P) 2500 bar @ 280 k

Figure 108: Principal Component Analysis side-by-side - High Hydrostatic Pressure and Low Temperature

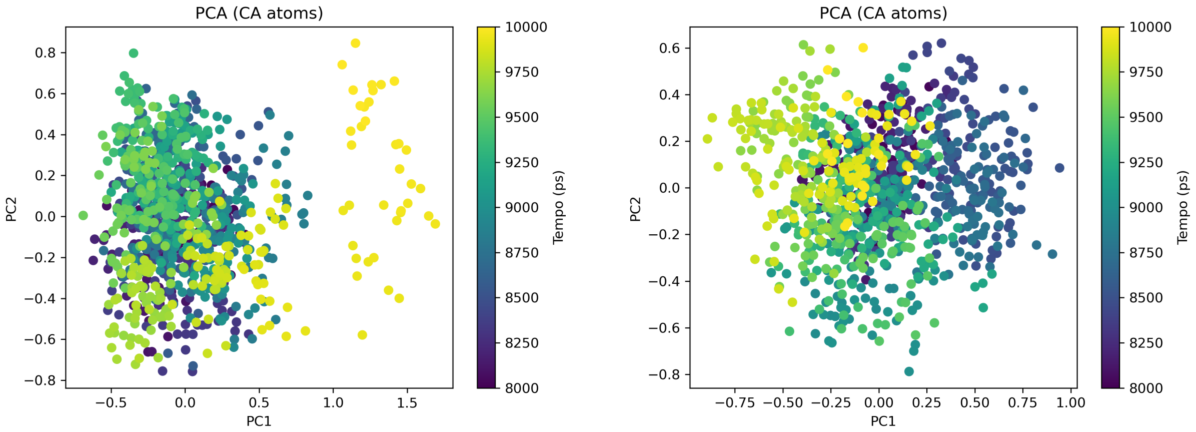

Q) 2500 bar @ 275 k

R) 2500 bar @ 270 k

Figure 109: Principal Component Analysis side-by-side - High Hydrostatic Pressure and Low Temperature

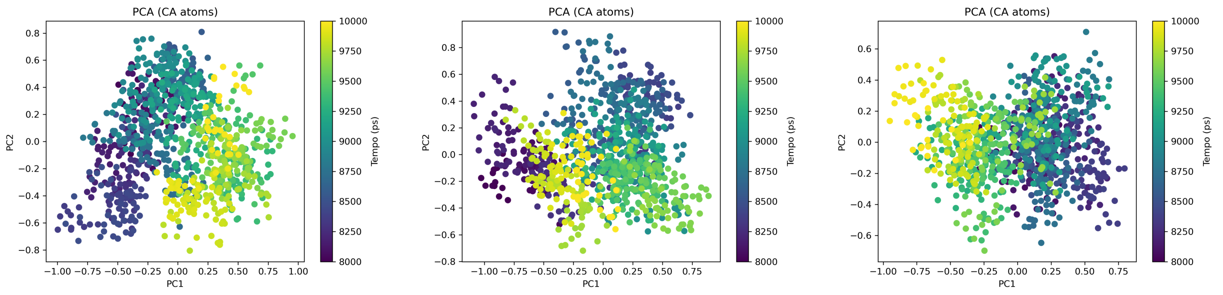

S) 2500 bar @ 265 k T) 2500 bar @ 260 k U) 2500 bar @ 255 k

Figure 110: Principal Component Analysis side-by-side - High Hydrostatic Pressure and Low Temperature
