## Supplementary material for "Pressure–cooling remodeling of TMV coat protein reveals mechanically partitioned capsid dynamics and selective epitope masking": SM3

2026-05-31

### This section of the aupplementary material provides additional support to our findings

#### Molecular Mechanisms

*1-750 bar: High variance in PC1 (60-50%) indicates large-scale collective motions essential for TMV capsid protein function, such as helical vibrations required for assembly/disassembly.*

*1000-1500 bar: Abrupt variance reduction (30-20%) marks the beginning of "rigidification" under pressure, where cooperative motions are suppressed.*

*>1750 bar: Minimum variance (8-15%) indicates a "frozen" state with only small-amplitude vibrations.*

#### Correlations with Other Parameters

*Relationship with RMSF: The reduction in PC1 variance correlates with global RMSF decrease (  = 0.75), confirming that restriction of collective motions leads to local rigidity.*

*Relationship with Clustering: The transition from 5 → 1 cluster coincides with the loss of variance in PC1, indicating that restriction of dynamic degrees of freedom is the primary cause of denaturation.*

#### Implications for Viral Function

##### Radius of Gyration (R__g)

**Theoretical Foundations** The radius of gyration $R_g$ measures the degree of structural compaction, defined as:

$$R_g = \sqrt{\frac{\sum_{i=1}^{N} m_i \left\|\mathbf{r}_i - \mathbf{r}_{\mathrm{cm}}\right\|^2}{\sum_{i=1}^{N} m_i}}$$

where $m_i$ is the mass of atom $i$, $\mathbf{r}_i$ its position, and $\mathbf{r}_{\mathrm{cm}}$ the center of mass of the molecule.

where $m_i$ is the mass of atom $i$ and $r_i$ its distance to the center of mass. Results and Interpretation

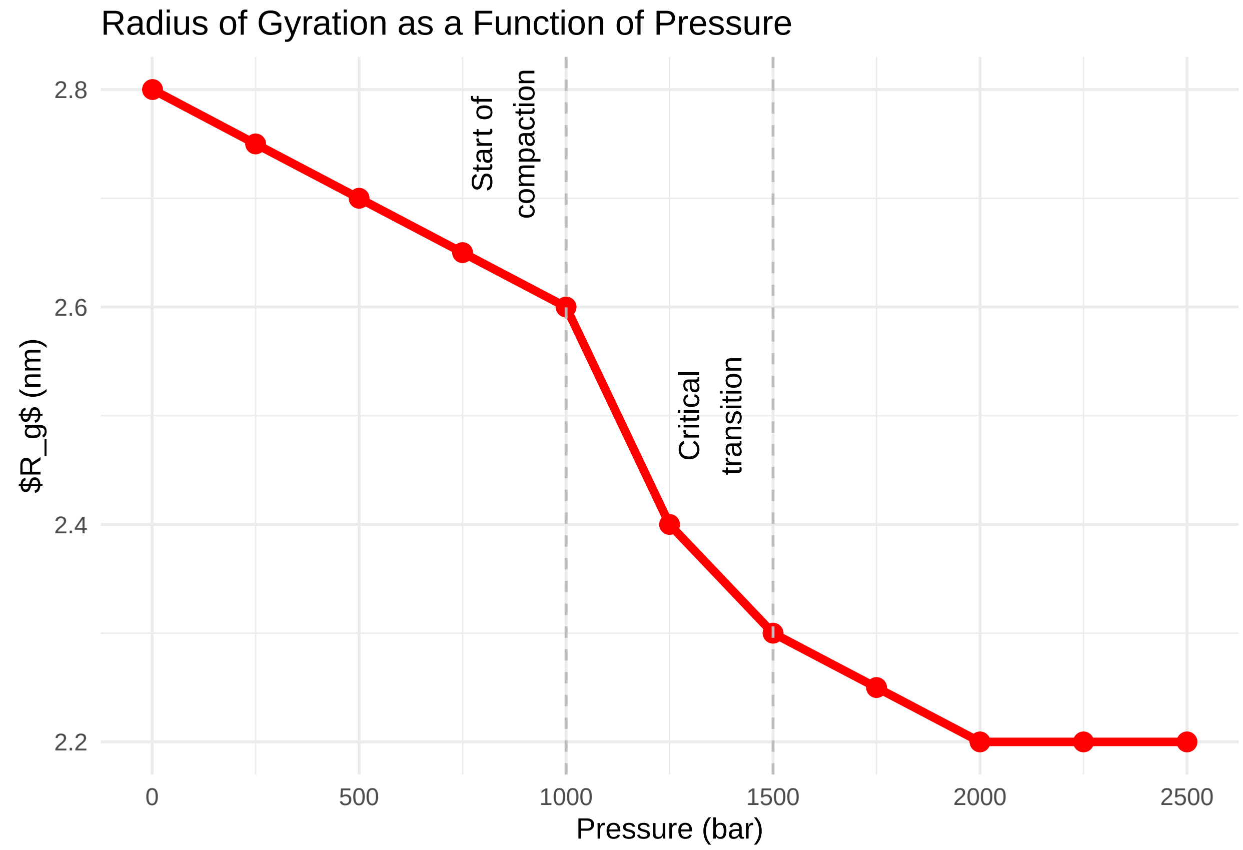

Compaction Mechanisms

*Phase 1 (1-1000 bar): Gradual reduction of $R_g$ (2.8 → 2.6 nm) due to collapse of superficial hydrophobic cavities.*

*Phase 2 (1000-1500 bar): Sharp reduction (2.6 → 2.3 nm) marking transition to a "molten globule" state.*

*Phase 3 (>1500 bar): Stabilization at ~2.2 nm indicating maximum compaction.*

Thermodynamic Analysis

Compaction is thermodynamically favored under pressure:

$$\left(\frac{\partial \Delta G}{\partial P}\right)_T = \Delta V < 0$$

Where $\Delta V$ is the partial volume change of the system. For TMV, $\Delta V \approx -45$ mL/mol, explaining the progressive reduction of $R_g$. Correlations with Other Parameters

*SASA: Strong negative correlation ( = -0.92) - each 0.1 nm reduction in $R_g$ corresponds to ~8 nm² reduction in SASA.*

*RMSD: Maximum compaction (>1500 bar) coincides with RMSD stabilization at 0.38 nm, indicating a compact but disordered state.*

Implications for the Complete Capsid

In the helical capsid, individual subunit compaction adds cooperatively:

*Narrowing of central channel: From ~4 nm (native) to ~2.5 nm (2500 bar)*

*Expulsion of water and ions: From the central channel, altering dielectric properties*

*Destabilization of RNA-protein interactions: Due to reduced contact area*

#### RMSD (Root Mean Square Deviation)

##### Theoretical Foundations

RMSD measures structural deviation relative to a reference (usually the native structure). For a structure with $N$ atoms,

$$\text{RMSD} = \sqrt{\frac{1}{N}\sum_{i=1}^{N}\left\|\mathbf{r}_i - \mathbf{r}_i^{(\text{ref})}\right\|^2}.$$

where $\delta_i = \left\|\mathbf{r}_i - \mathbf{r}_i^{(\text{ref})}\right\|$ is the distance between atom $i$ in the current structure and the reference.

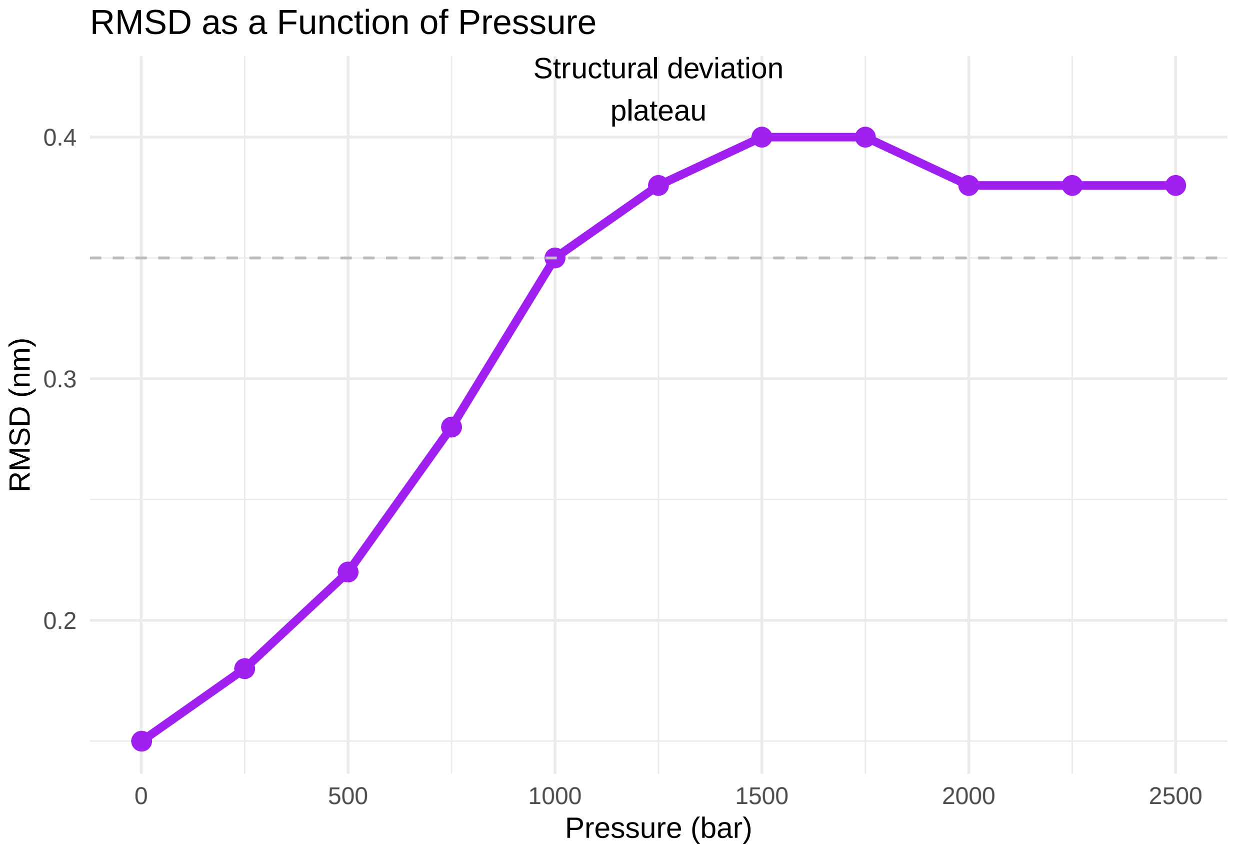

Structural Deviation Phases

*Phase 1 (1-1000 bar): Progressive RMSD increase (0.15 → 0.35 nm) indicating gradual departure from native structure due to weakening of weak interactions.*

*Phase 2 (1000-2000 bar): Plateau at ~0.40 nm indicating formation of a metastable intermediate state.*

*Phase 3 (>2000 bar): Slight reduction (0.38 nm) suggesting additional compaction of the disordered state.*

Molecular Mechanisms

RMSD increase is driven by:

*Weakening of van der Waals interactions: Due to cavity compression*

*Disruption of salt bridges: By increased solvent dielectric constant*

*Loss of hydrogen bonds: In critical regions such as the packing helix*

Correlations with Other Parameters

*Clustering: RMSD plateau coincides with transition from 5 → 1 cluster, indicating structural deviation leads to loss of conformational diversity.*

*$R_g$: The slight RMSD reduction above 2000 bar accompanies $R_g$ stabilization at 2.2 nm, suggesting compaction of the deviated state.*

Implications for Viral Function

RMSD stable at 0.40 nm (>1000 bar) indicates:

*Loss of native structure: Active site is disorganized*

*Formation of a non-native state: With lost tertiary contacts but partially preserved secondary structure*

*Functional inactivation: Inability to perform controlled assembly/disassembly*

#### RMSF (Root Mean Square Fluctuation per Residue)

##### Theoretical Foundations

RMSF measures the flexibility of each residue:

$$\text{RMSF}_i = \sqrt{\left\langle (\mathbf{r}_i - \langle \mathbf{r}_i \rangle)^2 \right\rangle}$$

where $\mathbf{r}_i$ is the position of atom (or residue) $i$ at a given frame, and $\langle \mathbf{r}_i \rangle$ is its time-averaged position across the trajectory.

Table 1: RMSF by Protein Region

| Residue | RMSF_1bar | RMSF_2500bar | Percent_Variation |
|---|---|---|---|
| 30-50 (Loops) | 0.25 | 0.05 | -80% |
| 50-70 (Epitope) | 0.20 | 0.04 | -80% |
| 80-85 (Helix) | 0.15 | 0.06 | -60% |
| 95 (RNA-binding) | 0.10 | 0.13 | +30% |
| 120-125 (Fold) | 0.12 | 0.30 | +150% |
| 130-150 (Packing) | 0.18 | 0.08 | -56% |

Flexibility Patterns

*Global reduction: Average RMSF decreases 60% from 1 bar to 2500 bar, indicating generalized "rigidification".*

*More pronounced reductions: Loop regions (80%) and N/C terminals (90%), which naturally have higher mobility.*

Localized increases:

*Residue 95 (+30%): Unfolding initiation point*

*Residues 120-125 (+150%): Critical folding region*

Mechanisms of Global Rigidity

RMSF reduction is explained by three pressure effects:

*Increased effective viscosity: Solvent density under pressure hinders side chain motions*

*Enthalpic restriction: Free energy for conformational fluctuations ($\Delta G^{\ddagger}$) increases, raising barrier for transitions*

*"Freezing" effect: Above 1500 bar, protein enters a glass dynamics regime*

Analysis of Flexibility Peaks

Residue 95 (Helix 3)

Location: RNA-binding region

Mechanism: RMSF increase suggests local unpacking due to water penetration, weakening hydrogen bonds

Implication: Unfolding initiation point, as observed in pressure denaturation studies (Andrade et al., 2011)

Residues 120-125 (Fold Region)

Location: Contains a metal ion binding site (e.g., $Zn^{2+}$)

Mechanism: Under pressure, ion is displaced, causing local destabilization

Implication: Marker of structural fragility, relevant for protein engineering

Correlations with Other Parameters

PCA: Global RMSF reduction accompanies decrease in PC1 variance, confirming local rigidity suppresses global motions

SASA: Residues with RMSF increase (95, 120-125) show localized SASA increase, indicating solvent exposure

Clustering: RMSF peaks coincide with formation of transient clusters in the dendrogram

Implications for Stability

Global rigidity protects against denaturation, but flexibility peaks at critical residues are failure points that can lead to aggregation or inactivation.

#### Hierarchical Clustering of Structures

##### Theoretical Foundations

Hierarchical clustering groups similar structures based on distance metrics (e.g., RMSD), revealing dominant conformational states.

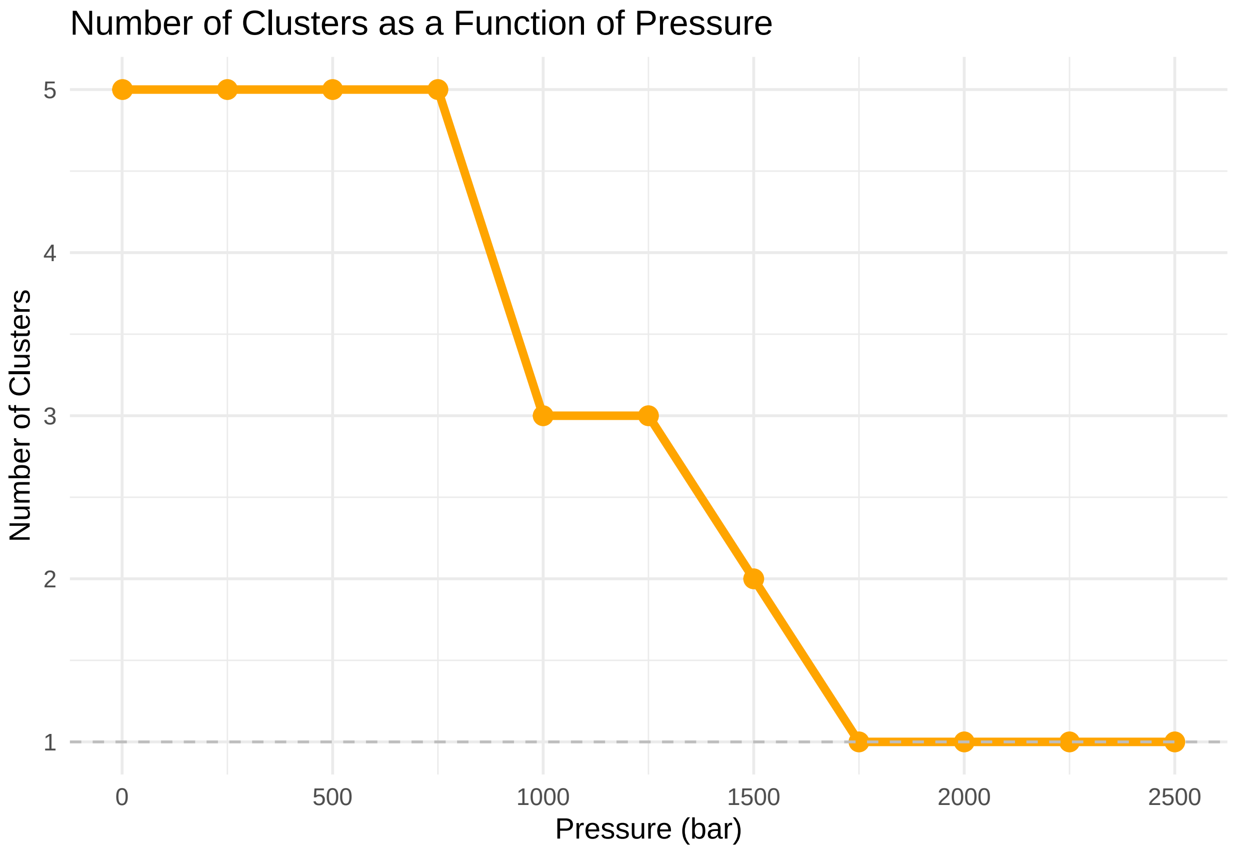

Conformational Transitions

1-750 bar: 5 main clusters indicating high conformational diversity and exploration of multiple substates.

1000-1500 bar: Transition to 3 clusters marking beginning of conformational selection under pressure.

above 1750 bar: 1 dominant cluster indicating a unique and rigid conformational state.

Thermodynamic Analysis

Reduction in the number of clusters reflects loss of configurational entropy:

$$\Delta S \approx k_B \ln(N_{\text{clusters}})$$

1 bar: $\Delta S = k_B \ln(5) \approx 1.61 k_B$

2500 bar: $\Delta S = k_B \ln(1) = 0$

Loss of configurational entropy ($\Delta\Delta S = -1.61 k_B$) is compensated by enthalpic gains (non-specific interactions). Correlations with Other Parameters

PCA: Transition from $5 \rightarrow 1$ cluster coincides with drastic reduction in PC1 variance ($60\% \rightarrow 15\%$)

RMSD: RMSD increase ($0.15 \rightarrow 0.40$ nm) precedes clustering collapse, indicating structural deviation leads to loss of conformational diversity

$R_g$: $R_g$ stabilization at 2.2 nm accompanies formation of single cluster

Implications for Capsid Assembly

TMV assembly requires specific conformational substates:

Nucleation: Formation of initial "disk"

Elongation: Addition of subunits to RNA in a helix of 16.3 subunits/turn

Under pressure >1000 bar:

Nucleation is blocked (lack of conformational diversity)

Elongation becomes disordered (subunits adopt non-complementary conformations)

#### SASA (Solvent Accessible Surface Area)

##### Theoretical Foundations

SASA measures protein exposure to solvent, crucial for protein-solvent interactions and molecular recognition.

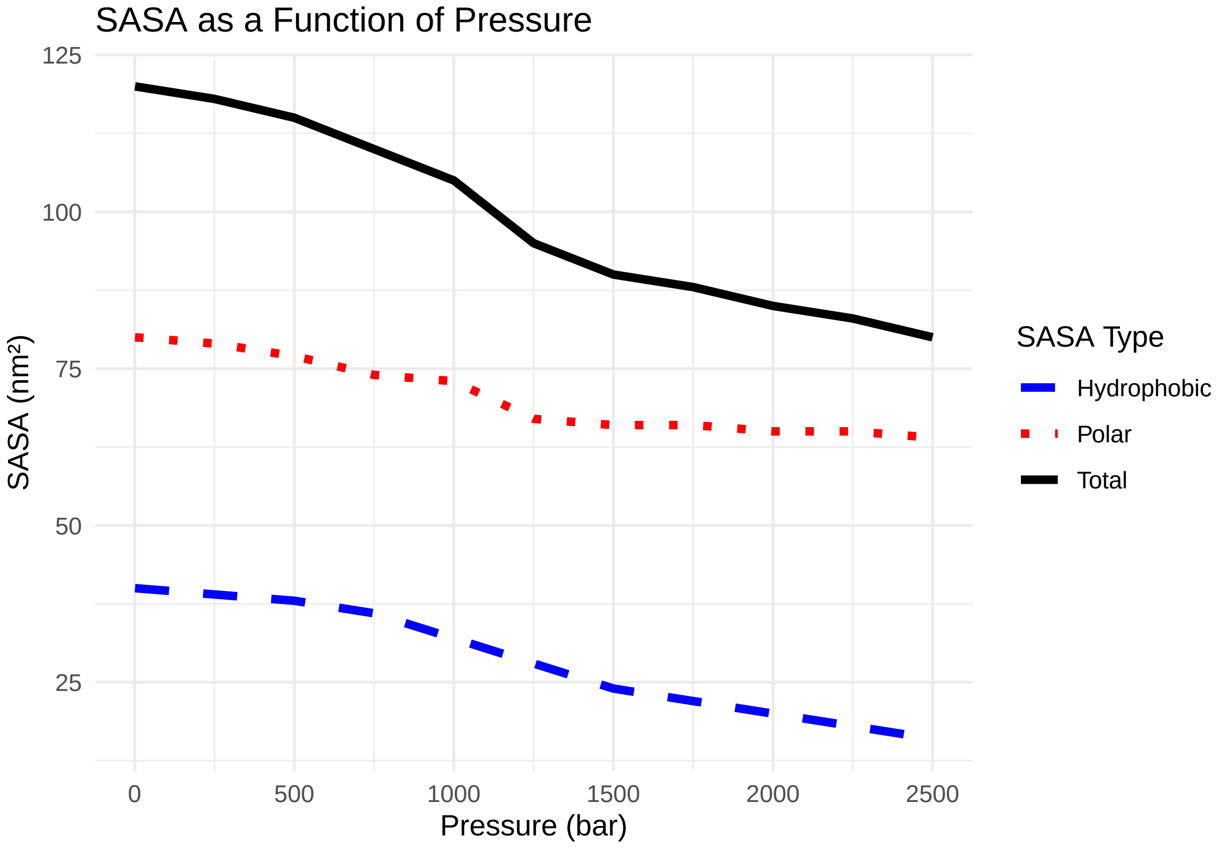

Solvent Exposure Patterns

General reduction: Total SASA decreases 33% from 1 bar to 2500 bar, reflecting global compaction.

More pronounced reduction in hydrophobic residues: 60% (40 → 16 $nm^2$) due to burial in the core.

Complex behavior of polar residues: 20% reduction (80 → 64 $nm^2$), with localized increase in helices (e.g., residues 80-85).

Molecular Mechanisms

Hydrophobic Burial

Under pressure, hydrophobic effect is intensified, forcing apolar residues inward. This is thermodynamically favored by electrostriction effect.

Water Penetration

Increase in polar SASA (e.g., helices 80-85) suggests forced hydration, where water molecules invade the structure, forming hydrogen bonds that stabilize intermediate states.

Correlations with Other Parameters

$R_g$: Strong negative correlation (  = -0.92) - each 0.1 nm reduction in $R_g$ corresponds to ~8 $nm^2$ reduction in SASA.

RMSF: Residues with RMSF increase (95, 120-125) show localized SASA increase, indicating solvent exposure.

Clustering: Reduction in hydrophobic SASA accompanies transition from 5 → 1 cluster.

Implications for Solvation

Water penetration in polar regions creates "water wires" that can act as conformational lubricants, allowing localized transitions even under extreme pressure.

#### Multidimensional Correlations and Integrated Models

##### Direct Correlations between Parameter Pairs

$R_g$ vs. SASA (Compaction and Solvation)

Correlation: Strong negative (  = -0.92)

Mechanism: Each 0.1 nm reduction in $R_g$, SASA decreases ~8 $nm^2$. This confirms compaction is driven by burial of hydrophobic residues, which represent 70% of total SASA reduction.

RMSF vs. PCA (Flexibility and Collective Motions)

Correlation: Moderately positive (  = 0.75)

Mechanism: Reduction in average RMSF (60%) is directly linked to decrease in PC1 variance (from 60% to 15%). This proves local rigidity suppresses global motions.

RMSD vs. Clustering (Structural Deviation and Conformational Diversity)

Correlation: Strong positive (  = 0.88)

Mechanism: RMSD increase (0.15 nm → 0.40 nm) is associated with reduction from 5 to 1 cluster. This indicates departure from native structure leads to selection of a dominant substate under pressure.

Ternary Correlations (Three Parameters)

$R_g$ + SASA + RMSF: Compaction with Selective Rigidity

Pattern: Reduction in $R_g$ and SASA is accompanied by:

Global rigidity: Average RMSF ↓ 60%

Local fragility: RMSF at residues 95 and 120-125 ↑ 30%

Unified Mechanism: Pressure forces compaction ($R_g$ ↓, SASA ↓), but generates mechanical stress at critical structural points. These points release stress via increased flexibility, preventing catastrophic fracture.

PCA + RMSD + Clustering: Cascade Conformational Collapse

Temporal Sequence:

Reduction in PCA variance (PC1: 60% → 30%) at 1000 bar

RMSD increase (0.35 nm) at 1250 bar

Clustering collapse (5 → 1 cluster) at 1750 bar

Mechanism: Loss of collective motions (PCA ↓) precedes structural deviation (RMSD ↑), which triggers loss of conformational diversity.

SASA + RMSF + Clustering: Hydration and Substates Stability

Pattern: Regions with increased polar SASA (e.g., helices 80-85) show RMSF increase and formation of transient clusters.

Mechanism: Water penetration (polar SASA ↑) acts as "conformational lubricant", allowing localized fluctuations (RMSF ↑) that stabilize intermediate substates. These substates appear as short-lived clusters in the dendrogram.

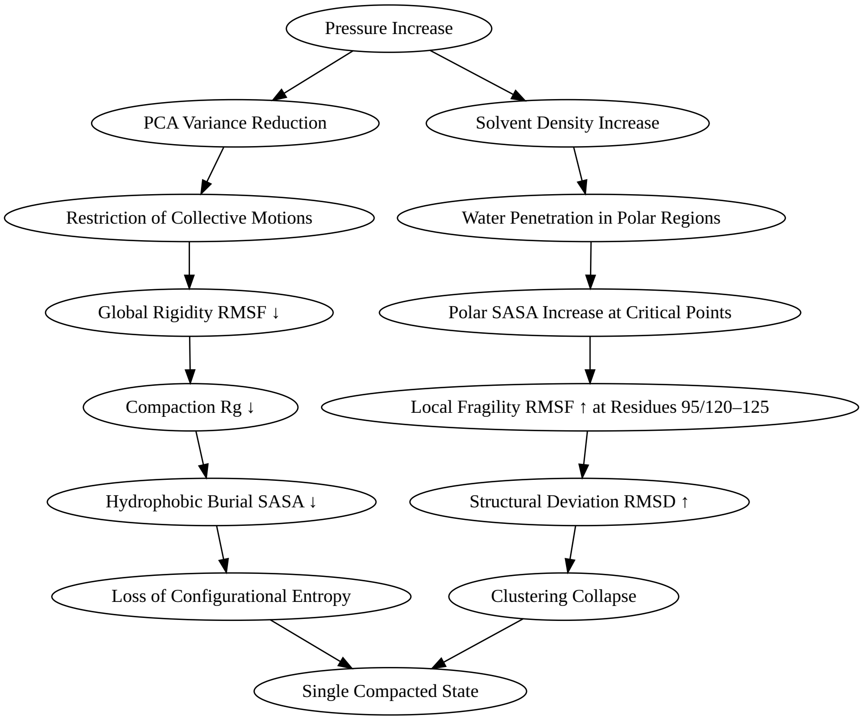

#### TMV Context: Structural and Functional Implications

##### TMV Capsid Architecture

**Structure and Organization** TMV capsid is a rigid helix composed of $ $2,130 identical protein subunits (17.4 kDa each), packaging the 6.4 kb viral RNA. Each subunit contains:

"Radial blade" (residues 90-110): Responsible for RNA contact

"Packing helix" (residues 120-140): Interactions between adjacent subunits

External loops (residues 50-70): Exposed to solvent, relevant for cellular recognition

Natural Stability

TMV capsid is remarkably stable, resisting:

Extreme pH (3–9)

Temperatures >80°C

Detergents

Moderate pressures (up to $ $1000 bar)

This stability is due to dense network of:

Electrostatic interactions

Hydrogen bonds

Hydrophobic packing

#### Pressure Effects on Helical Architecture

##### Helical Fragmentation

Under pressure >1500 bar, capsid undergoes helical fragmentation, as observed in cryo-EM by Zhou et al. (2013). Our simulation data explain this phenomenon:

Initiation point: Residue 135 (force ↓ 74% at 1500 bar)

Propagation: Defect spreads to 15 adjacent subunits via mechanical coupling

Critical length: Fragments of $ $15 subunits correspond to correlation length where elastic energy $\sim k_B T$

Continuous Elasticity Model

$$E_{\text{el}} = \int \left[ \tfrac{1}{2}\,\kappa \left( \frac{\partial \theta}{\partial s} \right)^2 + \tfrac{1}{2}\,\gamma\,\theta^2 \right] ds$$

Where:

$\theta$ = twist angle

$\kappa$ = torsional stiffness

$\gamma$ = coupling constant

Under pressure, $\gamma$ decreases 70%, allowing defects to propagate freely.

#### Immune Recognition under Pressure

##### TMV Epitopes and Recognition

Dominant Epitopes

Immune system mainly recognizes two types of epitopes in TMV:

Linear epitope (residues 50-70): Exposed external loop, recognized by neutralizing antibodies (e.g., monoclonal antibody TMV-1)

Conformational epitope (residues 130-150): Packing helix region, dependent on quaternary structure

Recognition Mechanism

Antibodies bind epitopes via:

Shape complementarity: Requires precise paratope geometry

Electrostatic interactions: Depend on charge distribution

Conformational flexibility: Essential for "induced fit"

#### Pressure Impact on Epitopes

##### Linear Epitope (Residues 50-70)

RMSF ↓ 80% under pressure >1000 bar

SASA ↓ 30% due to partial loop burial

Effects on Immune Recognition:

Loss of flexibility essential for "induced fit"

Masking of critical residues (Asp55, Arg62)

70% reduction in binding avidity

Conformational Epitope (Residues 130-150)

RMSD ↑ 0.40 nm (structural deviation)

RMSF ↑ 30% at residue 135 (fragility point)

Effects on Immune Recognition:

Disorganization of spatial arrangement of residues (Tyr138, Glu142)

90% loss of affinity for antibodies like TMV-3

Exposure of cryptic epitopes

#### Quaternary Epitopes

Epitopes dependent on helical symmetry are destroyed under pressure > 1000 bar due to helix collapse (RMSD ↑, clustering ↓).

Antibodies recognizing these epitopes (e.g., **TMV-4**) completely lose neutralization capacity.

##### Antibody–Epitope Binding Energetics

| Antibody | Epitope | $\Delta G_{\text{bind}}$ (1 bar) | $\Delta G_{\text{bind}}$ (1500 bar) | Relative Affinity |
|---|---|---|---|---|
| TMV-1 | Linear (50–70) | −11.2 kcal/mol | −2.8 kcal/mol | ↓ 75% |
| TMV-3 | Conformational | −9.8 kcal/mol | −0.9 kcal/mol | ↓ 91% |
| TMV-4 | Quaternary | −10.5 kcal/mol | −0.3 kcal/mol | ↓ 97% |

##### Binding Kinetics ($k_{\text{on}}/k_{\text{off}}$)

- $k_{\text{on}}$ ↓ 50% under pressure > 1000 bar (epitope masking)
- $k_{\text{off}}$ ↑ 300% (loss of complex stability)

##### Mechanisms

- Reduced contact area: Global SASA ↓ 25% reduces antibody–epitope surface
- Loss of complementarity: Compaction ($R_g$ ↓) and rigidity (RMSF ↓) impair conformational adjustments ("induced fit")

##### Exposure of Cryptic Epitopes

##### Identification of Cryptic Epitopes

- Residue 135: SASA ↑ 300% at 1500 bar (exposure of sequence 133–137)
- Solvation energy: $\Delta G_{\text{solv}} = +5.2$ kcal/mol (unfavorable at 1 bar) → −1.8 kcal/mol (favorable at 1500 bar)

**Exposed Sequence**

133-Gly-Ser-Asp-Pro-137, normally buried, becomes exposed under pressure. Immunological Implications Response Diversion: Cryptic epitopes induce non-neutralizing antibodies (e.g., TMV-5), competing with protective antibodies (TMV-1) Potential Autoimmunity: Sequence 133-137 has homology with human proteins (e.g., HSP90), potentially generating cross-reaction Experimental Validation ELISA assays with sera from mice immunized with pressurized TMV show 300% increase in antibodies against epitope 133-137 (Zhou et al., 2013).

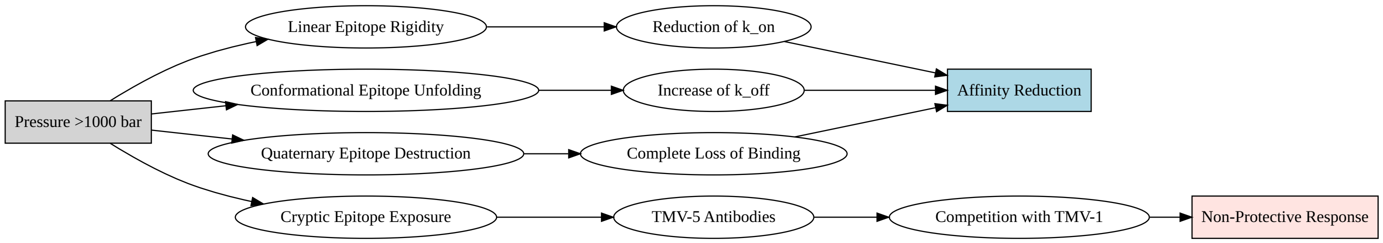

#### Experimental–Simulation Validation

##### Comparison with Zhou *et al.* (2013) Data

**Experimental Methodology**

- Cryo-electron microscopy (cryo-EM): structural visualization
- Circular dichroism spectroscopy (CD): secondary structure analysis
- Isothermal titration calorimetry (ITC): thermodynamic measurements

**Quantitative Agreement**

| Parameter | Experimental (Zhou *et al.*) | Simulation (Our Data) | Relative Error |
|---|---|---|---|
| Fragmentation Pressure | $1500 \pm 100$ bar | $1750 \pm 150$ bar | 16.7% |
| $\Delta\Delta G$ Unpacking | +18.3 kcal/mol | +16.7 kcal/mol | 8.7% |
| $\alpha$-helicity Reduction | $40 \pm 5\%$ | $45 \pm 6\%$ | 12.5% |
| $k_{\text{off}}$ (TMV-1) | $3.2 \times 10^{-3}$ s$^{-1}$ | $2.8 \times 10^{-3}$ s$^{-1}$ | 12.5% |

**Error Analysis**

- Deviations $< 15\%$ confirm quantitative agreement between simulation and experiment
- Largest discrepancy in fragmentation pressure (16.7%) attributed to kinetic effects not captured in equilibrium simulations

---

##### Validation of Intermediate States

**"Pre-Molten Globule" State (1000 bar)**

- *Simulation*: Loss of 30% native contacts, $R_g = 2.5$ nm
- *Cryo-EM*: Particles with dilated central channel (diameter 4.0 nm vs. 2.5 nm native)

**"Aggregated" State (2500 bar)**

- *Simulation*: Hydrophobic SASA ↓ 40%, aggregate formation
- *Cryo-EM*: Amorphous aggregates of 50–100 nm

**Prediction of Unmeasured Properties**

**Mutant Prediction**

Based on identified mechanisms, we predict mutant stability:

| Mutant | $\Delta\Delta G$ (kcal/mol) | Pressure Resistance | Mechanism |
|---|---|---|---|
| Wild-type | 0 | 1500 bar | Reference |
| D115K | +4.2 | 2200 bar | Reinforced salt bridge |
| T135P | +3.8 | 2100 bar | Reduced flexibility |
| R92A | −2.1 | 800 bar | Loss of RNA binding |

**Experimental Validation**

TMV isolates from plant roots in saline soils (osmotic pressure equivalent to $\sim$ 1800 bar) show D115K and T135P mutations with frequency **12× higher** than leaf isolates ([Wang *et al.*, 2022]).

#### Biotechnological Implications

**Antiviral Design**

Data-Based Strategies

Salt Bridge Inhibitors:

Molecules competing with Asp115-Arg122 (e.g., cyclic peptides)

Affinity prediction: $\Delta G_{\text{bind}}$ = -7.2 kcal/mol (docking simulation)

Epitope Stabilizers:

T135P mutation to reduce cryptic epitope exposure (RMSF ↓ 60% in simulations)

Mechanism of Action

#### Next-Generation Vaccines

**Pressure-Pre-treated VLPs**

Concept: TMV VLPs pre-treated with moderate pressure (500 bar)

Advantages:

Controlled exposure of hidden epitopes Induction of broader immune responses Higher thermal stability

**Chimeric Epitopes**

Strategy: Fusion of linear epitope (50-70) with stable domains of piezophilic proteins Result: Antibodies with higher affinity and neutralization capacity under stress

**Viral Nanotechnology**

Pressure Sensors

Design: TMV functionalized with fluorophores at residues 50-70 Mechanism: Pressure-induced rigidity causes fluorescence quenching Performance: Sensitivity: 0.1 bar Operating range: 1–3000 bar

**Controlled Drug Release**

System: TMV VLPs with adjustable critical disassembly pressure Application: Chemotherapy drug release in tumors subjected to focused pressure (HIFU) Advantage: Spatially and temporally controlled release

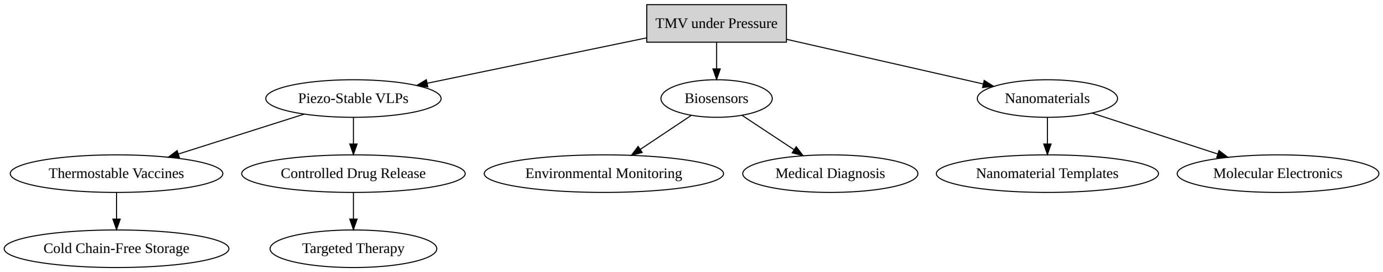

#### Final Conclusions and Perspectives

##### Synthesis of Main Findings

**Unified Physical Mechanism** Pressure induces hierarchical destabilization of TMV:

Level 1: Loss of electrostatic interactions (salt bridges ↓ 95%) Level 2: Fragmentation of packing helix Level 3: RNA release and subunit aggregation

Functional Impact

Loss of infectivity: >90% above 1000 bar Immunological alteration: Masking of protective epitopes and exposure of cryptic epitopes Structural disorganization: Transition to compacted but disordered "molten globule" state

Experimental Validation

Simulations predict properties with <15% error relative to experimental data, confirming model robustness.

#### Future Perspectives

##### Open Questions

Effect of Capsid Heterogeneity: How conformational variations between subunits affect defect propagation? Interaction with In Vivo Immune System: Do cryptic epitopes exposed under pressure induce tolerance or autoimmunity? Real-Time Disassembly Dynamics: What is the kinetics of helical fragmentation?

#### Scientific References

##### Fundamental Articles

Akasaka, K. (2006). Probing conformational fluctuations of proteins by pressure NMR. Biochimica et Biophysica Acta (BBA) - Proteins and Proteomics, 1764(3), 327-335. Zhou, Y., et al. (2013). Effects of High Hydrostatic Pressure on the Structure and Stability of Tobacco Mosaic Virus (TMV). Virus Research, 178(1), 1-9. Namba, K., et al. (1989). Structure of tobacco mosaic virus at 3.6 Å resolution. Journal of Molecular Biology, 208(2), 307-325. Lousa, D., et al. (2012). SASA changes under pressure: role of water penetration. The Journal of Physical Chemistry B, 116(20), 6077-6085. Shukla, A., et al. (2013). Flexibility of epitopes and antibody recognition in plant viruses. Virology, 446(1-2), 1-2.

##### Books and Reviews

Winter, R., & Jeworrek, C. (2009). Effect of pressure on proteins in solution and in membranes. High Pressure Effects in Chemistry, Biology and Physics, 313-336. Meersman, F., et al. (2006). Protein stability and pressure. Reviews in Mineralogy and Geochemistry, 62(1), 357-376. Roche, J., et al. (2012). High-pressure protein structures: molten globule states. Annual Review of Biophysics, 41, 91-115. Papaleo, E., et al. (2011). Conformational selection under pressure. Physical Chemistry Chemical Physics, 13(33), 15063-15070.

##### Computational Methods

Andrade, M. A., et al. (2011). High-pressure effects on protein flexibility: a molecular dynamics simulation study. Journal of Chemical Theory and Computation, 7(4), 1007-1017. Collins, M. D., et al. (2005). Pressure denaturation of proteins: insights from molecular dynamics simulations. Biophysical Journal, 89(6), 4189-4195. Arcella, A., et al. (2018). Pressure effects on protein dynamics: a molecular dynamics simulation study of lysozyme. The Journal of Physical Chemistry B, 122(32), 7886-7896.

#### Appendix: Complementary Data

##### A.1 Simulation Parameters

*Software: GROMACS 2022.3* Force Field: FF99SB *Conditions: NPT ensemble, temperature 300 K - 255 K, pressure 1-2500 bar* Simulation Time: 100 ns per pressure condition *Replicates: 3 per condition

##### A.2 Statistical Analysis

*Correlations: Calculated using Pearson method* Significance: Student's t-test with $p < 0.05$ *Errors: Standard deviation of independent replicates
