## Supplementary material for "Pressure–cooling remodeling of TMV coat protein reveals mechanically partitioned capsid dynamics and selective epitope masking": SM1

2026-05-31

### Conformational Epitope Prediction

Conformational B-cell epitope prediction with DiscoTope is fundamentally a structure-based exercise that couples residue-level propensities with measures of surface exposure to infer discontinuous antigenic patches. The original method (DiscoTope 1.0) introduced the idea of combining 3D neighborhood information with amino-acid propensities derived from curated antigen–antibody complexes; DiscoTope 2.0 refined this by adopting an improved surface metric (half-sphere exposure), revised spatial neighborhoods, and re-benchmarked thresholds, leading to more reliable performance across independent test sets. Mechanistically, DiscoTope flags residues that are simultaneously (i) enriched for epitope-favored chemistries and (ii) sufficiently accessible on the native fold—features that map well onto the known enrichment of conformational epitopes in flexible, solvent-exposed loops and protrusions. These modeling choices are well documented in the method's primary papers and the DTU server description, and they align with broader reviews showing that accessibility and local dynamics are the strongest single-feature correlates of epitope location.

A central variable in that framework is relative solvent accessibility (RSA)—a normalized expression of solvent-accessible surface area (SASA)—which is typically calculated using Shrake–Rupley "rolling-probe" geometry. Because DiscoTope's scoring depends on surface exposure, any perturbation that shifts the protein's solvent-exposed landscape will alter predicted epitope patches. High pressure is exactly such a perturbation: it favors conformations with reduced system volume (protein + solvent), thereby encouraging the collapse of internal cavities, water penetration into hydrophobic cores, and redistribution of hydrogen-bond/salt-bridge networks. NMR/crystallographic and simulation studies show that pressure-driven unfolding or partial unfolding is often initiated at pre-existing voids and flexible segments, which can change RSA in a residue-specific manner—precisely the type of signal DiscoTope reads. Thus, as hydrostatic pressure increases, one expects non-monotonic RSA changes across the surface: modest pressures unmask cryptic but native-adjacent patches; higher pressures promote larger rearrangements that create entirely new non-native antigenic surfaces. This picture is well supported by the protein-under-pressure literature on cavity collapse and hydration, and by quantitative analyses linking unfolding volume changes to internal voids.

TMV provides a well-characterized testbed for these ideas. Structurally, TMV is a helical nucleoprotein rod composed of ~2,130 copies of a 158-residue coat protein wrapped around a single-stranded RNA; decades of fiber diffraction and, more recently, high-resolution cryo-EM have established atomic models for both the intact virion and assembly intermediates. This precise structural knowledge is ideal for surface-based epitope prediction and for modeling pressure responses, because it anchors RSA estimates to a realistic 3D framework. In practical terms, DiscoTope run on atomistic TMV models will report different epitope landscapes as pressure shifts loop flexibilities, side-chain rotamers, and inter-subunit contacts that define the solvent-exposed helical grooves and lateral interfaces.

Experimentally, these expectations were borne out in the TMV literature: exposure to HHP altered the pattern of antibody recognition relative to native virions, with moderate pressures revealing previously masked epitopes and higher pressures enabling recognition of neo-exposed patches. In the Virology Journal study by Ferreira de Lima Neto and Bonafé, pressure (alone and in combination with low temperature or urea) reshaped the serological fingerprint of TMV, consistent with a hierarchical destabilization of structural domains under pressure and a graded unmasking of antigenic sites. The concordance between (i) the direction and residue-level logic of RSA changes predicted by structure-based tools and (ii) the experimental epitope mapping across pressure conditions validates the conceptual bridge between DiscoTope outputs and pressure-modulated antigenicity.

At the level of physical chemistry, the pressure dependence of protein conformations naturally extends to antibody–antigen complexes: pressures in the 1–4 kbar range can reversibly dissociate oligomers and even some antigen–antibody pairs, reflecting positive volume changes of association. This thermodynamic lens helps interpret pressure-dependent serological assays and suggests that certain antibody interactions will be more pressure-labile than others, depending on interfacial packing, trapped solvent, and cavity formation upon binding. When such complexes dissociate under pressure, epitope exposure measured on the free antigen may no longer mirror the epitope as presented in the bound state—an important caveat that encourages combining DiscoTope with explicit modeling of binding interfaces and, where possible, wet-lab pressure perturbations in sera or monoclonals.

Mechanistically, several linked processes coherently explain the stepwise changes we observe in TMV under pressure and their readout by DiscoTope. First, small increases in pressure perturb marginal tertiary contacts and side-chain packing, often at pre-existing surface grooves or inter-subunit seams, causing local RSA increases without wholesale unfolding; DiscoTope registers these as strengthened or expanded patches around native-adjacent epitopes. Second, further pressure drives water into shallow cavities and along hydrogen-bond networks, increasing local compressibility and micro-hydration; flexible loops and termini become more dynamic, sometimes flipping out to expose new surfaces that score as epitopes. Third, at still higher pressure, partial unfolding or inter-subunit slippage modifies the helical lattice, generating non-native protrusions and de-novo patches that only become DiscoTope-positive in these states. The literature on pressure-induced cavity collapse, hydration, and the primacy of voids in pressure unfolding supports each link of this cascade.

This HHP-tunable antigenic landscape has practical consequences. For discovery, pressure acts as a controlled perturbation to reveal cryptic epitopes—regions immunologically silent at ambient conditions yet structurally predisposed to exposure. For engineering, pressure-responsive "weak points" can be mapped and then stabilized or destabilized (e.g., by targeted mutations or formulation) to bias epitope display in vaccines or diagnostics. For analytical virology, pairing DiscoTope (or related structure-based tools such as ElliPro/SEPPA) with pressure series and RSA tracking offers a quantitative route to rank epitope robustness across environmental stresses relevant to processing, inactivation, or storage. The broader HHP virology literature—though focused mostly on foodborne pathogens—reinforces that capsid proteins are pressure-sensitive in ways that change exposure and immunoreactivity, mirroring what is observed in TMV at the structural/epitope level.

Placing these observations in a methodological context, it is worth recalling both the strengths and limits of structure-based epitope predictors. DiscoTope's reliance on 3D structure and exposure metrics offers specificity for conformational epitopes, outperforming single-feature scales that track only hydrophilicity or flexibility. Yet no single predictor captures the full immunogenic reality: bound vs. unbound conformations, quaternary arrangements, glycosylation or ion occupancy (e.g., $Ca^{2}$ effects in TMV assemblies), and pressure-coupled dynamics can all modulate the true paratope–epitope interface. Hence, convergent evidence—structure-based predictions under multiple conditions, complementary tools, and experimental mapping—provides the most reliable guide for identifying pressure-sensitive antigenic sites.

#### Structural Models and Epitope Prediction

Protein structures were obtained in PDB format from molecular dynamics trajectories and used as input for conformational B-cell epitope prediction. Conformational epitopes were identified with DiscoTope 3.0, which provides per-residue prediction scores calibrated to surface exposure and residue statistics. The output was tabulated in .csv files, containing residue identity, position, chain information, DiscoTope score, calibrated score, epitope classification, relative solvent accessibility (RSA), and confidence values (pLDDT).

#### Mapping of Epitope Scores to B-Factors

To enable three-dimensional visualization, DiscoTope scores were programmatically mapped to the B-factor field of the corresponding PDB coordinate files. A custom Python script was developed to (i) parse .csv outputs, (ii) match residues to their structural counterparts in the PDB, and (iii) overwrite the B-factor field with normalized DiscoTope values. Only residues present in both the prediction table and the structural file were annotated. This approach allowed downstream molecular viewers to interpret epitope propensities as temperature-factor–like values without altering the structural coordinates.

#### PyMOL Visualization and Selective Coloring

Visualization and figure preparation were performed using PyMOL 2.0. All structures were initially rendered in cartoon representation and colored uniformly in light gray to emphasize highlighted regions. Residues with DiscoTope scores below 0.35 were displayed using a blue–white–red gradient proportional to their score distribution. Residues with scores 0.35, considered higher-confidence epitope candidates, were isolated as a separate selection and colored using a yellow–orange–red ramp, with increasing intensity reflecting higher scores. Non-epitope regions were partially transparent to improve visual contrast. Epitope residues were also displayed as spheres to facilitate identification.

#### Automated Figure Generation

To standardize figure production across multiple experimental conditions, a PyMOL automation pipeline was implemented. For each PDB annotated with epitope scores, a .pml script was generated that loaded the structure, applied the color ramps, oriented the view, and exported publication-quality images (.png). For each object, two views were systematically generated: a front orientation and a 180° rotation along the Z-axis, saved at 2000×2000 pixels with ray tracing enabled. Outputs were stored in a structured directory (figs/) with filenames encoding experimental identifiers.

#### Comparative Visualization in R Markdown

Processed images were integrated into a reproducible R Markdown workflow. Using the cowplot package in R, images corresponding to different pressure–temperature conditions were assembled into multi-panel figures (plot_grid). Each panel was annotated with the corresponding condition (e.g., "A) 1 bar", "B) 250 bar") using draw_label. Figures were arranged in two-column layouts for balanced comparison, with residual plots centered when odd numbers of panels were present. This ensured consistency and comparability across up to 21 independent conditions.

##### Data Analysis and Statistical Evaluation

In parallel to structural visualization, DiscoTope scores and associated residue-level metrics were analyzed using Python (pandas, NumPy, SciPy, Matplotlib). For each experiment, the correlation between DiscoTope scores and relative solvent accessibility (RSA) was calculated using Pearson's correlation coefficient. This step provided an internal validation of whether epitope prediction was influenced by solvent exposure. In addition, distributions of DiscoTope scores were evaluated by plotting histograms, cumulative frequency plots, and boxplots, allowing the identification of threshold-dependent subsets of residues. Comparisons across experimental conditions (e.g., pressure or temperature series) were made by aggregating CSV results and calculating summary statistics per replicate (mean, median, interquartile range). Outliers were detected by Tukey's rule and retained for visualization, as they may correspond to structurally relevant epitope hot-spots. Statistical outputs were saved in tabular .csv form and linked to the graphical representations. By combining quantitative statistical analysis of DiscoTope predictions with three-dimensional mapping and side-by-side figure layouts, the methodology ensured both numerical rigor and visual interpretability of predicted epitope landscapes under different biophysical conditions.

#### Discontinuous Epitope predicitons

##### HHP

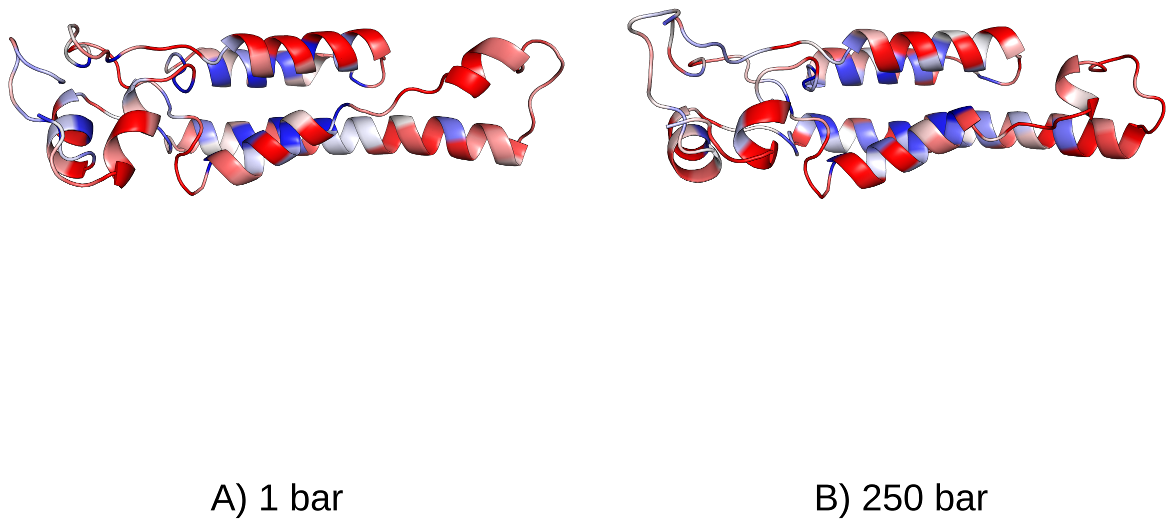

Figure 1: Epitope Preditions side-by-side - High Hydrostatic Pressure

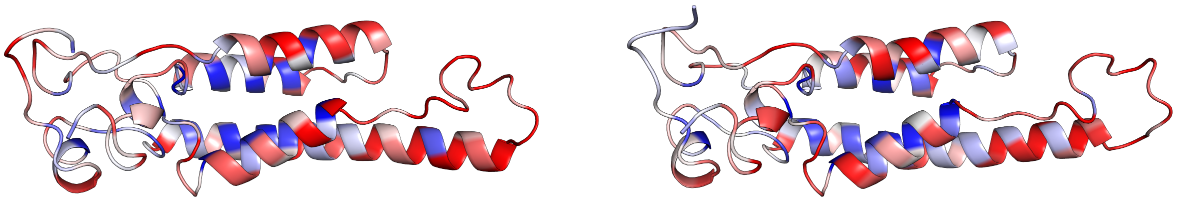

C) 500 bar D) 750 bar

Figure 2: Epitope Preditions side-by-side - High Hydrostatic Pressure

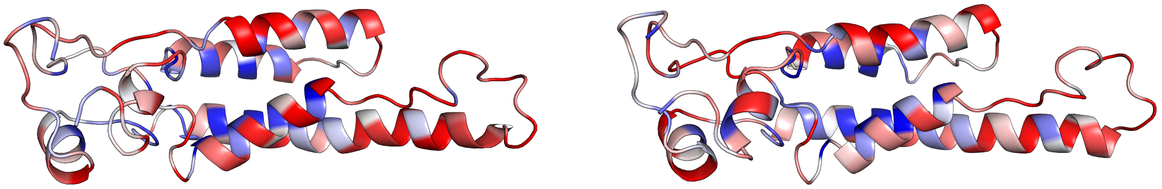

E) 1000 bar F) 1250 bar

Figure 3: Epitope Preditions side-by-side - High Hydrostatic Pressure

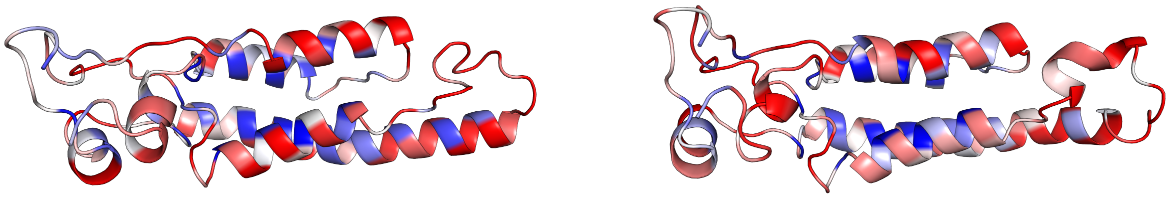

G) 1500 bar H) 1750 bar

Figure 4: Epitope Preditions side-by-side - High Hydrostatic Pressure

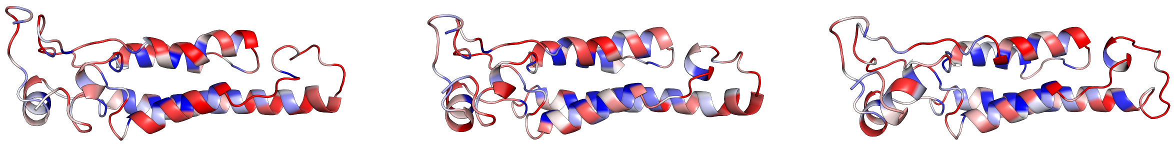

I) 2000 bar J) 2250 bar K) 2500 bar

Figure 5: Epitope Preditions side-by-side - High Hydrostatic Pressure

#### HHP and low temperature

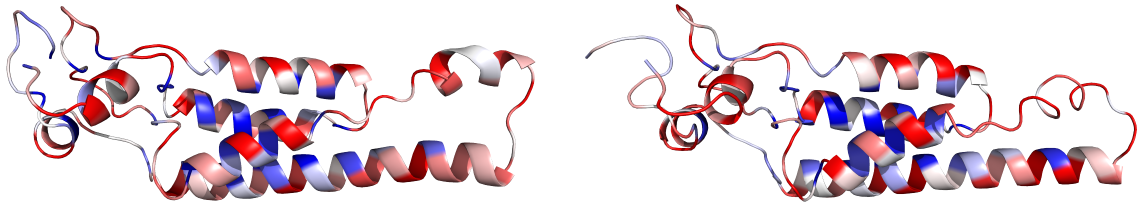

A) 1 bar B) 250 bar

Figure 6: Epitope Preditions side-by-side - High Hydrostatic Pressure and Low Temperature

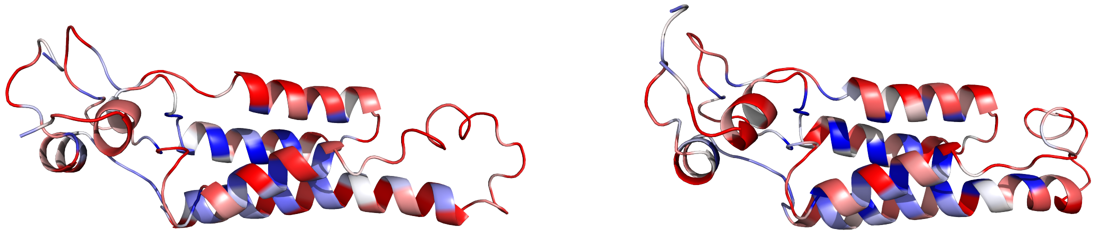

C) 500 bar D) 750 bar

Figure 7: Epitope Preditions side-by-side - High Hydrostatic Pressure and Low Temperature

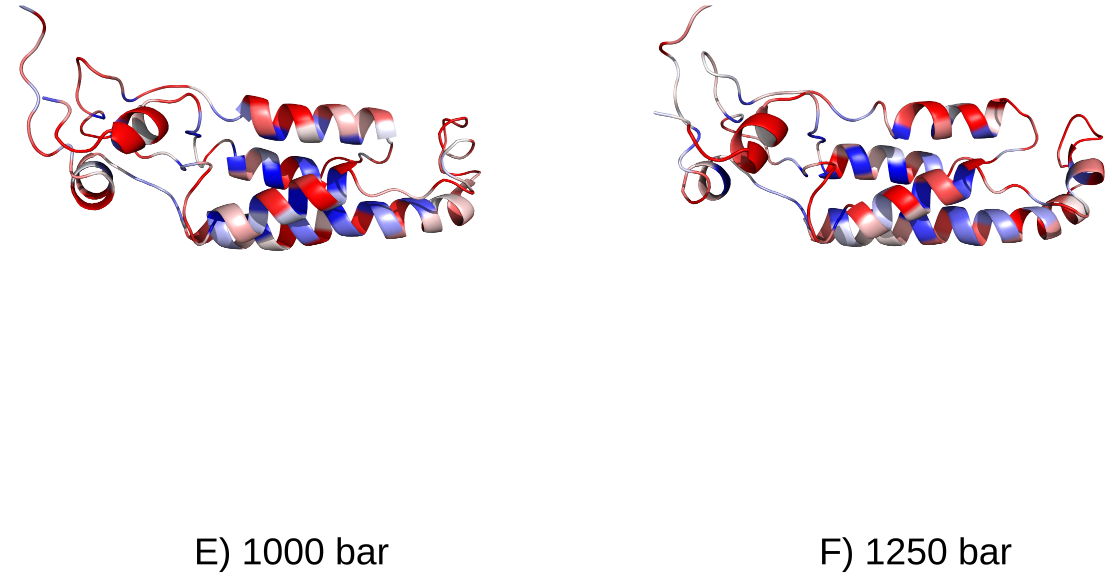

Figure 8: Epitope Preditions side-by-side - High Hydrostatic Pressure and Low Temperature

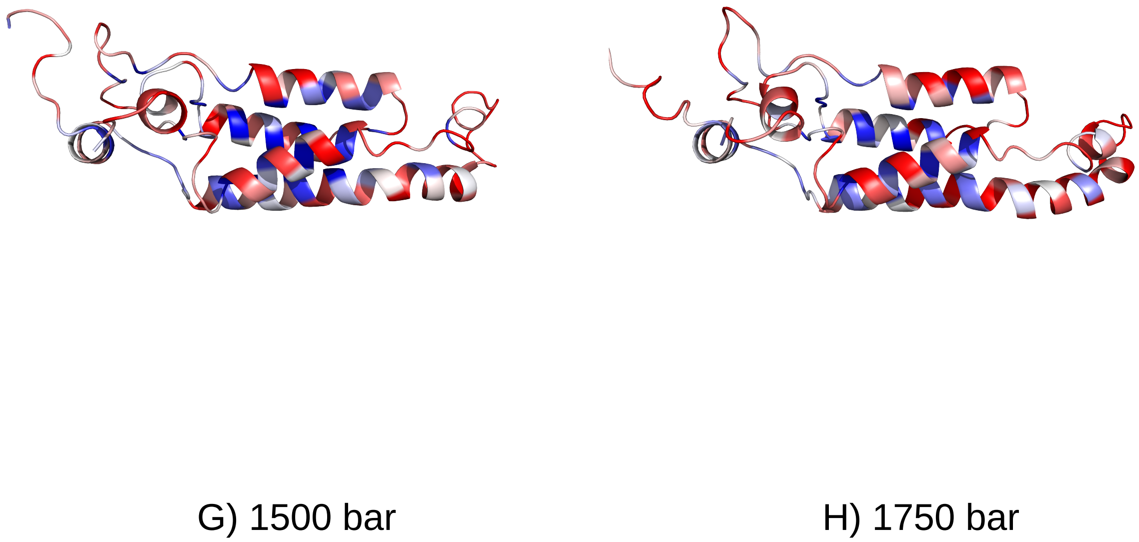

Figure 9: Epitope Preditions side-by-side - High Hydrostatic Pressure and Low Temperature

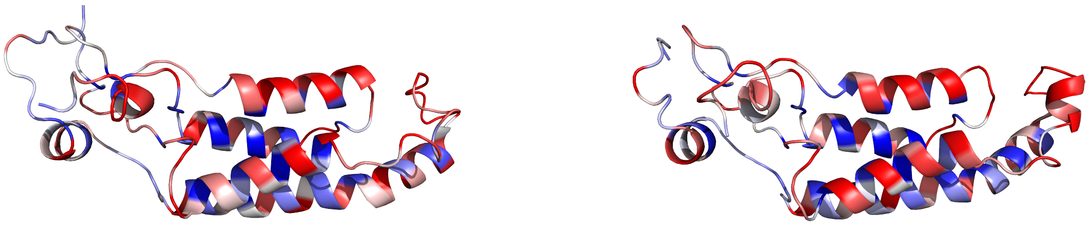

I) 2000 bar J) 2250 bar

Figure 10: Epitope Preditions side-by-side - High Hydrostatic Pressure and Low Temperature

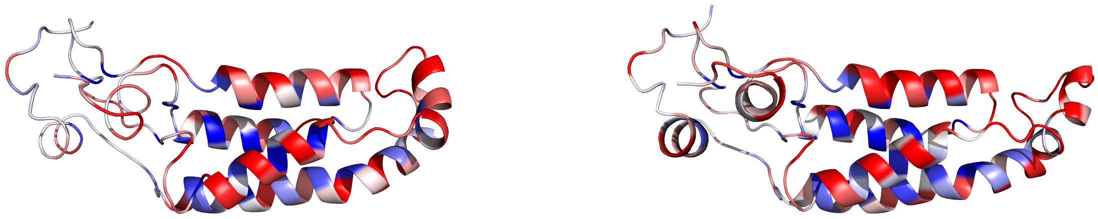

K) 2500 bar L) 2500 bar @ 300 k

Figure 11: Epitope Preditions side-by-side - High Hydrostatic Pressure and Low Temperature

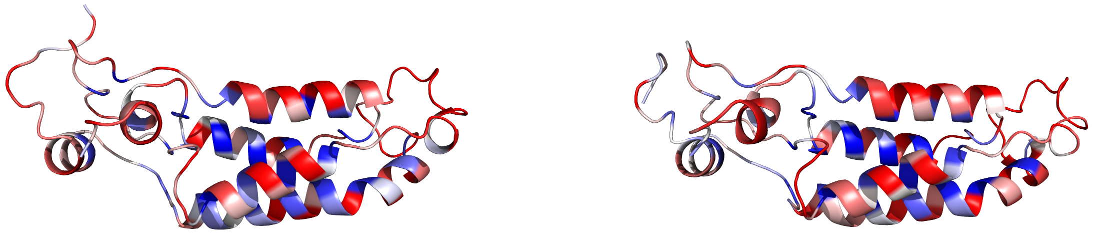

M) 2500 bar @ 295 k

N) 2500 bar @ 290 k

Figure 12: Epitope Preditions side-by-side - High Hydrostatic Pressure and Low Temperature

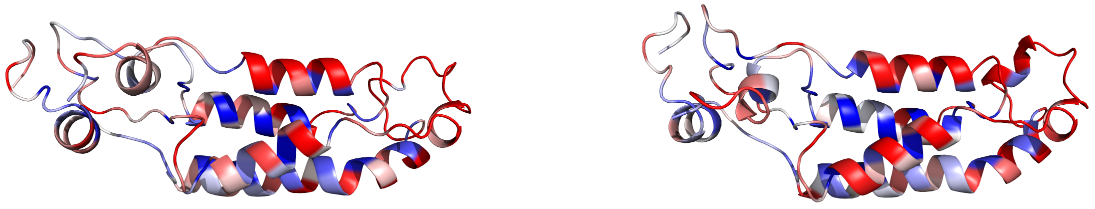

O) 2500 bar @ 285 k

P) 2500 bar @ 280 k

Figure 13: Epitope Preditions side-by-side - High Hydrostatic Pressure and Low Temperature

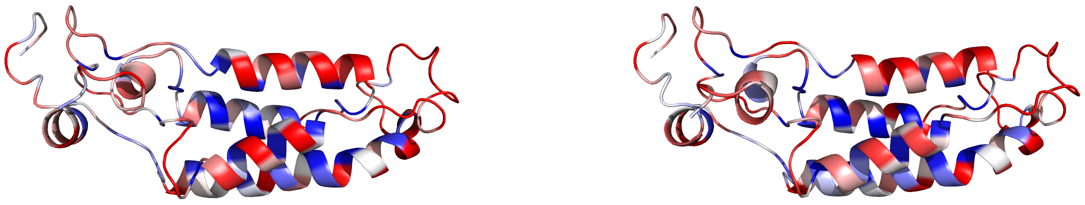

Q) 2500 bar @ 275 k R) 2500 bar @ 270 k

Figure 14: Epitope Preditions side-by-side - High Hydrostatic Pressure and Low Temperature

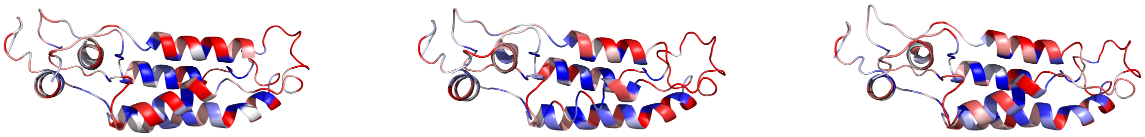

S) 2500 bar @ 265 k T) 2500 bar @ 260 k U) 2500 bar @ 255 k

Figure 15: Epitope Preditions side-by-side - High Hydrostatic Pressure and Low Temperature

### Discotope tabular results

#### HHP

Residues with DiscoTope ≥ 0.35 — md_0_1_A_discotope3.csv (n = 17)

| PDB | Chain | ResID | AA | DiscoTope | Calibrated | Epitope | RSA | pLDDT | Length | AF2 flag |
|---|---|---|---|---|---|---|---|---|---|---|
| md_0_1_A | A | 92 | R | 0.52813 | 2.15947 | TRUE | 0.64816 | 100 | 155 | 0 |
| md_0_1_A | A | 46 | R | 0.50507 | 1.99032 | TRUE | 0.58819 | 100 | 155 | 0 |
| md_0_1_A | A | 72 | Y | 0.46701 | 1.71113 | TRUE | 0.42973 | 100 | 155 | 0 |
| md_0_1_A | A | 73 | N | 0.44853 | 1.57558 | TRUE | 0.14330 | 100 | 155 | 0 |
| md_0_1_A | A | 64 | D | 0.42569 | 1.40804 | TRUE | 0.64877 | 100 | 155 | 0 |
| md_0_1_A | A | 95 | E | 0.41221 | 1.30916 | TRUE | 0.74165 | 100 | 155 | 0 |
| md_0_1_A | A | 147 | S | 0.40082 | 1.22561 | TRUE | 0.58847 | 100 | 155 | 0 |
| md_0_1_A | A | 50 | E | 0.39528 | 1.18497 | TRUE | 0.69301 | 100 | 155 | 0 |
| md_0_1_A | A | 7 | P | 0.39110 | 1.15431 | TRUE | 0.80617 | 100 | 155 | 0 |
| md_0_1_A | A | 10 | F | 0.39020 | 1.14771 | TRUE | 0.59925 | 100 | 155 | 0 |
| md_0_1_A | A | 39 | Q | 0.38660 | 1.12130 | TRUE | 0.63616 | 100 | 155 | 0 |
| md_0_1_A | A | 91 | N | 0.38358 | 1.09915 | TRUE | 0.20280 | 100 | 155 | 0 |
| md_0_1_A | A | 53 | K | 0.38293 | 1.09438 | TRUE | 0.60781 | 100 | 155 | 0 |
| md_0_1_A | A | 90 | R | 0.37395 | 1.02851 | TRUE | 0.59534 | 100 | 155 | 0 |
| md_0_1_A | A | 42 | T | 0.37386 | 1.02785 | TRUE | 0.29709 | 100 | 155 | 0 |
| md_0_1_A | A | 54 | P | 0.36824 | 0.98662 | TRUE | 0.43247 | 100 | 155 | 0 |
| md_0_1_A | A | 74 | A | 0.35988 | 0.92530 | TRUE | 0.82693 | 100 | 155 | 0 |

Figure 16: Residues with DiscoTope above the threshold of 0.35 — md-0-1-A-discotope3.csv (n = 17)

Residues with DiscoTope ≥ 0.35 — md_0_2_A_discotope3.csv (n = 25)

| PDB | Chain | ResID | AA | DiscoTope | Calibrated | Epitope | RSA | pLDDT | Length | AF2 flag |
|---|---|---|---|---|---|---|---|---|---|---|
| md_0_2_A | A | 93 | I | 0.49887 | 1.94972 | TRUE | 0.59715 | 100 | 155 | 0 |
| md_0_2_A | A | 103 | T | 0.49848 | 1.94685 | TRUE | 0.45711 | 100 | 155 | 0 |
| md_0_2_A | A | 72 | Y | 0.48512 | 1.84860 | TRUE | 0.57653 | 100 | 155 | 0 |
| md_0_2_A | A | 108 | L | 0.43786 | 1.50107 | TRUE | 0.64370 | 100 | 155 | 0 |
| md_0_2_A | A | 91 | N | 0.41601 | 1.34039 | TRUE | 0.13431 | 100 | 155 | 0 |
| md_0_2_A | A | 102 | P | 0.41440 | 1.32855 | TRUE | 0.89200 | 100 | 155 | 0 |
| md_0_2_A | A | 99 | Q | 0.40570 | 1.26457 | TRUE | 0.95334 | 100 | 155 | 0 |
| md_0_2_A | A | 90 | R | 0.40470 | 1.25722 | TRUE | 0.45870 | 100 | 155 | 0 |
| md_0_2_A | A | 104 | T | 0.39450 | 1.18221 | TRUE | 0.79505 | 100 | 155 | 0 |
| md_0_2_A | A | 77 | D | 0.38887 | 1.14081 | TRUE | 0.26426 | 100 | 155 | 0 |
| md_0_2_A | A | 81 | T | 0.38678 | 1.12544 | TRUE | 0.53012 | 100 | 155 | 0 |
| md_0_2_A | A | 109 | D | 0.38597 | 1.11948 | TRUE | 0.44171 | 100 | 155 | 0 |
| md_0_2_A | A | 100 | A | 0.38490 | 1.11161 | TRUE | 0.70926 | 100 | 155 | 0 |
| md_0_2_A | A | 15 | S | 0.37806 | 1.06131 | TRUE | 0.52621 | 100 | 155 | 0 |
| md_0_2_A | A | 112 | R | 0.37502 | 1.03896 | TRUE | 0.69976 | 100 | 155 | 0 |
| md_0_2_A | A | 92 | R | 0.37326 | 1.02602 | TRUE | 0.81143 | 100 | 155 | 0 |
| md_0_2_A | A | 74 | A | 0.37145 | 1.01271 | TRUE | 0.80167 | 100 | 155 | 0 |
| md_0_2_A | A | 126 | N | 0.36862 | 0.99189 | TRUE | 0.56531 | 100 | 155 | 0 |
| md_0_2_A | A | 46 | R | 0.36593 | 0.97211 | TRUE | 0.61600 | 100 | 155 | 0 |
| md_0_2_A | A | 123 | S | 0.35919 | 0.92255 | TRUE | 0.45534 | 100 | 155 | 0 |
| md_0_2_A | A | 39 | Q | 0.35885 | 0.92005 | TRUE | 0.69155 | 100 | 155 | 0 |
| md_0_2_A | A | 31 | L | 0.35714 | 0.90747 | TRUE | 0.35853 | 100 | 155 | 0 |
| md_0_2_A | A | 37 | T | 0.35476 | 0.88997 | FALSE | 0.63131 | 100 | 155 | 0 |
| md_0_2_A | A | 8 | S | 0.35377 | 0.88269 | FALSE | 0.52364 | 100 | 155 | 0 |
| md_0_2_A | A | 143 | S | 0.35357 | 0.88122 | FALSE | 0.36643 | 100 | 155 | 0 |

Figure 17: Residues with DiscoTope above the threshold of 0.35 — md-0-2-A-discotope3.csv (n = 25)

Residues with DiscoTope ≥ 0.35 — md_0_3_A_discotope3.csv (n = 23)

| PDB | Chain | ResID | AA | DiscoTope | Calibrated | Epitope | RSA | pLDDT | Length | AF2 flag |
|---|---|---|---|---|---|---|---|---|---|---|
| md_0_3_A | A | 112 | R | 0.53664 | 2.15527 | TRUE | 0.76469 | 100 | 155 | 0 |
| md_0_3_A | A | 99 | Q | 0.52612 | 2.08042 | TRUE | 0.81277 | 100 | 155 | 0 |
| md_0_3_A | A | 46 | R | 0.46022 | 1.61152 | TRUE | 0.60281 | 100 | 155 | 0 |
| md_0_3_A | A | 93 | I | 0.45997 | 1.60974 | TRUE | 0.88079 | 100 | 155 | 0 |
| md_0_3_A | A | 111 | T | 0.44858 | 1.52869 | TRUE | 0.48502 | 100 | 155 | 0 |
| md_0_3_A | A | 72 | Y | 0.41166 | 1.26599 | TRUE | 0.56827 | 100 | 155 | 0 |
| md_0_3_A | A | 92 | R | 0.40948 | 1.25048 | TRUE | 0.71155 | 100 | 155 | 0 |
| md_0_3_A | A | 96 | V | 0.39946 | 1.17919 | TRUE | 0.86230 | 100 | 155 | 0 |
| md_0_3_A | A | 97 | E | 0.39478 | 1.14589 | TRUE | 0.60762 | 100 | 155 | 0 |
| md_0_3_A | A | 109 | D | 0.39074 | 1.11714 | TRUE | 0.40563 | 100 | 155 | 0 |
| md_0_3_A | A | 102 | P | 0.38175 | 1.05317 | TRUE | 0.93103 | 100 | 155 | 0 |
| md_0_3_A | A | 122 | R | 0.37812 | 1.02734 | TRUE | 0.38897 | 100 | 155 | 0 |
| md_0_3_A | A | 115 | D | 0.37779 | 1.02499 | TRUE | 0.47676 | 100 | 155 | 0 |
| md_0_3_A | A | 91 | N | 0.37247 | 0.98714 | TRUE | 0.10493 | 100 | 155 | 0 |
| md_0_3_A | A | 88 | D | 0.37200 | 0.98380 | TRUE | 0.57465 | 100 | 155 | 0 |
| md_0_3_A | A | 123 | S | 0.36921 | 0.96394 | TRUE | 0.42498 | 100 | 155 | 0 |
| md_0_3_A | A | 108 | L | 0.36699 | 0.94815 | TRUE | 0.73677 | 100 | 155 | 0 |
| md_0_3_A | A | 134 | R | 0.36582 | 0.93982 | TRUE | 0.59791 | 100 | 155 | 0 |
| md_0_3_A | A | 54 | P | 0.36493 | 0.93349 | TRUE | 0.59188 | 100 | 155 | 0 |
| md_0_3_A | A | 29 | N | 0.36402 | 0.92702 | TRUE | 0.52502 | 100 | 155 | 0 |
| md_0_3_A | A | 33 | N | 0.35615 | 0.87102 | FALSE | 0.51885 | 100 | 155 | 0 |
| md_0_3_A | A | 105 | A | 0.35538 | 0.86554 | FALSE | 0.74988 | 100 | 155 | 0 |
| md_0_3_A | A | 103 | T | 0.35198 | 0.84135 | FALSE | 0.51201 | 100 | 155 | 0 |

Figure 18: Residues with DiscoTope above the threshold of 0.35 — md-0-3-A-discotope3.csv (n = 23)

Residues with DiscoTope ≥ 0.35 — md_0_4_A_discotope3.csv (n = 12)

| PDB | Chain | ResID | AA | DiscoTope | Calibrated | Epitope | RSA | pLDDT | Length | AF2 flag |
|---|---|---|---|---|---|---|---|---|---|---|
| md_0_4_A | A | 46 | R | 0.47236 | 1.87992 | TRUE | 0.48240 | 100 | 155 | 0 |
| md_0_4_A | A | 115 | D | 0.45979 | 1.78089 | TRUE | 0.53223 | 100 | 155 | 0 |
| md_0_4_A | A | 126 | N | 0.42662 | 1.51957 | TRUE | 0.57599 | 100 | 155 | 0 |
| md_0_4_A | A | 108 | L | 0.41829 | 1.45394 | TRUE | 0.61249 | 100 | 155 | 0 |
| md_0_4_A | A | 54 | P | 0.41426 | 1.42219 | TRUE | 0.68942 | 100 | 155 | 0 |
| md_0_4_A | A | 112 | R | 0.41224 | 1.40628 | TRUE | 0.60236 | 100 | 155 | 0 |
| md_0_4_A | A | 104 | T | 0.40393 | 1.34081 | TRUE | 0.99249 | 100 | 155 | 0 |
| md_0_4_A | A | 72 | Y | 0.40289 | 1.33262 | TRUE | 0.48711 | 100 | 155 | 0 |
| md_0_4_A | A | 99 | Q | 0.40059 | 1.31450 | TRUE | 0.86071 | 100 | 155 | 0 |
| md_0_4_A | A | 93 | I | 0.38341 | 1.17915 | TRUE | 0.67923 | 100 | 155 | 0 |
| md_0_4_A | A | 32 | G | 0.37094 | 1.08091 | TRUE | 0.76503 | 100 | 155 | 0 |
| md_0_4_A | A | 111 | T | 0.35568 | 0.96069 | TRUE | 0.48816 | 100 | 155 | 0 |

Figure 19: Residues with DiscoTope above the threshold of 0.35 — md-0-4A-discotope3.csv (n = 12)

Residues with DiscoTope ≥ 0.35 — md_0_5_A_discotope3.csv (n = 20)

| PDB | Chain | ResID | AA | DiscoTope | Calibrated | Epitope | RSA | pLDDT | Length | AF2 flag |
|---|---|---|---|---|---|---|---|---|---|---|
| md_0_5_A | A | 112 | R | 0.48284 | 1.82296 | TRUE | 0.58276 | 100 | 155 | 0 |
| md_0_5_A | A | 93 | I | 0.47997 | 1.80196 | TRUE | 0.83083 | 100 | 155 | 0 |
| md_0_5_A | A | 92 | R | 0.46760 | 1.71143 | TRUE | 0.75455 | 100 | 155 | 0 |
| md_0_5_A | A | 103 | T | 0.46001 | 1.65589 | TRUE | 0.62788 | 100 | 155 | 0 |
| md_0_5_A | A | 99 | Q | 0.45188 | 1.59639 | TRUE | 0.73454 | 100 | 155 | 0 |
| md_0_5_A | A | 46 | R | 0.44895 | 1.57495 | TRUE | 0.62748 | 100 | 155 | 0 |
| md_0_5_A | A | 54 | P | 0.43657 | 1.48435 | TRUE | 0.69158 | 100 | 155 | 0 |
| md_0_5_A | A | 72 | Y | 0.43490 | 1.47213 | TRUE | 0.48299 | 100 | 155 | 0 |
| md_0_5_A | A | 49 | S | 0.42675 | 1.41249 | TRUE | 0.30841 | 100 | 155 | 0 |
| md_0_5_A | A | 108 | L | 0.41937 | 1.35848 | TRUE | 0.78428 | 100 | 155 | 0 |
| md_0_5_A | A | 55 | S | 0.41442 | 1.32226 | TRUE | 0.28058 | 100 | 155 | 0 |
| md_0_5_A | A | 88 | D | 0.39209 | 1.15884 | TRUE | 0.52195 | 100 | 155 | 0 |
| md_0_5_A | A | 147 | S | 0.38206 | 1.08544 | TRUE | 0.54112 | 100 | 155 | 0 |
| md_0_5_A | A | 123 | S | 0.37755 | 1.05244 | TRUE | 0.47171 | 100 | 155 | 0 |
| md_0_5_A | A | 74 | A | 0.37738 | 1.05120 | TRUE | 0.85232 | 100 | 155 | 0 |
| md_0_5_A | A | 90 | R | 0.36949 | 0.99346 | TRUE | 0.50007 | 100 | 155 | 0 |
| md_0_5_A | A | 115 | D | 0.36251 | 0.94238 | TRUE | 0.44420 | 100 | 155 | 0 |
| md_0_5_A | A | 85 | G | 0.36183 | 0.93740 | TRUE | 0.50924 | 100 | 155 | 0 |
| md_0_5_A | A | 102 | P | 0.36168 | 0.93630 | TRUE | 0.96794 | 100 | 155 | 0 |
| md_0_5_A | A | 50 | E | 0.35993 | 0.92349 | TRUE | 0.56567 | 100 | 155 | 0 |

Figure 20: Residues with DiscoTope above the threshold of 0.35 — md-0-5-A-discotope3.csv (n = 20)

Residues with DiscoTope ≥ 0.35 — md_0_6_A_discotope3.csv (n = 17)

| PDB | Chain | ResID | AA | DiscoTope | Calibrated | Epitope | RSA | pLDDT | Length | AF2 flag |
|---|---|---|---|---|---|---|---|---|---|---|
| md_0_6_A | A | 54 | P | 0.47856 | 1.86845 | TRUE | 0.63013 | 100 | 155 | 0 |
| md_0_6_A | A | 21 | I | 0.45816 | 1.71276 | TRUE | 0.80112 | 100 | 155 | 0 |
| md_0_6_A | A | 146 | S | 0.45756 | 1.70818 | TRUE | 0.78592 | 100 | 155 | 0 |
| md_0_6_A | A | 111 | T | 0.41716 | 1.39985 | TRUE | 0.43173 | 100 | 155 | 0 |
| md_0_6_A | A | 64 | D | 0.41192 | 1.35986 | TRUE | 0.70138 | 100 | 155 | 0 |
| md_0_6_A | A | 99 | Q | 0.40949 | 1.34132 | TRUE | 0.90009 | 100 | 155 | 0 |
| md_0_6_A | A | 115 | D | 0.40204 | 1.28446 | TRUE | 0.30498 | 100 | 155 | 0 |
| md_0_6_A | A | 55 | S | 0.39292 | 1.21486 | TRUE | 0.27762 | 100 | 155 | 0 |
| md_0_6_A | A | 72 | Y | 0.39113 | 1.20120 | TRUE | 0.54704 | 100 | 155 | 0 |
| md_0_6_A | A | 65 | S | 0.39075 | 1.19830 | TRUE | 0.60540 | 100 | 155 | 0 |
| md_0_6_A | A | 46 | R | 0.38888 | 1.18403 | TRUE | 0.59187 | 100 | 155 | 0 |
| md_0_6_A | A | 66 | D | 0.38858 | 1.18174 | TRUE | 0.60124 | 100 | 155 | 0 |
| md_0_6_A | A | 141 | R | 0.36289 | 0.98568 | TRUE | 0.35925 | 100 | 155 | 0 |
| md_0_6_A | A | 47 | Q | 0.35908 | 0.95660 | TRUE | 0.36491 | 100 | 155 | 0 |
| md_0_6_A | A | 38 | Q | 0.35167 | 0.90005 | TRUE | 0.55254 | 100 | 155 | 0 |
| md_0_6_A | A | 112 | R | 0.35089 | 0.89409 | FALSE | 0.51036 | 100 | 155 | 0 |
| md_0_6_A | A | 93 | I | 0.35026 | 0.88929 | FALSE | 0.82284 | 100 | 155 | 0 |

Figure 21: Residues with DiscoTope above the threshold of 0.35 — md-0-6-A-discotope3.csv (n = 17)

#### HHP and low tempeterature

Residues with DiscoTope ≥ 0.35 — md_0_7_A_discotope3.csv (n = 23)

| PDB | Chain | ResID | AA | DiscoTope | Calibrated | Epitope | RSA | pLDDT | Length | AF2 flag |
|---|---|---|---|---|---|---|---|---|---|---|
| md_0_7_A | A | 92 | R | 0.52084 | 2.02543 | TRUE | 0.64418 | 100 | 155 | 0 |
| md_0_7_A | A | 55 | S | 0.51581 | 1.98994 | TRUE | 0.38190 | 100 | 155 | 0 |
| md_0_7_A | A | 50 | E | 0.50078 | 1.88391 | TRUE | 0.47635 | 100 | 155 | 0 |
| md_0_7_A | A | 54 | P | 0.48335 | 1.76095 | TRUE | 0.60662 | 100 | 155 | 0 |
| md_0_7_A | A | 93 | I | 0.45110 | 1.53343 | TRUE | 0.80969 | 100 | 155 | 0 |
| md_0_7_A | A | 46 | R | 0.44027 | 1.45703 | TRUE | 0.50047 | 100 | 155 | 0 |
| md_0_7_A | A | 103 | T | 0.43819 | 1.44236 | TRUE | 0.51374 | 100 | 155 | 0 |
| md_0_7_A | A | 123 | S | 0.43463 | 1.41724 | TRUE | 0.47936 | 100 | 155 | 0 |
| md_0_7_A | A | 65 | S | 0.42396 | 1.34197 | TRUE | 0.42979 | 100 | 155 | 0 |
| md_0_7_A | A | 81 | T | 0.39960 | 1.17012 | TRUE | 0.51372 | 100 | 155 | 0 |
| md_0_7_A | A | 106 | E | 0.38250 | 1.04948 | TRUE | 0.45887 | 100 | 155 | 0 |
| md_0_7_A | A | 100 | A | 0.37375 | 0.98775 | TRUE | 0.78383 | 100 | 155 | 0 |
| md_0_7_A | A | 72 | Y | 0.37345 | 0.98564 | TRUE | 0.47759 | 100 | 155 | 0 |
| md_0_7_A | A | 73 | N | 0.37113 | 0.96927 | TRUE | 0.11515 | 100 | 155 | 0 |
| md_0_7_A | A | 78 | P | 0.37031 | 0.96348 | TRUE | 0.55117 | 100 | 155 | 0 |
| md_0_7_A | A | 64 | D | 0.36916 | 0.95537 | TRUE | 0.74232 | 100 | 155 | 0 |
| md_0_7_A | A | 39 | Q | 0.35878 | 0.88214 | FALSE | 0.77288 | 100 | 155 | 0 |
| md_0_7_A | A | 97 | E | 0.35875 | 0.88193 | FALSE | 0.33216 | 100 | 155 | 0 |
| md_0_7_A | A | 99 | Q | 0.35873 | 0.88179 | FALSE | 0.85098 | 100 | 155 | 0 |
| md_0_7_A | A | 66 | D | 0.35729 | 0.87163 | FALSE | 0.73958 | 100 | 155 | 0 |
| md_0_7_A | A | 109 | D | 0.35394 | 0.84800 | FALSE | 0.52453 | 100 | 155 | 0 |
| md_0_7_A | A | 147 | S | 0.35085 | 0.82620 | FALSE | 0.50368 | 100 | 155 | 0 |
| md_0_7_A | A | 53 | K | 0.35074 | 0.82542 | FALSE | 0.62327 | 100 | 155 | 0 |

Figure 22: Residues with DiscoTope above the threshold of 0.35 — md-0-7-A-discotope3.csv (n = 23)

Residues with DiscoTope ≥ 0.35 — md_0_8_A_discotope3.csv (n = 20)

| PDB | Chain | ResID | AA | DiscoTope | Calibrated | Epitope | RSA | pLDDT | Length | AF2 flag |
|---|---|---|---|---|---|---|---|---|---|---|
| md_0_8_A | A | 92 | R | 0.48539 | 1.80218 | TRUE | 0.51860 | 100 | 155 | 0 |
| md_0_8_A | A | 54 | P | 0.48449 | 1.79573 | TRUE | 0.47587 | 100 | 155 | 0 |
| md_0_8_A | A | 64 | D | 0.48224 | 1.77962 | TRUE | 0.60834 | 100 | 155 | 0 |
| md_0_8_A | A | 103 | T | 0.45223 | 1.56471 | TRUE | 0.65013 | 100 | 155 | 0 |
| md_0_8_A | A | 93 | I | 0.44116 | 1.48543 | TRUE | 0.60411 | 100 | 155 | 0 |
| md_0_8_A | A | 72 | Y | 0.43917 | 1.47118 | TRUE | 0.52489 | 100 | 155 | 0 |
| md_0_8_A | A | 142 | S | 0.43360 | 1.43129 | TRUE | 0.69077 | 100 | 155 | 0 |
| md_0_8_A | A | 99 | Q | 0.43015 | 1.40659 | TRUE | 0.86055 | 100 | 155 | 0 |
| md_0_8_A | A | 112 | R | 0.42076 | 1.33934 | TRUE | 0.52080 | 100 | 155 | 0 |
| md_0_8_A | A | 100 | A | 0.41916 | 1.32788 | TRUE | 0.78550 | 100 | 155 | 0 |
| md_0_8_A | A | 141 | R | 0.39655 | 1.16596 | TRUE | 0.41878 | 100 | 155 | 0 |
| md_0_8_A | A | 15 | S | 0.38875 | 1.11011 | TRUE | 0.53449 | 100 | 155 | 0 |
| md_0_8_A | A | 102 | P | 0.37735 | 1.02847 | TRUE | 0.85309 | 100 | 155 | 0 |
| md_0_8_A | A | 46 | R | 0.37216 | 0.99130 | TRUE | 0.63383 | 100 | 155 | 0 |
| md_0_8_A | A | 65 | S | 0.37007 | 0.97633 | TRUE | 0.47786 | 100 | 155 | 0 |
| md_0_8_A | A | 67 | F | 0.36663 | 0.95170 | TRUE | 0.35241 | 100 | 155 | 0 |
| md_0_8_A | A | 12 | F | 0.36301 | 0.92577 | TRUE | 0.20035 | 100 | 155 | 0 |
| md_0_8_A | A | 108 | L | 0.35511 | 0.86920 | FALSE | 0.94120 | 100 | 155 | 0 |
| md_0_8_A | A | 63 | P | 0.35102 | 0.83991 | FALSE | 0.30816 | 100 | 155 | 0 |
| md_0_8_A | A | 73 | N | 0.35069 | 0.83754 | FALSE | 0.17021 | 100 | 155 | 0 |

Figure 23: Residues with DiscoTope above the threshold of 0.35 — md-0-8-A-discotope3.csv (n = 20)

Residues with DiscoTope ≥ 0.35 — md_0_9_A_discotope3.csv (n = 22)

| PDB | Chain | ResID | AA | DiscoTope | Calibrated | Epitope | RSA | pLDDT | Length | AF2 flag |
|---|---|---|---|---|---|---|---|---|---|---|
| md_0_9_A | A | 92 | R | 0.61171 | 2.69769 | TRUE | 0.63313 | 100 | 155 | 0 |
| md_0_9_A | A | 103 | T | 0.53001 | 2.11458 | TRUE | 0.46542 | 100 | 155 | 0 |
| md_0_9_A | A | 93 | I | 0.52303 | 2.06476 | TRUE | 0.69592 | 100 | 155 | 0 |
| md_0_9_A | A | 46 | R | 0.51146 | 1.98218 | TRUE | 0.64099 | 100 | 155 | 0 |
| md_0_9_A | A | 91 | N | 0.48739 | 1.81039 | TRUE | 0.24956 | 100 | 155 | 0 |
| md_0_9_A | A | 38 | Q | 0.46494 | 1.65015 | TRUE | 0.32972 | 100 | 155 | 0 |
| md_0_9_A | A | 15 | S | 0.43810 | 1.45859 | TRUE | 0.51228 | 100 | 155 | 0 |
| md_0_9_A | A | 108 | L | 0.42569 | 1.37002 | TRUE | 0.65616 | 100 | 155 | 0 |
| md_0_9_A | A | 102 | P | 0.41050 | 1.26160 | TRUE | 0.92515 | 100 | 155 | 0 |
| md_0_9_A | A | 99 | Q | 0.41018 | 1.25932 | TRUE | 0.90499 | 100 | 155 | 0 |
| md_0_9_A | A | 116 | D | 0.38808 | 1.10159 | TRUE | 0.56881 | 100 | 155 | 0 |
| md_0_9_A | A | 85 | G | 0.38438 | 1.07518 | TRUE | 0.42006 | 100 | 155 | 0 |
| md_0_9_A | A | 61 | R | 0.37291 | 0.99331 | TRUE | 0.38115 | 100 | 155 | 0 |
| md_0_9_A | A | 88 | D | 0.36888 | 0.96455 | TRUE | 0.56971 | 100 | 155 | 0 |
| md_0_9_A | A | 39 | Q | 0.36743 | 0.95420 | TRUE | 0.82230 | 100 | 155 | 0 |
| md_0_9_A | A | 64 | D | 0.36576 | 0.94228 | TRUE | 0.68628 | 100 | 155 | 0 |
| md_0_9_A | A | 153 | T | 0.36217 | 0.91666 | TRUE | 0.44634 | 100 | 155 | 0 |
| md_0_9_A | A | 89 | T | 0.36130 | 0.91045 | TRUE | 0.64323 | 100 | 155 | 0 |
| md_0_9_A | A | 90 | R | 0.36033 | 0.90353 | TRUE | 0.28051 | 100 | 155 | 0 |
| md_0_9_A | A | 72 | Y | 0.35857 | 0.89097 | FALSE | 0.47130 | 100 | 155 | 0 |
| md_0_9_A | A | 122 | R | 0.35749 | 0.88326 | FALSE | 0.28378 | 100 | 155 | 0 |
| md_0_9_A | A | 59 | T | 0.35245 | 0.84729 | FALSE | 0.56231 | 100 | 155 | 0 |

Figure 24: Residues with DiscoTope above the threshold of 0.35 — md-0-9-A-discotope3.csv (n = 22)

Residues with DiscoTope ≥ 0.35 — md_0_10_A_discotope3.csv (n = 13)

| PDB | Chain | ResID | AA | DiscoTope | Calibrated | Epitope | RSA | pLDDT | Length | AF2 flag |
|---|---|---|---|---|---|---|---|---|---|---|
| md_0_10_A | A | 91 | N | 0.47046 | 1.83359 | TRUE | 0.29035 | 100 | 155 | 0 |
| md_0_10_A | A | 11 | V | 0.42690 | 1.49618 | TRUE | 0.51784 | 100 | 155 | 0 |
| md_0_10_A | A | 15 | S | 0.42618 | 1.49061 | TRUE | 0.47874 | 100 | 155 | 0 |
| md_0_10_A | A | 93 | I | 0.41611 | 1.41261 | TRUE | 0.47725 | 100 | 155 | 0 |
| md_0_10_A | A | 102 | P | 0.41601 | 1.41183 | TRUE | 0.83486 | 100 | 155 | 0 |
| md_0_10_A | A | 108 | L | 0.41463 | 1.40114 | TRUE | 0.91160 | 100 | 155 | 0 |
| md_0_10_A | A | 54 | P | 0.38117 | 1.14197 | TRUE | 0.49052 | 100 | 155 | 0 |
| md_0_10_A | A | 103 | T | 0.37965 | 1.13020 | TRUE | 0.45148 | 100 | 155 | 0 |
| md_0_10_A | A | 65 | S | 0.37752 | 1.11370 | TRUE | 0.50500 | 100 | 155 | 0 |
| md_0_10_A | A | 116 | D | 0.36085 | 0.98458 | TRUE | 0.56682 | 100 | 155 | 0 |
| md_0_10_A | A | 7 | P | 0.35708 | 0.95538 | TRUE | 0.81965 | 100 | 155 | 0 |
| md_0_10_A | A | 72 | Y | 0.35297 | 0.92354 | TRUE | 0.58386 | 100 | 155 | 0 |
| md_0_10_A | A | 147 | S | 0.35208 | 0.91665 | TRUE | 0.45864 | 100 | 155 | 0 |

Figure 25: Residues with DiscoTope above the threshold of 0.35 — md-0-10-A-discotope3.csv (n = 13)

Residues with DiscoTope ≥ 0.35 — md_0_11_A_discotope3.csv (n = 20)

| PDB | Chain | ResID | AA | DiscoTope | Calibrated | Epitope | RSA | pLDDT | Length | AF2 flag |
|---|---|---|---|---|---|---|---|---|---|---|
| md_0_11_A | A | 55 | S | 0.48808 | 1.90204 | TRUE | 0.24249 | 100 | 155 | 0 |
| md_0_11_A | A | 102 | P | 0.48735 | 1.89658 | TRUE | 0.76541 | 100 | 155 | 0 |
| md_0_11_A | A | 99 | Q | 0.48689 | 1.89314 | TRUE | 0.86205 | 100 | 155 | 0 |
| md_0_11_A | A | 91 | N | 0.45048 | 1.62086 | TRUE | 0.15310 | 100 | 155 | 0 |
| md_0_11_A | A | 92 | R | 0.43107 | 1.47570 | TRUE | 0.59247 | 100 | 155 | 0 |
| md_0_11_A | A | 93 | I | 0.42670 | 1.44302 | TRUE | 0.21978 | 100 | 155 | 0 |
| md_0_11_A | A | 64 | D | 0.42116 | 1.40160 | TRUE | 0.71078 | 100 | 155 | 0 |
| md_0_11_A | A | 103 | T | 0.41853 | 1.38193 | TRUE | 0.41994 | 100 | 155 | 0 |
| md_0_11_A | A | 88 | D | 0.41457 | 1.35231 | TRUE | 0.60095 | 100 | 155 | 0 |
| md_0_11_A | A | 46 | R | 0.41352 | 1.34446 | TRUE | 0.65749 | 100 | 155 | 0 |
| md_0_11_A | A | 54 | P | 0.40980 | 1.31664 | TRUE | 0.51406 | 100 | 155 | 0 |
| md_0_11_A | A | 38 | Q | 0.39539 | 1.20888 | TRUE | 0.27712 | 100 | 155 | 0 |
| md_0_11_A | A | 90 | R | 0.39031 | 1.17089 | TRUE | 0.48600 | 100 | 155 | 0 |
| md_0_11_A | A | 98 | N | 0.38749 | 1.14980 | TRUE | 0.90763 | 100 | 155 | 0 |
| md_0_11_A | A | 96 | V | 0.38402 | 1.12385 | TRUE | 0.58623 | 100 | 155 | 0 |
| md_0_11_A | A | 72 | Y | 0.38069 | 1.09895 | TRUE | 0.58485 | 100 | 155 | 0 |
| md_0_11_A | A | 104 | T | 0.38042 | 1.09693 | TRUE | 0.95742 | 100 | 155 | 0 |
| md_0_11_A | A | 97 | E | 0.37165 | 1.03135 | TRUE | 0.44173 | 100 | 155 | 0 |
| md_0_11_A | A | 112 | R | 0.37012 | 1.01991 | TRUE | 0.46176 | 100 | 155 | 0 |
| md_0_11_A | A | 50 | E | 0.36791 | 1.00338 | TRUE | 0.54313 | 100 | 155 | 0 |

Figure 26: Residues with DiscoTope above the threshold of 0.35 — md-0-11-A-discotope3.csv (n = 20)

Residues with DiscoTope ≥ 0.35 — md_0_1_A_discotope3.csv (n = 17)

| PDB | Chain | ResID | AA | DiscoTope | Calibrated | Epitope | RSA | pLDDT | Length | AF2 flag |
|---|---|---|---|---|---|---|---|---|---|---|
| md_0_1_A | A | 72 | Y | 0.53656 | 2.28066 | TRUE | 0.42510 | 100 | 155 | 0 |
| md_0_1_A | A | 46 | R | 0.51568 | 2.12340 | TRUE | 0.59664 | 100 | 155 | 0 |
| md_0_1_A | A | 147 | S | 0.41198 | 1.34241 | TRUE | 0.49080 | 100 | 155 | 0 |
| md_0_1_A | A | 92 | R | 0.40762 | 1.30957 | TRUE | 0.63739 | 100 | 155 | 0 |
| md_0_1_A | A | 54 | P | 0.40640 | 1.30038 | TRUE | 0.47444 | 100 | 155 | 0 |
| md_0_1_A | A | 111 | T | 0.40476 | 1.28803 | TRUE | 0.51017 | 100 | 155 | 0 |
| md_0_1_A | A | 85 | G | 0.39070 | 1.18214 | TRUE | 0.57838 | 100 | 155 | 0 |
| md_0_1_A | A | 10 | F | 0.38750 | 1.15804 | TRUE | 0.62927 | 100 | 155 | 0 |
| md_0_1_A | A | 112 | R | 0.37948 | 1.09764 | TRUE | 0.46038 | 100 | 155 | 0 |
| md_0_1_A | A | 53 | K | 0.37844 | 1.08981 | TRUE | 0.59923 | 100 | 155 | 0 |
| md_0_1_A | A | 39 | Q | 0.37410 | 1.05712 | TRUE | 0.62076 | 100 | 155 | 0 |
| md_0_1_A | A | 64 | D | 0.36856 | 1.01540 | TRUE | 0.68842 | 100 | 155 | 0 |
| md_0_1_A | A | 15 | S | 0.36829 | 1.01336 | TRUE | 0.48169 | 100 | 155 | 0 |
| md_0_1_A | A | 6 | T | 0.36445 | 0.98444 | TRUE | 0.54451 | 100 | 155 | 0 |
| md_0_1_A | A | 123 | S | 0.36199 | 0.96592 | TRUE | 0.41236 | 100 | 155 | 0 |
| md_0_1_A | A | 14 | S | 0.35895 | 0.94302 | TRUE | 0.32119 | 100 | 155 | 0 |
| md_0_1_A | A | 21 | I | 0.35686 | 0.92728 | TRUE | 0.72317 | 100 | 155 | 0 |

Figure 27: Residues with DiscoTope above the threshold of 0.35 — md-0-1-A-discotope3.csv (n = 17)

Residues with DiscoTope ≥ 0.35 — md_0_2_A_discotope3.csv (n = 22)

| PDB | Chain | ResID | AA | DiscoTope | Calibrated | Epitope | RSA | pLDDT | Length | AF2 flag |
|---|---|---|---|---|---|---|---|---|---|---|
| md_0_2_A | A | 72 | Y | 0.48491 | 1.78270 | TRUE | 0.48138 | 100 | 155 | 0 |
| md_0_2_A | A | 8 | S | 0.47443 | 1.70832 | TRUE | 0.67595 | 100 | 155 | 0 |
| md_0_2_A | A | 46 | R | 0.44962 | 1.53223 | TRUE | 0.55449 | 100 | 155 | 0 |
| md_0_2_A | A | 111 | T | 0.44340 | 1.48808 | TRUE | 0.59339 | 100 | 155 | 0 |
| md_0_2_A | A | 122 | R | 0.44166 | 1.47573 | TRUE | 0.54198 | 100 | 155 | 0 |
| md_0_2_A | A | 7 | P | 0.43277 | 1.41264 | TRUE | 0.94366 | 100 | 155 | 0 |
| md_0_2_A | A | 42 | T | 0.42963 | 1.39035 | TRUE | 0.50326 | 100 | 155 | 0 |
| md_0_2_A | A | 39 | Q | 0.41721 | 1.30220 | TRUE | 0.88608 | 100 | 155 | 0 |
| md_0_2_A | A | 61 | R | 0.41218 | 1.26650 | TRUE | 0.47368 | 100 | 155 | 0 |
| md_0_2_A | A | 92 | R | 0.40842 | 1.23981 | TRUE | 0.50524 | 100 | 155 | 0 |
| md_0_2_A | A | 77 | D | 0.40614 | 1.22363 | TRUE | 0.25022 | 100 | 155 | 0 |
| md_0_2_A | A | 109 | D | 0.40611 | 1.22342 | TRUE | 0.34110 | 100 | 155 | 0 |
| md_0_2_A | A | 99 | Q | 0.40564 | 1.22008 | TRUE | 0.68130 | 100 | 155 | 0 |
| md_0_2_A | A | 147 | S | 0.38995 | 1.10872 | TRUE | 0.42777 | 100 | 155 | 0 |
| md_0_2_A | A | 112 | R | 0.38916 | 1.10312 | TRUE | 0.38079 | 100 | 155 | 0 |
| md_0_2_A | A | 32 | G | 0.38902 | 1.10212 | TRUE | 0.75942 | 100 | 155 | 0 |
| md_0_2_A | A | 10 | F | 0.38786 | 1.09389 | TRUE | 0.57975 | 100 | 155 | 0 |
| md_0_2_A | A | 5 | T | 0.38705 | 1.08814 | TRUE | 0.89541 | 100 | 155 | 0 |
| md_0_2_A | A | 6 | T | 0.36748 | 0.94924 | TRUE | 0.43084 | 100 | 155 | 0 |
| md_0_2_A | A | 64 | D | 0.36607 | 0.93923 | TRUE | 0.61786 | 100 | 155 | 0 |
| md_0_2_A | A | 85 | G | 0.35677 | 0.87323 | FALSE | 0.36382 | 100 | 155 | 0 |
| md_0_2_A | A | 93 | I | 0.35196 | 0.83909 | FALSE | 0.92978 | 100 | 155 | 0 |

Figure 28: Residues with DiscoTope above the threshold of 0.35 — md-0-2-A-discotope3.csv (n = 22)

Residues with DiscoTope ≥ 0.35 — md_0_3_A_discotope3.csv (n = 21)

| PDB | Chain | ResID | AA | DiscoTope | Calibrated | Epitope | RSA | pLDDT | Length | AF2 flag |
|---|---|---|---|---|---|---|---|---|---|---|
| md_0_3_A | A | 46 | R | 0.48956 | 1.80627 | TRUE | 0.70207 | 100 | 155 | 0 |
| md_0_3_A | A | 81 | T | 0.48560 | 1.77831 | TRUE | 0.52678 | 100 | 155 | 0 |
| md_0_3_A | A | 92 | R | 0.46407 | 1.62630 | TRUE | 0.44802 | 100 | 155 | 0 |
| md_0_3_A | A | 99 | Q | 0.45770 | 1.58132 | TRUE | 0.86250 | 100 | 155 | 0 |
| md_0_3_A | A | 7 | P | 0.45143 | 1.53705 | TRUE | 0.80081 | 100 | 155 | 0 |
| md_0_3_A | A | 93 | I | 0.44223 | 1.47209 | TRUE | 0.89269 | 100 | 155 | 0 |
| md_0_3_A | A | 72 | Y | 0.41744 | 1.29706 | TRUE | 0.44171 | 100 | 155 | 0 |
| md_0_3_A | A | 54 | P | 0.40900 | 1.23747 | TRUE | 0.54410 | 100 | 155 | 0 |
| md_0_3_A | A | 112 | R | 0.40793 | 1.22991 | TRUE | 0.65364 | 100 | 155 | 0 |
| md_0_3_A | A | 71 | R | 0.40711 | 1.22413 | TRUE | 0.23334 | 100 | 155 | 0 |
| md_0_3_A | A | 98 | N | 0.40632 | 1.21855 | TRUE | 0.27942 | 100 | 155 | 0 |
| md_0_3_A | A | 36 | Q | 0.38855 | 1.09308 | TRUE | 0.52009 | 100 | 155 | 0 |
| md_0_3_A | A | 119 | V | 0.38855 | 1.09308 | TRUE | 0.57886 | 100 | 155 | 0 |
| md_0_3_A | A | 115 | D | 0.38854 | 1.09301 | TRUE | 0.40272 | 100 | 155 | 0 |
| md_0_3_A | A | 152 | W | 0.38341 | 1.05679 | TRUE | 0.86363 | 100 | 155 | 0 |
| md_0_3_A | A | 52 | W | 0.37716 | 1.01266 | TRUE | 0.22466 | 100 | 155 | 0 |
| md_0_3_A | A | 58 | V | 0.37659 | 1.00863 | TRUE | 0.68879 | 100 | 155 | 0 |
| md_0_3_A | A | 100 | A | 0.37622 | 1.00602 | TRUE | 0.72171 | 100 | 155 | 0 |
| md_0_3_A | A | 97 | E | 0.36490 | 0.92610 | TRUE | 0.52653 | 100 | 155 | 0 |
| md_0_3_A | A | 151 | V | 0.35534 | 0.85860 | FALSE | 0.84815 | 100 | 155 | 0 |
| md_0_3_A | A | 49 | S | 0.35196 | 0.83473 | FALSE | 0.61196 | 100 | 155 | 0 |

Figure 29: Residues with DiscoTope above the threshold of 0.35 — md-0-3-A-discotope3.csv (n = 21)

Residues with DiscoTope ≥ 0.35 — md_0_4_A_discotope3.csv (n = 21)

| PDB | Chain | ResID | AA | DiscoTope | Calibrated | Epitope | RSA | pLDDT | Length | AF2 flag |
|---|---|---|---|---|---|---|---|---|---|---|
| md_0_4_A | A | 92 | R | 0.49669 | 1.88454 | TRUE | 0.85693 | 100 | 155 | 0 |
| md_0_4_A | A | 6 | T | 0.45693 | 1.59959 | TRUE | 0.50128 | 100 | 155 | 0 |
| md_0_4_A | A | 10 | F | 0.43211 | 1.42171 | TRUE | 0.52851 | 100 | 155 | 0 |
| md_0_4_A | A | 97 | E | 0.40556 | 1.23143 | TRUE | 0.70839 | 100 | 155 | 0 |
| md_0_4_A | A | 46 | R | 0.40359 | 1.21731 | TRUE | 0.59103 | 100 | 155 | 0 |
| md_0_4_A | A | 88 | D | 0.39822 | 1.17883 | TRUE | 0.61402 | 100 | 155 | 0 |
| md_0_4_A | A | 108 | L | 0.39504 | 1.15604 | TRUE | 0.70674 | 100 | 155 | 0 |
| md_0_4_A | A | 72 | Y | 0.39350 | 1.14500 | TRUE | 0.39599 | 100 | 155 | 0 |
| md_0_4_A | A | 37 | T | 0.39016 | 1.12106 | TRUE | 0.26308 | 100 | 155 | 0 |
| md_0_4_A | A | 39 | Q | 0.38672 | 1.09641 | TRUE | 0.66524 | 100 | 155 | 0 |
| md_0_4_A | A | 93 | I | 0.38147 | 1.05878 | TRUE | 0.88457 | 100 | 155 | 0 |
| md_0_4_A | A | 146 | S | 0.38006 | 1.04868 | TRUE | 0.61808 | 100 | 155 | 0 |
| md_0_4_A | A | 14 | S | 0.37490 | 1.01169 | TRUE | 0.37224 | 100 | 155 | 0 |
| md_0_4_A | A | 103 | T | 0.37338 | 1.00080 | TRUE | 0.79568 | 100 | 155 | 0 |
| md_0_4_A | A | 147 | S | 0.36525 | 0.94253 | TRUE | 0.59056 | 100 | 155 | 0 |
| md_0_4_A | A | 102 | P | 0.36357 | 0.93049 | TRUE | 0.85029 | 100 | 155 | 0 |
| md_0_4_A | A | 90 | R | 0.36023 | 0.90656 | TRUE | 0.31696 | 100 | 155 | 0 |
| md_0_4_A | A | 8 | S | 0.35661 | 0.88061 | FALSE | 0.54386 | 100 | 155 | 0 |
| md_0_4_A | A | 152 | W | 0.35550 | 0.87266 | FALSE | 0.75713 | 100 | 155 | 0 |
| md_0_4_A | A | 100 | A | 0.35538 | 0.87180 | FALSE | 0.69687 | 100 | 155 | 0 |
| md_0_4_A | A | 32 | G | 0.35147 | 0.84378 | FALSE | 0.52664 | 100 | 155 | 0 |

Figure 30: Residues with DiscoTope above the threshold of 0.35 — md-0-4A-discotope3.csv (n = 21)

Residues with DiscoTope ≥ 0.35 — md_0_5_A_discotope3.csv (n = 23)

| PDB | Chain | ResID | AA | DiscoTope | Calibrated | Epitope | RSA | pLDDT | Length | AF2 flag |
|---|---|---|---|---|---|---|---|---|---|---|
| md_0_5_A | A | 147 | S | 0.49415 | 1.79855 | TRUE | 0.49100 | 100 | 155 | 0 |
| md_0_5_A | A | 72 | Y | 0.47987 | 1.69992 | TRUE | 0.41202 | 100 | 155 | 0 |
| md_0_5_A | A | 123 | S | 0.42486 | 1.32000 | TRUE | 0.45877 | 100 | 155 | 0 |
| md_0_5_A | A | 10 | F | 0.41064 | 1.22179 | TRUE | 0.54765 | 100 | 155 | 0 |
| md_0_5_A | A | 112 | R | 0.40842 | 1.20645 | TRUE | 0.56850 | 100 | 155 | 0 |
| md_0_5_A | A | 29 | N | 0.40784 | 1.20245 | TRUE | 0.65165 | 100 | 155 | 0 |
| md_0_5_A | A | 102 | P | 0.40536 | 1.18532 | TRUE | 1.01620 | 100 | 155 | 0 |
| md_0_5_A | A | 5 | T | 0.40443 | 1.17890 | TRUE | 0.73329 | 100 | 155 | 0 |
| md_0_5_A | A | 46 | R | 0.40405 | 1.17627 | TRUE | 0.57956 | 100 | 155 | 0 |
| md_0_5_A | A | 32 | G | 0.40074 | 1.15341 | TRUE | 0.75772 | 100 | 155 | 0 |
| md_0_5_A | A | 11 | V | 0.39948 | 1.14471 | TRUE | 0.52149 | 100 | 155 | 0 |
| md_0_5_A | A | 146 | S | 0.39736 | 1.13007 | TRUE | 0.73435 | 100 | 155 | 0 |
| md_0_5_A | A | 7 | P | 0.39643 | 1.12364 | TRUE | 0.85634 | 100 | 155 | 0 |
| md_0_5_A | A | 85 | G | 0.39307 | 1.10044 | TRUE | 0.39340 | 100 | 155 | 0 |
| md_0_5_A | A | 88 | D | 0.38964 | 1.07675 | TRUE | 0.58381 | 100 | 155 | 0 |
| md_0_5_A | A | 8 | S | 0.38293 | 1.03041 | TRUE | 0.52400 | 100 | 155 | 0 |
| md_0_5_A | A | 50 | E | 0.38192 | 1.02343 | TRUE | 0.68869 | 100 | 155 | 0 |
| md_0_5_A | A | 6 | T | 0.37657 | 0.98648 | TRUE | 0.46378 | 100 | 155 | 0 |
| md_0_5_A | A | 156 | P | 0.37268 | 0.95961 | TRUE | 0.86162 | 100 | 155 | 0 |
| md_0_5_A | A | 92 | R | 0.36340 | 0.89552 | FALSE | 0.74380 | 100 | 155 | 0 |
| md_0_5_A | A | 126 | N | 0.35583 | 0.84324 | FALSE | 0.50360 | 100 | 155 | 0 |
| md_0_5_A | A | 99 | Q | 0.35399 | 0.83053 | FALSE | 0.41451 | 100 | 155 | 0 |
| md_0_5_A | A | 65 | S | 0.35253 | 0.82045 | FALSE | 0.75294 | 100 | 155 | 0 |

Figure 31: Residues with DiscoTope above the threshold of 0.35 — md-0-5-A-discotope3.csv (n = 23)

Residues with DiscoTope ≥ 0.35 — md_0_6_A_discotope3.csv (n = 29)

| PDB | Chain | ResID | AA | DiscoTope | Calibrated | Epitope | RSA | pLDDT | Length | AF2 flag |
|---|---|---|---|---|---|---|---|---|---|---|
| md_0_6_A | A | 72 | Y | 0.54050 | 2.12887 | TRUE | 0.45333 | 100 | 155 | 0 |
| md_0_6_A | A | 7 | P | 0.48016 | 1.71012 | TRUE | 0.81149 | 100 | 155 | 0 |
| md_0_6_A | A | 34 | Q | 0.46837 | 1.62830 | TRUE | 0.60123 | 100 | 155 | 0 |
| md_0_6_A | A | 99 | Q | 0.44671 | 1.47799 | TRUE | 0.43703 | 100 | 155 | 0 |
| md_0_6_A | A | 6 | T | 0.44561 | 1.47035 | TRUE | 0.42091 | 100 | 155 | 0 |
| md_0_6_A | A | 46 | R | 0.43820 | 1.41893 | TRUE | 0.61055 | 100 | 155 | 0 |
| md_0_6_A | A | 108 | L | 0.43529 | 1.39873 | TRUE | 0.74692 | 100 | 155 | 0 |
| md_0_6_A | A | 10 | F | 0.43403 | 1.38999 | TRUE | 0.51185 | 100 | 155 | 0 |
| md_0_6_A | A | 115 | D | 0.42921 | 1.35654 | TRUE | 0.26367 | 100 | 155 | 0 |
| md_0_6_A | A | 102 | P | 0.42473 | 1.32545 | TRUE | 0.99251 | 100 | 155 | 0 |
| md_0_6_A | A | 105 | A | 0.42376 | 1.31872 | TRUE | 0.84614 | 100 | 155 | 0 |
| md_0_6_A | A | 103 | T | 0.42223 | 1.30810 | TRUE | 0.48935 | 100 | 155 | 0 |
| md_0_6_A | A | 106 | E | 0.41807 | 1.27923 | TRUE | 0.72817 | 100 | 155 | 0 |
| md_0_6_A | A | 101 | N | 0.41676 | 1.27014 | TRUE | 0.25296 | 100 | 155 | 0 |
| md_0_6_A | A | 8 | S | 0.41378 | 1.24946 | TRUE | 0.51854 | 100 | 155 | 0 |
| md_0_6_A | A | 81 | T | 0.41070 | 1.22809 | TRUE | 0.58696 | 100 | 155 | 0 |
| md_0_6_A | A | 88 | D | 0.40832 | 1.21157 | TRUE | 0.65992 | 100 | 155 | 0 |
| md_0_6_A | A | 97 | E | 0.40784 | 1.20824 | TRUE | 0.59979 | 100 | 155 | 0 |
| md_0_6_A | A | 5 | T | 0.40576 | 1.19380 | TRUE | 0.88462 | 100 | 155 | 0 |
| md_0_6_A | A | 42 | T | 0.40092 | 1.16021 | TRUE | 0.44124 | 100 | 155 | 0 |
| md_0_6_A | A | 32 | G | 0.40023 | 1.15543 | TRUE | 0.72303 | 100 | 155 | 0 |
| md_0_6_A | A | 11 | V | 0.39180 | 1.09692 | TRUE | 0.45414 | 100 | 155 | 0 |
| md_0_6_A | A | 154 | S | 0.37677 | 0.99262 | TRUE | 0.85476 | 100 | 155 | 0 |
| md_0_6_A | A | 14 | S | 0.37271 | 0.96444 | TRUE | 0.39739 | 100 | 155 | 0 |
| md_0_6_A | A | 112 | R | 0.37034 | 0.94800 | TRUE | 0.55402 | 100 | 155 | 0 |
| md_0_6_A | A | 73 | N | 0.36948 | 0.94203 | TRUE | 0.15774 | 100 | 155 | 0 |
| md_0_6_A | A | 147 | S | 0.36100 | 0.88318 | FALSE | 0.69378 | 100 | 155 | 0 |
| md_0_6_A | A | 100 | A | 0.35780 | 0.86097 | FALSE | 0.80101 | 100 | 155 | 0 |
| md_0_6_A | A | 38 | Q | 0.35348 | 0.83099 | FALSE | 0.59419 | 100 | 155 | 0 |

Figure 32: Residues with DiscoTope above the threshold of 0.35 — md-0-6-A-discotope3.csv (n = 29)

Residues with DiscoTope ≥ 0.35 — md_0_7_A_discotope3.csv (n = 14)

| PDB | Chain | ResID | AA | DiscoTope | Calibrated | Epitope | RSA | pLDDT | Length | AF2 flag |
|---|---|---|---|---|---|---|---|---|---|---|
| md_0_7_A | A | 46 | R | 0.49347 | 1.94219 | TRUE | 0.60173 | 100 | 155 | 0 |
| md_0_7_A | A | 108 | L | 0.48058 | 1.84580 | TRUE | 0.90332 | 100 | 155 | 0 |
| md_0_7_A | A | 72 | Y | 0.48022 | 1.84311 | TRUE | 0.43029 | 100 | 155 | 0 |
| md_0_7_A | A | 104 | T | 0.42920 | 1.46160 | TRUE | 0.86744 | 100 | 155 | 0 |
| md_0_7_A | A | 93 | I | 0.41880 | 1.38384 | TRUE | 0.77583 | 100 | 155 | 0 |
| md_0_7_A | A | 7 | P | 0.38934 | 1.16354 | TRUE | 0.85638 | 100 | 155 | 0 |
| md_0_7_A | A | 103 | T | 0.38895 | 1.16063 | TRUE | 0.52343 | 100 | 155 | 0 |
| md_0_7_A | A | 85 | G | 0.38559 | 1.13550 | TRUE | 0.35487 | 100 | 155 | 0 |
| md_0_7_A | A | 50 | E | 0.37052 | 1.02282 | TRUE | 0.63089 | 100 | 155 | 0 |
| md_0_7_A | A | 92 | R | 0.36852 | 1.00786 | TRUE | 0.65661 | 100 | 155 | 0 |
| md_0_7_A | A | 90 | R | 0.36137 | 0.95439 | TRUE | 0.33010 | 100 | 155 | 0 |
| md_0_7_A | A | 8 | S | 0.36022 | 0.94580 | TRUE | 0.79437 | 100 | 155 | 0 |
| md_0_7_A | A | 102 | P | 0.35676 | 0.91992 | TRUE | 1.03681 | 100 | 155 | 0 |
| md_0_7_A | A | 116 | D | 0.35065 | 0.87423 | FALSE | 0.61045 | 100 | 155 | 0 |

Figure 33: Residues with DiscoTope above the threshold of 0.35 — md-0-7-A-discotope3.csv (n = 14)

Residues with DiscoTope ≥ 0.35 — md_0_8_A_discotope3.csv (n = 25)

| PDB | Chain | ResID | AA | DiscoTope | Calibrated | Epitope | RSA | pLDDT | Length | AF2 flag |
|---|---|---|---|---|---|---|---|---|---|---|
| md_0_8_A | A | 108 | L | 0.49513 | 1.81617 | TRUE | 0.68210 | 100 | 155 | 0 |
| md_0_8_A | A | 37 | T | 0.48560 | 1.74996 | TRUE | 0.55845 | 100 | 155 | 0 |
| md_0_8_A | A | 46 | R | 0.44960 | 1.49983 | TRUE | 0.53153 | 100 | 155 | 0 |
| md_0_8_A | A | 99 | Q | 0.44040 | 1.43590 | TRUE | 0.69718 | 100 | 155 | 0 |
| md_0_8_A | A | 152 | W | 0.43500 | 1.39838 | TRUE | 0.79094 | 100 | 155 | 0 |
| md_0_8_A | A | 115 | D | 0.42708 | 1.34336 | TRUE | 0.38813 | 100 | 155 | 0 |
| md_0_8_A | A | 103 | T | 0.42191 | 1.30743 | TRUE | 0.49764 | 100 | 155 | 0 |
| md_0_8_A | A | 106 | E | 0.41794 | 1.27985 | TRUE | 0.63167 | 100 | 155 | 0 |
| md_0_8_A | A | 100 | A | 0.41195 | 1.23823 | TRUE | 0.76979 | 100 | 155 | 0 |
| md_0_8_A | A | 112 | R | 0.41105 | 1.23198 | TRUE | 0.54239 | 100 | 155 | 0 |
| md_0_8_A | A | 102 | P | 0.40739 | 1.20655 | TRUE | 1.08217 | 100 | 155 | 0 |
| md_0_8_A | A | 104 | T | 0.40418 | 1.18425 | TRUE | 0.69481 | 100 | 155 | 0 |
| md_0_8_A | A | 72 | Y | 0.40067 | 1.15986 | TRUE | 0.43072 | 100 | 155 | 0 |
| md_0_8_A | A | 88 | D | 0.39402 | 1.11365 | TRUE | 0.70691 | 100 | 155 | 0 |
| md_0_8_A | A | 119 | V | 0.39190 | 1.09892 | TRUE | 0.47984 | 100 | 155 | 0 |
| md_0_8_A | A | 59 | T | 0.39086 | 1.09170 | TRUE | 0.89864 | 100 | 155 | 0 |
| md_0_8_A | A | 61 | R | 0.38948 | 1.08211 | TRUE | 0.35067 | 100 | 155 | 0 |
| md_0_8_A | A | 32 | G | 0.37535 | 0.98393 | TRUE | 0.77973 | 100 | 155 | 0 |
| md_0_8_A | A | 77 | D | 0.37109 | 0.95434 | TRUE | 0.29272 | 100 | 155 | 0 |
| md_0_8_A | A | 34 | Q | 0.36625 | 0.92071 | TRUE | 0.69549 | 100 | 155 | 0 |
| md_0_8_A | A | 122 | R | 0.36140 | 0.88701 | FALSE | 0.41399 | 100 | 155 | 0 |
| md_0_8_A | A | 153 | T | 0.36003 | 0.87749 | FALSE | 0.31026 | 100 | 155 | 0 |
| md_0_8_A | A | 126 | N | 0.35530 | 0.84463 | FALSE | 0.47186 | 100 | 155 | 0 |
| md_0_8_A | A | 42 | T | 0.35393 | 0.83511 | FALSE | 0.46003 | 100 | 155 | 0 |
| md_0_8_A | A | 5 | T | 0.35051 | 0.81134 | FALSE | 0.73788 | 100 | 155 | 0 |

Figure 34: Residues with DiscoTope above the threshold of 0.35 — md-0-8-A-discotope3.csv (n = 25)

Residues with DiscoTope ≥ 0.35 — md_0_9_A_discotope3.csv (n = 26)

| PDB | Chain | ResID | AA | DiscoTope | Calibrated | Epitope | RSA | pLDDT | Length | AF2 flag |
|---|---|---|---|---|---|---|---|---|---|---|
| md_0_9_A | A | 72 | Y | 0.50176 | 1.99711 | TRUE | 0.45068 | 100 | 155 | 0 |
| md_0_9_A | A | 46 | R | 0.48668 | 1.88475 | TRUE | 0.60805 | 100 | 155 | 0 |
| md_0_9_A | A | 115 | D | 0.45526 | 1.65063 | TRUE | 0.29201 | 100 | 155 | 0 |
| md_0_9_A | A | 100 | A | 0.44696 | 1.58878 | TRUE | 0.94406 | 100 | 155 | 0 |
| md_0_9_A | A | 15 | S | 0.43071 | 1.46770 | TRUE | 0.46928 | 100 | 155 | 0 |
| md_0_9_A | A | 112 | R | 0.42746 | 1.44348 | TRUE | 0.64573 | 100 | 155 | 0 |
| md_0_9_A | A | 106 | E | 0.42000 | 1.38790 | TRUE | 0.69877 | 100 | 155 | 0 |
| md_0_9_A | A | 8 | S | 0.41589 | 1.35727 | TRUE | 0.85347 | 100 | 155 | 0 |
| md_0_9_A | A | 101 | N | 0.40409 | 1.26935 | TRUE | 0.29287 | 100 | 155 | 0 |
| md_0_9_A | A | 39 | Q | 0.40170 | 1.25154 | TRUE | 0.74012 | 100 | 155 | 0 |
| md_0_9_A | A | 47 | Q | 0.40035 | 1.24148 | TRUE | 0.44466 | 100 | 155 | 0 |
| md_0_9_A | A | 102 | P | 0.39134 | 1.17434 | TRUE | 0.64177 | 100 | 155 | 0 |
| md_0_9_A | A | 103 | T | 0.38877 | 1.15519 | TRUE | 0.45531 | 100 | 155 | 0 |
| md_0_9_A | A | 34 | Q | 0.38806 | 1.14990 | TRUE | 0.70298 | 100 | 155 | 0 |
| md_0_9_A | A | 105 | A | 0.37538 | 1.05542 | TRUE | 0.97061 | 100 | 155 | 0 |
| md_0_9_A | A | 104 | T | 0.37529 | 1.05475 | TRUE | 0.44735 | 100 | 155 | 0 |
| md_0_9_A | A | 33 | N | 0.37231 | 1.03255 | TRUE | 0.65544 | 100 | 155 | 0 |
| md_0_9_A | A | 9 | Q | 0.37097 | 1.02256 | TRUE | 0.45763 | 100 | 155 | 0 |
| md_0_9_A | A | 88 | D | 0.36949 | 1.01153 | TRUE | 0.53607 | 100 | 155 | 0 |
| md_0_9_A | A | 123 | S | 0.36669 | 0.99067 | TRUE | 0.48912 | 100 | 155 | 0 |
| md_0_9_A | A | 42 | T | 0.36351 | 0.96697 | TRUE | 0.42422 | 100 | 155 | 0 |
| md_0_9_A | A | 74 | A | 0.36049 | 0.94447 | TRUE | 0.82183 | 100 | 155 | 0 |
| md_0_9_A | A | 97 | E | 0.36035 | 0.94343 | TRUE | 0.53501 | 100 | 155 | 0 |
| md_0_9_A | A | 85 | G | 0.35841 | 0.92897 | TRUE | 0.30716 | 100 | 155 | 0 |
| md_0_9_A | A | 32 | G | 0.35475 | 0.90170 | TRUE | 0.72157 | 100 | 155 | 0 |
| md_0_9_A | A | 7 | P | 0.35326 | 0.89060 | FALSE | 0.93472 | 100 | 155 | 0 |

Figure 35: Residues with DiscoTope above the threshold of 0.35 — md-0-9-A-discotope3.csv (n = 26)

Residues with DiscoTope ≥ 0.35 — md_0_10_A_discotope3.csv (n = 18)

| PDB | Chain | ResID | AA | DiscoTope | Calibrated | Epitope | RSA | pLDDT | Length | AF2 flag |
|---|---|---|---|---|---|---|---|---|---|---|
| md_0_10_A | A | 99 | Q | 0.57888 | 2.53298 | TRUE | 0.62786 | 100 | 155 | 0 |
| md_0_10_A | A | 46 | R | 0.52448 | 2.13374 | TRUE | 0.68060 | 100 | 155 | 0 |
| md_0_10_A | A | 102 | P | 0.49740 | 1.93500 | TRUE | 0.93350 | 100 | 155 | 0 |
| md_0_10_A | A | 72 | Y | 0.46418 | 1.69120 | TRUE | 0.46202 | 100 | 155 | 0 |
| md_0_10_A | A | 37 | T | 0.45640 | 1.63411 | TRUE | 0.18986 | 100 | 155 | 0 |
| md_0_10_A | A | 103 | T | 0.44234 | 1.53092 | TRUE | 0.41521 | 100 | 155 | 0 |
| md_0_10_A | A | 101 | N | 0.43048 | 1.44388 | TRUE | 0.31865 | 100 | 155 | 0 |
| md_0_10_A | A | 34 | Q | 0.43029 | 1.44249 | TRUE | 0.67257 | 100 | 155 | 0 |
| md_0_10_A | A | 39 | Q | 0.41636 | 1.34026 | TRUE | 0.70015 | 100 | 155 | 0 |
| md_0_10_A | A | 32 | G | 0.41389 | 1.32213 | TRUE | 0.78900 | 100 | 155 | 0 |
| md_0_10_A | A | 105 | A | 0.40320 | 1.24368 | TRUE | 0.73821 | 100 | 155 | 0 |
| md_0_10_A | A | 42 | T | 0.39226 | 1.16339 | TRUE | 0.42613 | 100 | 155 | 0 |
| md_0_10_A | A | 81 | T | 0.38609 | 1.11811 | TRUE | 0.54759 | 100 | 155 | 0 |
| md_0_10_A | A | 152 | W | 0.37885 | 1.06497 | TRUE | 0.74493 | 100 | 155 | 0 |
| md_0_10_A | A | 147 | S | 0.37811 | 1.05954 | TRUE | 0.48421 | 100 | 155 | 0 |
| md_0_10_A | A | 106 | E | 0.37737 | 1.05411 | TRUE | 0.70597 | 100 | 155 | 0 |
| md_0_10_A | A | 104 | T | 0.37495 | 1.03635 | TRUE | 0.50421 | 100 | 155 | 0 |
| md_0_10_A | A | 54 | P | 0.35541 | 0.89295 | FALSE | 0.76402 | 100 | 155 | 0 |

Figure 36: Residues with DiscoTope above the threshold of 0.35 — md-0-10-A-discotope3.csv (n = 18)

Residues with DiscoTope ≥ 0.35 — md_0_11_A_discotope3.csv (n = 21)

| PDB | Chain | ResID | AA | DiscoTope | Calibrated | Epitope | RSA | pLDDT | Length | AF2 flag |
|---|---|---|---|---|---|---|---|---|---|---|
| md_0_11_A | A | 72 | Y | 0.48959 | 1.90884 | TRUE | 0.44136 | 100 | 155 | 0 |
| md_0_11_A | A | 7 | P | 0.47847 | 1.82587 | TRUE | 0.60836 | 100 | 155 | 0 |
| md_0_11_A | A | 46 | R | 0.46495 | 1.72501 | TRUE | 0.59776 | 100 | 155 | 0 |
| md_0_11_A | A | 85 | G | 0.44066 | 1.54379 | TRUE | 0.49718 | 100 | 155 | 0 |
| md_0_11_A | A | 92 | R | 0.42637 | 1.43717 | TRUE | 0.66902 | 100 | 155 | 0 |
| md_0_11_A | A | 115 | D | 0.42595 | 1.43404 | TRUE | 0.30772 | 100 | 155 | 0 |
| md_0_11_A | A | 104 | T | 0.41304 | 1.33772 | TRUE | 0.56673 | 100 | 155 | 0 |
| md_0_11_A | A | 108 | L | 0.41129 | 1.32467 | TRUE | 0.78001 | 100 | 155 | 0 |
| md_0_11_A | A | 112 | R | 0.40998 | 1.31489 | TRUE | 0.64451 | 100 | 155 | 0 |
| md_0_11_A | A | 33 | N | 0.40709 | 1.29333 | TRUE | 0.67487 | 100 | 155 | 0 |
| md_0_11_A | A | 106 | E | 0.38987 | 1.16486 | TRUE | 0.75449 | 100 | 155 | 0 |
| md_0_11_A | A | 102 | P | 0.38876 | 1.15658 | TRUE | 0.95539 | 100 | 155 | 0 |
| md_0_11_A | A | 119 | V | 0.37599 | 1.06130 | TRUE | 0.53316 | 100 | 155 | 0 |
| md_0_11_A | A | 50 | E | 0.37269 | 1.03668 | TRUE | 0.56288 | 100 | 155 | 0 |
| md_0_11_A | A | 105 | A | 0.37204 | 1.03183 | TRUE | 0.69187 | 100 | 155 | 0 |
| md_0_11_A | A | 99 | Q | 0.37080 | 1.02258 | TRUE | 0.48551 | 100 | 155 | 0 |
| md_0_11_A | A | 81 | T | 0.36814 | 1.00274 | TRUE | 0.53147 | 100 | 155 | 0 |
| md_0_11_A | A | 77 | D | 0.36776 | 0.99990 | TRUE | 0.32878 | 100 | 155 | 0 |
| md_0_11_A | A | 54 | P | 0.36550 | 0.98304 | TRUE | 0.49838 | 100 | 155 | 0 |
| md_0_11_A | A | 39 | Q | 0.36538 | 0.98215 | TRUE | 0.48604 | 100 | 155 | 0 |
| md_0_11_A | A | 32 | G | 0.36124 | 0.95126 | TRUE | 0.74303 | 100 | 155 | 0 |

Figure 37: Residues with DiscoTope above the threshold of 0.35 — md-0-11-A-discotope3.csv (n = 21)

Residues with DiscoTope ≥ 0.35 — md_0_12_A_discotope3.csv (n = 22)

| PDB | Chain | ResID | AA | DiscoTope | Calibrated | Epitope | RSA | pLDDT | Length | AF2 flag |
|---|---|---|---|---|---|---|---|---|---|---|
| md_0_12_A | A | 72 | Y | 0.50460 | 1.96551 | TRUE | 0.41458 | 100 | 155 | 0 |
| md_0_12_A | A | 108 | L | 0.49198 | 1.87394 | TRUE | 0.93495 | 100 | 155 | 0 |
| md_0_12_A | A | 46 | R | 0.47366 | 1.74100 | TRUE | 0.59982 | 100 | 155 | 0 |
| md_0_12_A | A | 99 | Q | 0.45789 | 1.62656 | TRUE | 0.58769 | 100 | 155 | 0 |
| md_0_12_A | A | 85 | G | 0.45206 | 1.58426 | TRUE | 0.48139 | 100 | 155 | 0 |
| md_0_12_A | A | 106 | E | 0.44412 | 1.52664 | TRUE | 0.73658 | 100 | 155 | 0 |
| md_0_12_A | A | 93 | I | 0.43197 | 1.43848 | TRUE | 0.72536 | 100 | 155 | 0 |
| md_0_12_A | A | 112 | R | 0.42779 | 1.40814 | TRUE | 0.65439 | 100 | 155 | 0 |
| md_0_12_A | A | 90 | R | 0.42751 | 1.40611 | TRUE | 0.64557 | 100 | 155 | 0 |
| md_0_12_A | A | 37 | T | 0.42184 | 1.36497 | TRUE | 0.15868 | 100 | 155 | 0 |
| md_0_12_A | A | 92 | R | 0.42094 | 1.35844 | TRUE | 0.39213 | 100 | 155 | 0 |
| md_0_12_A | A | 42 | T | 0.41451 | 1.31178 | TRUE | 0.58256 | 100 | 155 | 0 |
| md_0_12_A | A | 116 | D | 0.40725 | 1.25910 | TRUE | 0.62777 | 100 | 155 | 0 |
| md_0_12_A | A | 39 | Q | 0.40219 | 1.22238 | TRUE | 0.64046 | 100 | 155 | 0 |
| md_0_12_A | A | 59 | T | 0.39193 | 1.14793 | TRUE | 0.58207 | 100 | 155 | 0 |
| md_0_12_A | A | 152 | W | 0.38992 | 1.13334 | TRUE | 0.78244 | 100 | 155 | 0 |
| md_0_12_A | A | 103 | T | 0.38863 | 1.12398 | TRUE | 0.42472 | 100 | 155 | 0 |
| md_0_12_A | A | 96 | V | 0.38595 | 1.10453 | TRUE | 0.58992 | 100 | 155 | 0 |
| md_0_12_A | A | 107 | T | 0.38506 | 1.09807 | TRUE | 0.36103 | 100 | 155 | 0 |
| md_0_12_A | A | 8 | S | 0.38233 | 1.07826 | TRUE | 0.73085 | 100 | 155 | 0 |
| md_0_12_A | A | 77 | D | 0.37423 | 1.01949 | TRUE | 0.27241 | 100 | 155 | 0 |
| md_0_12_A | A | 105 | A | 0.36041 | 0.91920 | TRUE | 0.58861 | 100 | 155 | 0 |

Figure 38: Residues with DiscoTope above the threshold of 0.35 — md-0-12-A-discotope3.csv (n = 22)

Residues with DiscoTope ≥ 0.35 — md_0_13_A_discotope3.csv (n = 24)

| PDB | Chain | ResID | AA | DiscoTope | Calibrated | Epitope | RSA | pLDDT | Length | AF2 flag |
|---|---|---|---|---|---|---|---|---|---|---|
| md_0_13_A | A | 108 | L | 0.51922 | 1.98066 | TRUE | 0.48027 | 100 | 155 | 0 |
| md_0_13_A | A | 46 | R | 0.51656 | 1.96220 | TRUE | 0.63070 | 100 | 155 | 0 |
| md_0_13_A | A | 109 | D | 0.48906 | 1.77141 | TRUE | 0.63210 | 100 | 155 | 0 |
| md_0_13_A | A | 102 | P | 0.48171 | 1.72042 | TRUE | 0.94261 | 100 | 155 | 0 |
| md_0_13_A | A | 101 | N | 0.47845 | 1.69780 | TRUE | 0.38150 | 100 | 155 | 0 |
| md_0_13_A | A | 103 | T | 0.45420 | 1.52955 | TRUE | 0.35940 | 100 | 155 | 0 |
| md_0_13_A | A | 100 | A | 0.44370 | 1.45671 | TRUE | 0.96003 | 100 | 155 | 0 |
| md_0_13_A | A | 106 | E | 0.44272 | 1.44991 | TRUE | 0.66730 | 100 | 155 | 0 |
| md_0_13_A | A | 43 | V | 0.43469 | 1.39420 | TRUE | 0.57149 | 100 | 155 | 0 |
| md_0_13_A | A | 54 | P | 0.43202 | 1.37567 | TRUE | 0.63063 | 100 | 155 | 0 |
| md_0_13_A | A | 98 | N | 0.41869 | 1.28319 | TRUE | 0.35494 | 100 | 155 | 0 |
| md_0_13_A | A | 123 | S | 0.41853 | 1.28208 | TRUE | 0.43790 | 100 | 155 | 0 |
| md_0_13_A | A | 96 | V | 0.41337 | 1.24628 | TRUE | 0.46625 | 100 | 155 | 0 |
| md_0_13_A | A | 92 | R | 0.40860 | 1.21319 | TRUE | 0.35258 | 100 | 155 | 0 |
| md_0_13_A | A | 112 | R | 0.40207 | 1.16788 | TRUE | 0.57715 | 100 | 155 | 0 |
| md_0_13_A | A | 152 | W | 0.39454 | 1.11564 | TRUE | 0.80740 | 100 | 155 | 0 |
| md_0_13_A | A | 97 | E | 0.37872 | 1.00588 | TRUE | 0.72887 | 100 | 155 | 0 |
| md_0_13_A | A | 99 | Q | 0.37223 | 0.96085 | TRUE | 0.51660 | 100 | 155 | 0 |
| md_0_13_A | A | 72 | Y | 0.37014 | 0.94635 | TRUE | 0.45012 | 100 | 155 | 0 |
| md_0_13_A | A | 119 | V | 0.36817 | 0.93269 | TRUE | 0.55806 | 100 | 155 | 0 |
| md_0_13_A | A | 5 | T | 0.36087 | 0.88204 | FALSE | 0.82467 | 100 | 155 | 0 |
| md_0_13_A | A | 8 | S | 0.35606 | 0.84867 | FALSE | 0.65636 | 100 | 155 | 0 |
| md_0_13_A | A | 85 | G | 0.35274 | 0.82563 | FALSE | 0.55755 | 100 | 155 | 0 |
| md_0_13_A | A | 126 | N | 0.35004 | 0.80690 | FALSE | 0.54976 | 100 | 155 | 0 |

Figure 39: Residues with DiscoTope above the threshold of 0.35 — md-0-13-A-discotope3.csv (n = 24)

Residues with DiscoTope ≥ 0.35 — md_0_14_A_discotope3.csv (n = 29)

| PDB | Chain | ResID | AA | DiscoTope | Calibrated | Epitope | RSA | pLDDT | Length | AF2 flag |
|---|---|---|---|---|---|---|---|---|---|---|
| md_0_14_A | A | 99 | Q | 0.54888 | 2.15519 | TRUE | 0.69417 | 100 | 155 | 0 |
| md_0_14_A | A | 106 | E | 0.52800 | 2.01240 | TRUE | 0.60579 | 100 | 155 | 0 |
| md_0_14_A | A | 107 | T | 0.51204 | 1.90325 | TRUE | 0.36621 | 100 | 155 | 0 |
| md_0_14_A | A | 112 | R | 0.50750 | 1.87221 | TRUE | 0.53505 | 100 | 155 | 0 |
| md_0_14_A | A | 109 | D | 0.46489 | 1.58081 | TRUE | 0.60716 | 100 | 155 | 0 |
| md_0_14_A | A | 39 | Q | 0.45743 | 1.52979 | TRUE | 0.68431 | 100 | 155 | 0 |
| md_0_14_A | A | 103 | T | 0.45685 | 1.52582 | TRUE | 0.32263 | 100 | 155 | 0 |
| md_0_14_A | A | 108 | L | 0.45149 | 1.48917 | TRUE | 0.39890 | 100 | 155 | 0 |
| md_0_14_A | A | 102 | P | 0.45114 | 1.48677 | TRUE | 0.90699 | 100 | 155 | 0 |
| md_0_14_A | A | 119 | V | 0.44188 | 1.42345 | TRUE | 0.53588 | 100 | 155 | 0 |
| md_0_14_A | A | 72 | Y | 0.42960 | 1.33947 | TRUE | 0.49797 | 100 | 155 | 0 |
| md_0_14_A | A | 152 | W | 0.42794 | 1.32811 | TRUE | 0.75698 | 100 | 155 | 0 |
| md_0_14_A | A | 100 | A | 0.42312 | 1.29515 | TRUE | 0.95136 | 100 | 155 | 0 |
| md_0_14_A | A | 29 | N | 0.41293 | 1.22546 | TRUE | 0.38180 | 100 | 155 | 0 |
| md_0_14_A | A | 98 | N | 0.40804 | 1.19202 | TRUE | 0.27278 | 100 | 155 | 0 |
| md_0_14_A | A | 32 | G | 0.40488 | 1.17041 | TRUE | 0.74080 | 100 | 155 | 0 |
| md_0_14_A | A | 101 | N | 0.40203 | 1.15092 | TRUE | 0.37067 | 100 | 155 | 0 |
| md_0_14_A | A | 43 | V | 0.39547 | 1.10606 | TRUE | 0.37748 | 100 | 155 | 0 |
| md_0_14_A | A | 97 | E | 0.38985 | 1.06762 | TRUE | 0.61707 | 100 | 155 | 0 |
| md_0_14_A | A | 115 | D | 0.38925 | 1.06352 | TRUE | 0.19195 | 100 | 155 | 0 |
| md_0_14_A | A | 34 | Q | 0.38518 | 1.03569 | TRUE | 0.31886 | 100 | 155 | 0 |
| md_0_14_A | A | 33 | N | 0.37744 | 0.98275 | TRUE | 0.53687 | 100 | 155 | 0 |
| md_0_14_A | A | 105 | A | 0.37292 | 0.95184 | TRUE | 0.87185 | 100 | 155 | 0 |
| md_0_14_A | A | 81 | T | 0.36505 | 0.89802 | FALSE | 0.51947 | 100 | 155 | 0 |
| md_0_14_A | A | 74 | A | 0.36466 | 0.89536 | FALSE | 0.80788 | 100 | 155 | 0 |
| md_0_14_A | A | 47 | Q | 0.36013 | 0.86438 | FALSE | 0.37792 | 100 | 155 | 0 |
| md_0_14_A | A | 46 | R | 0.35668 | 0.84078 | FALSE | 0.53624 | 100 | 155 | 0 |
| md_0_14_A | A | 73 | N | 0.35107 | 0.80242 | FALSE | 0.11299 | 100 | 155 | 0 |
| md_0_14_A | A | 58 | V | 0.35094 | 0.80153 | FALSE | 0.42653 | 100 | 155 | 0 |

Figure 40: Residues with DiscoTope above the threshold of 0.35 — md-0-14-A-discotope3.csv (n = 29)

Residues with DiscoTope ≥ 0.35 — md_0_15_A_discotope3.csv (n = 34)

| PDB | Chain | ResID | AA | DiscoTope | Calibrated | Epitope | RSA | pLDDT | Length | AF2 flag |
|---|---|---|---|---|---|---|---|---|---|---|
| md_0_15_A | A | 92 | R | 0.52334 | 1.95572 | TRUE | 0.43520 | 100 | 155 | 0 |
| md_0_15_A | A | 102 | P | 0.51824 | 1.92128 | TRUE | 0.97367 | 100 | 155 | 0 |
| md_0_15_A | A | 46 | R | 0.50707 | 1.84585 | TRUE | 0.60616 | 100 | 155 | 0 |
| md_0_15_A | A | 99 | Q | 0.48915 | 1.72483 | TRUE | 0.73087 | 100 | 155 | 0 |
| md_0_15_A | A | 98 | N | 0.47925 | 1.65798 | TRUE | 0.30342 | 100 | 155 | 0 |
| md_0_15_A | A | 72 | Y | 0.47043 | 1.59841 | TRUE | 0.50650 | 100 | 155 | 0 |
| md_0_15_A | A | 107 | T | 0.46883 | 1.58761 | TRUE | 0.30379 | 100 | 155 | 0 |
| md_0_15_A | A | 109 | D | 0.46364 | 1.55256 | TRUE | 0.66586 | 100 | 155 | 0 |
| md_0_15_A | A | 49 | S | 0.45910 | 1.52190 | TRUE | 0.32088 | 100 | 155 | 0 |
| md_0_15_A | A | 100 | A | 0.45099 | 1.46713 | TRUE | 0.89679 | 100 | 155 | 0 |
| md_0_15_A | A | 96 | V | 0.44640 | 1.43614 | TRUE | 0.48196 | 100 | 155 | 0 |
| md_0_15_A | A | 90 | R | 0.44465 | 1.42432 | TRUE | 0.88336 | 100 | 155 | 0 |
| md_0_15_A | A | 97 | E | 0.43837 | 1.38191 | TRUE | 0.63009 | 100 | 155 | 0 |
| md_0_15_A | A | 106 | E | 0.43489 | 1.35841 | TRUE | 0.63092 | 100 | 155 | 0 |
| md_0_15_A | A | 93 | I | 0.42716 | 1.30621 | TRUE | 0.63917 | 100 | 155 | 0 |
| md_0_15_A | A | 39 | Q | 0.42154 | 1.26825 | TRUE | 0.60239 | 100 | 155 | 0 |
| md_0_15_A | A | 101 | N | 0.41965 | 1.25549 | TRUE | 0.37146 | 100 | 155 | 0 |
| md_0_15_A | A | 108 | L | 0.41811 | 1.24509 | TRUE | 0.53823 | 100 | 155 | 0 |
| md_0_15_A | A | 103 | T | 0.41683 | 1.23645 | TRUE | 0.41058 | 100 | 155 | 0 |
| md_0_15_A | A | 32 | G | 0.41173 | 1.20201 | TRUE | 0.85451 | 100 | 155 | 0 |
| md_0_15_A | A | 50 | E | 0.39846 | 1.11239 | TRUE | 0.50997 | 100 | 155 | 0 |
| md_0_15_A | A | 113 | R | 0.39397 | 1.08207 | TRUE | 0.46700 | 100 | 155 | 0 |
| md_0_15_A | A | 91 | N | 0.38742 | 1.03784 | TRUE | 0.48092 | 100 | 155 | 0 |
| md_0_15_A | A | 152 | W | 0.37840 | 0.97693 | TRUE | 0.69775 | 100 | 155 | 0 |
| md_0_15_A | A | 29 | N | 0.37424 | 0.94883 | TRUE | 0.57609 | 100 | 155 | 0 |
| md_0_15_A | A | 38 | Q | 0.36889 | 0.91270 | TRUE | 0.36737 | 100 | 155 | 0 |
| md_0_15_A | A | 85 | G | 0.36875 | 0.91176 | TRUE | 0.66018 | 100 | 155 | 0 |
| md_0_15_A | A | 123 | S | 0.36631 | 0.89528 | FALSE | 0.39675 | 100 | 155 | 0 |
| md_0_15_A | A | 116 | D | 0.35975 | 0.85098 | FALSE | 0.64473 | 100 | 155 | 0 |
| md_0_15_A | A | 73 | N | 0.35774 | 0.83741 | FALSE | 0.15295 | 100 | 155 | 0 |
| md_0_15_A | A | 43 | V | 0.35420 | 0.81350 | FALSE | 0.58076 | 100 | 155 | 0 |
| md_0_15_A | A | 105 | A | 0.35078 | 0.79040 | FALSE | 0.52897 | 100 | 155 | 0 |
| md_0_15_A | A | 33 | N | 0.35064 | 0.78946 | FALSE | 0.43323 | 100 | 155 | 0 |
| md_0_15_A | A | 42 | T | 0.35056 | 0.78892 | FALSE | 0.44201 | 100 | 155 | 0 |

Figure 41: Residues with DiscoTope above the threshold of 0.35 — md-0-15-A-discotope3.csv (n = 34)

Residues with DiscoTope ≥ 0.35 — md_0_16_A_discotope3.csv (n = 29)

| PDB | Chain | ResID | AA | DiscoTope | Calibrated | Epitope | RSA | pLDDT | Length | AF2 flag |
|---|---|---|---|---|---|---|---|---|---|---|
| md_0_16_A | A | 107 | T | 0.54202 | 2.14708 | TRUE | 0.30152 | 100 | 155 | 0 |
| md_0_16_A | A | 39 | Q | 0.51066 | 1.92867 | TRUE | 0.45822 | 100 | 155 | 0 |
| md_0_16_A | A | 93 | I | 0.49231 | 1.80087 | TRUE | 0.66438 | 100 | 155 | 0 |
| md_0_16_A | A | 109 | D | 0.47692 | 1.69368 | TRUE | 0.68425 | 100 | 155 | 0 |
| md_0_16_A | A | 108 | L | 0.47642 | 1.69020 | TRUE | 0.38915 | 100 | 155 | 0 |
| md_0_16_A | A | 72 | Y | 0.47454 | 1.67711 | TRUE | 0.45599 | 100 | 155 | 0 |
| md_0_16_A | A | 101 | N | 0.46191 | 1.58915 | TRUE | 0.40677 | 100 | 155 | 0 |
| md_0_16_A | A | 46 | R | 0.45995 | 1.57549 | TRUE | 0.61299 | 100 | 155 | 0 |
| md_0_16_A | A | 98 | N | 0.45353 | 1.53078 | TRUE | 0.36326 | 100 | 155 | 0 |
| md_0_16_A | A | 102 | P | 0.45332 | 1.52932 | TRUE | 0.73315 | 100 | 155 | 0 |
| md_0_16_A | A | 85 | G | 0.42460 | 1.32929 | TRUE | 0.52019 | 100 | 155 | 0 |
| md_0_16_A | A | 100 | A | 0.42290 | 1.31745 | TRUE | 1.01795 | 100 | 155 | 0 |
| md_0_16_A | A | 106 | E | 0.41578 | 1.26787 | TRUE | 0.67826 | 100 | 155 | 0 |
| md_0_16_A | A | 92 | R | 0.40496 | 1.19251 | TRUE | 0.37397 | 100 | 155 | 0 |
| md_0_16_A | A | 96 | V | 0.39778 | 1.14250 | TRUE | 0.57131 | 100 | 155 | 0 |
| md_0_16_A | A | 90 | R | 0.39624 | 1.13178 | TRUE | 0.94817 | 100 | 155 | 0 |
| md_0_16_A | A | 38 | Q | 0.39557 | 1.12711 | TRUE | 0.45236 | 100 | 155 | 0 |
| md_0_16_A | A | 103 | T | 0.39495 | 1.12279 | TRUE | 0.48553 | 100 | 155 | 0 |
| md_0_16_A | A | 91 | N | 0.39430 | 1.11827 | TRUE | 0.43032 | 100 | 155 | 0 |
| md_0_16_A | A | 6 | T | 0.39196 | 1.10197 | TRUE | 0.48543 | 100 | 155 | 0 |
| md_0_16_A | A | 97 | E | 0.38745 | 1.07056 | TRUE | 0.73848 | 100 | 155 | 0 |
| md_0_16_A | A | 99 | Q | 0.37900 | 1.01171 | TRUE | 0.50416 | 100 | 155 | 0 |
| md_0_16_A | A | 105 | A | 0.37798 | 1.00460 | TRUE | 0.36455 | 100 | 155 | 0 |
| md_0_16_A | A | 32 | G | 0.37397 | 0.97667 | TRUE | 0.76935 | 100 | 155 | 0 |
| md_0_16_A | A | 81 | T | 0.37169 | 0.96080 | TRUE | 0.50441 | 100 | 155 | 0 |
| md_0_16_A | A | 42 | T | 0.37129 | 0.95801 | TRUE | 0.50067 | 100 | 155 | 0 |
| md_0_16_A | A | 5 | T | 0.36616 | 0.92228 | TRUE | 0.86982 | 100 | 155 | 0 |
| md_0_16_A | A | 113 | R | 0.36264 | 0.89776 | FALSE | 0.52963 | 100 | 155 | 0 |
| md_0_16_A | A | 115 | D | 0.35673 | 0.85660 | FALSE | 0.16669 | 100 | 155 | 0 |

Figure 42: Residues with DiscoTope above the threshold of 0.35 — md-0-16-A-discotope3.csv (n = 29)

Residues with DiscoTope ≥ 0.35 — md_0_17_A_discotope3.csv (n = 28)

| PDB | Chain | ResID | AA | DiscoTope | Calibrated | Epitope | RSA | pLDDT | Length | AF2 flag |
|---|---|---|---|---|---|---|---|---|---|---|
| md_0_17_A | A | 72 | Y | 0.52861 | 2.07112 | TRUE | 0.43410 | 100 | 155 | 0 |
| md_0_17_A | A | 112 | R | 0.51382 | 1.96724 | TRUE | 0.57870 | 100 | 155 | 0 |
| md_0_17_A | A | 46 | R | 0.50072 | 1.87523 | TRUE | 0.58528 | 100 | 155 | 0 |
| md_0_17_A | A | 92 | R | 0.49598 | 1.84194 | TRUE | 0.39533 | 100 | 155 | 0 |
| md_0_17_A | A | 147 | S | 0.47622 | 1.70315 | TRUE | 0.37514 | 100 | 155 | 0 |
| md_0_17_A | A | 73 | N | 0.44076 | 1.45409 | TRUE | 0.28150 | 100 | 155 | 0 |
| md_0_17_A | A | 152 | W | 0.44043 | 1.45177 | TRUE | 0.73749 | 100 | 155 | 0 |
| md_0_17_A | A | 99 | Q | 0.43279 | 1.39811 | TRUE | 0.50761 | 100 | 155 | 0 |
| md_0_17_A | A | 85 | G | 0.42980 | 1.37711 | TRUE | 0.73842 | 100 | 155 | 0 |
| md_0_17_A | A | 90 | R | 0.42509 | 1.34402 | TRUE | 0.78013 | 100 | 155 | 0 |
| md_0_17_A | A | 113 | R | 0.42030 | 1.31038 | TRUE | 0.48061 | 100 | 155 | 0 |
| md_0_17_A | A | 49 | S | 0.41663 | 1.28460 | TRUE | 0.35056 | 100 | 155 | 0 |
| md_0_17_A | A | 39 | Q | 0.41008 | 1.23860 | TRUE | 0.66831 | 100 | 155 | 0 |
| md_0_17_A | A | 91 | N | 0.40831 | 1.22616 | TRUE | 0.51141 | 100 | 155 | 0 |
| md_0_17_A | A | 81 | T | 0.40793 | 1.22349 | TRUE | 0.50745 | 100 | 155 | 0 |
| md_0_17_A | A | 101 | N | 0.40605 | 1.21029 | TRUE | 0.29331 | 100 | 155 | 0 |
| md_0_17_A | A | 32 | G | 0.40593 | 1.20945 | TRUE | 0.73092 | 100 | 155 | 0 |
| md_0_17_A | A | 97 | E | 0.40236 | 1.18437 | TRUE | 0.66159 | 100 | 155 | 0 |
| md_0_17_A | A | 109 | D | 0.40063 | 1.17222 | TRUE | 0.77406 | 100 | 155 | 0 |
| md_0_17_A | A | 108 | L | 0.38655 | 1.07333 | TRUE | 0.36301 | 100 | 155 | 0 |
| md_0_17_A | A | 100 | A | 0.38475 | 1.06068 | TRUE | 0.99902 | 100 | 155 | 0 |
| md_0_17_A | A | 102 | P | 0.37415 | 0.98623 | TRUE | 0.97038 | 100 | 155 | 0 |
| md_0_17_A | A | 98 | N | 0.36926 | 0.95189 | TRUE | 0.37123 | 100 | 155 | 0 |
| md_0_17_A | A | 103 | T | 0.36359 | 0.91206 | TRUE | 0.39713 | 100 | 155 | 0 |
| md_0_17_A | A | 116 | D | 0.35828 | 0.87476 | FALSE | 0.65137 | 100 | 155 | 0 |
| md_0_17_A | A | 93 | I | 0.35504 | 0.85201 | FALSE | 0.62776 | 100 | 155 | 0 |
| md_0_17_A | A | 42 | T | 0.35469 | 0.84955 | FALSE | 0.40389 | 100 | 155 | 0 |
| md_0_17_A | A | 126 | N | 0.35173 | 0.82876 | FALSE | 0.54947 | 100 | 155 | 0 |

Figure 43: Residues with DiscoTope above the threshold of 0.35 — md-0-17-A-discotope3.csv (n = 28)

Residues with DiscoTope ≥ 0.35 — md_0_18_A_discotope3.csv (n = 28)

| PDB | Chain | ResID | AA | DiscoTope | Calibrated | Epitope | RSA | pLDDT | Length | AF2 flag |
|---|---|---|---|---|---|---|---|---|---|---|
| md_0_18_A | A | 123 | S | 0.53197 | 1.98516 | TRUE | 0.46449 | 100 | 155 | 0 |
| md_0_18_A | A | 99 | Q | 0.52575 | 1.94375 | TRUE | 0.41606 | 100 | 155 | 0 |
| md_0_18_A | A | 72 | Y | 0.52180 | 1.91746 | TRUE | 0.41053 | 100 | 155 | 0 |
| md_0_18_A | A | 98 | N | 0.48458 | 1.66971 | TRUE | 0.41686 | 100 | 155 | 0 |
| md_0_18_A | A | 101 | N | 0.47471 | 1.60401 | TRUE | 0.40062 | 100 | 155 | 0 |
| md_0_18_A | A | 100 | A | 0.47280 | 1.59130 | TRUE | 1.01161 | 100 | 155 | 0 |
| md_0_18_A | A | 103 | T | 0.46910 | 1.56667 | TRUE | 0.41340 | 100 | 155 | 0 |
| md_0_18_A | A | 109 | D | 0.46202 | 1.51954 | TRUE | 0.58869 | 100 | 155 | 0 |
| md_0_18_A | A | 96 | V | 0.45158 | 1.45005 | TRUE | 0.65197 | 100 | 155 | 0 |
| md_0_18_A | A | 102 | P | 0.45152 | 1.44965 | TRUE | 0.90625 | 100 | 155 | 0 |
| md_0_18_A | A | 46 | R | 0.42121 | 1.24790 | TRUE | 0.56293 | 100 | 155 | 0 |
| md_0_18_A | A | 143 | S | 0.41606 | 1.21361 | TRUE | 0.39115 | 100 | 155 | 0 |
| md_0_18_A | A | 113 | R | 0.40096 | 1.11310 | TRUE | 0.49473 | 100 | 155 | 0 |
| md_0_18_A | A | 106 | E | 0.39337 | 1.06258 | TRUE | 0.67414 | 100 | 155 | 0 |
| md_0_18_A | A | 119 | V | 0.39106 | 1.04721 | TRUE | 0.55402 | 100 | 155 | 0 |
| md_0_18_A | A | 107 | T | 0.38741 | 1.02291 | TRUE | 0.32095 | 100 | 155 | 0 |
| md_0_18_A | A | 152 | W | 0.38654 | 1.01712 | TRUE | 0.71380 | 100 | 155 | 0 |
| md_0_18_A | A | 50 | E | 0.38422 | 1.00168 | TRUE | 0.57075 | 100 | 155 | 0 |
| md_0_18_A | A | 54 | P | 0.38318 | 0.99475 | TRUE | 0.62407 | 100 | 155 | 0 |
| md_0_18_A | A | 91 | N | 0.37945 | 0.96992 | TRUE | 0.46602 | 100 | 155 | 0 |
| md_0_18_A | A | 90 | R | 0.37576 | 0.94536 | TRUE | 0.73206 | 100 | 155 | 0 |
| md_0_18_A | A | 47 | Q | 0.36941 | 0.90309 | TRUE | 0.48091 | 100 | 155 | 0 |
| md_0_18_A | A | 29 | N | 0.36477 | 0.87221 | FALSE | 0.54323 | 100 | 155 | 0 |
| md_0_18_A | A | 105 | A | 0.35667 | 0.81829 | FALSE | 0.47087 | 100 | 155 | 0 |
| md_0_18_A | A | 92 | R | 0.35559 | 0.81110 | FALSE | 0.37161 | 100 | 155 | 0 |
| md_0_18_A | A | 39 | Q | 0.35558 | 0.81104 | FALSE | 0.52740 | 100 | 155 | 0 |
| md_0_18_A | A | 93 | I | 0.35168 | 0.78508 | FALSE | 0.66389 | 100 | 155 | 0 |
| md_0_18_A | A | 97 | E | 0.35046 | 0.77696 | FALSE | 0.63083 | 100 | 155 | 0 |

Figure 44: Residues with DiscoTope above the threshold of 0.35 — md-0-18-A-discotope3.csv (n = 28)

Residues with DiscoTope ≥ 0.35 — md_0_19_A_discotope3.csv (n = 23)

| PDB | Chain | ResID | AA | DiscoTope | Calibrated | Epitope | RSA | pLDDT | Length | AF2 flag |
|---|---|---|---|---|---|---|---|---|---|---|
| md_0_19_A | A | 72 | Y | 0.49238 | 1.91558 | TRUE | 0.44788 | 100 | 155 | 0 |
| md_0_19_A | A | 106 | E | 0.46074 | 1.68124 | TRUE | 0.66735 | 100 | 155 | 0 |
| md_0_19_A | A | 100 | A | 0.45621 | 1.64769 | TRUE | 1.06917 | 100 | 155 | 0 |
| md_0_19_A | A | 101 | N | 0.44595 | 1.57170 | TRUE | 0.45503 | 100 | 155 | 0 |
| md_0_19_A | A | 107 | T | 0.44352 | 1.55371 | TRUE | 0.30134 | 100 | 155 | 0 |
| md_0_19_A | A | 46 | R | 0.43252 | 1.47224 | TRUE | 0.56871 | 100 | 155 | 0 |
| md_0_19_A | A | 73 | N | 0.41742 | 1.36040 | TRUE | 0.15379 | 100 | 155 | 0 |
| md_0_19_A | A | 98 | N | 0.41209 | 1.32093 | TRUE | 0.31501 | 100 | 155 | 0 |
| md_0_19_A | A | 96 | V | 0.41147 | 1.31634 | TRUE | 0.57559 | 100 | 155 | 0 |
| md_0_19_A | A | 99 | Q | 0.39614 | 1.20280 | TRUE | 0.39373 | 100 | 155 | 0 |
| md_0_19_A | A | 81 | T | 0.39250 | 1.17584 | TRUE | 0.48697 | 100 | 155 | 0 |
| md_0_19_A | A | 103 | T | 0.38939 | 1.15281 | TRUE | 0.44044 | 100 | 155 | 0 |
| md_0_19_A | A | 43 | V | 0.38507 | 1.12081 | TRUE | 0.57402 | 100 | 155 | 0 |
| md_0_19_A | A | 49 | S | 0.38460 | 1.11733 | TRUE | 0.30125 | 100 | 155 | 0 |
| md_0_19_A | A | 113 | R | 0.38289 | 1.10467 | TRUE | 0.54951 | 100 | 155 | 0 |
| md_0_19_A | A | 109 | D | 0.37464 | 1.04356 | TRUE | 0.63947 | 100 | 155 | 0 |
| md_0_19_A | A | 85 | G | 0.36763 | 0.99165 | TRUE | 0.62298 | 100 | 155 | 0 |
| md_0_19_A | A | 102 | P | 0.36634 | 0.98209 | TRUE | 0.93547 | 100 | 155 | 0 |
| md_0_19_A | A | 112 | R | 0.35986 | 0.93410 | TRUE | 0.55682 | 100 | 155 | 0 |
| md_0_19_A | A | 119 | V | 0.35888 | 0.92684 | TRUE | 0.54678 | 100 | 155 | 0 |
| md_0_19_A | A | 32 | G | 0.35676 | 0.91114 | TRUE | 0.85049 | 100 | 155 | 0 |
| md_0_19_A | A | 42 | T | 0.35607 | 0.90603 | TRUE | 0.51818 | 100 | 155 | 0 |
| md_0_19_A | A | 78 | P | 0.35526 | 0.90003 | TRUE | 0.52605 | 100 | 155 | 0 |

Figure 45: Residues with DiscoTope above the threshold of 0.35 — md-0-19-A-discotope3.csv (n = 23)

Residues with DiscoTope ≥ 0.35 — md_0_20_A_discotope3.csv (n = 31)

| PDB | Chain | ResID | AA | DiscoTope | Calibrated | Epitope | RSA | pLDDT | Length | AF2 flag |
|---|---|---|---|---|---|---|---|---|---|---|
| md_0_20_A | A | 46 | R | 0.49601 | 1.81831 | TRUE | 0.55440 | 100 | 155 | 0 |
| md_0_20_A | A | 93 | I | 0.48011 | 1.70808 | TRUE | 0.64051 | 100 | 155 | 0 |
| md_0_20_A | A | 90 | R | 0.47386 | 1.66475 | TRUE | 0.73215 | 100 | 155 | 0 |
| md_0_20_A | A | 106 | E | 0.46975 | 1.63625 | TRUE | 0.72079 | 100 | 155 | 0 |
| md_0_20_A | A | 92 | R | 0.46762 | 1.62149 | TRUE | 0.42802 | 100 | 155 | 0 |
| md_0_20_A | A | 102 | P | 0.46406 | 1.59681 | TRUE | 0.95918 | 100 | 155 | 0 |
| md_0_20_A | A | 100 | A | 0.45898 | 1.56159 | TRUE | 0.93031 | 100 | 155 | 0 |
| md_0_20_A | A | 99 | Q | 0.45487 | 1.53309 | TRUE | 0.72378 | 100 | 155 | 0 |
| md_0_20_A | A | 91 | N | 0.44650 | 1.47506 | TRUE | 0.43035 | 100 | 155 | 0 |
| md_0_20_A | A | 96 | V | 0.44577 | 1.47000 | TRUE | 0.49090 | 100 | 155 | 0 |
| md_0_20_A | A | 152 | W | 0.43838 | 1.41877 | TRUE | 0.71920 | 100 | 155 | 0 |
| md_0_20_A | A | 112 | R | 0.42444 | 1.32212 | TRUE | 0.58852 | 100 | 155 | 0 |
| md_0_20_A | A | 103 | T | 0.42237 | 1.30777 | TRUE | 0.46383 | 100 | 155 | 0 |
| md_0_20_A | A | 109 | D | 0.42110 | 1.29897 | TRUE | 0.70228 | 100 | 155 | 0 |
| md_0_20_A | A | 116 | D | 0.41987 | 1.29044 | TRUE | 0.66631 | 100 | 155 | 0 |
| md_0_20_A | A | 101 | N | 0.41049 | 1.22541 | TRUE | 0.36467 | 100 | 155 | 0 |
| md_0_20_A | A | 107 | T | 0.40820 | 1.20953 | TRUE | 0.31600 | 100 | 155 | 0 |
| md_0_20_A | A | 32 | G | 0.40734 | 1.20357 | TRUE | 0.70219 | 100 | 155 | 0 |
| md_0_20_A | A | 6 | T | 0.40587 | 1.19338 | TRUE | 0.46444 | 100 | 155 | 0 |
| md_0_20_A | A | 39 | Q | 0.40492 | 1.18679 | TRUE | 0.52782 | 100 | 155 | 0 |
| md_0_20_A | A | 43 | V | 0.39455 | 1.11490 | TRUE | 0.52714 | 100 | 155 | 0 |
| md_0_20_A | A | 98 | N | 0.38279 | 1.03337 | TRUE | 0.40247 | 100 | 155 | 0 |
| md_0_20_A | A | 73 | N | 0.38247 | 1.03115 | TRUE | 0.21553 | 100 | 155 | 0 |
| md_0_20_A | A | 38 | Q | 0.37733 | 0.99552 | TRUE | 0.38930 | 100 | 155 | 0 |
| md_0_20_A | A | 105 | A | 0.37164 | 0.95607 | TRUE | 0.45887 | 100 | 155 | 0 |
| md_0_20_A | A | 72 | Y | 0.37079 | 0.95017 | TRUE | 0.44124 | 100 | 155 | 0 |
| md_0_20_A | A | 97 | E | 0.36524 | 0.91170 | TRUE | 0.62185 | 100 | 155 | 0 |
| md_0_20_A | A | 108 | L | 0.36497 | 0.90983 | TRUE | 0.30963 | 100 | 155 | 0 |
| md_0_20_A | A | 119 | V | 0.36086 | 0.88133 | FALSE | 0.55336 | 100 | 155 | 0 |
| md_0_20_A | A | 33 | N | 0.35911 | 0.86920 | FALSE | 0.34884 | 100 | 155 | 0 |
| md_0_20_A | A | 115 | D | 0.35338 | 0.82947 | FALSE | 0.12017 | 100 | 155 | 0 |

Figure 46: Residues with DiscoTope above the threshold of 0.35 — md-0-20-A-discotope3.csv (n = 31)

Residues with DiscoTope ≥ 0.35 — md_0_21_A_discotope3.csv (n = 17)

| PDB | Chain | ResID | AA | DiscoTope | Calibrated | Epitope | RSA | pLDDT | Length | AF2 flag |
|---|---|---|---|---|---|---|---|---|---|---|
| md_0_21_A | A | 72 | Y | 0.48804 | 1.91082 | TRUE | 0.51879 | 100 | 155 | 0 |
| md_0_21_A | A | 100 | A | 0.47417 | 1.80660 | TRUE | 0.98278 | 100 | 155 | 0 |
| md_0_21_A | A | 109 | D | 0.47237 | 1.79308 | TRUE | 0.65815 | 100 | 155 | 0 |
| md_0_21_A | A | 108 | L | 0.46049 | 1.70381 | TRUE | 0.41812 | 100 | 155 | 0 |
| md_0_21_A | A | 103 | T | 0.45138 | 1.63536 | TRUE | 0.39768 | 100 | 155 | 0 |
| md_0_21_A | A | 106 | E | 0.43629 | 1.52197 | TRUE | 0.71179 | 100 | 155 | 0 |
| md_0_21_A | A | 46 | R | 0.43577 | 1.51806 | TRUE | 0.70497 | 100 | 155 | 0 |
| md_0_21_A | A | 50 | E | 0.42852 | 1.46359 | TRUE | 0.67623 | 100 | 155 | 0 |
| md_0_21_A | A | 85 | G | 0.41078 | 1.33029 | TRUE | 0.79369 | 100 | 155 | 0 |
| md_0_21_A | A | 101 | N | 0.40102 | 1.25696 | TRUE | 0.33843 | 100 | 155 | 0 |
| md_0_21_A | A | 98 | N | 0.37439 | 1.05686 | TRUE | 0.22547 | 100 | 155 | 0 |
| md_0_21_A | A | 15 | S | 0.35940 | 0.94423 | TRUE | 0.48513 | 100 | 155 | 0 |
| md_0_21_A | A | 33 | N | 0.35833 | 0.93619 | TRUE | 0.42157 | 100 | 155 | 0 |
| md_0_21_A | A | 112 | R | 0.35335 | 0.89877 | FALSE | 0.56220 | 100 | 155 | 0 |
| md_0_21_A | A | 54 | P | 0.35277 | 0.89441 | FALSE | 0.83967 | 100 | 155 | 0 |
| md_0_21_A | A | 152 | W | 0.35261 | 0.89321 | FALSE | 0.74441 | 100 | 155 | 0 |
| md_0_21_A | A | 29 | N | 0.35034 | 0.87615 | FALSE | 0.38198 | 100 | 155 | 0 |

Figure 47: Residues with DiscoTope above the threshold of 0.35 — md-0-21-A-discotope3.csv (n = 17)
