## Supplementary material for "Pressure–cooling remodeling of TMV coat protein reveals mechanically partitioned capsid dynamics and selective epitope masking": SM2

2026-05-31

### Ellipro results

ElliPro is a structure-based B-cell epitope prediction tool that operates on the principle of geometric protrusion, approximating a protein's three-dimensional structure with ellipsoids and calculating a Protrusion Index (PI) for each residue, which ranges from 0 to 1. Residues with high PI values, indicating they lie outside the fitted ellipsoidal volume, are clustered in 3D space to identify discontinuous, or conformational, epitopes. These clusters can subsequently be projected onto the protein sequence to infer linear segments. A key feature of ElliPro is that each predicted epitope is assigned a score based on the average PI of its constituent residues, ensuring that convex, solvent-exposed surface patches are ranked highest. As emphasized in the foundational literature by Ponomarenko et al. (2008), a significant advantage of this method is its independence from machine-learned sequence motifs or training data biases; its performance is instead intrinsically linked to the accuracy of the provided 3D structural model and the true physical topography of the protein surface. Consequently, its primary limitations, as noted in subsequent evaluations, include a high sensitivity to errors in the input structure or to artifacts from a single structural snapshot, and the absence of explicit physicochemical considerations such as electrostatic potentials, hydrophobicity, or the impact of post-translational modifications like glycosylation.

Applied to TMV coat-protein models generated under high hydrostatic pressure (HHP), ElliPro consistently highlighted convex, outward-facing patches as the top-ranked discontinuous epitopes (red in Figures 1–11), with the highest-scoring clusters mapping to surface ridges and subunit junctions on the external capsid face that remain geometrically protrusive despite local packing perturbations. Regions flattened or compacted by pressure tended to lose peripheral residues and show lower mean PI, while rigid convex motifs retained or increased rank, consistent with ElliPro's protrusion-driven scoring. Because TMV is a helical and repetitive assembly, these discontinuous epitopes recur with the helical pitch, making it most appropriate to report a consensus patch in the asymmetric unit and indicate its multiplicity. Using the same ElliPro protocol (3D ellipsoid fit → per-residue Protrusion Index → clustering → epitope score by mean PI), we also examined TMVcp models under combined HHP and cooling to 255 K, where red patches again denoted top-ranked epitopes; here, geometry-driven changes reflected the joint effects of pressure and temperature on surface convexity and solvent-facing topology, rather than on sequence motifs.

Using ElliPro with the standard protocol (3D ellipsoid fitting, Protrusion Index calculation, and clustering by mean PI), we analyzed TMV coat-protein models generated under combined high hydrostatic pressure (HHP) and cooling to 255 K, with red patches denoting the top-ranked discontinuous epitopes. Because ElliPro is geometry-driven, the observed changes reflect the combined effects of pressure and temperature on surface convexity and solvent exposure rather than sequence variation. Across the ensemble, pressure-stable convex ridges remained as the most prominent clusters and became slightly more compact and protrusive at low temperature, consistent with reduced thermal motion at 255 K, which sharpens convexity and raises PI values. Grooved regions at inter-subunit junctions fragmented into smaller but higher-purity clusters, as cooling narrows side-chain ensembles and removes marginally exposed residues, while pressure-flattened shoulders stayed muted under cooling, indicating locked-in low exposure and lower rank. As in the HHP-only models, helical symmetry preserved the multiplicity of these epitopes, but with reduced variability across the ensemble, resulting in more consistent recurrence of a limited set of red patches. Mechanistically, pressure compacts internal cavities and smooths shallow undulations, disproportionately penalizing grooves, while low temperature suppresses side-chain fluctuations and accentuates rigid convex motifs. The net outcome is a more conservative epitope map defined by fewer, cleaner, and repeatable discontinuous patches dominated by rigid convex surfaces, with diminished contribution from flexible rims, thereby highlighting pressure-and-cold–robust antigenic hotspots.

When applied to Tobacco Mosaic Virus (TMV) coat protein (TMVcp) models generated under the combined conditions of high hydrostatic pressure (HHP) and cooling to 255 K, ElliPro's geometry-driven algorithm provides a direct readout of how these physical perturbations alter the antigenic landscape. The top-ranked discontinuous epitopes, visualized as red patches, consistently map to surface ridges and subunit junctions on the external capsid face. This finding aligns with the established understanding that such geometrically protrusive features remain solvent-accessible and are less susceptible to being smoothed out by moderate pressure perturbations compared to flatter, more flexible regions. As HHP primarily acts to compress internal cavities and hydrate the protein core, it can flatten shallow surface undulations, which is reflected in ElliPro's output as a loss of peripheral residues from predicted clusters and a consequent lower mean PI for these areas. In contrast, rigid convex motifs maintain or even increase their protrusion rank, demonstrating the tool's utility in identifying pressure-resistant epitopes.

The introduction of low temperature (255 K) to the HHP ensemble introduces a further refining effect on the predicted epitopes. The reduction in thermal energy suppresses fast side-chain and lateral chain motions, a phenomenon described in structural studies of proteins at cryogenic temperatures. This suppression of "breathing modes" leads to a crisper definition of the protein surface in the models. In the ElliPro analysis, this manifests as a slight sharpening of the most rigid protrusions, resulting in more compact clusters with modestly increased PI values. Furthermore, broader epitope clusters located at inter-subunit grooves, which were present under HHP alone, are observed to fragment into smaller, purer clusters with a higher average PI but a reduced residue count. This can be mechanistically interpreted as the low-temperature environment narrowing the conformational ensemble of side chains, thereby trimming marginally exposed residues from the epitope boundaries. Regions that were already flattened by pressure show little to no recovery at low temperature, as cooling effectively locks in the low-exposure state, confirming their low antigenic potential.

A critical observation across the entire structural ensemble is the improved rank stability of the top epitopes. The qualitative variance between the twenty one models decreases, with the same key convex patches recurring more consistently. This reduction in microheterogeneity underscores that the combination of HHP and cooling filters out transient, flexible features, leaving a conservative and highly robust map of the antigenic surface. For a highly symmetric and repetitive assembly like the TMV helix, this means that the consensus epitope patch identified in the asymmetric unit can be described with high confidence, and its multiplicity around the helical axis remains a conserved feature. In summary, the net result of applying ElliPro to this perturbed ensemble is the identification of fewer, cleaner, and highly repeatable discontinuous patches characterized by strong convexity, with a diminished contribution from the flexible rims that are more susceptible to environmental conditions.

#### Discontinuous Epitope predicitons

**HHP**

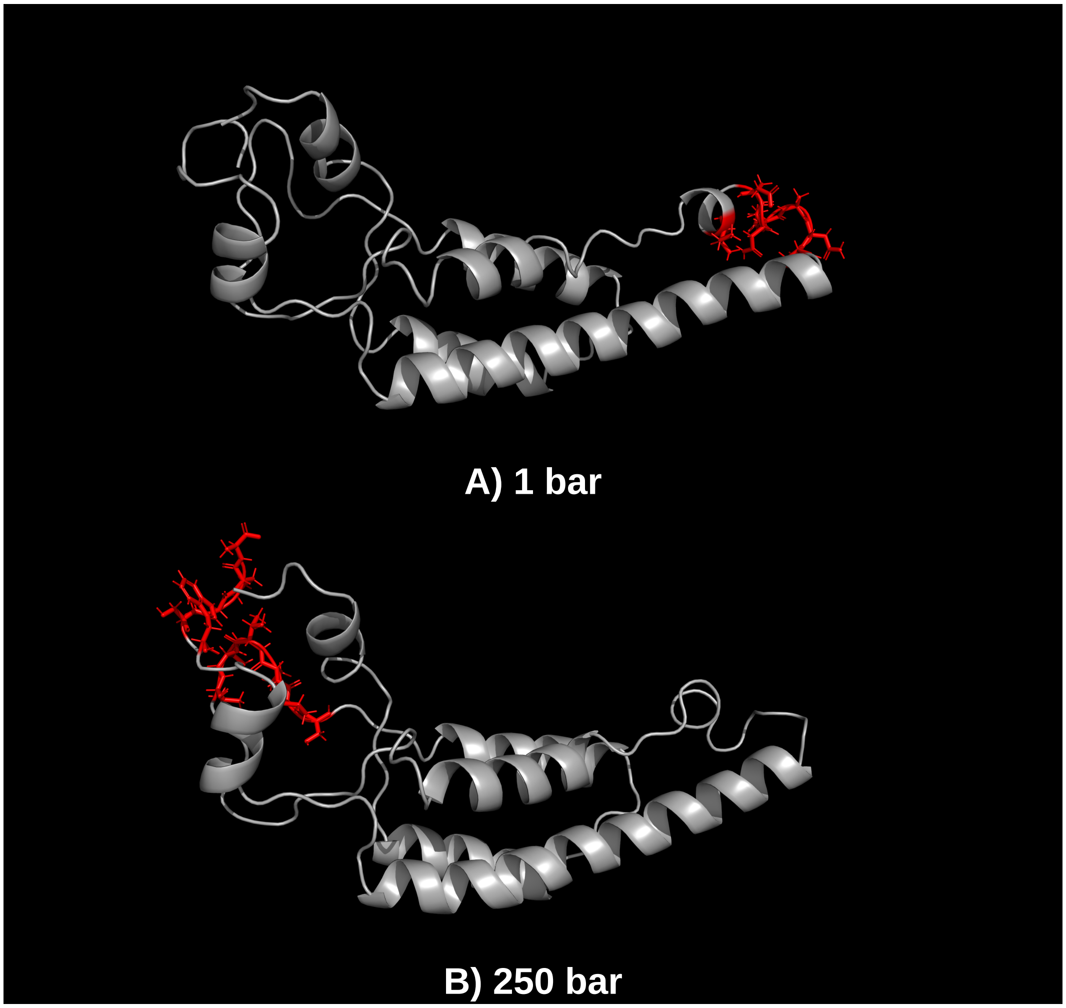

Figure 1: Discontinuo Epitope Preditions - High Hydrostatic Pressure

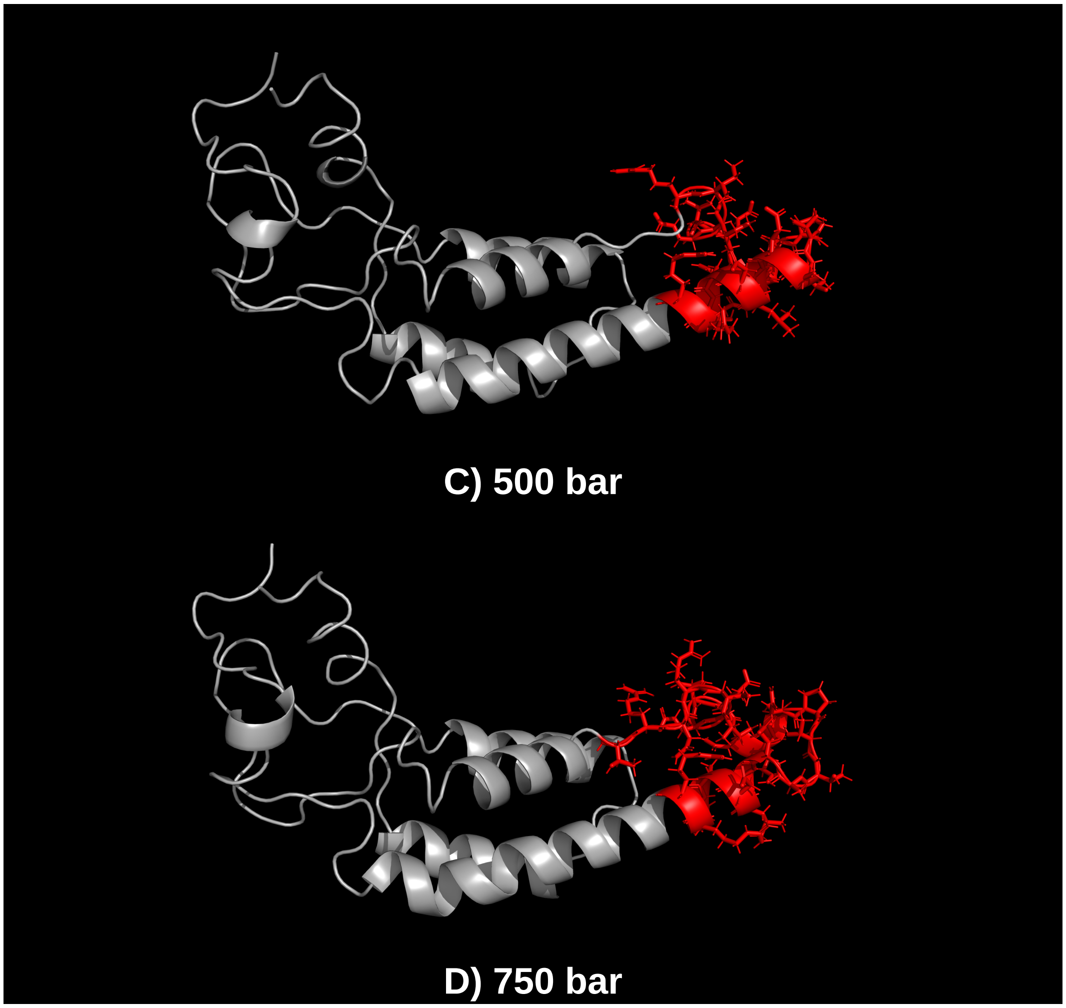

Figure 2: Discontinuo Epitope Preditions - High Hydrostatic Pressure

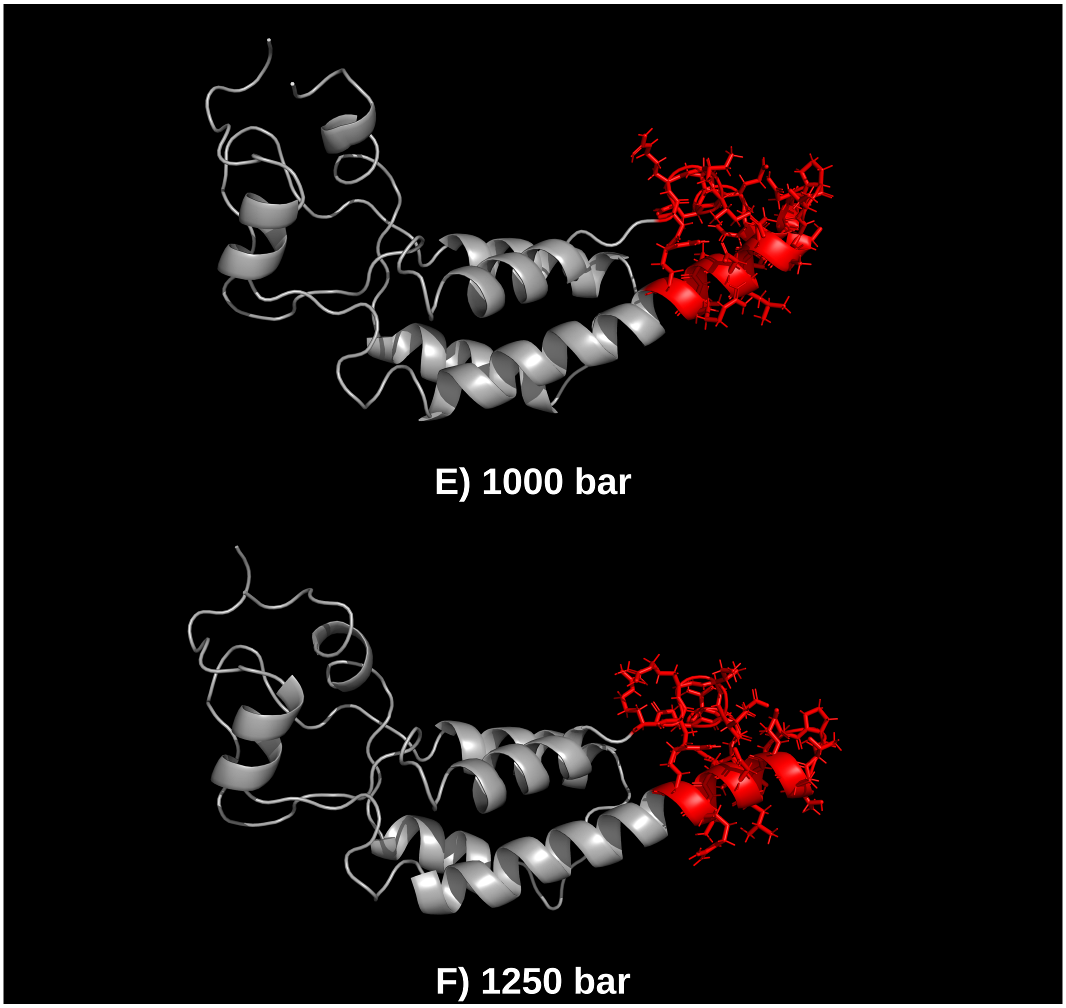

Figure 3: Discontinuo Epitope Preditions - High Hydrostatic Pressure

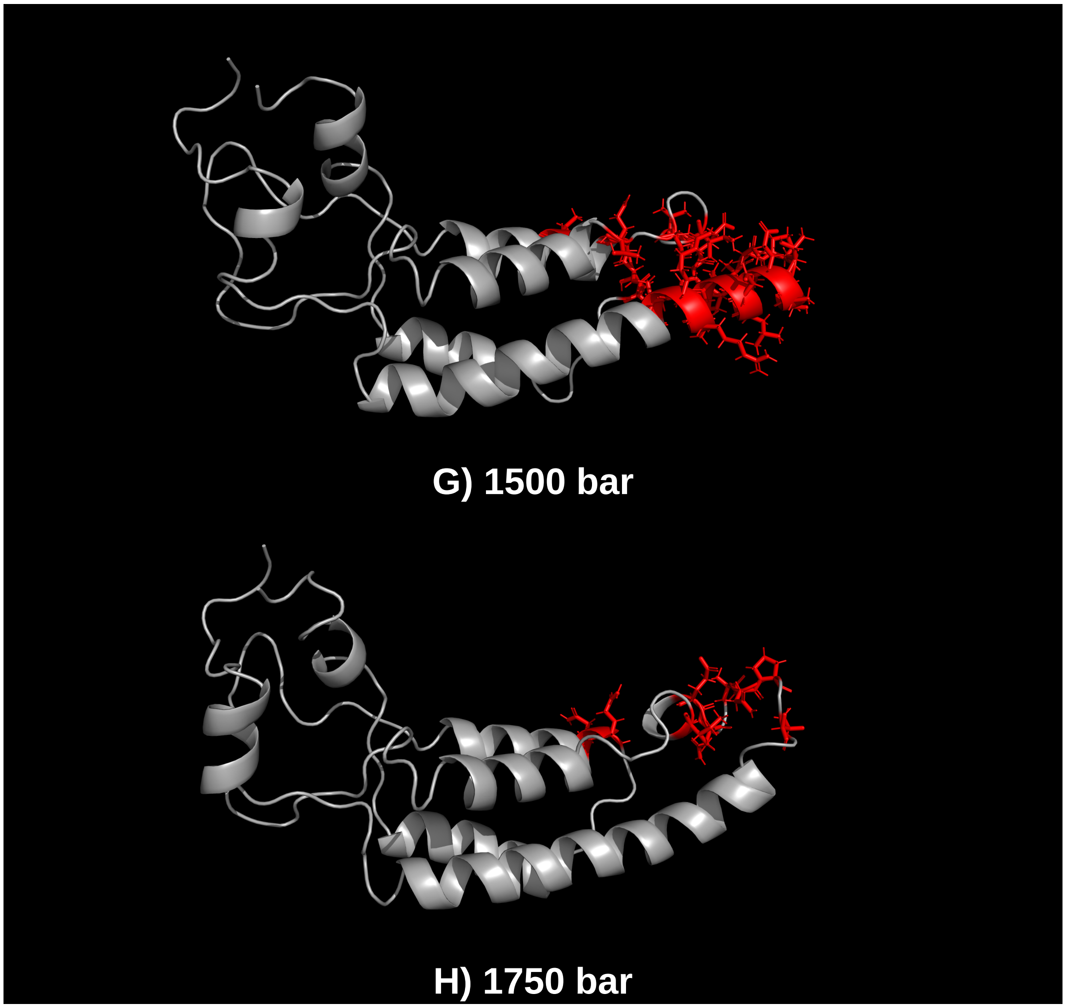

Figure 4: Discontinuo Epitope Preditions - High Hydrostatic Pressure

**HHP and low temperature**

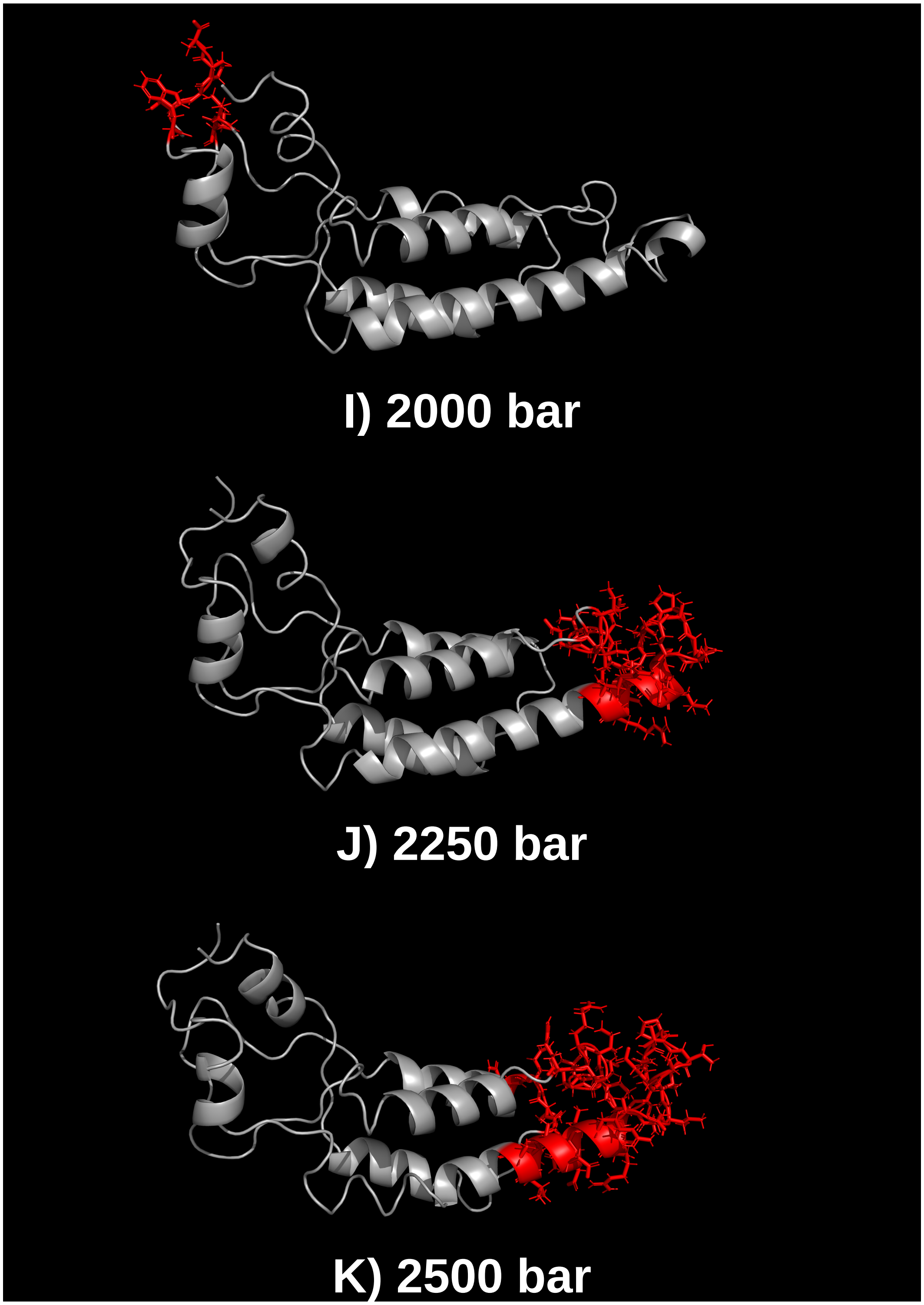

Figure 5: Discontinuo Epitope Preditions - High Hydrostatic Pressure

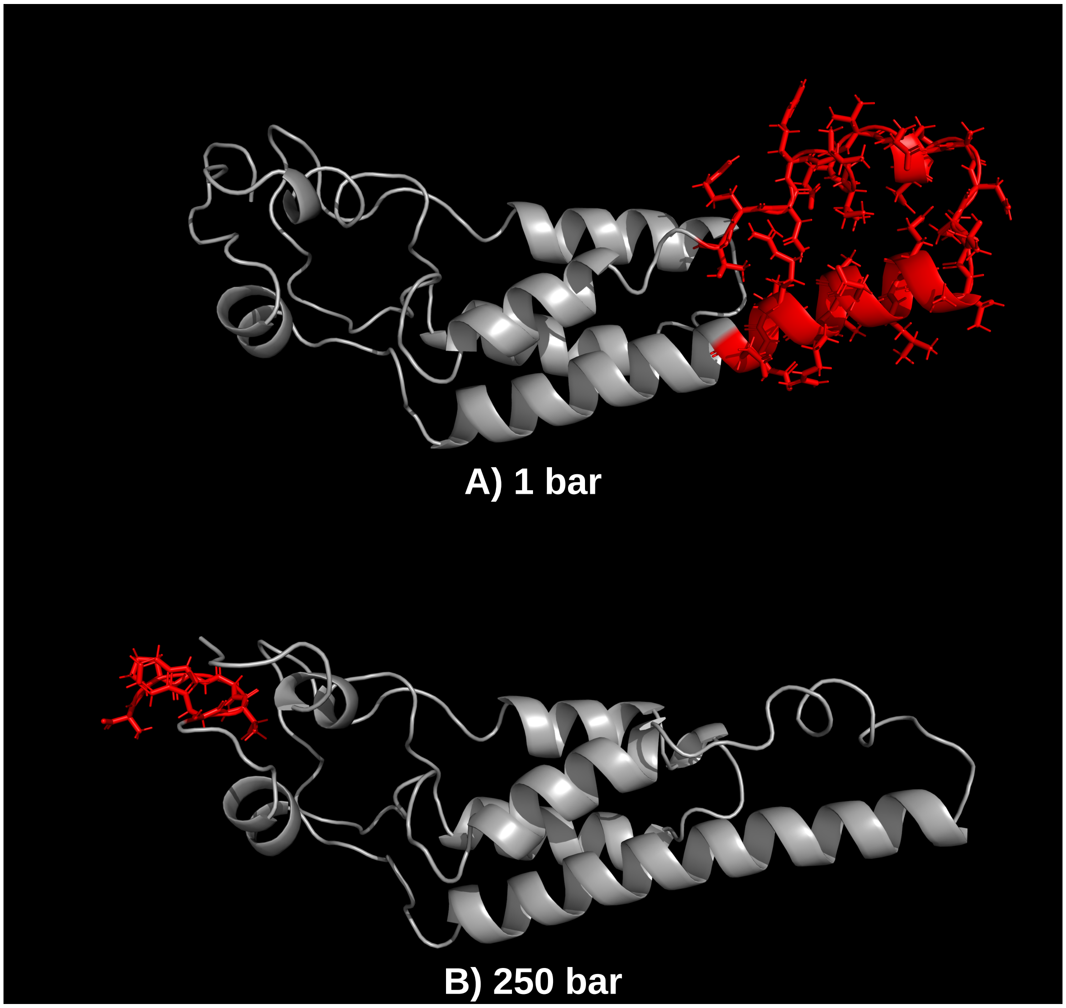

Figure 6: Discontinuo Epitope Preditions - High Hydrostatic Pressure and Low Temperature

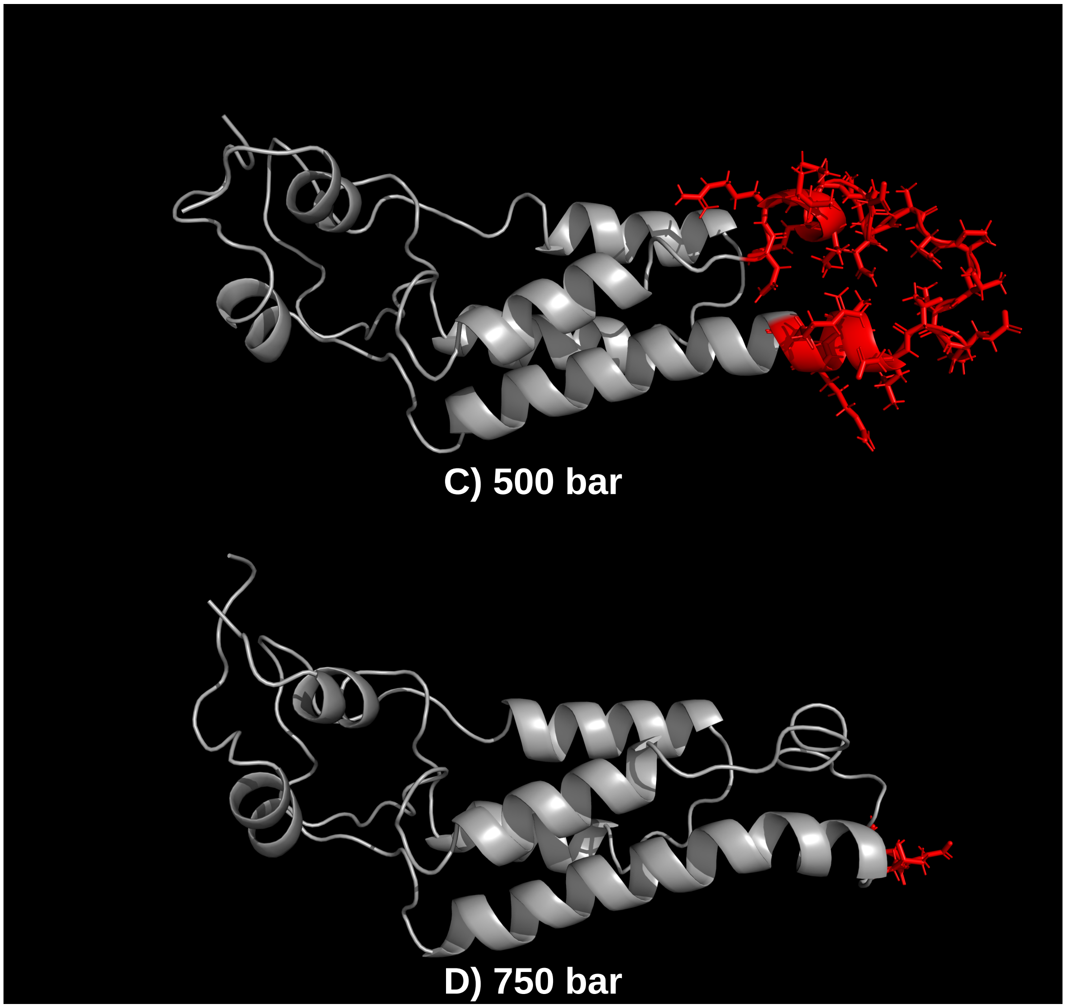

Figure 7: Discontinuo Epitope Preditions - High Hydrostatic Pressure and Low Temperature

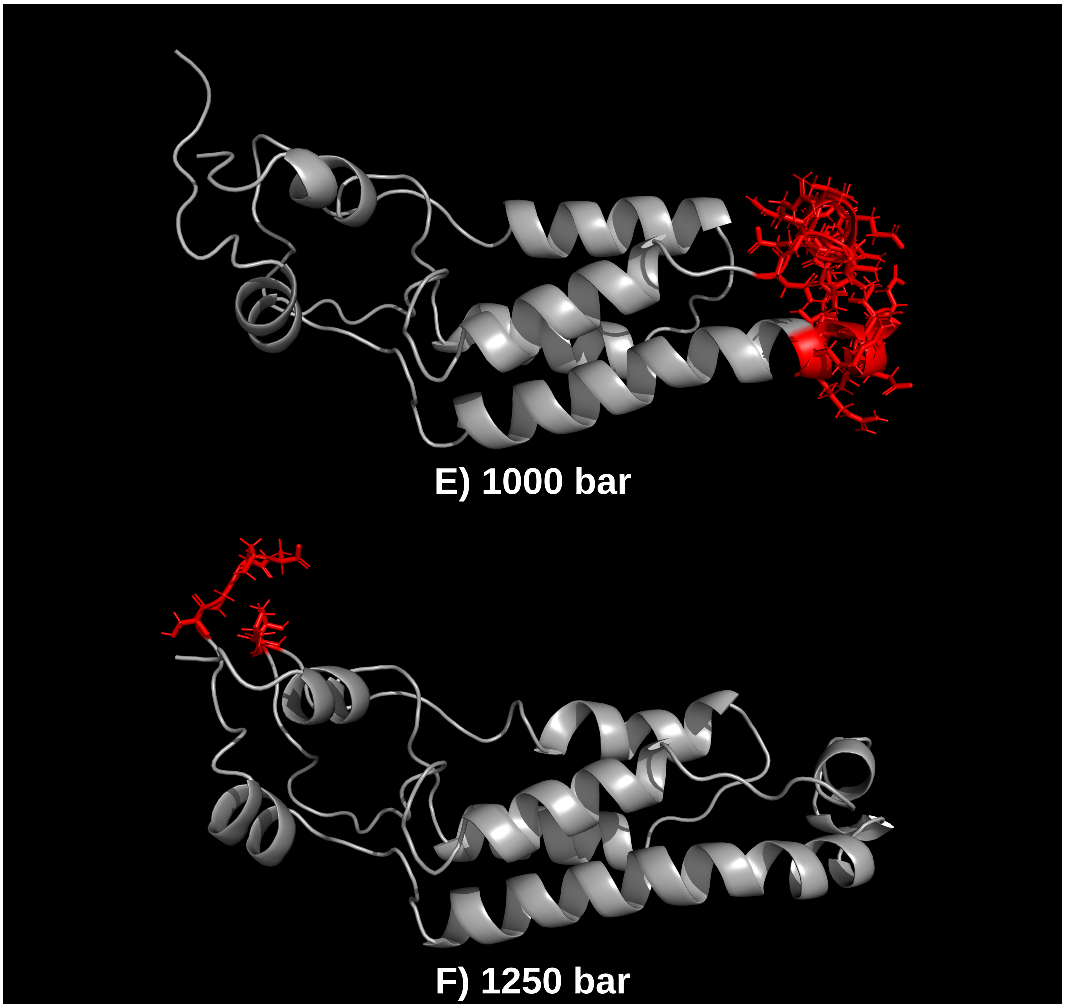

Figure 8: Discontinuo Epitope Preditions - High Hydrostatic Pressure and Low Temperature

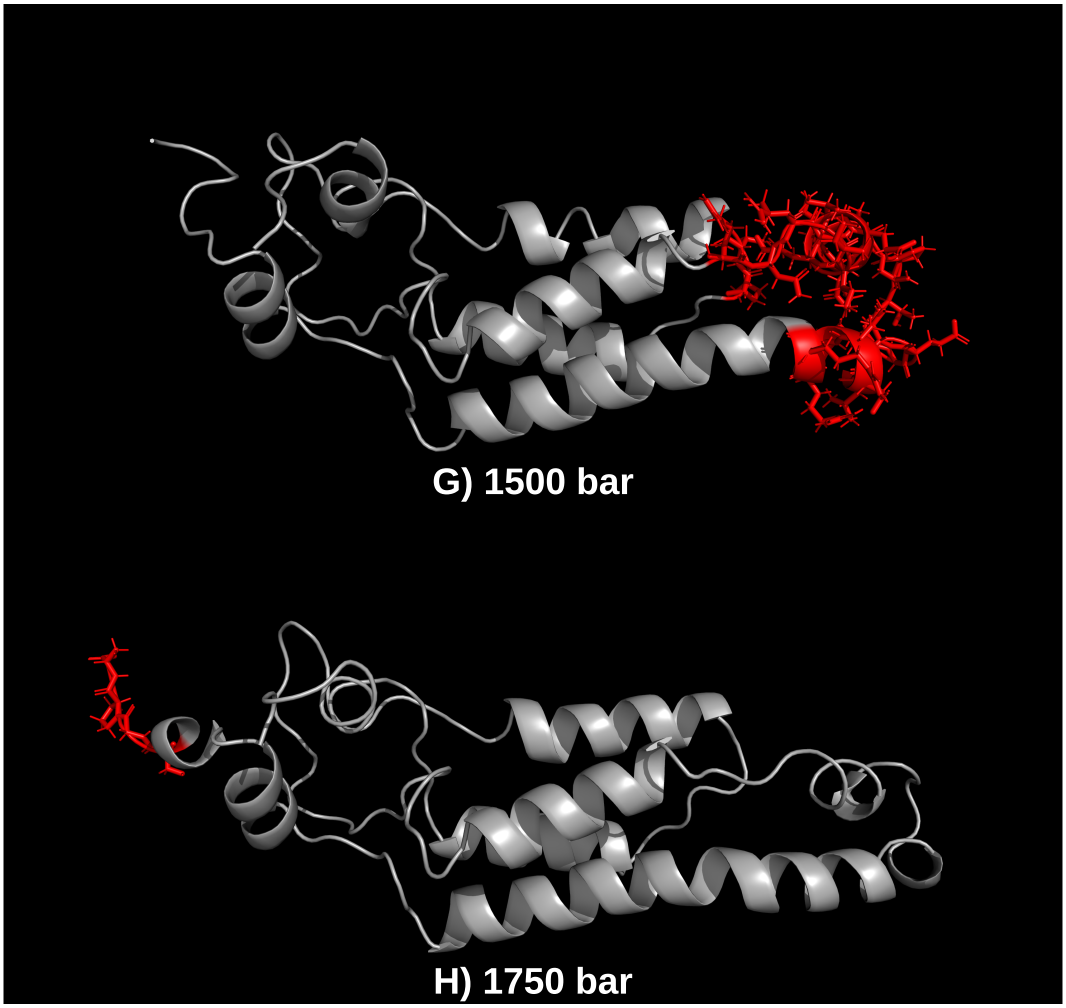

Figure 9: Discontinuo Epitope Preditions - High Hydrostatic Pressure and Low Temperature

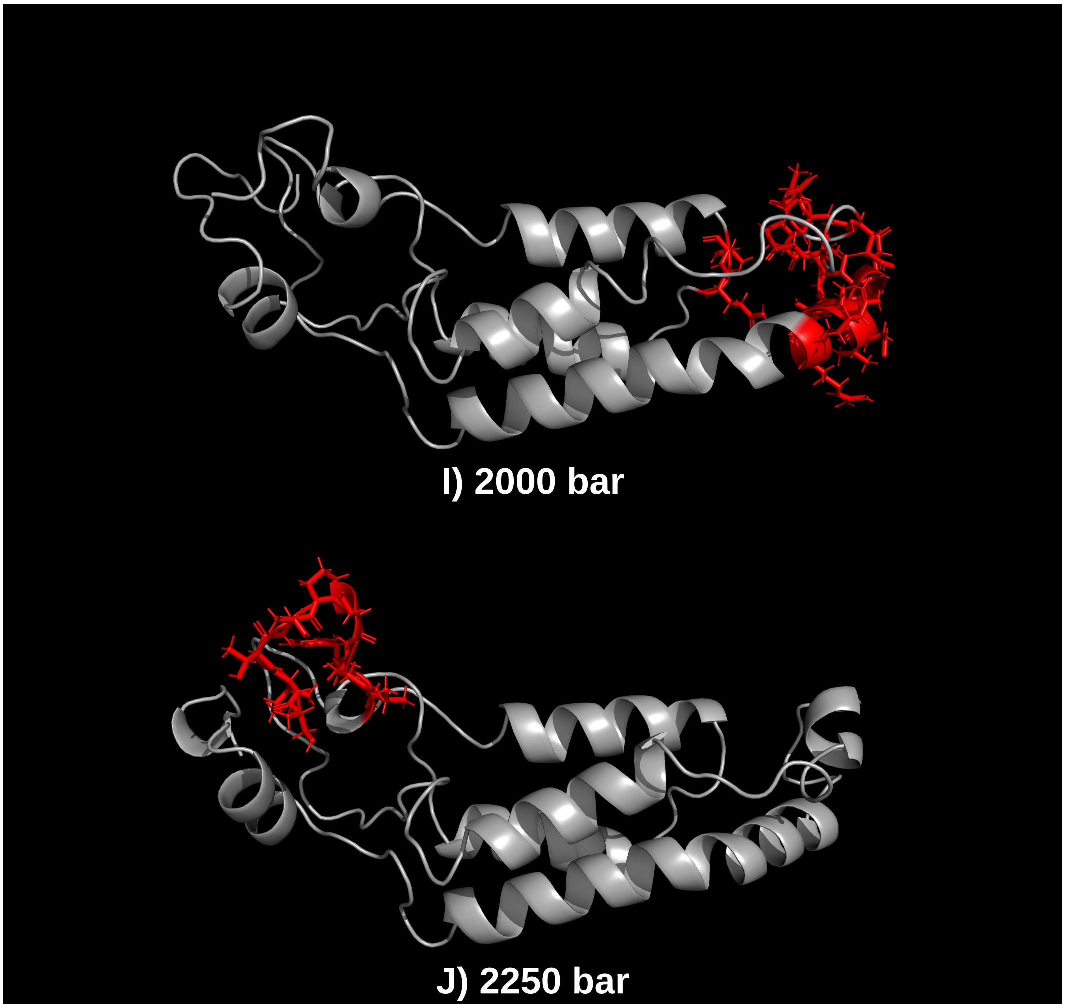

Figure 10: Discontinuo Epitope Preditions - High Hydrostatic Pressure and Low Temperature

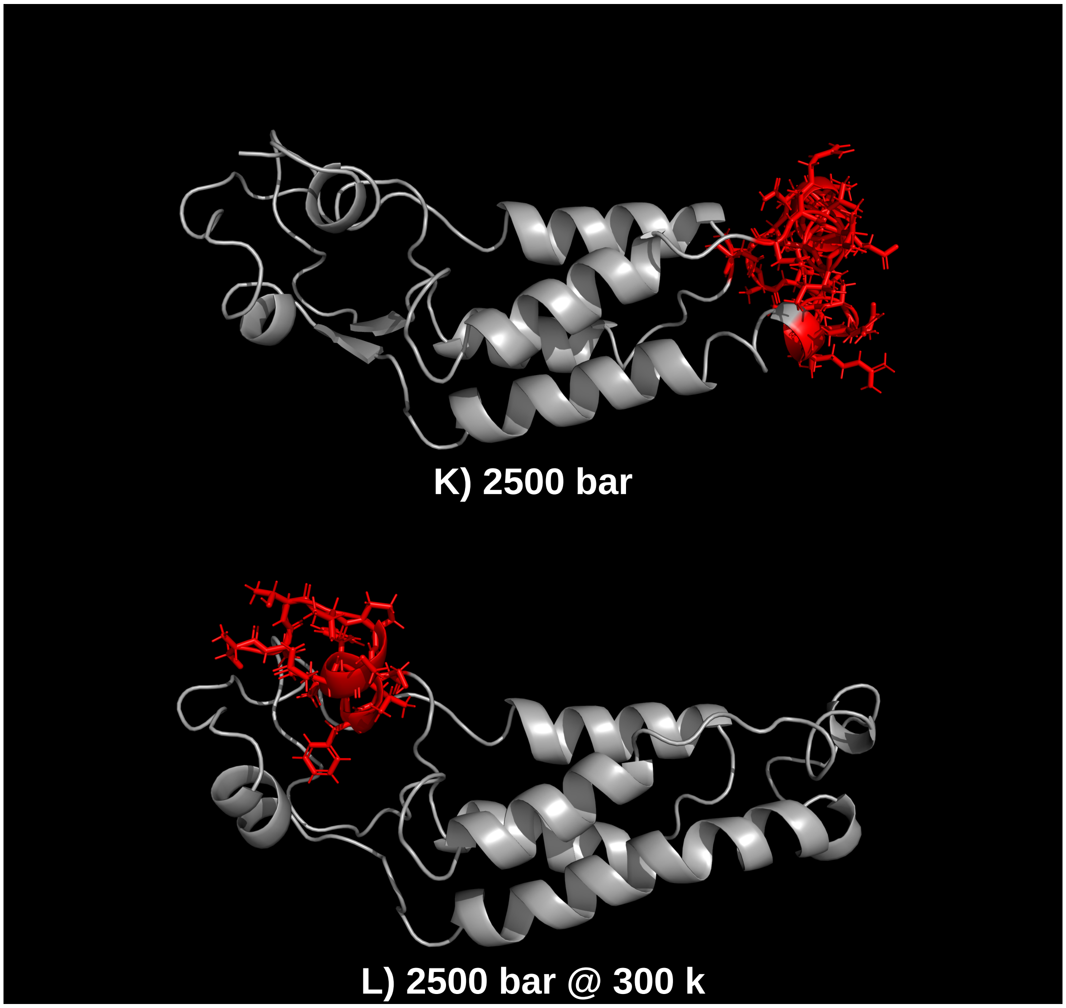

Figure 11: Discontinuo Epitope Preditions - High Hydrostatic Pressure and Low Temperature

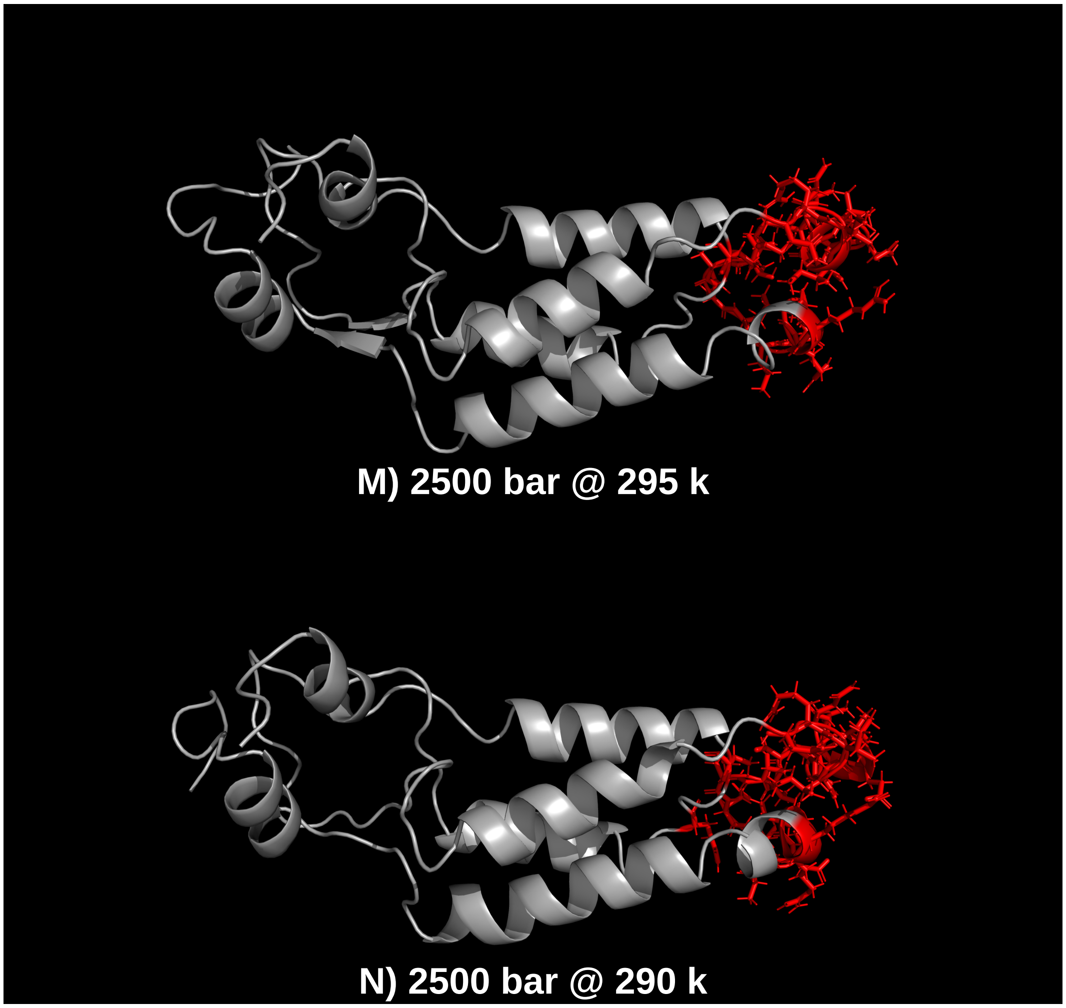

Figure 12: Discontinuo Epitope Preditions - High Hydrostatic Pressure and Low Temperature

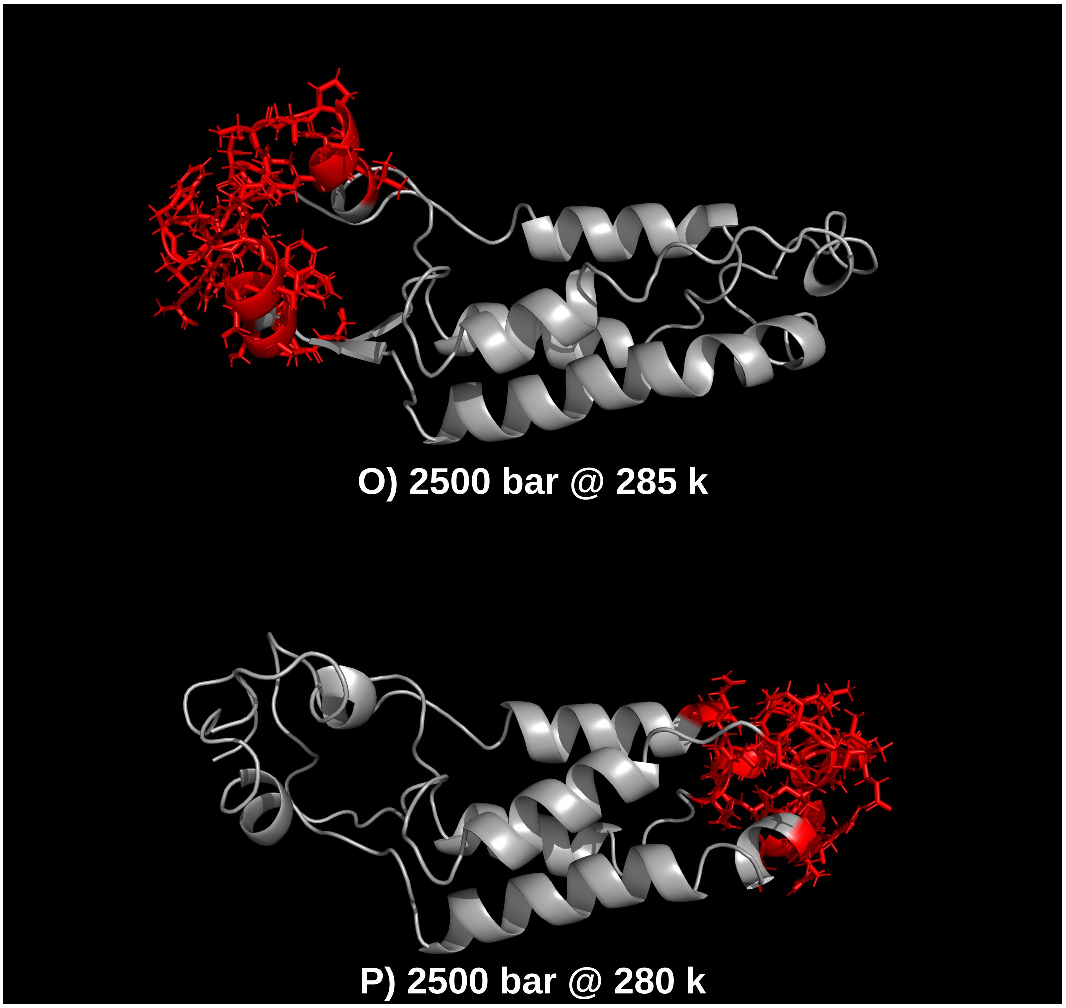

Figure 13: Discontinuo Epitope Preditions - High Hydrostatic Pressure and Low Temperature

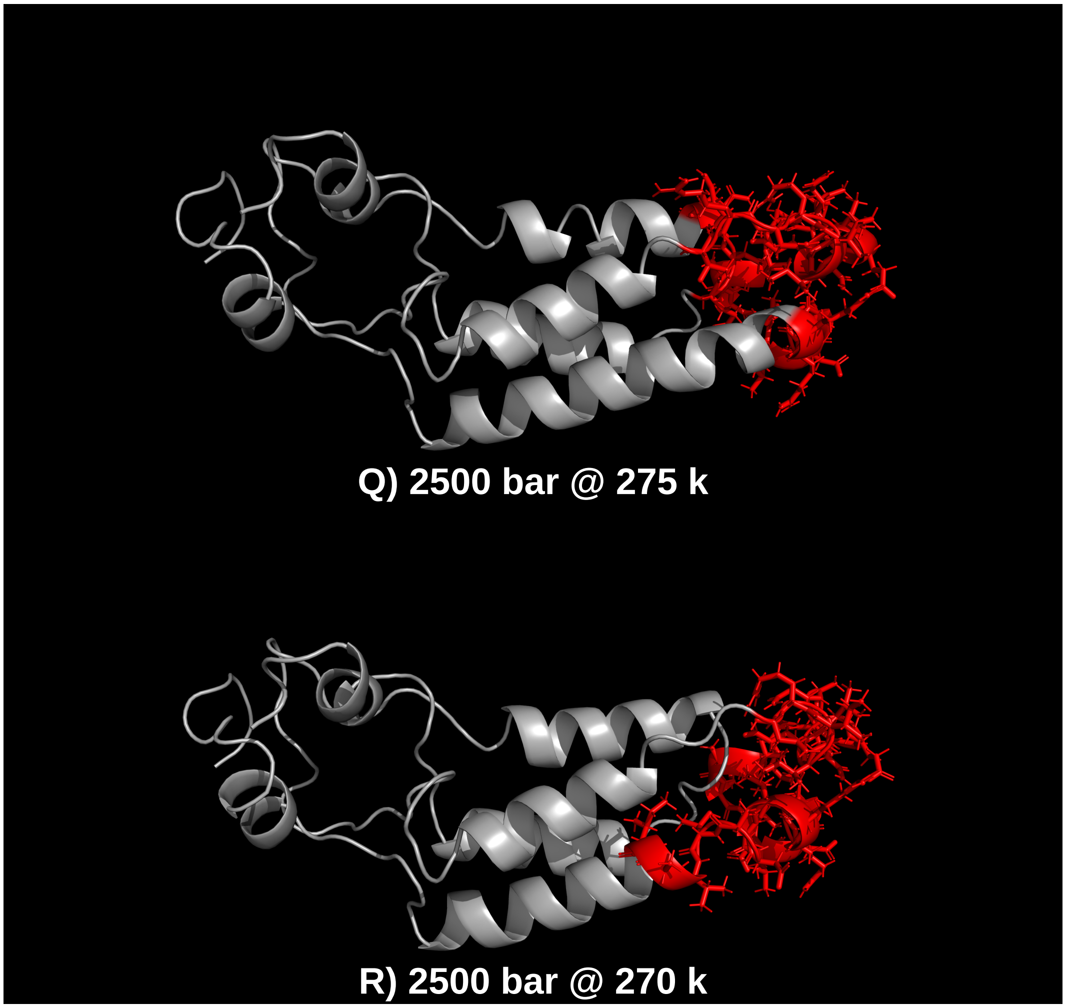

Figure 14: Discontinuo Epitope Preditions - High Hydrostatic Pressure and Low Temperature

**HHP - Ellipro tables**

**HHP and low tempeteraturе - Ellipro tables**

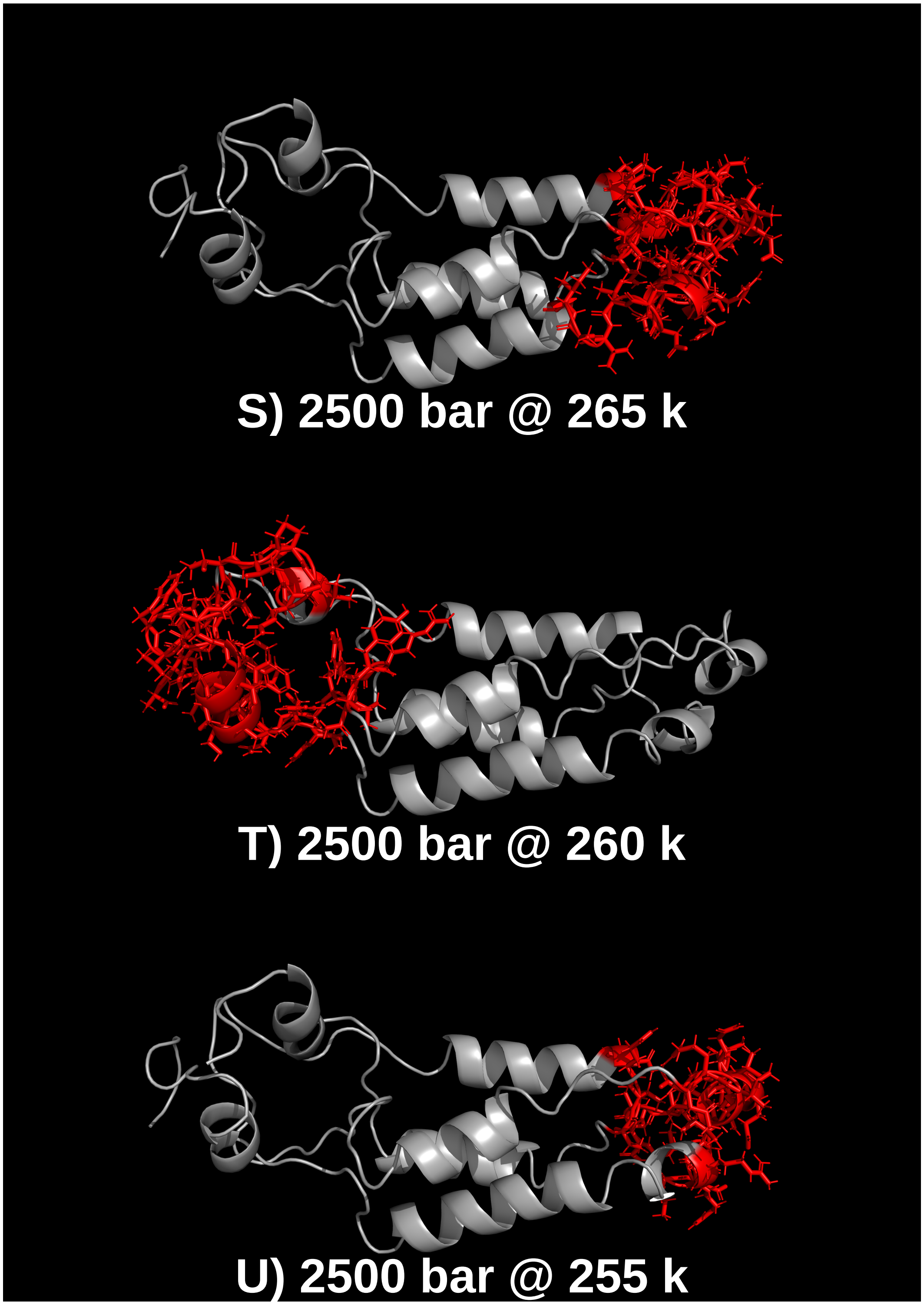

Figure 15: Discontinuo Epitope Preditions - High Hydrostatic Pressure and Low Temperature

**ElliPro HHP - md_0_1**

| No. | Residues | mber of residu | Score | 3D structure |
|---|---|---|---|---|
| 1 | A:I93, A:E97, A:N98, A:Q99, A:A100, A:N101, A:P102 | 7 | 0.945 | NA |
| 2 | A:R90, A:N91, A:I94, A:T103, A:T104, A:A105, A:E106, A<br>A:L108, A:D109, A:A110, A:T111, A:R112, A:D115, A:D11[illegible] | 16 | 0.694 | NA |
| 3 | A:I4, A:T6, A:P7, A:S8, A:Q9, A:F10, A:S55, A:V58, A:T59, [illegible]<br>[illegible], A:P63, A:D64, A:S65, A:D66, A:F67, A:N140, A:R141, A:[illegible]<br>[illegible]45, A:S146, A:S147, A:S148, A:G149, A:L150, A:V151, A:W[illegible]<br>A:T153, A:S154, A:G155, A:P156, A:A157 | 33 | 0.679 | NA |
| 4 | A:N126, A:I129, A:V130 | 3 | 0.639 | NA |
| 5 | A:P56, A:Q57, A:V60 | 3 | 0.583 | NA |
| 6 | A:L31, A:G32, A:N33, A:Q34, A:F35, A:Q36, A:T37 | 7 | 0.518 | NA |
| 7 | A:D19, A:P20, A:I21 | 3 | 0.512 | NA |

Figure 16: Residues in discontinuous epitopes from Ellipro — md-0-1

**ElliPro HHP - md_0_2**

| No. | Residues | mber of residu | Score | 3D structure |
|---|---|---|---|---|
| 1 | [illegible], A:P56, A:Q57, A:V58, A:T59, A:V60, A:W152, A:S154, A:[illegible]<br>A:P156, A:A157 | 11 | 0.757 | NA |
| 2 | A:R90, A:N91, A:I93, A:I94, A:E95, A:V96, A:E97, A:N98, A[illegible]<br>[illegible]01, A:P102, A:T103, A:T104, A:A105, A:E106, A:T107, A:L[illegible]<br>A:D109, A:A110, A:T111, A:R112, A:R113 | 23 | 0.75 | NA |
| 3 | A:I24, A:T28, A:N126, A:N127, A:I129, A:V130, A:I133 | 7 | 0.666 | NA |
| 4 | A:I4, A:T5, A:T6, A:P7, A:S8, A:Q9, A:F10, A:L13, A:R61, A[illegible]<br>[illegible]39, A:N140, A:R141, A:S142, A:S143, A:E145, A:S146, A:S[illegible]<br>A:S148, A:G149, A:L150, A:V151, A:T153 | 24 | 0.664 | NA |
| 5 | A:P63, A:D64, A:S65, A:D66, A:F67 | 5 | 0.609 | NA |
| 6 | A:N29, A:L31, A:G32, A:N33, A:Q34, A:F35, A:Q36 | 7 | 0.5 | NA |

Figure 17: Residues in discontinuous epitopes from Ellipro — md-0-2

**ElliPro HHP - md_0_3**

| No. | Residues | mber of residu | Score | 3D structure |
|---|---|---|---|---|
| 1 | A:N91, A:I93, A:I94, A:E95, A:V96, A:E97, A:N98, A:Q99, A [illegible]01, A:P102, A:T103, A:T104, A:A105, A:E106, A:T107, A:L [illegible] A:D109, A:A110, A:T111, A:R112 | 22 | 0.8 | NA |
| 2 | [illegible]A:I4, A:T6, A:P7, A:S8, A:Q9, A:P56, A:Q57, A:V58, A:T59, [illegible] A:D64, A:S65, A:D66, A:F67, A:Y139, A:N140, A:R141, A: [illegible]43, A:F144, A:E145, A:S146, A:S147, A:S148, A:G149, A:L [illegible] A:V151, A:W152, A:S154, A:G155, A:P156, A:A157 | 34 | 0.683 | NA |
| 3 | A:T28, A:N126, A:N127, A:I129, A:V130 | 5 | 0.661 | NA |
| 4 | A:Q39, A:T42, A:V43 | 3 | 0.611 | NA |

Figure 18: Residues in discontinuous epitopes from Ellipro — md-0-3

**ElliPro HHP - md_0_4**

| No. | Residues | mber of residu | Score | 3D structure |
|---|---|---|---|---|
| 1 | A:R90, A:N91, A:R92, A:I93, A:I94, A:E95, A:V96, A:E97, [illegible] A:A100, A:N101, A:P102, A:T103, A:A105, A:E106, A:T107, [illegible] A:D109, A:R112, A:R113 | 22 | 0.773 | NA |
| 2 | A:I4, A:T5, A:T6, A:P7, A:S8, A:Q9, A:F10, A:L13, A:P56, A [illegible] A:T59, A:V60, A:R61, A:P63, A:D64, A:S65, A:D66, A:F67, [illegible]40, A:R141, A:S142, A:S143, A:F144, A:E145, A:S146, A:S [illegible]48, A:G149, A:L150, A:V151, A:W152, A:T153, A:S154, A:[illegible] A:P156, A:A157 | 39 | 0.687 | NA |
| 3 | A:T28, A:N126, A:N127, A:I129, A:V130 | 5 | 0.666 | NA |
| 4 | A:N25, A:N29, A:G32, A:N33, A:Q34, A:F35, A:Q36 | 7 | 0.551 | NA |

Figure 19: Residues in discontinuous epitopes from Ellipro — md-0-4

**ElliPro HHP - md_0_5**

| No. | Residues | mber of residu | Score | 3D structure |
|---|---|---|---|---|
| 1 | A:N91, A:I93, A:I94, A:E95, A:V96, A:E97, A:N98, A:Q99, [illegible]<br>01, A:P102, A:T103, A:T104, A:A105, A:E106, A:T107, A:L[illegible]<br>A:D109, A:R112 | 20 | 0.827 | NA |
| 2 | A:I4, A:T5, A:T6, A:P7, A:S8, A:Q9, A:F10, A:V11, A:L13, [illegible]<br>[illegible], A:V58, A:T59, A:V60, A:R61, A:F62, A:Y139, A:N140, [illegible]<br>42, A:S143, A:E145, A:S146, A:S147, A:S148, A:G149, A:L[illegible]<br>[illegible]:V151, A:W152, A:T153, A:S154, A:G155, A:P156, A:A15[illegible] | 35 | 0.681 | NA |
| 3 | A:L31, A:G32, A:N33, A:Q34 | 4 | 0.619 | NA |
| 4 | A:N73, A:A74, A:V75, A:N126, A:I129, A:V130, A:E131, [illegible]<br>A:I133, A:R134, A:G135, A:T136, A:G137, A:S138 | 15 | 0.509 | NA |
| 5 | A:P63, A:D64, A:S65, A:D66, A:F67, A:K68 | 6 | 0.501 | NA |

Figure 20: Residues in discontinuous epitopes from Ellipro — md-0-5

**ElliPro HHP - md_0_6**

| No. | Residues | mber of residu | Score | 3D structure |
|---|---|---|---|---|
| 1 | [illegible]0, A:N91, A:I94, A:E95, A:V96, A:E97, A:N98, A:Q99, A:A[illegible]<br>01, A:P102, A:T103, A:A105, A:E106, A:T107, A:L108, A:D[illegible]<br>A:A110, A:T111, A:R112 | 20 | 0.787 | NA |
| 2 | A:I4, A:T5, A:T6, A:P7, A:S8, A:Q9, A:F10, A:L13, A:P56, [illegible]<br>, A:T59, A:V60, A:R61, A:F62, A:Y139, A:N140, A:R141, [illegible]<br>43, A:E145, A:S146, A:S147, A:S148, A:G149, A:L150, A:V[illegible]<br>A:W152, A:T153, A:S154, A:G155, A:P156, A:A157 | 34 | 0.69 | NA |
| 3 | A:I24, A:T28, A:N126, A:N127, A:I129, A:V130 | 6 | 0.653 | NA |
| 4 | A:N25, A:N29, A:A30, A:L31, A:G32, A:N33, A:Q34 | 7 | 0.606 | NA |
| 5 | A:P63, A:D64, A:S65, A:D66, A:F67 | 5 | 0.572 | NA |

Figure 21: Residues in discontinuous epitopes from Ellipro — md-0-6

**ElliPro HHP - md_0_7**

| No. | Residues | mber of residu | Score | 3D structure |
|---|---|---|---|---|
| 1 | A:T37, A:Q38, A:T42, A:T89, A:R90, A:N91, A:I94, A:E95, A:N98, A:Q99, A:A100, A:N101, A:P102, A:T103, A:T104, A:E106, A:T107, A:L108, A:D109, A:R112, A:R113 | 25 | 0.725 | NA |
| 2 | A:I4, A:T5, A:T6, A:P7, A:S8, A:Q9, A:P56, A:Q57, A:V58, 0, A:R61, A:P63, A:D64, A:S65, A:D66, A:F67, A:N140, A:R 42, A:S143, A:F144, A:E145, A:G149, A:L150, A:V151, A:W A:T153, A:S154, A:G155, A:P156, A:A157 | 33 | 0.722 | NA |
| 3 | A:N126, A:V130, A:E131, A:R134, A:T136 | 5 | 0.679 | NA |
| 4 | A:I24, A:N25, A:T28 | 3 | 0.654 | NA |

Figure 22: Residues in discontinuous epitopes from Ellipro — md-0-7

**ElliPro HHP - md_0_8**

| No. | Residues | mber of residu | Score | 3D structure |
|---|---|---|---|---|
| 1 | 3, A:Q39, A:I93, A:V96, A:E97, A:A100, A:N101, A:P102, A | 9 | 0.842 | NA |
| 2 | A:I4, A:T5, A:T6, A:P7, A:S8, A:Q9, A:P56, A:Q57, A:V58, A:F62, A:N140, A:R141, A:S142, A:S143, A:E145, A:S146, A:G149, A:L150, A:V151, A:W152, A:S154, A:G155, A:P15 | 29 | 0.741 | NA |
| 3 | , A:I94, A:N98, A:Q99, A:T103, A:A105, A:E106, A:T107, A 09, A:A110, A:T111, A:R112, A:R113, A:D115, A:D116, A: | 17 | 0.741 | NA |
| 4 | A:S123, A:N126, A:N127, A:V130 | 4 | 0.623 | NA |
| 5 | A:L31, A:G32, A:N33, A:Q34 | 4 | 0.584 | NA |
| 6 | A:P63, A:D64, A:S65, A:D66, A:F67, A:K68 | 6 | 0.531 | NA |

Figure 23: Residues in discontinuous epitopes from Ellipro — md-0-8

**ElliPro HHP - md_0_9**

| No. | Residues | mber of residu | Score | 3D structure |
|---|---|---|---|---|
| 1 | A:V58, A:T59, A:W152, A:S154, A:G155, A:P156, A:A157 | 7 | 0.938 | NA |
| 2 | 36, A:T37, A:N91, A:I93, A:I94, A:E97, A:N98, A:Q99, A:A1 01, A:P102, A:T103, A:T104, A:A105, A:E106, A:T107, A:L A:D109, A:A110, A:T111, A:R112, A:R113 | 22 | 0.762 | NA |
| 3 | A:N25, A:T28, A:N29 | 3 | 0.671 | NA |
| 4 | A:N126, A:N127, A:V130 | 3 | 0.654 | NA |
| 5 | A:I4, A:T5, A:T6, A:P7, A:S8, A:Q9, A:F10, A:V11, A:R61, A [illegible], A:D64, A:S65, A:D66, A:F67, A:K68, A:Y139, A:N140, A:[illegible] 42, A:S143, A:E145, A:S146, A:S147, A:S148, A:G149, A:L A:V151, A:T153 | 30 | 0.64 | NA |
| 6 | A:Q38, A:R90, A:R92, A:E95, A:V96 | 5 | 0.622 | NA |
| 7 | A:A74, A:V75, A:E131, A:R134, A:G135, A:T136, A:G137, | 9 | 0.506 | NA |
| 8 | A:L13, A:P56, A:Q57, A:V60 | 4 | 0.5 | NA |

Figure 24: Residues in discontinuous epitopes from Ellipro — md-0-9

**ElliPro HHP - md_0_10**

| No. | Residues | mber of residu | Score | 3D structure |
|---|---|---|---|---|
| 1 | 91, A:I93, A:I94, A:E95, A:V96, A:E97, A:N98, A:Q99, A:A1 01, A:P102, A:T103, A:A105, A:E106, A:T107, A:L108, A:D A:A110, A:T111, A:R112, A:R113 | 21 | 0.786 | NA |
| 2 | A:I4, A:T5, A:T6, A:P7, A:S8, A:Q9, A:F10, A:V11, A:F12, A A:Q57, A:V58, A:T59, A:V60, A:R61, A:F62, A:P63, A:D64, A:F67, A:Y139, A:N140, A:R141, A:S142, A:S143, A:E145, 47, A:S148, A:G149, A:L150, A:V151, A:W152, A:T153, A:S A:G155, A:P156, A:A157 | 41 | 0.672 | NA |
| 3 | A:D19, A:I21, A:I24, A:T28, A:L132 | 5 | 0.557 | NA |
| 4 | A:G32, A:N33, A:Q34 | 3 | 0.516 | NA |
| 5 | I73, A:A74, A:V75, A:R134, A:G135, A:T136, A:G137, A:S1 | 8 | 0.504 | NA |

Figure 25: Residues in discontinuous epitopes from Ellipro — md-0-10

**ElliPro HHP - Sheet11**

| No. | Residues | mber of residu | Score | 3D structure |
|---|---|---|---|---|
| 1 | A:T37, A:Q38, A:Q39, A:R90, A:N91, A:R92, A:I93, A:I94, A:E97, A:A100, A:N101, A:P102, A:T103, A:A105, A:E106, 08, A:D109, A:A110, A:T111, A:R112, A:R113, A:D116, A:[illegible] | 27 | 0.711 | NA |
| 2 | A:I4, A:T5, A:T6, A:P7, A:S8, A:Q9, A:P56, A:Q57, A:V58, A[illegible] 60, A:R61, A:F62, A:P63, A:D64, A:S65, A:D66, A:F67, A:Y[illegible] 40, A:R141, A:S142, A:S143, A:E145, A:S146, A:S147, A:S[illegible] 49, A:L150, A:V151, A:W152, A:S154, A:G155, A:P156, A:[illegible] | 36 | 0.69 | NA |
| 3 | 24, A:T28, A:N126, A:N127, A:I129, A:V130, A:L132, A:I1[illegible] | 8 | 0.601 | NA |
| 4 | A:N73, A:A74, A:V75, A:E131, A:R134, A:G135, A:T136 | 7 | 0.533 | NA |

Figure 26: Residues in discontinuous epitopes from Ellipro — md-0-11

**EllIPro HHP-LT - md_0_1_HT**

| No. | Residues | mber of residu | Score | 3D structure |
|---|---|---|---|---|
| 1 | 89, A:N91, A:I93, A:I94, A:E95, A:E97, A:N98, A:Q99, A:A1[illegible] 01, A:P102, A:T103, A:T104, A:A105, A:E106, A:T107, A:L[illegible] A:D109, A:A110, A:T111, A:R112, A:D115 | 22 | 0.796 | NA |
| 2 | A:N25, A:T28, A:N29 | 3 | 0.729 | NA |
| 3 | A:I4, A:T6, A:P7, A:S8, A:Q9, A:F10, A:V11, A:V58, A:T59, [illegible], A:P63, A:D64, A:S65, A:D66, A:F67, A:N140, A:R141, A:[illegible] 45, A:S146, A:S147, A:S148, A:G149, A:L150, A:V151, A:W[illegible] A:T153, A:S154, A:G155, A:P156, A:A157 | 33 | 0.683 | NA |
| 4 | A:G32, A:N33, A:Q34 | 3 | 0.613 | NA |
| 5 | A:P56, A:Q57, A:V60 | 3 | 0.57 | NA |

Figure 27: Residues in discontinuous epitopes from Ellipro — md-0-1

**ElliPro HHP-LT - md_0_2_HT**

| No. | Residues | mber of residu | Score | 3D structure |
|---|---|---|---|---|
| 1 | A:V151, A:W152, A:S154, A:G155, A:P156, A:A157 | 6 | 0.83 | NA |
| 2 | 0, A:N91, A:I94, A:E95, A:E97, A:N98, A:Q99, A:A100, A:N<br>02, A:T103, A:T104, A:A105, A:E106, A:T107, A:L108, A:D<br>A:A110, A:T111, A:R112, A:R113, A:D116, A:V119, A:R12 | 24 | 0.736 | NA |
| 3 | A:I4, A:T5, A:T6, A:P7, A:S8, A:Q9, A:F10, A:V11, A:R61, A<br>3, A:D64, A:S65, A:D66, A:F67, A:N73, A:A74, A:V75, A:E<br>34, A:G135, A:T136, A:G137, A:S138, A:Y139, A:N140, A:F<br>42, A:S143, A:S146, A:S147, A:S148, A:G149, A:L150, A:T | 36 | 0.623 | NA |
| 4 | A:S55, A:P56, A:Q57, A:V60 | 4 | 0.55 | NA |
| 5 | A:N29, A:G32, A:N33, A:Q34 | 4 | 0.542 | NA |

Figure 28: Residues in discontinuous epitopes from Ellipro — md-0-2

**ElliPro HHP-LT - md_0_3_HT**

| No. | Residues | mber of residu | Score | 3D structure |
|---|---|---|---|---|
| 1 | 1, A:I93, A:I94, A:E97, A:N98, A:Q99, A:A100, A:N101, A:F<br>03, A:T104, A:A105, A:E106, A:T107, A:L108, A:D109, A:A<br>A:T111, A:R112, A:R113 | 20 | 0.817 | NA |
| 2 | 3, A:I4, A:T5, A:T6, A:P7, A:S8, A:Q9, A:F10, A:S148, A:G1<br>A:L150, A:V151, A:W152, A:S154, A:G155, A:P156, A:A15 | 17 | 0.805 | NA |
| 3 | A:P56, A:Q57, A:V58, A:T59, A:V60 | 5 | 0.701 | NA |
| 4 | A:P63, A:D64, A:S65, A:D66, A:F67, A:K68, A:N73, A:A74,<br>31, A:R134, A:G135, A:T136, A:G137, A:S138, A:N140, A:F<br>A:S142, A:E145, A:T153 | 21 | 0.572 | NA |
| 5 | A:G32, A:N33, A:Q34, A:F35 | 4 | 0.561 | NA |

Figure 29: Residues in discontinuous epitopes from Ellipro — md-0-3

**ElliPro HHP-LT - md_0_4_HT**

| ontinuc | ...2 | ...3 | ...4 | ...5 |
|---|---|---|---|---|
| No. | Residues | umber of residue | Score | 3D structure |
| 1 | A:A105, A:E106, A:T107 | 3 | 0.935 | NA |
| 2 | A:L13, A:P56, A:Q57, A:V58, A:T59, A:G149, A:L150, A:V | 21 | 0.756 | NA |
| 3 | A:N98, A:Q99, A:A100, A:N101, A:P102, A:T103, A:T104, A | 22 | 0.745 | NA |
| 4 | A:S65, A:D66, A:F67, A:Y139, A:N140, A:R141, A:S142, A:S | 15 | 0.628 | NA |
| 5 | A:D19, A:I21, A:E22, A:N25 | 4 | 0.561 | NA |

Figure 30: Residues in discontinuous epitopes from Ellipro — md-0-4

**ElliPro HHP-LT - MD_0_5_HT**

| No. | Residues | mber of residu | Score | 3D structure |
|---|---|---|---|---|
| 1 | 1, A:R92, A:I93, A:I94, A:E95, A:V96, A:N98, A:A100, A:N 02, A:T103, A:T104, A:A105, A:E106, A:T107, A:L108, A:D A:A110, A:R112, A:R113 | 20 | 0.833 | NA |
| 2 | A:I4, A:T5, A:T6, A:P7, A:S8, A:Q9, A:F10, A:L13, A:P56, A :T59, A:V60, A:R61, A:F62, A:P63, A:D64, A:S65, A:D66, 1, A:S142, A:G149, A:L150, A:V151, A:W152, A:T153, A: A:G155, A:P156, A:A157 | 32 | 0.715 | NA |
| 3 | :A74, A:V75, A:V130, A:E131, A:R134, A:G135, A:T136, A:S138 | 10 | 0.529 | NA |

Figure 31: Residues in discontinuous epitopes from Ellipro — md-0-5

**ElliPro HHP-LT - MD_0_6_HT**

| No. | Residues | mber of residu | Score | 3D structure |
|---|---|---|---|---|
| 1 | A:V58, A:T59, A:S154, A:G155, A:P156, A:A157 | 6 | 0.94 | NA |
| 2 | 36, A:T37, A:N91, A:I93, A:I94, A:E95, A:N98, A:Q99, A:A1 01, A:P102, A:T103, A:T104, A:A105, A:E106, A:T107, A:L A:D109, A:A110, A:R112, A:R113 | 21 | 0.778 | NA |
| 3 | A:I4, A:T5, A:T6, A:P7, A:S8, A:Q9, A:F10, A:V11, A:R61, , A:D64, A:S65, A:D66, A:F67, A:K68, A:Y139, A:N140, A: 42, A:S143, A:E145, A:S146, A:S147, A:S148, A:G149, A:L A:V151, A:W152, A:T153 | 31 | 0.645 | NA |
| 4 | A:Q39, A:T42, A:V43 | 3 | 0.611 | NA |
| 5 | 73, A:A74, A:V75, A:V130, A:E131, A:R134, A:G135, A:T1 | 8 | 0.573 | NA |
| 6 | A:P56, A:Q57, A:V60 | 3 | 0.561 | NA |

Figure 32: Residues in discontinuous epitopes from Ellipro — md-0-6

**ElliPro HHP-LT - MD_0_7_HT**

| No. | Residues | mber of residu | Score | 3D structure |
|---|---|---|---|---|
| 1 | A:T37, A:R90, A:N91, A:R92, A:I93, A:I94, A:E95, A:V96, :Q99, A:A100, A:N101, A:P102, A:T103, A:A105, A:E106, A:L108, A:D109, A:A110, A:T111, A:R112, A:R113 | 25 | 0.77 | NA |
| 2 | A:I4, A:T5, A:T6, A:P7, A:S8, A:Q9, A:F10, A:L13, A:P56, A A:T59, A:V60, A:R61, A:F62, A:P63, A:D64, A:S65, A:D66, R141, A:G149, A:L150, A:V151, A:W152, A:T153, A:S154, A:P156, A:A157 | 32 | 0.715 | NA |

Figure 33: Residues in discontinuous epitopes from Ellipro — md-0-7

**ElliPro HHP-LT - MD_0_8_HT**

| No. | Residues | mber of residu | Score | 3D structure |
|---|---|---|---|---|
| 1 | A:S154, A:G155, A:P156, A:A157 | 4 | 0.911 | NA |
| 2 | A:E97, A:N98, A:Q99, A:A100, A:N101, A:P102, A:T103, A<br>A:E106, A:T107, A:L108, A:D109, A:A110, A:T111, A:R112 | 18 | 0.812 | NA |
| 3 | A:S3, A:I4, A:T5, A:T6, A:P7, A:S8, A:Q9, A:F10, A:V11 | 9 | 0.801 | NA |
| 4 | A:P56, A:Q57, A:V60 | 3 | 0.637 | NA |
| 5 | A:Q38, A:Q39, A:T42, A:R90, A:N91, A:R92, A:E95 | 7 | 0.615 | NA |
| 6 | 1, A:F62, A:P63, A:D64, A:S65, A:D66, A:F67, A:K68, A:S<br>A:R141, A:S142, A:S146, A:G149, A:L150, A:V151, A:W15 | 18 | 0.58 | NA |
| 7 | A:N73, A:A74, A:V75, A:R134, A:G135, A:T136, A:G137 | 7 | 0.501 | NA |

Figure 34: Residues in discontinuous epitopes from Ellipro — md-0-8

**ElliPro HHP-LT - MD_0_9_HT**

| No. | Residues | mber of residu | Score | 3D structure |
|---|---|---|---|---|
| 1 | , A:T37, A:E95, A:E97, A:N98, A:Q99, A:A100, A:N101, A:<br>03, A:T104, A:A105, A:E106, A:T107, A:L108, A:D109, A:A<br>A:T111, A:R112, A:R113 | 20 | 0.76 | NA |
| 2 | A:R90, A:N91, A:R92, A:I93, A:I94 | 5 | 0.735 | NA |
| 3 | A:I4, A:T5, A:T6, A:P7, A:S8, A:Q9, A:P56, A:Q57, A:V58, A<br>0, A:R61, A:F62, A:P63, A:D64, A:S65, A:D66, A:F67, A:Y<br>40, A:R141, A:S142, A:S143, A:E145, A:S146, A:S147, A:S<br>A:L150, A:V151, A:W152, A:T153, A:S154, A:G155, A:P15 | 37 | 0.695 | NA |
| 4 | A:N126, A:N127, A:V130 | 3 | 0.602 | NA |
| 5 | A:A74, A:V75, A:E131, A:R134, A:G135, A:T136, A:G137, | 9 | 0.513 | NA |

Figure 35: Residues in discontinuous epitopes from Ellipro — md-0-9

**ElliPro HHP-LT - MD_0_10_HT**

| No. | Residues | mber of residu | Score | 3D structure |
|---|---|---|---|---|
| 1 | A:S3, A:I4, A:P7, A:S8, A:Q9, A:F10 | 6 | 0.825 | NA |
| 2 | A:V96, A:E97, A:N98, A:Q99, A:A100, A:N101, A:P102, A<br>05, A:E106, A:T107, A:L108, A:D109, A:A110, A:R112, A:I | 17 | 0.822 | NA |
| 3 | A:R90, A:N91, A:R92, A:I93, A:I94 | 5 | 0.726 | NA |
| 4 | A:Q57, A:V58, A:T59, A:V60, A:R61, A:F62, A:P63, A:D64,<br>A:F67, A:N140, A:R141, A:S142, A:E145, A:S146, A:S147,<br>A:L150, A:V151, A:W152, A:T153, A:S154, A:G155, A:P15 | 28 | 0.685 | NA |
| 5 | A:G32, A:N33, A:Q34 | 3 | 0.533 | NA |
| 6 | A:A74, A:V75, A:L76, A:N127, A:V130, A:E131, A:R134, A<br>A:T136, A:G137, A:S138 | 12 | 0.521 | NA |

Figure 36: Residues in discontinuous epitopes from Ellipro — md-0-10

**ElliPro HHP-LT - MD_0_11_HT**

| No. | Residues | mber of residu | Score | 3D structure |
|---|---|---|---|---|
| 1 | A:N91, A:R92, A:I93, A:I94, A:E95, A:V96, A:E97, A:N98, A<br>00, A:N101, A:P102, A:T103, A:A105, A:E106, A:T107, A:L<br>A:D109, A:A110, A:R112, A:R113 | 22 | 0.811 | NA |
| 2 | A:I4, A:T5, A:T6, A:P7, A:S8, A:Q9, A:F10, A:V11, A:L13, A<br>A:V58, A:T59, A:V60, A:R61, A:F62, A:P63, A:D64, A:S65,<br>41, A:S142, A:E145, A:S147, A:S148, A:G149, A:L150, A:V<br>A:W152, A:T153, A:S154, A:G155, A:P156, A:A157 | 35 | 0.716 | NA |
| 3 | I73, A:A74, A:V75, A:E131, A:R134, A:G135, A:T136, A:G1 | 8 | 0.562 | NA |
| 4 | A:D115, A:D116, A:A117, A:V119 | 4 | 0.544 | NA |

Figure 37: Residues in discontinuous epitopes from Ellipro — md-0-11

**ElliPro HHP-LT - md_0_12_HT**

| No. | Residues | mber of residu | Score | 3D structure |
|---|---|---|---|---|
| 1 | A:S3, A:I4, A:T5, A:T6, A:P7, A:S8, A:Q9, A:F10, A:L13 | 9 | 0.814 | NA |
| 2 | A:E97, A:N98, A:Q99, A:A100, A:N101, A:P102, A:T103, A<br>05, A:E106, A:T107, A:L108, A:D109, A:A110, A:T111, A:I | 17 | 0.784 | NA |
| 3 | A:N91, A:R92, A:I93, A:I94, A:E95, A:R113 | 6 | 0.748 | NA |
| 4 | A:Q57, A:V58, A:T59, A:V60, A:R61, A:F62, A:P63, A:D64,<br>A:F67, A:N140, A:R141, A:S142, A:E145, A:S146, A:S147,<br>A:L150, A:V151, A:W152, A:T153, A:S154, A:G155, A:P15 | 28 | 0.689 | NA |
| 5 | A:A74, A:V75, A:E131, A:R134, A:G135, A:T136, A:G137, | 9 | 0.52 | NA |

Figure 38: Residues in discontinuous epitopes from Ellipro — md-0-12

**ElliPro HHP-LT - md_0_13_HT**

| No. | Residues | mber of residu | Score | 3D structure |
|---|---|---|---|---|
| 1 | A:N91, A:R92, A:I93, A:I94, A:E95, A:V96, A:E97, A:N98, ,<br>00, A:N101, A:P102, A:T103, A:T104, A:A105, A:E106, A:T<br>A:L108, A:D109, A:A110, A:T111, A:R112, A:R113 | 24 | 0.776 | NA |
| 2 | A:I4, A:T5, A:T6, A:P7, A:S8, A:Q9, A:P56, A:Q57, A:V60, A<br>, A:P63, A:D64, A:S65, A:D66, A:R141, A:S142, A:E145, A:<br>47, A:S148, A:G149, A:L150, A:V151, A:W152, A:T153, A:S<br>A:G155, A:P156, A:A157 | 31 | 0.728 | NA |
| 3 | A:N126, A:N127, A:V130 | 3 | 0.643 | NA |
| 4 | A:D116, A:A117, A:V119 | 3 | 0.617 | NA |
| 5 | A:D88, A:T89, A:R90 | 3 | 0.576 | NA |
| 6 | I73, A:A74, A:V75, A:E131, A:R134, A:G135, A:T136, A:G1 | 8 | 0.528 | NA |

Figure 39: Residues in discontinuous epitopes from Ellipro — md-0-13

**ElliPro HHP-LT - md_0_14_HT**

| No. | Residues | mber of residu | Score | 3D structure |
|---|---|---|---|---|
| 1 | A:T37, A:N91, A:R92, A:I93, A:I94, A:E95, A:V96, A:E97, ,<br>:A100, A:N101, A:P102, A:T103, A:T104, A:A105, A:E106,<br>A:L108, A:D109, A:A110, A:R112, A:R113 | 24 | 0.782 | NA |
| 2 | A:I4, A:T5, A:T6, A:P7, A:S8, A:Q9, A:F10, A:V11, A:P56, A<br>A:T59, A:V60, A:R61, A:F62, A:P63, A:D64, A:S65, A:D66, ,<br>41, A:S142, A:E145, A:G149, A:L150, A:V151, A:W152, A:T<br>A:S154, A:G155, A:P156, A:A157 | 33 | 0.749 | NA |
| 3 | A:A74, A:V75, A:N126, A:V130, A:E131, A:I133, A:R134, , | 9 | 0.609 | NA |

Figure 40: Residues in discontinuous epitopes from Ellipro — md-0-14

**ElliPro HHP-LT - md_0_15_HT**

| No. | Residues | mber of residu | Score | 3D structure |
|---|---|---|---|---|
| 1 | A:I4, A:T5, A:T6, A:P7, A:S8, A:Q9, A:F10, A:V11, A:P56, A<br>58, A:V60, A:R61, A:F62, A:P63, A:D64, A:S65, A:D66, A:R<br>42, A:E145, A:S148, A:G149, A:L150, A:V151, A:W152, A:T<br>A:S154, A:G155, A:P156, A:A157 | 32 | 0.739 | NA |
| 2 | A:Q36, A:T37, A:T89, A:N91, A:R92, A:I93, A:I94, A:E95,<br>A:N98, A:Q99, A:A100, A:N101, A:P102, A:T103, A:T104, A<br>06, A:T107, A:L108, A:D109, A:A110, A:T111, A:R112, A:R | 27 | 0.73 | NA |
| 3 | A:A74, A:V75, A:L76, A:N126, A:N127, A:V130, A:E131, A<br>A:G135, A:T136 | 11 | 0.535 | NA |

Figure 41: Residues in discontinuous epitopes from Ellipro — md-0-15

**ElliPro HHP-LT - md_0_16_HT**

| No. | Residues | mber of residu | Score | 3D structure |
|---|---|---|---|---|
| 1 | A:Q36, A:T37, A:Q39, A:N91, A:R92, A:I93, A:I94, A:E95,<br>A:N98, A:Q99, A:A100, A:N101, A:P102, A:T103, A:T104, A<br>06, A:T107, A:L108, A:D109, A:A110, A:T111, A:R112, A:R | 27 | 0.737 | NA |
| 2 | A:I4, A:T5, A:T6, A:P7, A:S8, A:Q9, A:V11, A:S14, A:P56, A<br>A:T59, A:V60, A:R61, A:F62, A:P63, A:D64, A:S65, A:D66,<br>40, A:R141, A:S142, A:E145, A:S146, A:S147, A:S148, A:G<br>50, A:V151, A:W152, A:T153, A:S154, A:G155, A:P156, A:A | 37 | 0.709 | NA |
| 3 | A:N126, A:N127, A:V130 | 3 | 0.589 | NA |

Figure 42: Residues in discontinuous epitopes from Ellipro — md-0-16

**ElliPro HHP-LT - md_0_17_HT**

| No. | Residues | mber of residu | Score | 3D structure |
|---|---|---|---|---|
| 1 | A:Q36, A:T37, A:T89, A:N91, A:R92, A:I93, A:I94, A:E95,<br>A:N98, A:Q99, A:A100, A:N101, A:P102, A:T103, A:T104, A<br>06, A:T107, A:L108, A:D109, A:A110, A:T111, A:R112, A:R | 27 | 0.738 | NA |
| 2 | A:I4, A:T5, A:T6, A:P7, A:S8, A:Q9, A:R61, A:F62, A:P63, A<br>A:D66, A:F67, A:N140, A:R141, A:S142, A:E145, A:S146, A<br>A:L150, A:V151, A:W152, A:T153, A:S154, A:G155, A:P15 | 29 | 0.736 | NA |
| 3 | A:S55, A:P56, A:Q57, A:V58, A:T59, A:V60 | 6 | 0.602 | NA |
| 4 | A:A74, A:V75, A:N126, A:N127, A:V130, A:E131, A:R134, A<br>A:T136 | 10 | 0.601 | NA |

Figure 43: Residues in discontinuous epitopes from Ellipro — md-0-17

**ElliPro HHP-LT - md_0_18_HT**

| No. | Residues | mber of residu | Score | 3D structure |
|---|---|---|---|---|
| 1 | A:T89, A:N91, A:R92, A:I93, A:I94, A:E95, A:V96, A:E97, A<br>:A100, A:N101, A:P102, A:T103, A:T104, A:T107, A:L108,<br>A:A110, A:T111, A:R112, A:R113, A:D116, A:A117, A:V11 | 26 | 0.742 | NA |
| 2 | A:P56, A:Q57, A:V58, A:T59, A:V60 | 5 | 0.738 | NA |
| 3 | A:I4, A:T5, A:T6, A:P7, A:S8, A:Q9, A:F10, A:R61, A:F62, A<br>A:S65, A:D66, A:F67, A:N140, A:R141, A:S142, A:E145, A<br>47, A:S148, A:G149, A:L150, A:V151, A:W152, A:T153, A:S<br>A:G155, A:P156, A:A157 | 31 | 0.726 | NA |
| 4 | A:A74, A:V75, A:N126, A:N127, A:V130, A:E131, A:R134, A<br>A:T136 | 10 | 0.576 | NA |

Figure 44: Residues in discontinuous epitopes from Ellipro — md-0-18

**ElliPro HHP-LT - md_0_19_HT**

| No. | Residues | mber of residu | Score | 3D structure |
|---|---|---|---|---|
| 1 | A:Q36, A:T37, A:T89, A:N91, A:R92, A:I93, A:I94, A:E95,<br>A:N98, A:Q99, A:A100, A:N101, A:P102, A:T103, A:T104, A<br>06, A:T107, A:L108, A:D109, A:A110, A:T111, A:R112, A:R<br>A:D116, A:A117, A:V119 | 30 | 0.717 | NA |
| 2 | A:I4, A:T5, A:T6, A:P7, A:S8, A:Q9, A:F10, A:V11, A:S14, A<br>A:V58, A:T59, A:V60, A:R61, A:F62, A:P63, A:D64, A:S65,<br>A:Y72, A:N140, A:R141, A:S142, A:E145, A:S146, A:S148,<br>50, A:V151, A:W152, A:T153, A:S154, A:G155, A:P156, A: | 38 | 0.704 | NA |
| 3 | A:N73, A:A74, A:V75, A:E131, A:I133, A:R134, A:T136 | 7 | 0.617 | NA |
| 4 | A:N126, A:N127, A:V130 | 3 | 0.6 | NA |
| 5 | A:G85, A:D88, A:R90 | 3 | 0.576 | NA |

Figure 45: Residues in discontinuous epitopes from Ellipro — md-0-19

**ElliPro HHP-LT - md_0_20_HT**

| No. | Residues | mber of residu | Score | 3D structure |
|---|---|---|---|---|
| 1 | A:I4, A:T5, A:T6, A:P7, A:S8, A:Q9, A:F10, A:V11, A:V60, A<br>2, A:P63, A:D64, A:S65, A:D66, A:F67, A:Y72, A:S138, A:N<br>41, A:S142, A:E145, A:S146, A:S148, A:G149, A:L150, A:V<br>A:W152, A:T153, A:S154, A:G155, A:P156, A:A157 | 34 | 0.71 | NA |
| 2 | A:P56, A:Q57, A:V58, A:T59 | 4 | 0.705 | NA |
| 3 | A:N33, A:Q34, A:Q36, A:T37, A:Q39, A:T89, A:N91, A:R92<br>, A:E95, A:V96, A:E97, A:N98, A:Q99, A:A100, A:N101, A:[illegible]<br>03, A:T104, A:A105, A:E106, A:T107, A:L108, A:D109, A:A<br>A:T111, A:R112, A:R113, A:D116, A:A117, A:V119 | 33 | 0.694 | NA |
| 4 | A:G85, A:D88, A:R90 | 3 | 0.57 | NA |
| 5 | A:A74, A:V75, A:S123, A:N126, A:V130, A:E131, A:R134, A<br>A:T136 | 10 | 0.546 | NA |

Figure 46: Residues in discontinuous epitopes from Ellipro — md-0-20

**ElliPro HHP-LT - md_0_21_HT**

| No. | Residues | mber of residu | Score | 3D structure |
|---|---|---|---|---|
| 1 | A:T37, A:N91, A:R92, A:I93, A:I94, A:E95, A:V96, A:E97,<br>A:A100, A:N101, A:P102, A:T103, A:T104, A:A105, A:E106,<br>A:L108, A:D109, A:A110, A:T111, A:R112, A:R113 | 25 | 0.754 | NA |
| 2 | A:I4, A:T5, A:P7, A:S8, A:Q9, A:F10, A:V11, A:P56, A:Q57,<br>A:V60, A:R61, A:F62, A:P63, A:D64, A:S65, A:D66, A:F67,<br>41, A:S142, A:E145, A:S146, A:S147, A:S148, A:G149, A:L<br>A:V151, A:W152, A:T153, A:S154, A:G155, A:P156, A:A15[illegible] | 36 | 0.721 | NA |
| 3 | A:D116, A:A117, A:V119 | 3 | 0.628 | NA |
| 4 | A:N73, A:A74, A:V75, A:E131, A:R134, A:G135, A:T136 | 7 | 0.523 | NA |

Figure 47: Residues in discontinuous epitopes from Ellipro — md-0-21
