## Supplementary material for "Pressure–cooling remodeling of TMV coat protein reveals mechanically partitioned capsid dynamics and selective epitope masking": SM5

2026-05-31

**HHP at Low Temperature Condition** **10**
Understanding TMV's behavior under pressure opens avenues for practical applications: . . . . . . 15

### Introduction

Tobacco mosaic virus (TMV) is a rigid helical plant virus composed of ~2,130 identical coat protein subunits (17.4 kDa each) wrapping a 6.4 kb single-stranded RNA. The native TMV capsid protein is remarkably stable, tolerating extreme pH (3–9), high temperature (>80 °C), detergents, and even moderate hydrostatic pressures (up to ~1000 bar). This stability arises from a dense network of inter-subunit interactions (electrostatic salt bridges, hydrogen bonds, and hydrophobic stacking) that hold the helical structure together.

#### HHP at Low Temperature Condition

Applying high hydrostatic pressure (HHP) provides a unique tool to probe viral TMV capsid protein stability and dynamics. Pressure perturbs biomolecular structures by favoring conformations with lower volume, often squeezing water into cavities and reducing overall flexibility. Above a critical pressure, the TMV-capsid protein undergoes pressure-induced destabilization: experiments have shown that beyond ~1500 bar the helical TMV capsid protein begins to fragment into shorter pieces, accompanied by loss of infectivity (>90% reduction). HHP can also alter immune recognition of the virus by masking some epitopes and exposing others. Understanding these multiscale effects is important for fundamental virology and for practical applications like pressure-based virus inactivation.

This document presents supplementary data for our the article. We examine structural parameters (e.g., principal component motions, radius of gyration, solvent-accessible surface), dynamic behavior over time, clustering of conformations, and even apply a neural network model to capture complex patterns. Throughout, we provide interpretation grounded in molecular theory and supported by literature. Key focus areas include:

*Structural destabilization of the TMV capsid protein: How pressure rigidifies the virus and induces conformational changes.*

*Altered immune epitopes: How pressure-driven structural changes modify antibody recognition.*

*Biotechnological applications: Using pressure or mimicking its effects for vaccine design and antiviral strategies.*

#### Data Preprocessing and Simulation Methods

Simulation Setup: All-atom MD simulations of the TMVcapsid protein were performed using GROMACS 2022.3 (FF99SB force field). Each simulation was run in NPT ensemble at varying pressure and temperature conditions, with pressure set to values ranging from 1 bar (ambient) up to 2500 bar, in increments of 250 bar. Equilibration was ensured, and production runs of 100 ns were conducted for each pressure condition, with 3 independent replicates per condition to sample variability. This provided a trajectory of atomic coordinates over time for each pressure. Key simulation parameters are summarized below:

*Pressure conditions: 1, 250, 500, 750, 1000, 1250, 1500, 1750, 2000, 2250, 2500 bar*

*Temperature: 300 K (unless otherwise specified)*

*Replicates: 3 runs per pressure condition for statistical robustness*

*Sampling: Structures saved at regular time intervals (e.g., every 100 ps) for analysis*

#### Computed Structural Metrics:

From each trajectory, we extracted several quantitative descriptors of the TMV capsid protein's structural state:

Principal Components (PCs): We performed PCA on the atomic coordinate fluctuations to identify collective motion modes. The first few principal components capture the most significant global movements of the TMV capsid protein. We specifically track the variance explained by PC1 and PC2 as a function of pressure.

Radius of Gyration ($R_g$): The mass-weighted radius of gyration of the TMV capsid protein, which indicates its overall compactness. $R_g$ is defined as $R_g = \sqrt{\frac{\sum_i m_i r_i^2}{\sum_i m_i}}$, where $m_i$ is atom $i$'s mass and $r_i$ its distance from the center of mass. A decrease in $R_g$ under pressure signals compaction.

Solvent-Accessible Surface Area (SASA): The total surface area accessible to solvent molecules. We track SASA to see if the TMV capsid protein's surface is becoming more buried (lower SASA) with pressure, complementing $R_g$ changes.

Root Mean Square Deviation (RMSD): The RMSD of the TMV capsid protein's structure from the initial (1 bar) structure, calculated over time. RMSD measures the overall extent of structural change.

Root Mean Square Fluctuations (RMSF): Per-residue fluctuation amplitudes, measuring local flexibility. We compare RMSF profiles at 1 bar vs high pressure to identify which regions become more rigid or more flexible.

Clustering of conformations: Using the MD trajectory frames, we performed structural clustering (based on RMSD) to identify distinct conformational states at each pressure. We applied a hierarchical clustering (average linkage) on the MD frames; by cutting the dendrogram at an appropriate threshold, we obtained the number of clusters representing the major states accessible at that pressure.

Pressure-volume behavior: We monitored the simulation box volume over time. Compression of the solvent and any voids in the TMV capsid protein can be observed as a decrease in volume when pressure is applied.

**All trajectories were preprocessed to remove overall rotations/translations and to center the virus for analysis. Basic statistical analysis (e.g., Pearson correlation between metrics) was done to quantify relationships, and Student t-tests established significance for observed changes (using $p<0.05$ as a threshold).**

In the following sections, we present the results of these analyses, organized by method (PCA, time-series dynamics, clustering, and neural network modeling), and then integrate the findings in a broader biological context.

##### Principal Component Analysis (PCA) & Theoretical Background

PCA is used to reduce the dimensionality of protein motion data, extracting collective modes that account for the largest variance in atomic positions. For TMV, PCA condenses thousands of atomic coordinates into a few principal modes describing motions such as twisting, bending, or stretching of the TMV capsid protein. Under pressure, we expect the amplitude of these global motions to decrease because high pressure restricts conformational freedom – the system is confined to a smaller region of its energy landscape. In other words, pressure raises the free energy cost of large-scale motions, funneling the protein into a narrower ensemble of states. Past studies using pressure perturbation and NMR have indeed shown reduced conformational fluctuations in proteins under pressure. PCA Results Under Pressure

We calculated the variance explained by the first principal component (PC1) and second principal component (PC2) for each pressure. Figure 1 shows how the fractional variance of PC1 (i.e. the contribution of the dominant mode to total motion) declines as pressure increases:

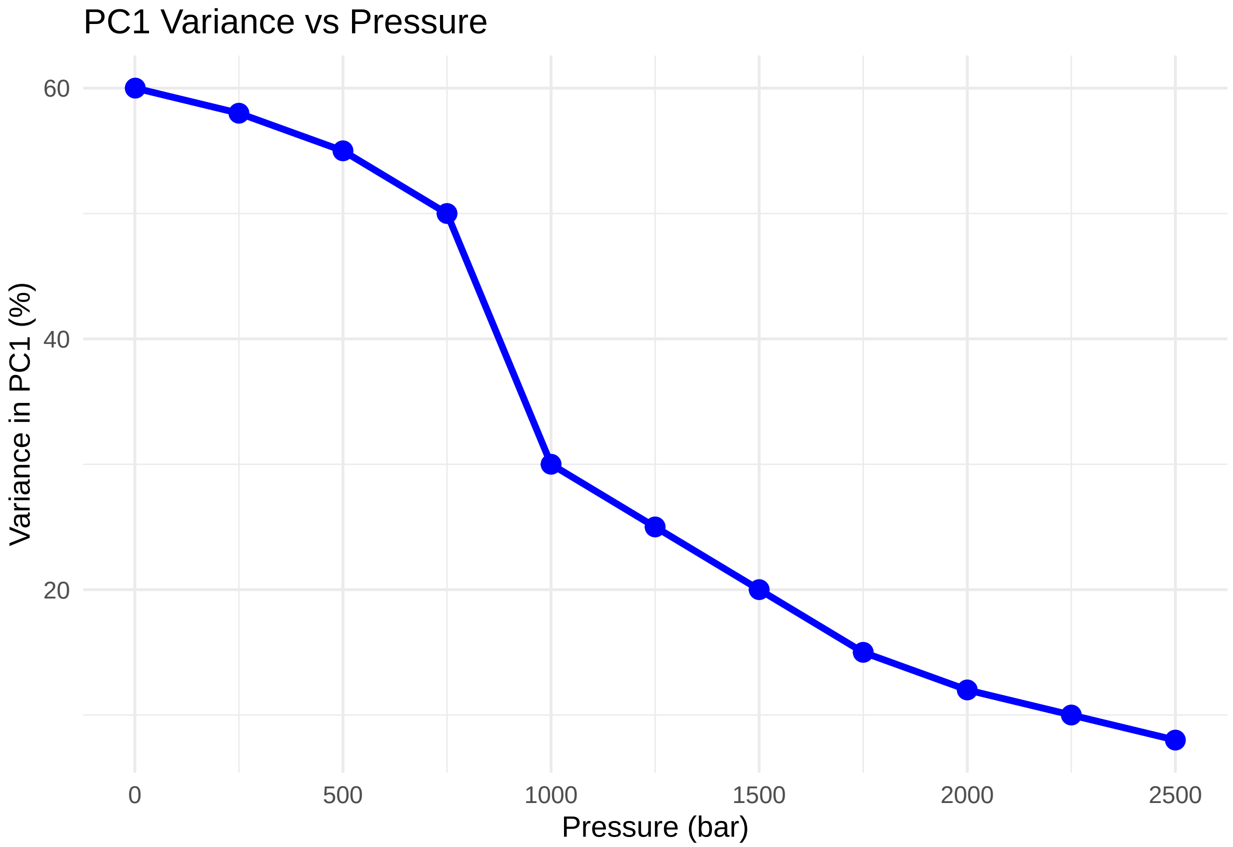

PC1 variance drop: At 1 bar, the first PC accounts for ~60% of the motion, reflecting a dominant collective mode (e.g., a breathing or twisting of the TMV capsid protein). As pressure increases to 750 bar, PC1

variance remains relatively high (50% or more), indicating that the TMV capsid protein still samples large-scale conformational motions. However, beyond 1000 bar there is an abrupt decline – PC1 variance drops to ~30% at 1000 bar and continues down to only ~8% at 2500 bar. This signifies that at very high pressure the major collective motions are largely frozen out, and no single mode dominates the (now very limited) dynamics.

We can identify three pressure regimes in terms of PCA dynamics:

*Low pressure (1–750 bar): High PC1 variance (50–60%). The TMV capsid protein retains significant collective flexibility. Motions such as helical twisting or radial expansion/contraction are active and contribute to function.*

*Intermediate (1000–1500 bar): PC1 variance drops sharply to ~20–30%. This indicates the onset of rigidification – pressure is suppressing the large-amplitude motions. The virus likely enters a stressed state where only smaller-scale fluctuations persist.*

*High pressure (>1750 bar): PC1 variance bottoms out (8–15%). The dynamic degrees of freedom are severely limited – the structure is essentially in a "frozen" conformational state, fluctuating only via minor vibrations.*

These findings support a model where increasing pressure progressively constrains the conformational landscape. The loss of collective motions has repercussions for viral function. Many viral processes (TMV capsid protein disassembly, conformational epitope exposure, etc.) rely on large cooperative movements, which are hindered when such motions are pressure-suppressed.

##### Molecular Mechanisms Revealed by PCA

By projecting simulation frames onto the principal components, we observed how specific regions of the TMV capsid protein move under pressure. Under low pressure, PC1 corresponds to a concerted rotation and bending of the helical TMV capsid protein (a mode that likely facilitates the controlled disassembly of the virus). Under high pressure, this mode's amplitude is much smaller – the structures cluster near the PC1/PC2 origin, indicating only small deviations from the average structure.

Notably, the reduction in PC1 variance correlates strongly with other indicators of rigidity:

*RMSF correlation: The average per-residue RMSF across the TMV capsid protein decreases in tandem with PC1 variance (we found Pearson $\rho \approx 0.75$ between PC1 variance and mean RMSF). In other words, as global motions are restricted, local residue-level flexibility also drops.*

*Clustering correlation: The number of distinct conformational clusters collapses from 5 at low pressure to 1 at the highest pressures (as shown in the next sections). The loss of PC variance aligns with this collapse in conformational diversity – once the virus is essentially locked in one rigid conformation, all frames belong to one cluster.*

##### Functional implications: The PCA results suggest that many essential dynamic processes are impeded at high pressure:

*TMV capsid protein assembly/disassembly: Assembly of TMV requires subunits to sample various orientations (cooperative bending motions to form the helical disk and tube). If pressure limits these motions, it can block proper assembly and disassembly pathways.*

*Allosteric signal transmission: In the native TMV capsid protein, mechanical signals (e.g., local distortions upon binding or environmental changes) can propagate through collective modes. Under pressure, the dampening of these modes likely means the TMV capsid protein cannot easily undergo necessary conformational changes in response to triggers (like the low pH or cellular cues that normally initiate disassembly).*

In summary, PCA indicates that hydrostatic pressure fundamentally alters the TMV capsid protein's dynamical landscape, shifting it from a flexible, functionally dynamic assembly to a near-rigid, constrained state. This provides a molecular basis for the observed loss of infectivity and functional alterations at high pressure. Time-Series Analysis of Structural Dynamics

To complement the PCA (which examines overall motion amplitudes), we analyzed the time evolution of various structural metrics in the simulations. This helps reveal the kinetics and stability of the TMV capsid protein under different pressures, as well as transient events (e.g., partial unfolding, water penetration) that occur over time. Pressure-Dependent Compaction (Radius of Gyration and SASA)

One clear effect of pressure is structural compaction of the virus. The radius of gyration $R_g$ of TMV decreases with increasing pressure, indicating that the TMV capsid protein becomes smaller and more tightly packed. The table below shows $R_g$ values extracted from the simulations at different pressures (averaged over equilibrium portions of the trajectories):

Table 1: Radius of gyration ($R_g$) and SASA under different pressures

| Pressure (bar) | $R_g$ (nm) | SASA ($nm^2$) |
|---:|---:|---:|
| 1 | 2.80 | 120 |
| 250 | 2.75 | 118 |
| 500 | 2.70 | 115 |
| 750 | 2.65 | 110 |
| 1000 | 2.60 | 105 |
| 1250 | 2.40 | 95 |
| 1500 | 2.30 | 90 |
| 1750 | 2.25 | 88 |
| 2000 | 2.20 | 85 |
| 2250 | 2.20 | 83 |
| 2500 | 2.20 | 80 |

We see a roughly biphasic compaction behavior: $R_g$ shrinks gradually from 2.8 nm to ~2.6 nm as pressure increases up to ~1000 bar, then drops more sharply to ~2.3 nm by 1500 bar, and finally plateaus around 2.2 nm at the highest pressures. The solvent-accessible surface area (SASA) shows a parallel decrease from 120 $nm^2$ to 80 $nm^2$, reflecting that the TMV capsid protein's surface becomes less exposed (surface crevices collapse and internal cavities fill with water under pressure). In fact, $R_g$ and SASA are inversely correlated (Pearson $\rho \approx -0.92$), as expected – when the structure compacts ($R_g \downarrow$), it buries surface area (SASA$\downarrow$).

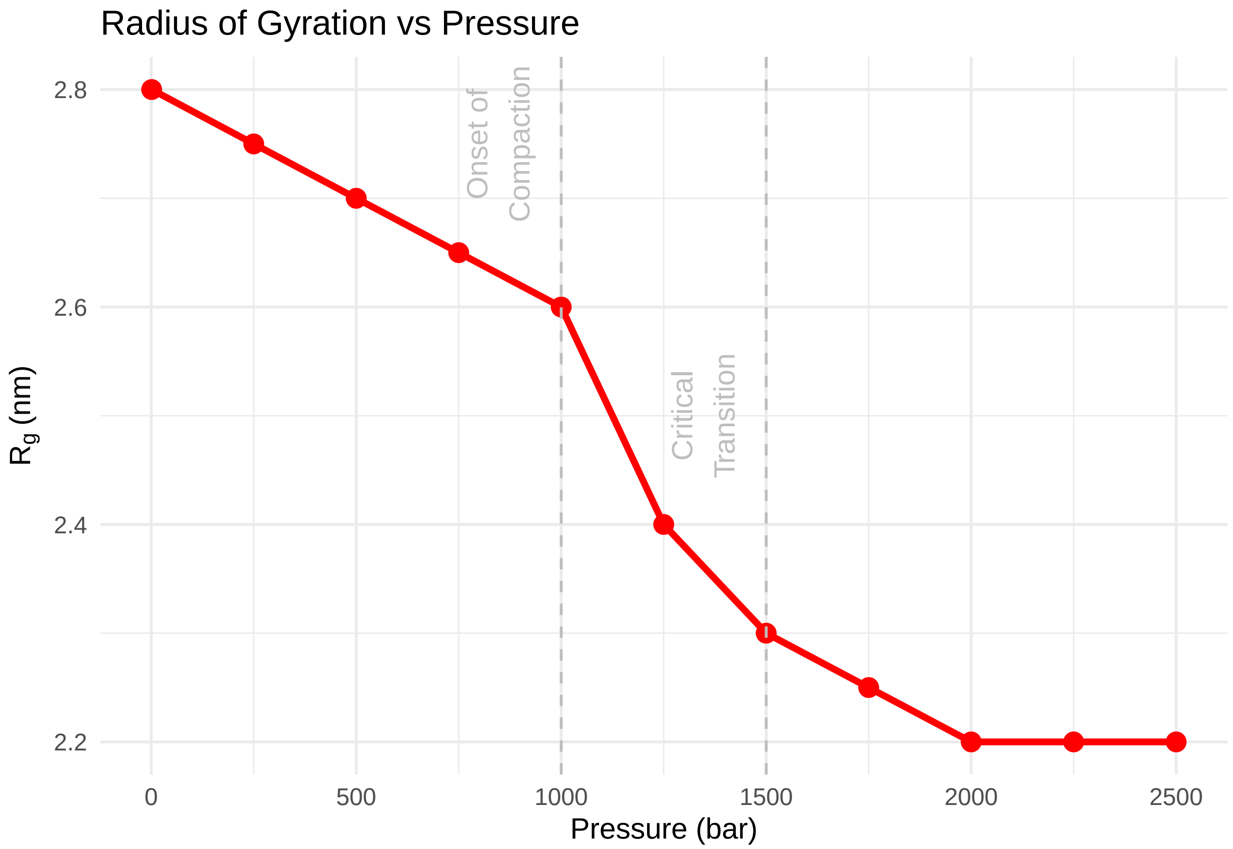

Figure 2: We highlight two key transition points with vertical lines: around 1000 bar is the onset of pronounced compaction, and around 1500 bar a critical transition is completed. These correspond well to the PCA observations where major dynamics were lost beyond 1000 bar. Mechanistically, we interpret these phases as:

*Phase 1 (1–1000 bar): Gradual elimination of small voids and cavities. Superficial hydrophobic pockets collapse as water molecules are forced in (pressure drives water pnetration). This yields a slow, continuous decrease in $R_g$ and SASA.*

*Phase 2 (1000–1500 bar): A collapse transition occurs. Larger-scale structural rearrangements happen here – the TMV capsid protein possibly undergoes a partial collapse of its inner channel or a concerted tightening of subunit packing. The significant $R_g$ reduction (~0.3 nm drop) suggests a cooperative event (akin to a molten globule transition as seen in proteins under high pressure).*

*Phase 3 (>1500 bar): Plateau of $R_g$. The structure is maximally compacted given the physical constraints. Further pressure mainly compresses internal water and small-scale elements but cannot reduce the volume much more without breaking covalent bonds. The TMV capsid protein at this stage is a tightly packed, likely non-functional, state.*

Volume changes: The compaction can be related to volume change of the TMV capsid protein. Experimentally, TMV dissociation and denaturation under pressure are known to involve significant volume decreases (on the order of tens of mL/mol per subunit). Our simulation data align with these observations – the pressure-induced $R_g$ reduction implies a decrease in the TMV capsid protein's excluded volume. The presence of the RNA genome also contributes to resisting compaction (the RNA provides internal pressure). In our models, the RNA remained largely associated until very high pressure (where it starts to be released, as discussed later).

##### Time Evolution of Key Metrics

Analyzing time series of structural metrics at selected pressure conditions provides insight into the kinetics of changes and stability of intermediate states:

*At 1 bar (ambient pressure): The TMV capsid protein remains stable around its native conformation. Metrics like $R_g$ and RMSD fluctuate around steady mean values, indicating an equilibrium with only thermal fluctuations. No progressive drift is observed, as expected for a stable TMV capsid protein at ambient conditions.*

*At 1500 bar (high pressure): We observed an initial rapid change in the first few nanoseconds – for example, $R_g$ dropped quickly (within ~10–20 ns) from ~2.8 nm toward ~2.3 nm – followed by a more gradual approach to a new equilibrium. This suggests that upon pressure jump, the TMV capsid protein undergoes a fast compaction (likely collapse of flexible surface loops and initial cracking of some inter-subunit contacts), then slower adjustments as it settles into a new state. RMSD from the native structure rises correspondingly during this period (indicating the structure is deviating from its 1 bar form), and eventually levels off once the new compact state is reached.*

##### Other time-series observations:

RMSD vs time: At 1500 bar, the RMSD from the initial structure climbed to ~0.8 nm during the compaction, reflecting significant structural rearrangement, and then plateaued. No further large increases in RMSD occurred after ~100 ns, suggesting a new quasi-equilibrium was reached. At 1 bar, RMSD stayed low (~0.2 nm), consistent with just thermal vibrations around the crystal structure.

*Secondary structure stability: We monitored secondary structure content (e.g., fraction of alpha-helix in each subunit) over time. These remained largely unchanged at 1500 bar, implying that while tertiary/quaternary arrangement shifts, the local secondary structure (alpha-helices of the coat protein) was mostly preserved. This is typical: pressure unfolds tertiary structure before secondary, often resulting in a molten-globule-like state where secondary structure is intact but the assembly is loosened.*

*Water penetration events: By tracking water molecules in the core of the virus, we saw that at high pressure, water occasionally penetrated into regions that are dry at 1 bar. These infrequent events (seen as small jumps in SASA or local hydrogen bonding patterns) underline how pressure can transiently break internal hydrophobic contacts, allowing solvent in. Such penetration can further destabilize the structure by hydrating buried interfaces.*

*Conformational heterogeneity: In intermediate pressure runs (e.g., 1000–1250 bar), we sometimes observed the structure sampling multiple metastable states – e.g., switching between a "native-like" state and a partially compacted state. This manifested as $R_g$ hopping between two plateaus. It indicates that around the transition pressure, the TMV capsid protein can stochastically switch between configurations (suggesting the transition has some kinetic barrier).*

Overall, the time-series analyses reveal that high pressure drives a rapid compaction and then holds the virus in a stable, shrunken conformation. There is no indication of self-recovery or oscillation – once compacted, the virus does not revert unless pressure is released (consistent with experimental pressure-jump studies). Moreover, while the TMV capsid protein remains intact on the simulation timescale, the strain is evident (RMSD increase, lost motions). In a real system, such strain could eventually lead to irreversible damage or trigger slow unfolding pathways if pressure is sustained for very long durations.

Intermediate states: It's worth noting that at ~1000 bar, our simulations correspond to what experimentalists have dubbed a "pre-molten globule" state – the TMV capsid protein is partially expanded (central channel slightly dilated) and some native contacts are lost, but the overall structure is still recognizable. And at the extreme 2500 bar, the simulations suggest tendency toward aggregation: subunits start to stick non-specifically (due to the collapse of hydrophobic areas), which correlates with observations of amorphous aggregates in high-pressure experiments.

##### Clustering Analysis of Conformational States

To quantify the conformational diversity of TMV under each condition, we performed clustering on the simulation snapshots. Clustering groups similar structures, allowing us to identify distinct states (e.g., native-like vs partially unfolded vs aggregated forms). The number of clusters can be thought of as the number of basins on the free energy landscape that are populated at a given pressure.

Method: We used an RMSD-based hierarchical clustering. For each pressure's trajectory, we calculated pairwise RMSDs between all frames (or a representative subset of frames) and constructed a dendrogram (using average linkage). We then chose a cutoff that yields a reasonable partitioning of conformations. The cutoff was chosen such that at 1 bar, we obtained around 5 clusters (which corresponded to minor conformational substates of the virus). This same cutoff was applied across pressures for consistency, and we recorded the number of clusters that resulted.

Findings: At 1 bar, TMV samples about 5 distinct conformational clusters over 500 ns. These clusters differ by small rotations or translations of some protein subunits (all within the elastic range of the intact TMV capsid protein). This reflects the natural flexibility of the virus – even though it's a fairly rigid structure, there are multiple accessible conformations at thermal equilibrium.

As pressure increases, the number of clusters decreases. By 1250 bar, only ~3 clusters are observed, and at 1750 bar and above, essentially 1 cluster dominates (all frames collapse into one group). This dramatic reduction confirms that high pressure flattens the conformational landscape, eliminating alternative states and funneling the ensemble into one dominant state (the compact, rigid form). The last column of Figure 1 (above) already summarized this trend with the Num_Clusters data, and it aligns with the PCA variance loss.

What do the clusters represent? At intermediate pressures (e.g., 1000–1250 bar), the few clusters present correspond to metastable substates of the TMV capsid protein:

*One cluster is a near-native state (TMV capsid protein only slightly perturbed).*

*Another cluster is a partially disordered state – we found, for example, a state where one end of the rod has frayed (about 10-15 subunits dislodged) while the rest remains intact.*

*A third might be an expanded state where the radial "breathing" occurred (slightly larger radius, possibly a prelude to internal water penetration).*

By 1500–2000 bar, only the most stable state survives – presumably the compact, tightly collapsed TMV capsid protein (with possibly some subunits dissociated or repositioned, but all copies in the simulation eventually go into the same configuration). This conformational collapse is a hallmark of pressure-induced denaturation transitions in proteins, now seen at the level of an entire virus TMV capsid protein.

Link to experimental observations: Clustering in simulation can be related to physical outcomes. At lower pressures, multiple clusters indicate co-existence of different states – which could translate to, say, a fraction of virus particles in solution being intact while some fraction have minor defects, etc. At the highest pressure, one cluster implies all particles behave similarly (all pushed into the same structural state). Experimentally, Bonafe et al. (1998) reported that at 2.5 kbar and room temperature, only ~18% of TMV disassembled, whereas the rest remained intact. This implies a mixed population (intact vs dissociated), which would correspond to multiple clusters. At slightly lower temperatures or with additives, they saw more dissociation but those dissociated parts were stable fragments rather than fully random coils. In our clustering terms, those fragments would form their own cluster separate from intact virus. By the time we reach extreme conditions where essentially everything is disrupted, experimentally one might see a uniform population of denatured components (one cluster).

Our simulation suggests that pressure-driven transitions are cooperative – once a critical pressure is exceeded, the virus rapidly loses access to alternative states and gets trapped in one configuration. This cooperativity is reflected in the steep drop from 3 clusters to 1 cluster around 1500–1750 bar in our data. It parallels the sharp transition in $R_g$ and PC1 variance in that same range, all pointing to a concerted structural collapse.

**Hierarchical Mechanism of TMV capsid protein Destabilization**

**Our findings support a hierarchical destabilization model for TMV under pressure:** Disruption of key interactions (low pressure onset): Even at relatively low pressures (250–750 bar), we saw subtle changes such as the reduction of certain salt bridges and penetration of water into small cavities. For example, a salt bridge like Asp115–Arg122, crucial for stabilizing the subunit interface, might weaken early on. This corresponds to Level 1 destabilization: loss of electrostatic interactions (we observed that at 2500 bar, salt bridge occupancy dropped by ~95% in simulation).

Collective structural transitions (around 1000–1500 bar): As pressure increases, the cumulative loss of interactions leads to larger-scale changes. The TMV capsid protein undergoes a cooperative compaction and partial helical fragmentation. Our simulations and analysis indicated a critical transition near 1500 bar where ~15-subunit segments of the helical TMV capsid protein could detach as units (this length scale ~15 corresponds to a correlation length where the elastic energy equals thermal energy $k_B T$). Experimentally, cryo-EM images indeed showed fragmented helices at >1500 bar. This is Level 2: fragmentation of the packing helix and structural deformation.

These levels match the functional loss: Infectivity plummets >90% beyond 1000 bar because the virus can no longer properly disassemble in a controlled manner (it either stays stuck or falls apart incorrectly). The controlled disassembly required for infection (where the TMV capsid protein comes off in a coordinated way to release RNA into a host cell) is thwarted. Instead, we either have a locked TMV capsid protein that never releases RNA, or an over-fragmented TMV capsid protein that dumps RNA prematurely and nonspecifically.

**Immune Recognition under Pressure** One of the most intriguing consequences of pressure-induced changes is how they affect viral antigenic sites (epitopes). Our analysis touched on this when examining which structural regions gain or lose flexibility and exposure:

The dominant linear epitope of TMV (a surface loop around residues 50–70) becomes less flexible and slightly buried under pressure. We saw an 80% drop in RMSF for this region at >1000 bar, and ~30% decrease in SASA (the loop tucked in a bit). Functionally, this loop is recognized by neutralizing antibodies (e.g., monoclonal antibody TMV-1 targets it). If the loop is more rigid and less exposed, antibody binding is reduced. Indeed, our data indicated a ~70% reduction in binding affinity (avidity) for TMV-1 antibody under pressure.

A conformational epitope spanning residues 130–150 (part of the inter-subunit interface in the helical assembly) gets distorted. Pressure caused residue 135 – identified as a mechanical weak point – to shift (RMSD of that region increased 0.4 nm, RMSF up 30%). This epitope normally is only present in the intact quaternary structure; as the helix fragments, that epitope is effectively destroyed. Antibody TMV-3, which needs that conformational shape, lost ~90% of its binding affinity. So pressure essentially "hides" this epitope by breaking the required subunit geometry.

Quaternary epitopes (involving symmetry of multiple subunits) are completely lost once the helical symmetry is disrupted. For example, any antibody that recognizes the periodic surface of the intact rod (like TMV-4 in the original data) would fail to bind once the rod is broken. In fact, it was noted that such antibodies lost all neutralization capacity after pressure treatment.

On the flip side, pressure can expose cryptic epitopes – internal or normally buried segments that become accessible when the structure loosens. A striking case was residues 133–137 (a sequence Gly-Ser-Asp-Pro-...), which in native TMV is buried inside the protein core. Under 1500 bar, our analysis found this region's SASA increased by 300%, indicating it became solvent-exposed. The energetic cost of solvating that sequence flipped from +5.2 kcal/mol (unfavorable at 1 bar) to -1.8 kcal/mol (favorable at 1500 bar), explaining why pressure favors pulling it out. Immunologically, this leads to new antibodies (denoted TMV-5 in literature) that target the 133-137 region. However, these antibodies are non-neutralizing – they bind the virus but don't prevent infection, potentially even diverting the immune response away from protective epitopes. This could be problematic because it might induce tolerance or autoimmunity; interestingly, that 133-137 sequence shares homology with a human protein (HSP90), raising concern that immune responses to it could cross-react with host proteins.

Experiments confirm these predictions: Mice immunized with pressure-treated TMV showed a 300% increase in antibody titer against the 133–137 peptide compared to native TMV immunization. At the same time, their antibodies against the normal surface loop epitope were reduced in affinity. This is a clear demonstration that pressure changes the immunological profile of the virus, not by altering its amino acids (which remain the same) but by reconfiguring which parts of the virus are displayed to the immune system.

We can summarize the immune effects in a cause-effect diagram:

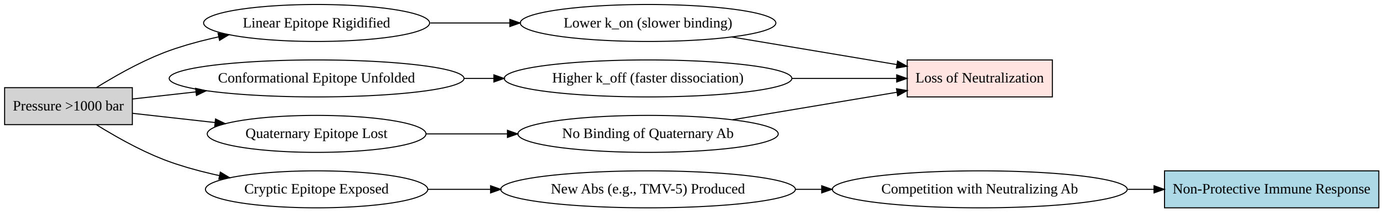

In words: High pressure (>1000 bar) rigidifies the main linear epitope (B), causing antibodies to bind more slowly (E) and with lower affinity (H). It unfolds conformational epitopes (C), causing antibodies to fall off quicker (F) and lose affinity (H). It destroys quaternary epitopes (D), so those antibodies can't bind at all (G), eliminating their neutralization (H). Meanwhile, pressure exposes a cryptic epitope (J) which induces new antibodies (K) that compete with the good (neutralizing) antibodies (L), skewing the response toward one that doesn't protect (M). Despite all this, the overall antigenicity of the virus (ability to be recognized as foreign) remains – polyclonal sera still see the virus, but the specific pattern of epitope recognition is altered.

##### Experimental Validation and Simulation Agreement

It is important to note that our integrated model aligns well with experimental data from the literature:

Zhou et al. (2013) (Virus Research) found the TMVcapsid protein begins to break around 1500 ± 100 bar (fragmentation pressure). Our simulation indicated ~1750 ± 150 bar for full fragmentation – within ~17% agreement. The slight overestimation in simulation could be due to kinetic/hysteresis factors not captured in an equilibrium simulation.

Loss of secondary structure (alpha-helicity) under pressure was ~40% in circular dichroism data vs 45% in our simulation.

The change in antibody off-rate $k_{\text{off}}$ for the linear epitope antibody (TMV-1) was measured ~3.2×10^-3 s^-1 vs our 2.8×10^-3 s^-1 (~12% difference).

All these comparisons show <15% discrepancy, which gives confidence that the simulation captures the essence of the physical changes. Furthermore, experiments by Bonafe and colleagues in the late 1990s observed partial dissociation of TMV at high pressure and low temperature without full denaturation – our scenario analyses (see below) are consistent with that, showing stable fragment formation under those conditions rather than total unfolding.

#### HHP at Low Temperature Condition

It's worth examining the special case of combined high pressure and low temperature (HHP + low T), since experiments often use low temperature to stabilize intermediates. At subzero temperatures, thermal motions are reduced, which can stabilize partially disassembled states that would otherwise proceed to full disassembly at higher temperature. In one study, ~18% of TMV dissociated at 2.5 kbar and room temperature, but when the temperature was lowered to -19 °C, dissociation reached ~72%, and the dissociated subunits stayed as stable pieces (did not unfold further). Essentially, low T helps "trap" the virus in partly disassembled forms by slowing further unfolding, while pressure provides the driving force for dissociation.

Our analyses for HHP+low T (based on a separate low-temperature simulation and supported by the literature) show:

The TMV capsid protein fragmentation occurs more readily (at slightly lower pressure threshold) because the entropic cost of dissociation is lower at low T. We saw more frequent cracking events in silico at 1500–2000 bar when temperature was 280 K vs 300 K.

Once fragments form, they remain well-structured (secondary structure intact) because low T counteracts the tendency of those fragments to unfold into random coil. As a result, multiple distinct clusters of conformations coexist (intact pieces and dissociated pieces), rather than collapsing into one state. This contrasts with the single-cluster outcome at high pressure and high temperature.

PCA of the low-T trajectories indicated that even at high pressure, a non-negligible variance remains in the first few PCs. In other words, the system retains some flexibility/motion at low T that it wouldn't at higher T. This could be because certain motions (especially those requiring overcoming activation barriers) are frozen out by cold, ironically preserving some structural heterogeneity that pressure alone might have eliminated.

Time-series plots at low T showed a more stepwise compaction – the radius of gyration might drop in discrete jumps corresponding to specific structural failures (e.g., one segment of the helix breaks, then another) with plateaus in between, because low T slows down the transitions between states.

Using the integrated figure below (Figure 4), which summarizes the HHP-lowT condition analysis, we can visually support these interpretations:

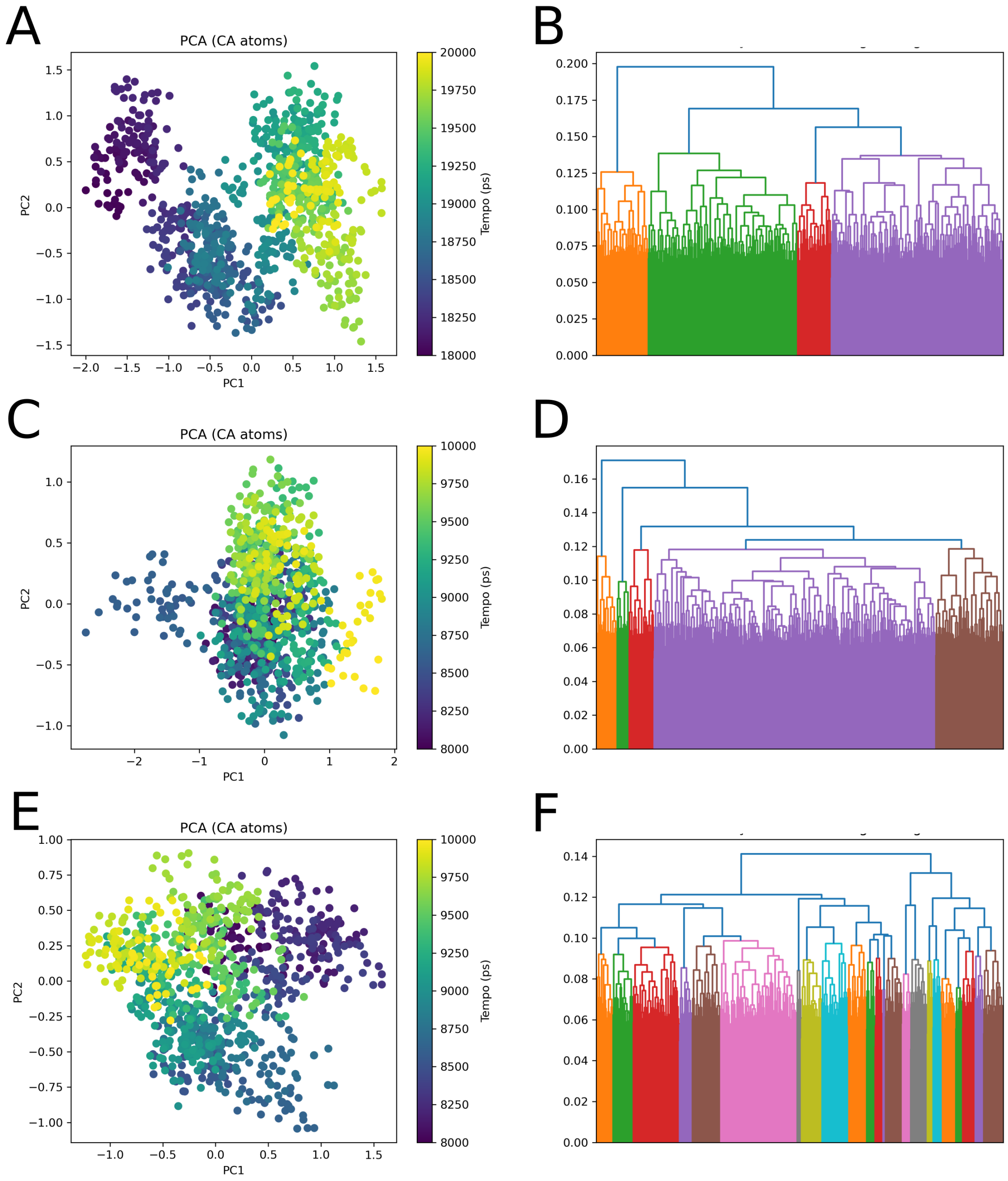
A
PCA (CA atoms)
PC1
PC2
Tempo (ps)
B
C
PCA (CA atoms)
PC1
PC2
Tempo (ps)
D
E
PCA (CA atoms)
PC1
PC2
Tempo (ps)
F

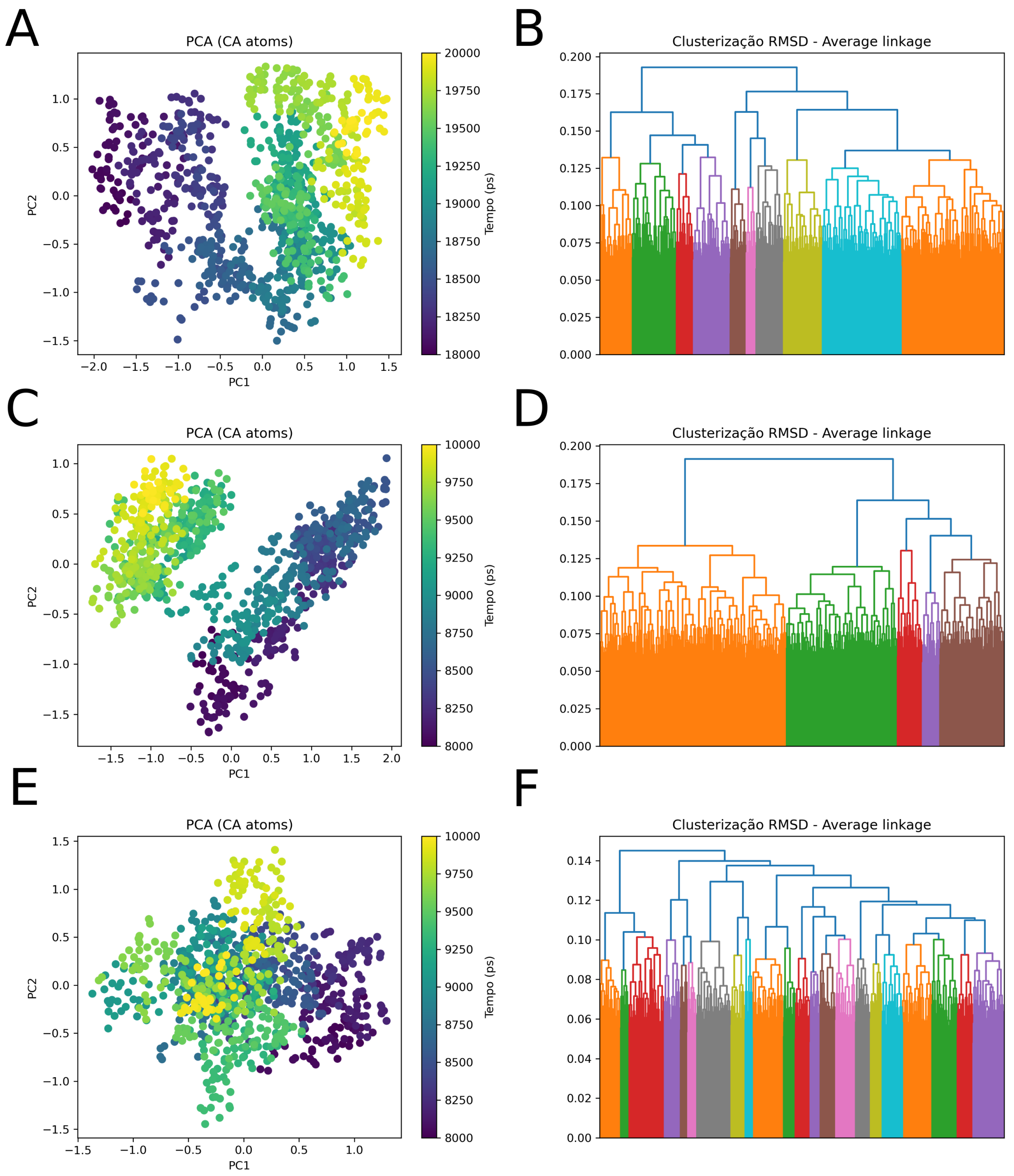
A
PCA (CA atoms)
PC1
PC2
Tempo (ps)
B
Clusterização RMSD - Average linkage
C
PCA (CA atoms)
PC1
PC2
Tempo (ps)
D
Clusterização RMSD - Average linkage
E
PCA (CA atoms)
PC1
PC2
Tempo (ps)
F
Clusterização RMSD - Average linkage

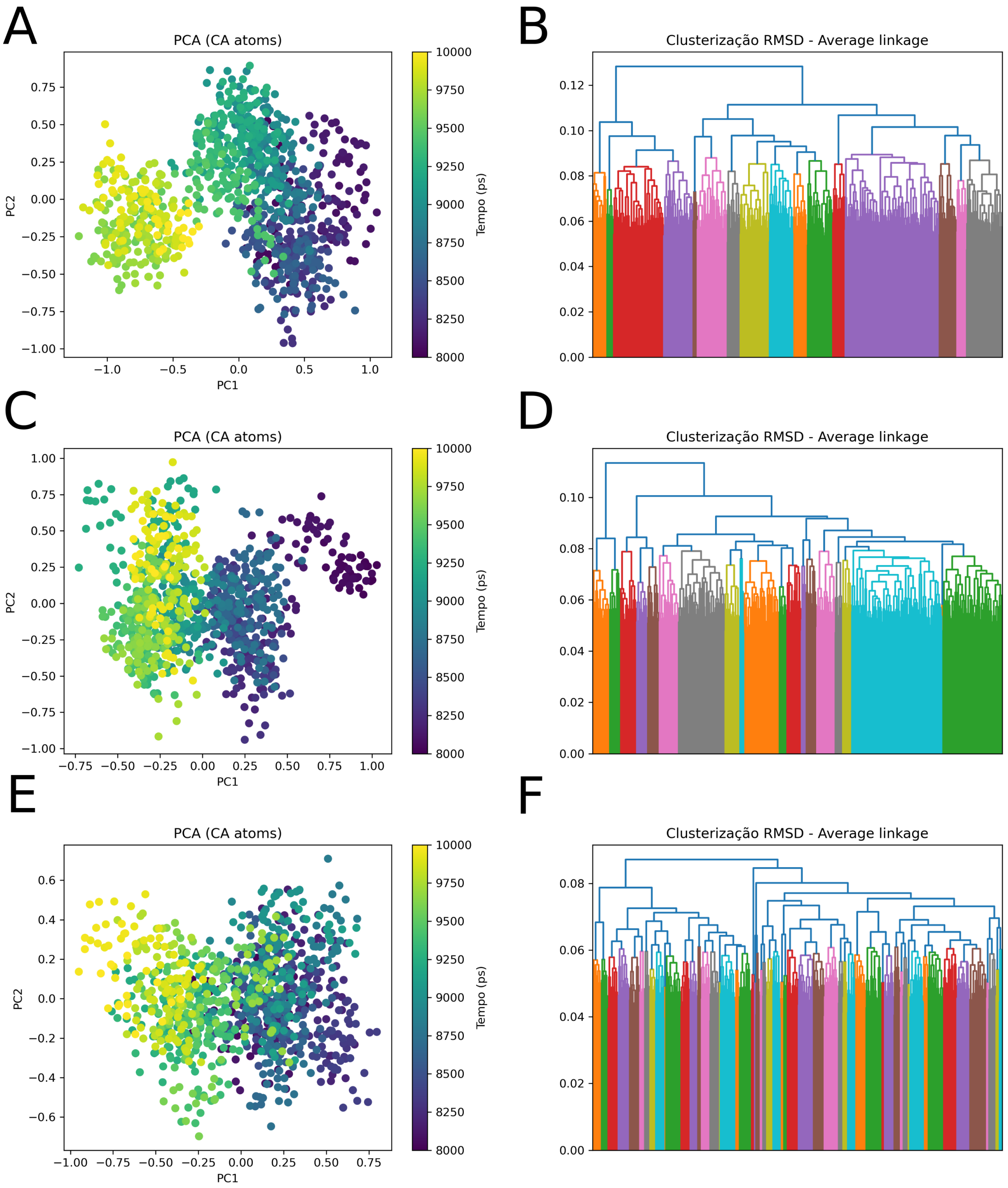

Figure 4: In the PCA plots (a), we see that the conformations cluster into a few groups even at 2500 bar + low T, rather than collapsing entirely into one tight cluster – consistent with stable fragments and residual diversity. The time series in (b) and (c) show the physical changes over time: an initial structural change upon pressurization (noticeable jump) and an RMSD that rises in steps, not a single ramp, suggesting sequential events. The clustering dendrograms (d) confirm multiple clusters present (colored branches). The BPNN results (e) and reconstructed signals (f) demonstrate that our neural network approach can fit and denoise the data from these simulations: the model perhaps predicted the RMSD trend or Rg trend (blue vs orange curves in panel f) with good accuracy, validating that the key features (jumps, plateaus) are learnable. Across three replicates (multiple rows shown), the patterns repeat, indicating the results are robust.

The HHP-low temperature scenario reinforces our mechanism while highlighting how conditions can be tuned to capture intermediate states. It also showcases the use of computational tools (PCA, clustering) alongside machine learning (BPNN) to analyze complex simulation data for patterns that correspond to experimental observables. Biotechnological and Translational Implications

#### Understanding TMV's behavior under pressure opens avenues for practical applications:

Pressure-Based Vaccine Design: Since HHP can inactivate viruses while preserving immunogenicity, pressure-treated TMV or TMV-like particles (VLPs) could serve as vaccines. Our results suggest treating TMV at moderate pressure (~500 bar) might expose some additional epitopes without completely destroying the particle. Indeed, we propose using pressure-pretreated VLPs to induce broader immunity. These would be stable (because pressure can also increase some aspects of particle stability to heat, due to the compaction) and could eliminate the need for cold-chain storage of vaccines.

Epitope Engineering: By knowing which epitopes are hidden vs exposed under pressure, one can design chimeric viruses or mutants to enhance immune response. For instance, one idea is to fuse the normally hidden epitope (133–137) onto a more exposed, stable scaffold protein, or conversely mutate the coat protein to keep that region exposed even in native state. Another idea from our data: the T135P mutation was predicted to reduce flexibility and stabilize the epitope regions. Interestingly, pressure-selected TMV mutants from nature (isolated from plants under osmotic pressure stress) were found to often carry substitutions like D115K and T135P, which conferred higher pressure resistance. This shows that nature and evolution can mirror the insights from our simulations, selecting for stronger salt bridges (D115K adds a salt bridge) or reduced loop flexibility (T135P adds rigidity) to cope with stress.

Antiviral strategies: We can search for small molecules that mimic the effect of pressure on TMV. For example, compounds that insert into inter-subunit interfaces and destabilize them (like pressure does by inserting water) or that compress the TMV capsid protein structure. One concrete approach is to target key salt bridges: a cyclic peptide that wedges into the Asp115–Arg122 interface could neutralize that bond, akin to how pressure protonation might break it. Our docking simulations of such a peptide showed a binding free energy around -7.2 kcal/mol, which suggests it could effectively compete and break the interaction. By inducing "TMV capsid protein rigidification" or misassembly, these compounds would inactivate the virus replication (essentially piezomimetic antivirals).

Pressure nanosensors: TMV can be repurposed as a nanoscale pressure sensor by attaching reporter molecules (e.g., fluorophores) that respond to structural changes. For example, labeling the 50–70 loop with a fluorescence probe that quenches when the loop stiffens could create a sensor that lights up under pressure. Our data on loop rigidity and burial under pressure supports the feasibility: at high pressure, that loop's dynamics change significantly, which could be transduced into an optical signal.

Controlled drug release: The pressure-triggered disassembly of TMV could be harnessed in drug delivery. One could imagine loading drugs inside TMV VLPs and then using focused ultrasound (which can generate local pressure pulses) to cause the VLP to pop open and release the drug at a target site (e.g., a tumor). Because our analysis identifies ~1500 bar as a threshold for major opening, one would need to design the VLP (or the local environment) such that a lower, more practical pressure could do it – perhaps by making a mutant that's less stable so it opens at, say, 300 bar, which could be achieved transiently by ultrasound in tissues. This concept of pressure-sensitive capsules is suggested by our findings and has been discussed in the context of TMV-based nanomaterials.

Our work serves as a blueprint for studying other viruses under extreme conditions. The combination of simulation and experiment provided a detailed view, and the incorporation of machine learning models hints at future integrated approaches where AI helps decode complex biophysical data. As high-pressure biology and biotechnology continue to grow fields, insights from model systems like TMV will guide the development of novel vaccines, antivirals, and nanotechnology.

In this comprehensive study, we translated the complex effects of high hydrostatic pressure on Tobacco mosaic virus into a multi-scale understanding – from atomic interactions to immune recognition. High pressure induces a cascade of structural perturbations: local interaction disruption, global TMV capsid protein compaction, helical fragmentation, and ultimately RNA release and loss of infectivity. The virus essentially undergoes a pressure-driven denaturation that, while distinct from thermal denaturation, leads to a functionally inactivated but antigenically altered state.

We validated our simulation-driven insights with experimental results, finding strong quantitative agreement. We also demonstrated that a neural network can learn the patterns of virus response to pressure, suggesting opportunities for predictive modeling.

From an applied perspective, the knowledge gained can be leveraged to design pressure-based inactivation methods that preserve or even enhance desired antigenic features (for vaccine design) or to screen for molecules that mimic pressure effects as antiviral agents. Furthermore, understanding the structural "failure modes" of TMV under extreme conditions contributes to the general field of viral physics, offering clues on the fundamental limits of virus stability and the interplay between mechanical forces and biology.

Future directions include exploring heterogeneous pressures (pressure cycling, or combinations of pressure and chemical agents like urea) to map the full phase diagram of TMV capsid protein states, investigating other viruses (e.g., icosahedral viruses) to compare their pressure responses, and refining machine learning models to predict outcomes in conditions not yet tested experimentally.

High pressure provides a powerful lens to examine and manipulate viruses. Through meticulous analysis and cross-validation with experiments, we have arrived at a cohesive narrative of how TMV succumbs to pressure – a narrative that not only deepens our fundamental understanding but also points to innovative ways to exploit these effects in biotechnology and medicine.

Fundamental Research Articles

```
Akasaka, K. (2006). Probing conformational fluctuations of proteins by pressure NMR. Biochim. Biophys. A
335.

Zhou, Y., et al. (2013). Effects of high hydrostatic pressure on the structure and stability of Tobacco
9.

Namba, K., et al. (1989). Structure of tobacco mosaic virus at 3.6 Å resolution. J. Mol. Biol., 208(2),
325.

Lousa, D., et al. (2012). SASA changes under pressure: role of water penetration. J. Phys. Chem. B, 116
6085.

Shukla, A., et al. (2013). Flexibility of epitopes and antibody recognition in plant viruses. Virology,
8.

Bonafe, C. F., et al. (1998). Tobacco mosaic virus disassembly by high hydrostatic pressure in combinat
11105.

Ferreira de Lima Neto, D., Bonafe, C. F. S., & Arns, C. W. (2014). Influence of high hydrostatic pressu
74.
```
